# Systematic Targeting of Protein Complexes with Molecular COUPLrs

**DOI:** 10.1101/2024.07.16.603666

**Authors:** Diane Yang, Stefan Andrew Harry, Harrison Byron Chong, Edwin Zhang, Natalie Shannon Nordenfelt, Nicholas Chen, Christine Lee, Stefan Kaluziak, Elizabeth Codd, Samay Trivedi, Magdy Gohar, Giovan McKnight, Dawn R. Mitchell, Maolin Ge, Chengzhuo Gao, Zavontae Holmes, Wenxin Yang, Abigail Elizabeth Smith, Alexander Daniel Carlin, Matthew J. Lazarov, Neha Khandelwal, Mariko Hara, Siwen Zhang, Herman Xin Yang Leong, Hector Martinez Luna, Zander Chearavanont, Kim Emonds, George Popoola, Idris Barakat, Maristela Onozato, Mohammed Mahamdeh, Toshio Fujino, Hyuk-Soo Seo, Sirano Dhe-Paganon, Zhen-Yu Jim Sun, Gregory J Heffron, Aaron Hata, Roy Jason Soberman, Brian B. Liau, A. John Iafrate, Liron Bar-Peled

## Abstract

Molecular glues that engage protein complexes have transformed the study of cell biology and have had a direct impact on clinical oncology. However, the identification of new glue classes and their corresponding protein complexes has remained largely serendipitous. To overcome this challenge, we report the development of molecular COUPLrs, elaborated small molecules flanked by two cysteine-reactive warheads, as well as CONNECT, an integrated chemical proteomic platform for target deconvolution. By profiling a library of molecular COUPLrs across 13 cancer cell lines, we uncovered hundreds of proteins that can be coupled together, including in some cases in mutant selective fashions. We develop an advanced COUPLr for the oncogene EML4-ALK, which engages the fusion outside of its kinase domain, restricts protein dynamics, and disrupts EML4-ALK signaling. Collectively, molecular COUPLrs substantially expand the scope of proteins that can be chemically connected, providing an unbiased approach to identify small molecules that target protein complexes.

## Introduction

Small molecule glues have transformed the study of biology. Molecular glues have also formed the basis for numerous FDA-approved medicines including rapamycin^1^ (immunosuppressant), thalidomide^2^ (multiple myeloma) and cyclosporin^3^ (organ rejection post-transplant). Despite their biological and clinical importance, the vast majority of molecular glue classes have been identified serendipitously^4–8^. While multiple derivatives of established glues continue to expand the number of E3 ubiquitin ligases and zinc fingers that can be engaged, the absolute scope of protein classes that undergo gluing remains unclear.

Protein crosslinkers have played an important role in the in vitro characterization of protein complexes^9–12^. Relying on two covalent warheads to bridge transient protein interactions, crosslinkers enable the biochemical dissection of multi-protein machines^13^. While the biochemical utility of crosslinkers is clear, their applicability to define new protein interactions in cells or to bind specific protein classes/complexes is highly limited^14^.

The privileged chemistry of cysteine has not only endowed this amino acid with a central role in protein function but over the last two decades has designated this residue as the preferred small molecule handle for covalent drug discovery^15–21^. With over 260,000 cysteines encoded in the human genome and many identified in disease-causing proteins, numerous chemical probes and medicines have been developed to target cysteines^22–36^. Taking advantage of the unique attributes of covalency has not only expanded the druggable genome but enabled the targeting of traditionally challenging protein classes including transcription factors and scaffolding proteins^37–54^. The concurrent development of cysteine-focused chemical proteomic technologies has now allowed the global dissection of cysteine-ligand interactions, providing an atlas of what is actionable across cancer^55–60^.

To enable the systematic identification of protein complexes that could be targeted by molecular glues, we combined design principles from glues, crosslinkers and covalency to develop molecular *COUPLrs* (COvalent Protein Ligators). These glue-like molecules are characterized by elaborated binding elements flanked by two cysteine (Cys-Cys, C^2^) reactive warheads that couple proteins together. Integrating a library of 33 C^2^-COUPLrs with CONNECT, a chemical proteomic discovery engine, enables the immediate characterization of proteins amenable to coupling. CONNECT combines cysteine focused chemical proteomics^61^ to identify the precise cysteines engaged by COUPLrs with mobility shift proteomics^62^ to define which proteins are coupled together. By first mapping the targets of our C^2^-COUPLr library in one cell line and then broadly profiling an *informer set* of C^2^-COUPLrs across 12 cell lines, we provide a systematic characterization of coupled proteins. This series of experiments identified 1000+ proteins amenable to coupling, including many classes of proteins not traditionally targeted by molecular glues including scaffolding proteins, transcriptional machinery and signaling molecules. We identify both common targets of COUPLrs, as well as distinct ones based on their restricted protein expression, mutational status, or the inherent chemical structure of the COUPLr employed. We proceed to identify a C^2^-COUPLr for EML4-ALK and through medicinal chemistry optimization arrive at an advanced probe which restricts EML4-ALK protein dynamics and disrupts the signaling of both wild-type and a kinase-domain resistance mutant. Collectively, we describe the development of a drug-discovery modality that systematically reveals which protein complexes can be permanently glued, substantially expanding the scope of proteins amenable to small molecule connectivity.

## Results

### Chemical proteomic characterization of a molecular COUPLr library using CONNECT

To date, many of the distinct classes of small molecule glues have originated from natural products^4, 63–70^. A systematic analysis of 400,000+ natural products revealed that greater than 40% contain a putative electrophilic warhead (e.g. α,β-unsaturated carbonyl, epoxide, etc.) (Extended Data Fig. 1a-b)^71–74^. Interestingly, ∼50,000+ (∼12%) natural products contain two or more electrophilic warheads that are potentially cysteine reactive, including cyanocycline D^75^, leucanthin B^76^, montadial A^77^, and manumycin polyketides^65^ (Extended Data Fig. 1b-c). Some of these natural products with multiple warheads have demonstrated covalent linkage of two proteins^78–80^. Building on these design principles, we sought to develop a library of synthetically tractable dual warhead biomimetics that we term molecular *COUPLrs*, given the permanent nature of their interactions with protein complexes. Accordingly, we constructed a 33-member library of C^2^-COUPLrs composed of structurally diverse binding elements flanked by two cysteine reactive warheads (e.g., chloroacetamides or acrylamides) (Fig. 1a, Extended Data Fig. 1d). In contrast to crosslinkers, which use simple linear chains to separate covalent warheads (Fig. 1a), COUPLrs feature more complex binding elements, such as polycyclic motifs, heterocycles, and a variety of non-covalent functional groups, which are flanked by electrophiles (Extended Data Fig. 1d-f)^81, 82^.

**Fig. 1:**
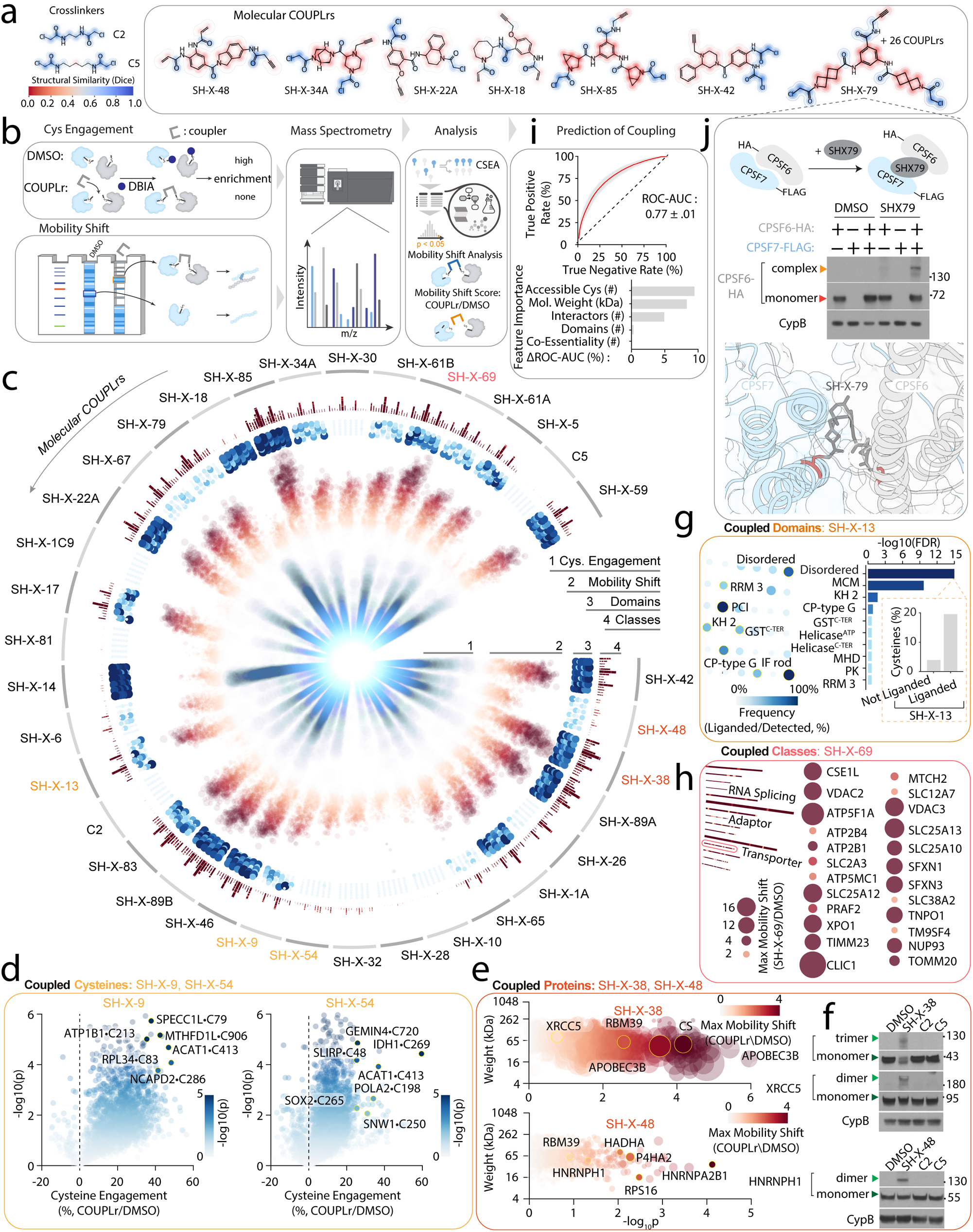
Chemical proteomic analysis of molecular COUPLrs reveals diverse protein targets. (a) Development of molecular COUPLrs. Representative members of C^2^-COUPLR library defined by two electrophilic warheads (see also Extended Data Fig. 1d) and color coded by their chemical fingerprints with the crosslinker C2 serving as a reference point (see Methods). (b) Schematic depicting CONNECT, a dual chemical proteomic platform for identifying the cysteine and protein targets of molecular COUPLrs. (c) Circular heatmap depicting differences among cysteine, protein, protein domain, and protein class coupling (moving from inner- to outermost layers) across 33 C^2^-COUPLrs profiled in U2OS cells (see also Extended Data Tables S1, Methods). Cysteine engagement is depicted in blue, and protein mobility shifting is depicted in red. (d) Examples of differences in cysteine coupling following treatment with SH-X-9 and SH-X-54. (e-f) Identification of coupled proteins. Scatter plots depict mobility shift ratios of different proteins following treatment with SH-X-38 and SH-X-48 (e). Targets of C^2^-COUPLrs. Lysate isolated from U2OS cells was treated with the indicated compounds (100 µM SH-X-38, SH-X-48, or cross-linkers for 1 hr) and coupling for HNRNPH1, XRCC5, and APOBEC3B was determined by immunoblot (f). (g) Protein domains coupled by SH-X-13. Inset, percentage of cysteines within disordered domains that are engaged by SH-X-13 (see also Methods). (h) Protein classes coupled by SH-X-69. (i) Machine learning enables prediction of proteins amenable to coupling with 77% ROC-AUC, (top) and features driving prediction (bottom) (see Methods). (j) COUPLrs target protein complexes. Top, representative immunoblot analysis of CPSF6-CPSF7 interaction following treatment with SH-X-79. HEK-293T cells transiently expressing the indicated FLAG- or HA-tagged proteins were lysed, and protein coupling was determined by immunoblot following treatment in lysate with 100 µM of SH-X-79. Bottom, structure of CPSF6-CPSF7 modeled^175^ with SH-X-79 (see also Extended Data Fig. 3k, methods).

To identify the targets of COUPLrs we employed a chemical proteomics platform termed CONNECT (<u>COuplr NEtwork deteCTor</u>), which integrates cysteine focused chemical proteomics with mobility shift proteomics (Fig. 1b)^83^. Cysteine chemical proteomics identifies the precise cysteine residues modified by electrophilic compounds^84–89^. With mobility shift proteomics, C^2^- COUPLr treated samples are denatured and run on a SDS-PAGE, which is then segmented into individual bands of defined molecular weights and analyzed by mass spectrometry (Fig. 1b). Coupled proteins are defined as those proteins that shift to a higher molecular weight band representing covalent linking to an additional protein(s) (Extended Data Fig. 2a-e). Using this platform, we systematically characterized the targets of the C^2^-COUPLr library in U2OS cell lysate, calculating a mobility shift and engagement score for each of the 5000+ proteins identified, finding a correlation between cysteines engaged and proteins coupled, in cells and in lysate (Fig. 1c-e, Extended Data Fig. 2a-g, Table S1). We consider a cysteine targeted by a COUPLr if it demonstrates 25% engagement with p < 0.05. A mobility shift is defined for individual proteins with ≥1.5-fold (p < 0.05) increase in abundance at an observed molecular weight which is greater than its expected molecular weight (see methods)^90^. We found that COUPLrs had varying reactivities, binding to both common as well as distinct targets (Fig. 1c, Extended Data Fig. 4a). Additionally, we identified certain proteins that were targeted by multiple C^2^-COUPLrs, including PRDX1 and EEF1A1, suggesting these proteins are more amenable to this form of chemical connectivity (Extended Data Fig. 4b). APOBEC3B, XRCC5 and HNRNPH1 are exemplary COUPLr targets (Fig. 1e-f). These proteins were not engaged by chloroacetamide containing cross-linkers (Fig. 1a, e-f). Many domain architectures, such as the ADF-H, ATP-grasp, disordered domains, and ABC transporter 2 domains, were targeted by several COUPLrs (Fig. 1h, Extended Data Fig. 3a). In aggregate, we identified 712 proteins which were coupled at least once, representative of 118 protein classes. Classes encompassing proteins such as transporters, translational machinery and transcription factors are frequently targeted (Fig. 1g, Extended Data Fig. 3b-e). Multiple pathways were targeted by COUPLrs, where the RAS signaling pathway exemplifies how differences in pathway member coupling can be measured and mapped to differences in underlying cysteine engagement (Extended Data Fig. 3f-h). In contrast to the target specificity of COUPLrs, we found that crosslinkers coupled very few proteins (Fig. 1c, Extended Data Table S1). These findings accord well with previous studies demonstrating that covalent small molecule protein interactions are favored by defined binding elements^59, 90, 91^. By leveraging an established protein-protein interaction database (see methods), we developed a network of those interactions that could potentially be coupled as measured by the CONNECT platform (Extended Data Fig. 3i). The RNA processing complex CPSF6-CPSF7 is coupled by SH-X-79 (Fig. 1j, Extended Data Fig. 3j-k) offering an example of how C^2^-COUPLrs stabilize hetero-protein complexes.

To determine if the target scope of COUPLrs was fundamentally different than compounds with monovalent warheads, we first compared cysteine reactivity changes between COUPLrs and a library of 152 previously profiled mono warhead ligands^83^, finding reasonable correlation between the two types of probes (Extended Data Fig. 4g). To further quantify differences between monovalent and dual warhead compounds, we constructed four pairs of dual– and mono-warhead ligands, where the second warhead is replaced with an acetamide group (Extended Data Fig. 4h). Comparison of cysteine reactivity changes between the pairs revealed good correlation between the target scope of each pair (Extended Data Fig. 4h), indicating that the addition of a second warhead does not fundamentally change protein reactivity.

The incorporation of stereospecificity into chemical probes takes advantage of the chiral nature of many protein binding pockets^43, 59, 92–101^. With this in mind, we designed two pairs of stereoisomer COUPLrs (SH-X-61A/B, and SH-X-89A/B), finding several oncogenes which are differentially engaged by SH-X-61A/B (Fig. 1c, Extended Data Fig. 4i). DNMT3A provides an example of stereo-selective coupling as it was targeted by SH-X-61B but not SH-X-61A (Extended Data Fig. 4i). Mapping all cysteines identified in this study to their corresponding structures and subsequent cysteine set enrichment analysis (CSEA)^83^ revealed that cysteines which are superficially exposed are targetable by COUPLrs (Extended Data Fig. 4f). Furthermore, we found that COUPLrs bind cysteines residing in protein pockets and protein-protein interfaces (Extended Data Fig. 4c,e). This observation is corroborated by mobility shift data which demonstrate that COUPLrs link proteins that are predicted to form homo-oligomers^102^ (Extended Data Fig. 4d). In search of a logic underlying the target scope of COUPLrs, we built a classifier aimed at predicting whether a protein is amenable to coupling by at least one of our molecular COUPLrs (Fig. 1i). This model achieves reasonable performance (77% ROC-AUC; 70% accuracy; 73% precision; 69% recall; 71% specificity; 0.71 F1 score) based on a streamlined set of features describing a single protein. Broadly, these define the known interactivity of a protein, the number and kind of its potential interacting interfaces, and the number of known reactive cysteines that it encodes (Fig. 1i). Encouragingly, the curation of these features did not require the generation of any new data, raising the possibility of prospectively assessing a protein’s amenability to molecular coupling.

Using density functional theory (DFT), we calculated (B3LYP/6-31+G(d,p)) the distances between the dual warheads (Extended Data Fig. 1e). Overlaying the distances between cysteines in proteins and their respective engagement scores, we identified hundreds of cysteine pairs that are possibly intra-molecularly coupled (Extended Data Fig. 4j). One prominent example of intra-protein coupling was NRAS by SH-X-14, where the C^2^-COUPLr possibly bridges Cys80 to Cys118 (Extended Data Fig. 4j)^103–112^. In support of this hypothesis, we find that treatment of the double mutant NRAS•C80A/C118A with SH-X-14 does not increase thermal stability compared to WT NRAS (Extended Data Fig. 4k-m). Single NRAS mutants (NRAS•C80A or NRAS•C118A) had an intermediate level of stabilization following SH-X-14 treatment, suggesting that distinct conformational changes occur upon intra-protein coupling in comparison to single site liganding events (Extended Data Fig. 4k-m). These results highlight the ability of COUPLrs to not only stabilize heterocomplex protein-protein interactions but also intra-protein connections^113^.

Collectively, by developing a library of diverse, doubly covalent molecular COUPLrs we are enabled to employ a chemical proteomic platform tailored for their analysis. We find that COUPLrs substantially expand the range of protein classes that undergo chemical connectivity, stabilizing proteins and the complexes they form.

### Lineage and mutation specific molecular coupling

To enable a broader dissection of C^2^-COUPLr targets we selected four probes of distinct molecular fingerprints (similarity average = 0.23) (Fig. 1a, Extended Data Fig. 1d-f) and reactivities (Fig. 1c) and profiled their targets in 12 additional cancer cell lines representative of 10 lineages using CONNECT. We identified 9606 cysteines and 3110 proteins that were coupled at least once across four COUPLrs and 12 cell lines (Fig. 2a, Extended Data Fig. 5a). Interestingly, of the 66 established cancer drivers detected in these models, we found that 36 were coupled at least once (Extended Data Fig. 5b). Clustering of coupled proteins in each cell line revealed examples of heterogeneous targeting among the different models profiled (Fig. 2a). For example, the ribosomal protein SA RPSA, whose expression was uniform throughout the models examined, was highly coupled in the UACC257 cell line but not others (Fig. 2a), which may be reflective differential complex formation or unique metabolic environments^90^.

**Fig. 2:**
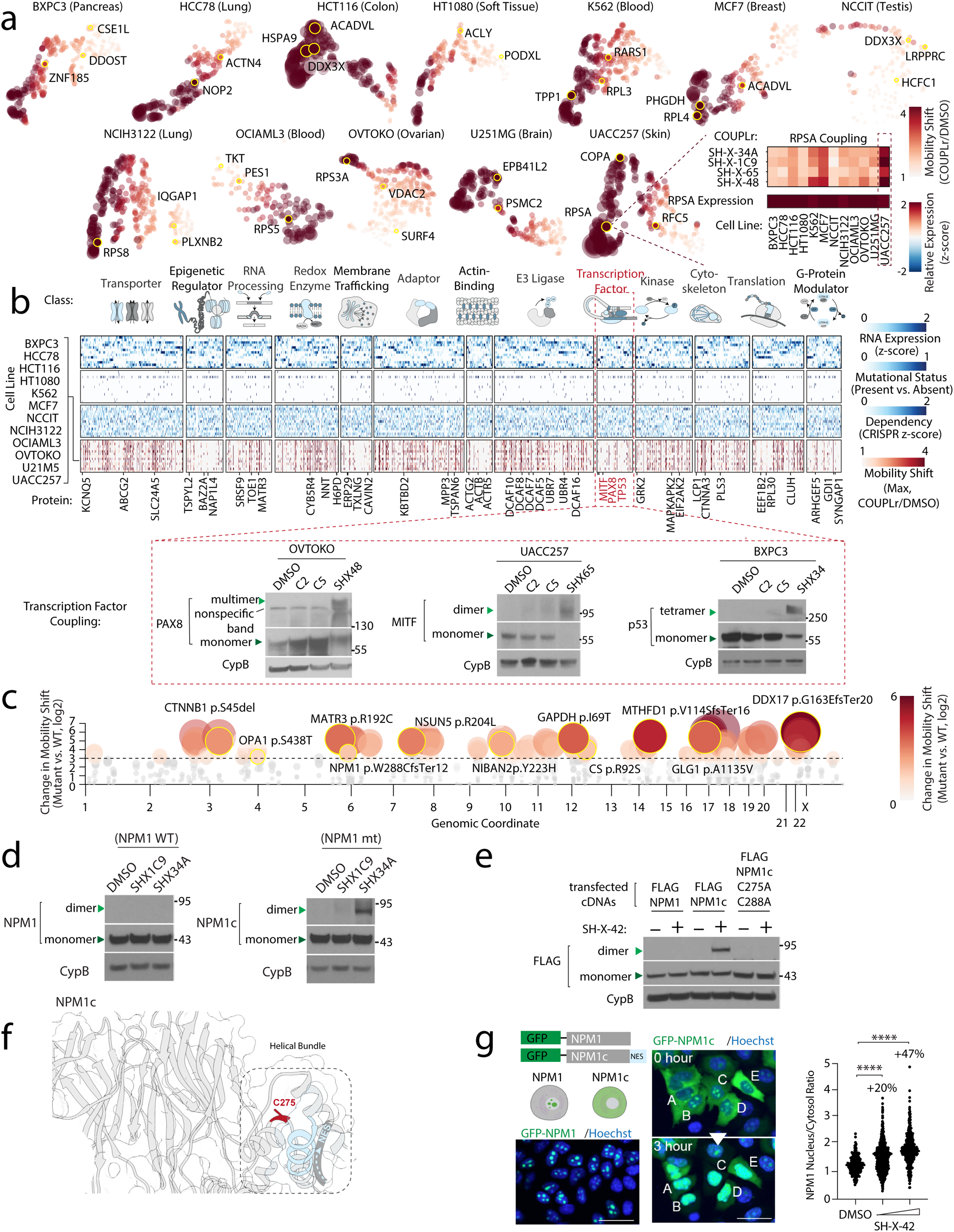
Defining Molecular COUPLr targets across cancer. (a) UMAP representation of coupled proteins (colored dots) in 12 cancer cell lines following treatment with SH-X-1C9. Heatmap inset depicts cell and COUPLr specific engagement for RPSA (top) and expression (bottom). (b) Multi-omic clustering of mRNA, gene essentiality, mutational status, and molecular coupling ordered by protein class across profiled cancer cell lines. Insets, PAX8 coupling by SH-X-48 in the OVTOKO ovarian cancer cell line, MITF coupling by SH-X-65 in UACC257 melanoma cell line and TP53 coupling by SH-X-34 in the BXPC3 pancreatic cancer cell line as determined by immunoblot. (c) Coupling plot identifying gene mutations associated with changes in coupling in cis and trans (see Methods). (d) Mutant NPM1 (NPM1c) is selectively coupled. Representative immunoblot analysis of WT NPM1 and NPM1c coupling in HT1080 and OCI-AML3 cell lines. (e) NPM1c coupling is dependent on Cys275 and Cys288. U2OS cells expressing the indicated NPM1c variants were treated with vehicle or 20 µM SH-X-42 for 3 hrs, and coupling was assessed as described in (b). (f) Model of NPM1c (see methods) highlighting location of Cys275 and mutant NES. (g) SH-X-42 coupling of NPM1c increases nuclear localization. Left, representative immunofluorescent images of NPM1-GFP or NPM1c-GFP following vehicle or SH-X-42 treatment in U2OS cells. Right, quantification of nucleus/cytoplasm ratio of FLAG-NPM1c following treatment with 12.5 and 25 µM SH-X-42 for 3 hrs. Scale bar = 50 µm. Data are represented as mean ± SD. ***p< 0.0001. Statistical significance determined by Student’s t-test (two-tailed, unpaired).

Leveraging the deep genomic and functional genomic characterization of each cell line profiled, we asked how coupling was correlated to expression, mutation or essentiality (Fig. 2b). Lineage restricted coupling was prominent for the transcription factor PAX8 in the OVTOKO ovarian cancer cell line, MITF in U257 melanoma cells and TP53•Y220C in BXPC3 pancreatic cancer cells (Fig. 2b). Aside from coupling of lineage restricted transcription factors, there was modest systematic correlation between expression/essentiality and mobility shifting; however, those targets exhibiting high correlation between essentiality and coupling may be reflective of established protein complexes^114^ (Extended Data Fig. 5c-d). Given that proximal mutations alter monovalent cysteine adduction^90^, we asked whether this might be the case for coupling. Systematic comparison of coupling between cell lines harboring mutant and wild-type copies of a protein revealed many examples of differential coupling which potentially associate with amino acid mutation (Fig. 2c). Coupling of mutant NPM1 (NPM1c) in OCI-AML3 drew our attention (Fig. 2c-d). NPM1c is mutated in 30% of acute myelogenous leukemia (AML)^115–118^ where a c-terminal frameshift mutation introduces a nuclear export signal (NES). The resulting translocation from the nucleolus to the cytosol is required for AML proliferation^115, 119–121^. Excitingly, we found that mutant NPM1 was coupled more than WT NPM1 by SH-X-34A (Fig. 2d). Profiling additional C^2^-COUPLrs led to the identification that SH-X-42 was a more potent engager of NPM1c. Initially, we hypothesized that engagement of the mutant Cys288 within the NES was driving coupling. However, mutation of each cysteine within NPM1c, revealed Cys275 was the main driver of coupling (Extended Data Fig. 5e). Mutation of both Cys275 and Cys288 to alanine completely abolished coupling (Fig. 2e, Extended Data Fig. 5f). This provides a striking example of how a proximal mutation alters ligandability^57^: the presence of the added NES domain increases the coupling of the proximal Cys275 (Fig. 2f). Expressing FLAG-tagged NPM1c in U2OS cells, revealed that treatment with SH-X-42 resulted in nuclear localization of NPM1c in a dose-dependent manner (Fig. 2g). Collectively, we demonstrate how comprehensive profiling of coupled targets in distinct cancer subtypes when integrated with multi-omic genomics data, allows for the identification of mutant specific coupling for an oncogenic driver.

### Development of a C^2^-COUPLr for EML4-ALK

We next leveraged functional genomic essentiality mapping in the cell lines profiled in this study to identify coupled dependencies. We focused on targets that were differentially essential across cell lines and were coupled by the least reactive COUPLr in our informer set, SH-X-1C9 (Fig. 3a, Extended Data Fig. 6a). This revealed EML4 as a target of high-interest (Fig. 3a). EML4 is a microtubule binding protein that is the most common fusion partner (>90%) of the ALK tyrosine kinase in non-small cell lung cancers (NSCLCs)^122–125^. EML4 undergoes oligomerization through its coiled-coil domain both as a wildtype protein and in the context of the EML4-ALK fusion. Structure-function analysis revealed that SH-X-1C9 preferentially couples EML4 as a fusion protein rather than full length EML4 (Fig. 3b), but within the fusion the EML4 domain is sufficient for coupling (Extended Data Fig. 6b). Because SH-X-1C9 targets EML4-ALK outside of its kinase domain, we were not surprised that the molecular COUPLr could engage common EML4-ALK resistant mutations arising in the clinic^123, 126–129^ (Fig. 3c). Analysis of other coiled-coil containing proteins including YAP1^130–132^ or KIF5B^133, 134^, revealed that SH-X-1C9 was selective for EML4 (Extended Data Fig. 6c). Treatment with SH-X-1C9 stabilized the fusion in thermal stability assays (Extended Data Fig. 6d) and mutation of all cysteines to alanine within the EML4 portion of EML4-ALK ablated coupling (Extended Data Fig. 6e). However, no one cysteine was itself sufficient to mediate SH-X-1C9 coupling (Extended Data Fig. 6f) likely reflecting the multivalent nature of this modality.

**Fig. 3:**
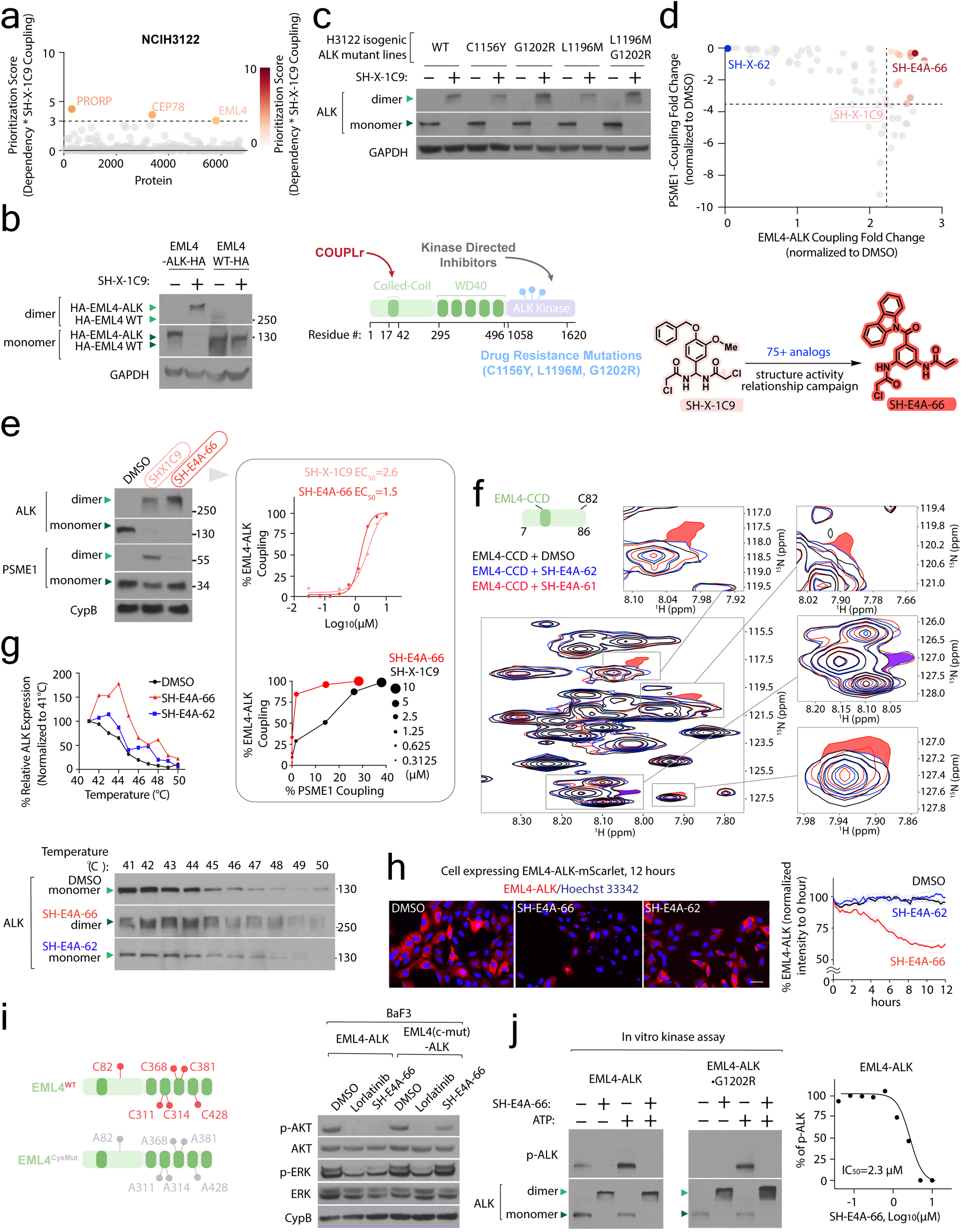
Development of a C^2^-COUPLr that disrupts EML4-ALK protein dynamics and blocks signaling. (a) Differentially coupled genetic dependencies. A prioritization score based on coupling with SH-X-1C9 and gene essentiality^176, 177^ highlights EML4 in NCI-H3122 cells as a target of interest. (b) SH-X-1C9 preferentially couples the EML4-ALK fusion compared to WT EML4. U2OS cells expressing WT EML4 or EML4-ALK were treated with 10 µM SH-X-1C9 for 3 hrs, and coupling was determined by immunoblot. (c) SH-X-1C9 couples EML4-ALK resistant mutations. Top, coupling of indicated EML4-ALK clinical variants^126, 178^ in isogenic H3122 cells lines was examined by immunoblot following treatment with 10 µM SH-X-1C9 (6 hr treatment). Bottom, schematic of EML4-ALK highlighting different domains and clinical mutations. (d) Development of an advanced EML4-ALK COUPLr. Top, structure-activity relationship (SAR) plot comparing EML4-ALK with off-target PSME1 coupling with different SH-X-1C9 analogs. Coupling was assessed by immunoblot following treatment in H3122 cell lysate with the indicated compounds (5 µM). Bottom, advancement of SH-X-1C9 to SH-E4A-66 (see also Extended Data Fig. S6i). (e) SH-E4A-66 couples EML4-ALK in cells, minimally targeting PSME1. H3122 cells were treated with 5 µM of the indicated compounds for 3 hrs, and coupling was assessed by immunoblot for EML4-ALK and PSME1. Inset, in vitro IC_50_ plot for SH-E4A-66 coupling of EML4-ALK and specificity over PSME1 targeting. (f) EML4 coupling alters protein conformations. 2D ^1^H-^15^N NMR analysis of EML4 (residues: 7-86) following treatment with vehicle (black), SH-E4A-62 (blue, mono-warhead analog), or SH-E4A-61 (red, dual-warhead analog). Insets display shared (purple) peak shifts for the mono and dual warhead analogs and those unique to the dual-warhead analog (red) (see also Methods). (g) EML4-ALK engagement by SH-E4A-66 increases thermal stability. H3122 cells were treated with 20 µM of the indicated compounds for 6 hrs, and EML4-ALK thermal stability was quantified (top) by immunoblot (bottom). (h) EML4-ALK coupling results in degradation. Left, representative immunofluorescent images of EML4-ALK-mScarlet expressed in U2OS cells following treatment with 2.5 µM of the indicated compounds for 12 hrs. Right, quantification of EML4-ALK-mScarlet levels following the indicated treatments over 12 hrs. (i) An EML4-ALK COUPLr disrupts signaling in a cysteine-dependent manner. BaF3 cells expressing EML4-ALK or EML4^CysMut^-ALK were treated with 5 µM of SH-E4A-66 for 2 hrs and pathway activity was determined by immunoblot. (j) SH-E4A-66 disrupts kinase activity of EML4-ALK and the EML4-ALK•G1202R mutant. Left, BaF3 cells expressing the indicated proteins were treated with vehicle or 10 µM of SH-E4A-66 for 3 hrs. Cells were lysed and EML4-ALK was immunoprecipitated with anti-HA beads and in vitro kinase was initiated following addition of ATP. Kinase activity was monitored by immunoblot for phosphorylated Tyr1278/1282/1283 in ALK. Right, SH-E4A-66 IC_50_ plot for EML4-ALK in vitro kinase activity. Scale bar= 50 µm. Data are represented as mean ± SD. ***p< 0.0001. Statistical significance determined by Student’s t-test (two-tailed, unpaired).

Given the structural simplicity of SH-X-1C9, we were not surprised to find additional proteins were also coupled by SH-X-1C9 in NCI-H3122 cells, including the proteasome subunit PSME1^61, 135–137^ (Extended Data Fig. 6g, Table S1). To increase the utility of this probe to study the consequences of EML4-ALK coupling, our primary focus in a structure-activity relationship (SAR) campaign was to improve potency for EML4-ALK while reducing PSME1 coupling. To this end, we generated 75 analogs varying warheads and substituents upon the central aryl ring, as well as their relative positions (e.g., exocyclic geminal dual-warheads vs 3,5-aryl dual-warhead). (Fig. 3d, Extended Data Fig. 6h). Moreover, we found the connectivity of the distal heterocyclic substituent (i.e., indole-SH-E4A-64 vs quinoline-SH-E4A-63 and carbazole-SH-E4A-59 vs diphenyl-SH-E4A-58) drastically enhanced or prevented coupling, respectively (Extended Data Fig. 6h). This suggests the necessity of molecular recognition and a structured pocket for coupling. Ultimately, this effort yielded SH-E4A-66 a C^2^-COUPLr with carbazole scaffold barring heterogeneous cysteine reactive warheads (acrylamide and chloroacetamide) (Fig. 3d, Extended Data Fig. 6h). SH-E4A-66 coupled EML4-ALK with an EC_50_=1.5 µM and displayed increased specificity in comparison to SH-X-1C9 (Fig. 3e). We synthesized a chemical control compound, SH-E4A-62, which maintains a single cysteine-reactive chloroacetamide warhead but has the second warhead replaced with a non-reactive acetamide group; allowing us to differentiate single liganding events with coupling. As expected, SH-E4A-62 could not couple EML4-ALK serving as a specificity control for coupling (Extended Data Fig. 6h). Importantly, crosslinkers did not couple EML4-ALK (Extended Data Fig. 6i). CONNECT analysis revealed similarly low levels of cysteine engagement between SH-E4A-66 and SH-E4A-62 (Extended Data Fig. 6j, Table S1) and moderate overall mobility shift by SH-E4A-66 (Extended Data Fig. 6j). Using a biotinylated version of SH-X-1C9 (SH-E4A-52), we found SH-E4A-66 could compete SH-E4A-52-based enrichment of EML4-ALK in NCI-H3122 cells (Extended Data Fig. 7a).

### Molecular coupling alters protein dynamics, EML4-ALK degradation and blockade of signaling

To begin to understand the biophysical impact of EML4-ALK molecular coupling, we determined structural changes by 2D NMR of the coiled-coil domain of EML4-ALK harboring Cys82 following treatment with SH-E4A-61, a bis-chloroacetamide analog of SH-E4A-66, and the mono warhead control SH-E4A-62 (Fig. 3d,f, Extended Data Fig. 7b). We observed a shared singular peak shift in EML4 following treatment with both the mono and dual warheads relative to vehicle control representing the likely engagement of EML4•C82 (Fig. 3f, purple region). We also detected distinct peak shifts emanating from the dual warhead SH-E4A-61, suggesting an alteration in protein dynamics (Fig. 3f, red regions). Thermal stability assays with EML4-ALK following in cell treatment with the dual warhead SH-E4A-66 revealed an increase in stability, whereas the mono warhead SH-E4A-62 was comparable to vehicle control (Fig. 3g). These results are suggestive of substantial changes in protein dynamics upon molecular coupling.

We hypothesized that one outcome of restricting protein dynamics in EML4-ALK might result in an alteration of protein stability^138–140^. To this end, we appended the mScarlet tag to EML4-ALK and monitored its stability following treatment with SH-X-1C9, SH-E4A-66 and SH-E4A-62. Dual warhead engagement led to a time-dependent degradation that was not observed with mono warhead liganding (Fig. 3h). An EML4-ALK cysteine mutant was not degraded by SH-E4A-66 or SH-X-1C9 and this coupling-mediated degradation was proteasome-dependent for both C^2^-COUPLrs (Extended Data Fig. 7c-d).

To determine whether coupling altered ALK signaling, we constructed a BaF3 cell line expressing WT EML4-ALK or EML4^CysMut^-ALK mutant, finding SH-E4A-66 inhibited AKT•S473 phosphorylation which was not observed with the EML4^CysMut^-ALK (Fig. 3i) Treatment with SH-E4A-62 did not disrupt ALK signaling in BaF3 cell lines (Extended Data Fig. 7e). To further explore the impact of coupling directly on ALK kinase activity, we adapted an *in vitro* kinase assay finding that SH-E4A-66 disrupts EML4-ALK kinase activity with an IC_50_=2.3 µM (Fig. 3j, Extended Data Fig. 7f-g). The blockade in ALK phosphorylation extended to an EML4-ALK variant identified in patients resistant to ALK-kinase directed inhibitors (Fig. 3j). Finally, we found partial disruption in ALK signaling in an EML4-ALK cell line (MGH-006) derived from a patient treated at MGH (Extended Data Fig. 7h). Collectively, we find that by restricting protein dynamics through coupling, EML4-ALK signaling is blocked and leads to protein degradation, suggesting a potentially new mode to inhibit this oncogenic fusion kinase, independent of ATP-directed kinase inhibitors.

## Discussion

In this study we demonstrate that molecular COUPLrs, when combined with CONNECT analysis, can systematically and unbiasedly identify proteins amenable to chemical connectivity. Historically, there have been limited rational approaches to identify both this class of target proteins and their corresponding small molecule probes^84, 141, 142^. Our development of a C^2^-COUPLr library and subsequent analysis of its targets in several cell lines reveals that proteins amenable to coupling are distributed throughout the proteome localizing to dozens of protein classes not traditionally appreciated as targets of existing molecular glues^143–149^. Thus, molecular COUPLrs and the CONNECT platform can be widely applied to uncover a fuller spectrum of chemically connected targets in many biological processes and disease states without prior knowledge. This approach may also lend itself well to the unbiased identification of protein-protein interactions and their interfaces that are amenable to small molecule binding. An advantage of COUPLr technology is that it defines a chemical lexicon which is in principle expandable to any permutation of reactive side chains, enabling a wider investigation of cellular biology at the residue level. On this understanding, we envision the future development of Cys-Lys, i.e. CK-COUPLrs, as well as Cys-Tyr, i.e. CY-COUPLrs, and more.

In comparison to their heterobifunctional counterparts both molecular glues and COUPLrs are characterized by a smaller chemical footprint and more drug-like properties. Moreover, the corresponding change in protein complex dynamics following binding of either class represents an exciting opportunity to target this fundamental biophysical property of proteins. With the aid of deep structural characterization, determining how protein dynamics are altered by molecular glues or COUPLrs will be essential. The biggest differentiator between these two modalities is the doubly covalent nature of COUPLrs, which provides for rapid target deconvolution, akin to other covalent modalities. This feature, along with structural resolution, will be central in rapidly deciphering the rules for coupling protein complexes, following a similar trajectory that has enabled our broader understanding of the chemical features required for gluing mTOR to FKBP12 or CRBN1 to IKZF1^150^.

What remains unknown at present, is whether molecular COUPLrs may also induce chemical proximity similar to molecular glues. To this end a promising future direction is to determine the interchangeability of molecular glues and COUPLrs: can one class of molecule be converted to the other? While powerful medicinal chemistry approaches are required for this to be borne out, if this is indeed the case, the potential scope of established and neo protein complexes that can be targeted with small molecules may be substantially expanded, providing a systematic approach to target protein complexes.

In recent years the drug discovery community has championed the use of covalency as a focused tool for both ligand discovery and drug development for several challenging protein classes^151–156^. The use of two covalent warheads in molecular COUPLrs may at first raise concerns of increased reactivity and non-selective interactions, reminiscent of crosslinkers. However, our comparison of COUPLr targets and those of crosslinkers, demonstrates that this form of chemical connectivity requires defined binding elements. This accords well with previous findings demonstrating that cysteine engagement by covalent fragments is disrupted by protein denaturation^91^. Here, our interpretation for the selectivity of COUPLrs when compared to crosslinkers, is that with the latter, a lack of a binding element precludes the required residence time for productive engagement^157–159^. A head-to-head comparison of matched mono-warhead and dual-warhead scaffolds revealed a near identical target scope. While the principles that govern molecular coupling are still emerging, the requirement for dual engagement likely constrains the scope of targets compared to mono warheads ligands. Future kinetic studies which define the precise order of engagement each warhead takes will be necessary to test this hypothesis.

Combining structural, chemical and genomic analysis reveals that coupling of 1000+ proteins is shaped by the unique binding element of a given COUPLr and the distinct conformation of its target. This former property of coupling is best exemplified by the distinct targets engaged between different COUPLr stereoisomer pairs, where we found that SH-X-61B, but not its isomer SH-X-61A, binds DNMT3A. We also found that coupling may be generally associated to protein conformations as shaped by amino acid mutations. NPM1c provided a striking example of mutant-specific coupling that arose from *de novo* encoding an aberrant nuclear export signal (NES). The increased liganding of NPM1c•C275 is an example of how proximal mutations^90^, in this case the introduction of an NES, changes the ligandability of cysteines within proteins. How the mutant NES alters NPM1 conformations remains to be determined but provides an example of how this approach reveals changes in protein conformations through the readout of coupling and its ability to be leveraged to target prominent mutations within cancer.

Leveraging the identification of coupled targets, we have begun to reveal how restricting the protein dynamics of EML4-ALK results in altered signaling. While it has been long appreciated that protein dynamics are required for protein function, the small molecule rationale for targeting these states is only now emerging^113, 160–165^, and at best has yet to be systematically applied. In an exciting turn, molecular COUPLrs may represent a modality that is uniquely suited to target these functional states within proteins through their bifunctional modification. The precise biophysical mechanism by which restricting EML4 dynamics leads to a decrease in ALK kinase activity remains to be determined and will likely benefit from further structural resolution and/or functional genomic approaches used to map residues required for small molecule regulation of protein function^166–171^. Protein complexation is a common form of biological regulation. Because multiple oncogenic kinases are activated by this mechanism and the data collected herein suggests coupling outside of their kinase domain is a possibility, this approach may lend itself well to disrupting signaling cascades outside of ATP-mediated competition and circumvent current mechanisms of resistance. While the EML4-ALK COUPLr functions as a starting point to explore these concepts, additional medicinal chemistry optimization afforded to drugs will be needed to carry these ideas to fruition.

The presence of two or more covalent warheads embedded in a substantial fraction of natural products, raises the tantalizing possibility that this modality is widely used in nature to control biological programs. While the full extent of biological processes impacted by dual warhead natural products remains to be determined, covalent modification by mono warhead natural products has demonstrated regulation of many signal transduction pathways, with some taken into clinical use^172–174^. It would not be surprising if dual warhead natural products also regulate a distinct set of biological processes through the control of protein dynamics, representing how molecular COUPLrs can shape cellular physiology.

## Acknowledgements

We thank all members of the Bar-Peled and Iafrate Labs, and David Sabatini and Keith Flaherty for helpful suggestions. We thank Julian Gruenewald, Ragan Szalay, and Keith Joung for help generating isogenic H3122 ALK mutant cell lines. This work was supported by the Damon Runyon Cancer Research Foundation (62-20 to L.B-P., 60S-20 to B.L.L.), the American Association for Cancer Research (19-20-45-BARP), the American Cancer Society (RSG-24-1199432-01-TBE), the Melanoma Research Alliance, the Ludwig Cancer Center of Harvard Medical School, the Lungevity Foundation, ALK Positive Foundation, V-Foundation, Mary Kay Foundation, Paula and Rodger Riney Foundation, the PEW-Stewart Trusts, Lisa and Mark Schwartz, the Krantz Center for Cancer Research Quantum Award, the NIH/NCI (1R21CA226082-01, R37CA260062, R01CA285415 to L.B-P., 1DP2GM137494, R35GM153476 to B.B.L) DF/HCC SPORE in Gastrointestinal Cancer, NIH/NCI (P50CA127003), and the DF/HCC SPORE in Lung Cancer.

## Author contributions

D.Y., S.H., A.J.I., and L.B-P conceived and designed the study. S.H. synthesized COUPLr compounds tested in the studies and analyzed with NMR with the assistance of N.C., C.L., G.M., H.X.Y.L., Z.C., I.B., under the guidance of B.B.L. and L.B.P. S.H., D.Y., E.Z., N.N, S.T. performed proteomic experiments with the assistance of M.H., K.E., I.B., and the LC/MS were performed by S.Z. and N.K.. D.Y., E.Z., N.N., performed most of the molecular biology and cloning experiments with the assistance of W.Y., C.G., Z.H. A.S., A.D.C., M.G., N.K., M.J.L., M.O.. H.B.C., E.C., D.M., and S.K. performed bioinformatics analysis and data visualization with the assistance of M.G. H.B.C. and G.P. developed the machine learning model to predict targets amenable to coupling. Z-Y.J.S., G.H., H-S.S, and S.D-P performed the protein structural analysis. M,M., C.G., and R.J.S. assisted with live cell imaging experiment. T.F. and A.N.H. generated EML4-ALK patient cells line and assisted with generation of the BaF3-EML4-ALK cell line. D.Y., S.H., H.B.C., A.J.I., and L.B-P. wrote the manuscript with assistance from all the coauthors. A.J.I., and L.B-P. supervised the studies.

## Methods

### Organic Chemistry

Unless specified otherwise: all reactions were performed in oven-dried or flame-dried glassware under a positive pressure of nitrogen unless otherwise noted. Column chromatography was performed with a Biotage Isolera system employing silica gel 60 (230-400 mesh, Silicycle). Celite filtration was performed using Celite® 545 (EMD Millipore). Analytical thin-layer chromatography (TLC) was performed using 0.25 mm silica gel 60 F254 (EMD Millipore). TLC plates were visualized by fluorescence quenching under ultraviolet light (UV) and exposure to a solution of ceric ammonium molybdate, *p*-anisaldehyde, or potassium permanganate stain followed by heating on a hot plate. Reversed-phase chromatography was performed on an Agilent 1200 series HPLC using water and acetonitrile as a mobile phase and Agilent Pursuit C18 as a stationary phase. Commercial reagents and solvents were used as received with the following exceptions: tetrahydrofuran, diethyl ether, dichloromethane, toluene, acetonitrile, and *N*,*N*-dimethylformamide were degassed with argon and passed through a solvent purification system (designed by Pure Process Technology) utilizing alumina columns.

NMR spectra were recorded with a Varian INOVA-500 spectrometer, are reported in parts per million (δ), and are calibrated using residual undeuterated solvent as an internal reference (CDCl_3_: δ 7.26 for ^1^H NMR; CD_3_CN: δ 1.94 for ^1^H NMR and δ 118.26; CD_3_OD: δ 3.31 for ^1^H NMR). Data for 1H NMR spectra are reported as follows: chemical shift (δ ppm) (multiplicity, coupling constant (Hz), integration). Multiplicities are reported as follows: s = singlet, d = doublet, t = triplet, q = quartet, m = multiplet, br = broad, or combinations thereof. High-resolution mass spectra (HRMS) were recorded using electrospray ionization (ESI) mass spectroscopy experiments on an Agilent 6210 TOF LC/MS. Please refer to **Extended Data Fig. 8** for NMR spectra.

### Synthesis of 3-(2-chloroacetamido)-4-(2-(2-chloroacetamido)phenoxy)-*N*-(prop-2-yn-1-yl)benzamide (SH-X-9)

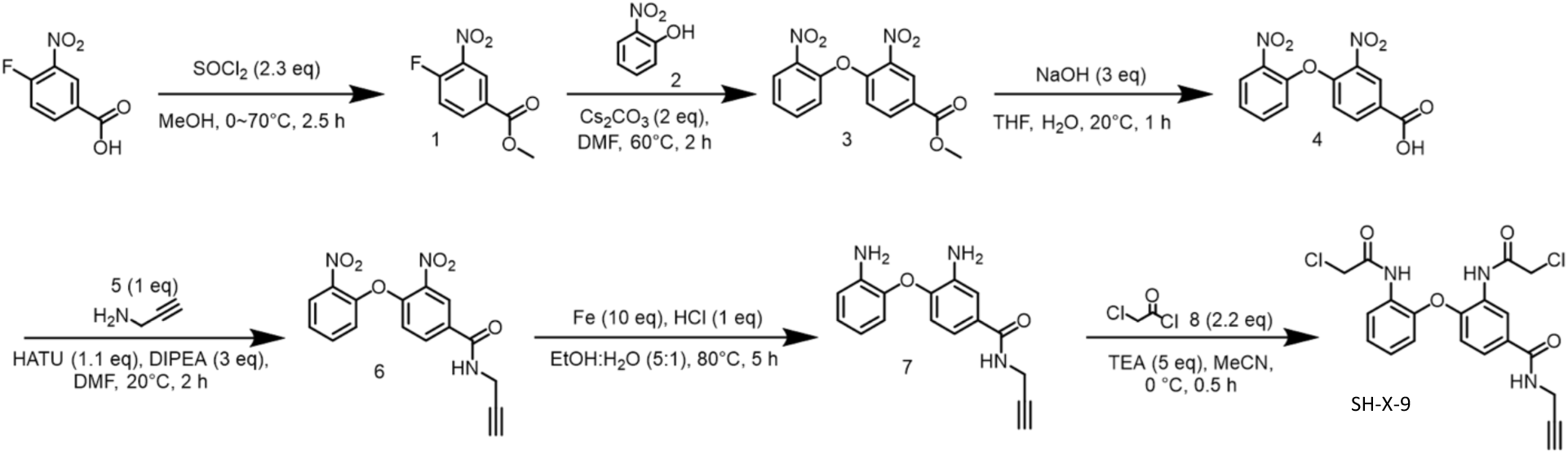

#### Step 1: Preparation of methyl 4-fluoro-3-nitrobenzoate (1)

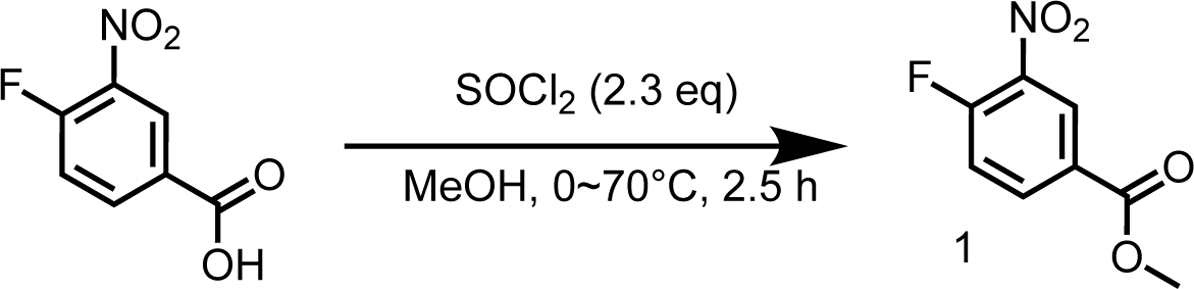

To a solution of 4-fluoro-3-nitrobenzoic acid (5.00 g, 27.0 mmol) in MeOH (100 mL) was added SOCl_2_ (7.39 g, 62.1 mmol, 4.51 mL) at 0 °C. The reaction mixture was stirred at 70 °C for 2 hrs. TLC (Petroleum ether: Ethyl acetate=5:1) indicated that 4-fluoro-3-nitrobenzoic acid was consumed completely and one new spot was formed. The residue was triturated with water and the mixture was filtered and the filter cake was concentrated to give methyl 4-fluoro-3-nitrobenzoate (5.00 g, 25.1 mmol, 92.9% yield) as a white solid.

^1^H NMR (400 MHz, DMSO-*d*_6_): *δ* 8.58 (dd, *J* = 2, 7.2 Hz, 1H), 8.39-8.30 (m, 1H), 7.75 (dd, *J* = 8.8, 11.2 Hz, 1H), 3.91 (s, 3H).

#### Step 2: Preparation of methyl 3-nitro-4-(2-nitrophenoxy)benzoate (3)

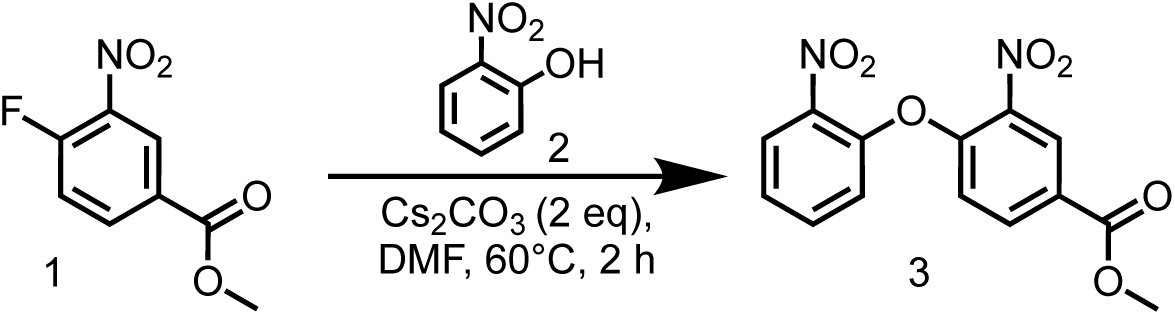

To a solution of methyl 4-fluoro-3-nitrobenzoate (5.00 g, 25.1 mmol) and 2-nitrophenol (3.49 g, 25.1 mmol) in DMF (100 mL) was added Cs_2_CO_3_ (16.4 g, 50.2 mmol). The reaction mixture was stirred at 60 °C for 2 hrs. TLC (Petroleum ether: Ethyl acetate=5:1) indicated that methyl 4-fluoro-3-nitrobenzoate was consumed completely and two new spots were formed. The reaction mixture was poured into H_2_O (200 mL) and extracted with EtOAc (200 mL×3). The organic layers were collected and washed with brine (200 mL×3), dried over Na_2_SO_4_, and concentrated to give a residue. The residue was purified by column chromatography on silica gel (eluted with petroleum ether: ethyl acetate = 10:0 to 1:1) to give methyl 3-nitro-4-(2-nitrophenoxy)benzoate (7.00 g, 22.0 mmol, 87.6% yield) as a yellow solid.

LC-MS: 341.0 [M+Na]^+^ / Ret time: 0.446 min / method: 5-95AB_0.8min.lcm

#### Step 3: Preparation of 3-nitro-4-(2-nitrophenoxy)benzoic acid (4)

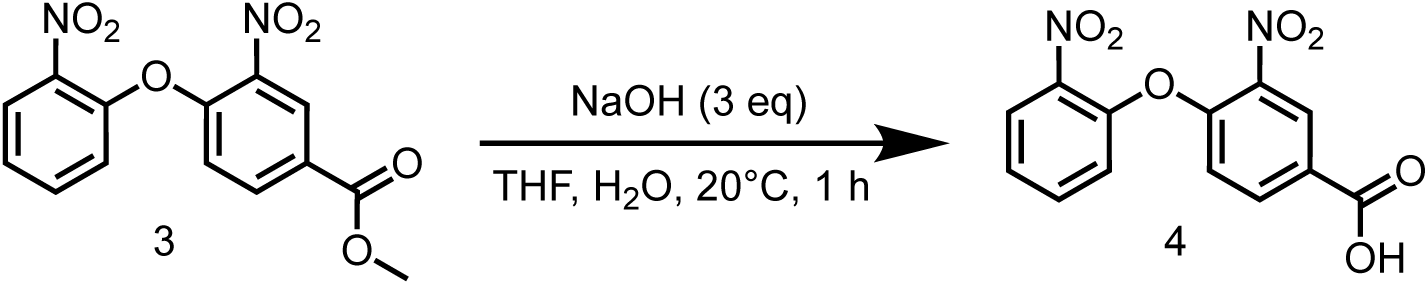

To a solution of methyl 3-nitro-4-(2-nitrophenoxy)benzoate (7.00 g, 22.0 mmol) in THF (70 mL) and H_2_O (20 mL) was added NaOH (2.64 g, 66.0 mmol). The mixture was stirred at 20 °C for 2 hrs. LCMS showed the reaction was completed. The reaction mixture was concentrated in vacuum to remove solvent, and then acidified with 1M HCl to pH=4. The reaction mixture was filtered, and the filter cake was collected, concentrated in vacuum to give 3-nitro-4-(2-nitrophenoxy)benzoic acid (6.40 g, 21.1 mmol, 95.6% yield) as a yellow solid.

LC-MS: 302.9 [M+H]^+^ / Ret time: 0.311 min / method: 5-95CD_1min_NEG.lcm

#### Step 4: Preparation of 3-nitro-4-(2-nitrophenoxy)-*N*-(prop-2-yn-1-yl)benzamide (6)

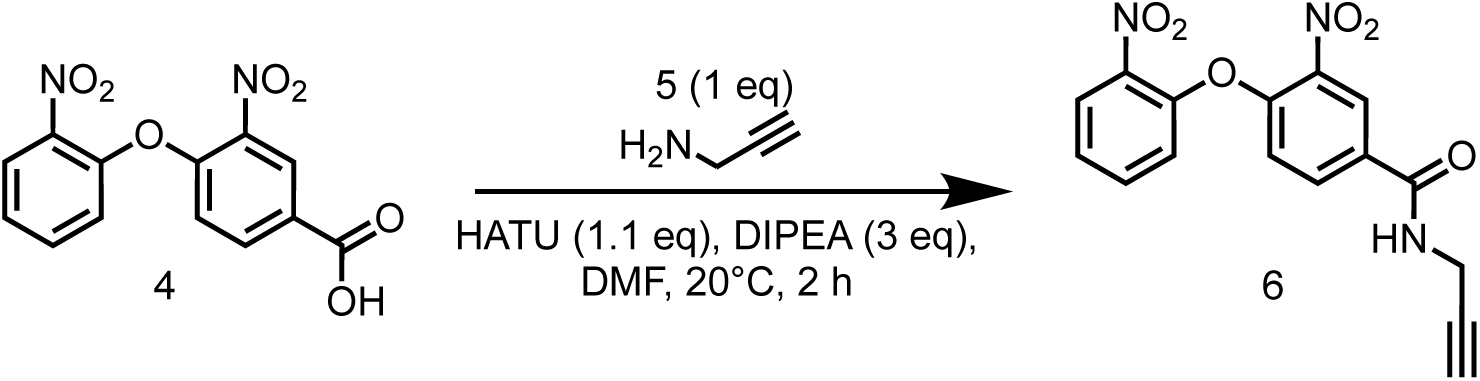

To a solution of 3-nitro-4-(2-nitrophenoxy)benzoic acid (6.40 g, 21.0 mmol) in DMF (65 mL) was added HATU (8.80 g, 23.1 mmol), DIPEA (8.16 g, 63.1 mmol, 10.9 mL), then the prop-2-yn-1-amine (1.16 g, 21.0 mmol, 1.35 mL) was added into the reaction mixture and stirred at 20 °C for 2 hrs. LCMS showed the reaction was completed. The reaction mixture was poured into H_2_O (150 mL), extracted with EtOAc (100 mL×3). The organic layers were collected and washed with brine (100 mL×3), dried over Na_2_SO_4_ and concentrated to give a residue. The residue was purified by column chromatography on silica gel (eluted with petroleum ether: ethyl acetate=5:1 to 1:2) to give 3-nitro-4-(2-nitrophenoxy)-*N*-(prop-2-yn-1-yl)benzamide (3.00 g, 8.79 mmol, 41.8% yield) as a yellow solid.

LC-MS: 342.1 [M+H]^+^ / Ret time: 0.396 min / method: 5-95AB_0.8min.lcm

#### Step 5: Preparation of 3-amino-4-(2-aminophenoxy)-*N*-(prop-2-yn-1-yl)benzamide (7)

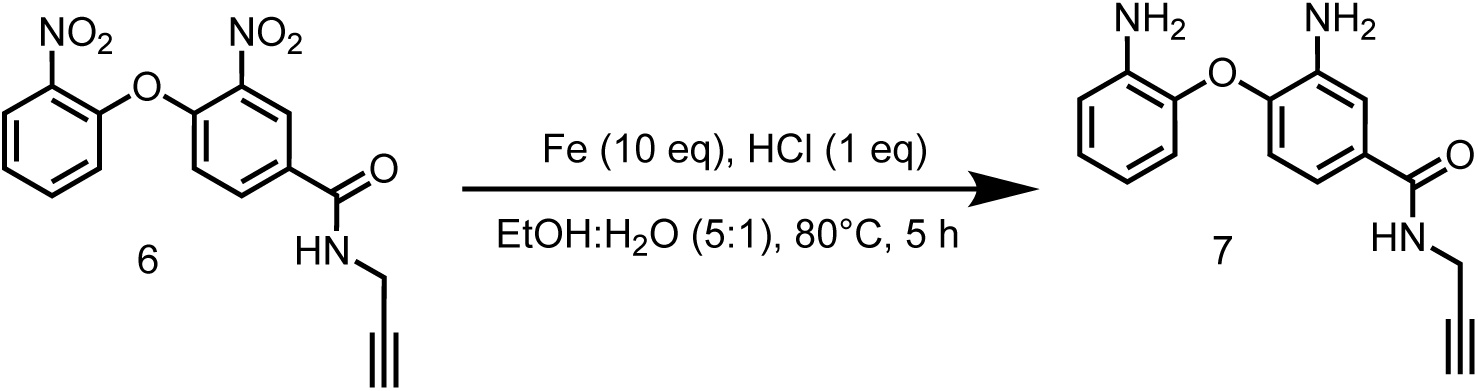

To a solution of 3-nitro-4-(2-nitrophenoxy)-*N*-(prop-2-yn-1-yl)benzamide (1.10 g, 3.22 mmol) and Fe (900 mg, 16.1 mmol) in EtOH (20 mL) and H_2_O (4 mL) was added concentrated HCl (12 M, 269 μL), and the resulting mixture was stirred at 80 °C for 4 hrs. TLC indicated 3-nitro-4-(2-nitrophenoxy)-*N*-(prop-2-yn-1-yl)benzamide was remained, and one major new spot with larger polarity was detected. The mixture was added Fe (900 mg, 16.1 mmol), then the resulting mixture was stirred at 80 °C for another 1 hr. LCMS showed the reaction was completed. The mixture was filtered and the filtrate was concentrated to a residue. The residue was purified by preparative– HPLC (column: Phenomenex luna C18 150*25mm* 10um; mobile phase: [water(FA)-MeCN]; gradient: 10%-40% B over 10 min) to give 3-amino-4-(2-aminophenoxy)-*N-*(prop-2-yn-1-yl)benzamide (45.0 mg, 160 μmol, 4.96% yield) as a brown solid.

LC-MS: 282.2 [M+H]^+^ / Ret time: 0.264 min / method: 5-95AB_0.8min.lcm

^1^H NMR (400 MHz, DMSO-*d*_6_): *δ* 8.61 (t, *J* = 5.6 Hz, 1H), 7.28 (d, *J* = 2.0 Hz, 1H), 6.97 (dd, *J* = 2.0, 8.4 Hz, 1H), 6.92-6.86 (m, 1H), 6.82-6.78 (m, 1H), 6.73 (d, *J* = 8.0 Hz, 1H), 6.57-6.51 (m, 2H), 5.15 (s, 2H), 4.92 (s, 2H), 3.99 (dd, *J* = 2.4, 5.6 Hz, 2H), 3.08 (t, *J* = 2.4 Hz, 1H).

#### Step 6: Preparation of 3-(2-chloroacetamido)-4-(2-(2-chloroacetamido)phenoxy)-*N*-(prop-2-yn-1-yl)benzamide (SH-X-9)

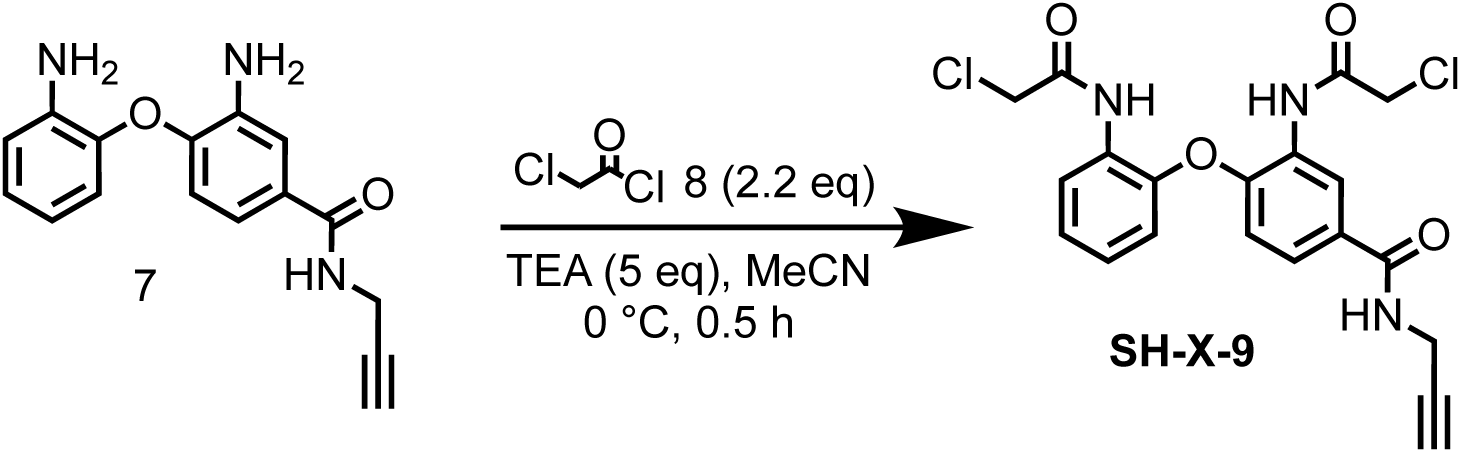

To a solution of 3-amino-4-(2-aminophenoxy)-*N*-(prop-2-yn-1-yl)benzamide (150 mg, 533 μmol) in MeCN (2 mL) was added TEA (270 mg, 2.67 mmol, 372 μL), then 2-chloroacetyl chloride (133 mg, 1.17 mmol, 93.4 μL) was added into the reaction mixture at 0 °C, The mixture was stirred at 0 °C for 0.5 hr. LCMS showed the reaction was completed. The reaction mixture was poured into H_2_O (5 mL), extracted with EtOAc (5 mL×3). The organic layers were collected and washed with brine (5 mL×3), dried over Na_2_SO_4_ and concentrated to give a residue. The residue was purified by preparative-TLC (petroleum ether: ethyl acetate=1:1) to give 3-(2-chloroacetamido)-4-(2-(2-chloroacetamido)phenoxy)-*N*-(prop-2-yn-1-yl)benzamide (20.9 mg, 46.9 μmol, 8.80% yield, 97.5% purity) as a white solid.

LC-MS: 434.2 [M+H]^+^ / Ret time: 0.348 min / method: 5-95AB_0.8min.lcm

^1^H NMR (400 MHz, DMSO-*d*_6_): *δ* 10.10 (s, 1H), 9.89 (s, 1H), 8.90 (s, 1H), 8.40 (s, 1H), 8.03 (d, *J* = 7.6 Hz, 1H), 7.64 (d, *J* = 8.8 Hz, 1H), 7.27-7.17 (m, 2H), 7.01 (d, *J* = 7.6 Hz, 1H), 6.83 (d, *J* = 8.4 Hz, 1H), 4.39 (s, 2H), 4.31 (s, 2H), 4.03 (d, *J* = 3.6 Hz, 2H), 3.11 (s, 1H)

### Synthesis of 3-acrylamido-4-(2-acrylamidophenoxy)-*N*-(prop-2-yn-1-yl)benzamide (SH-X-10)

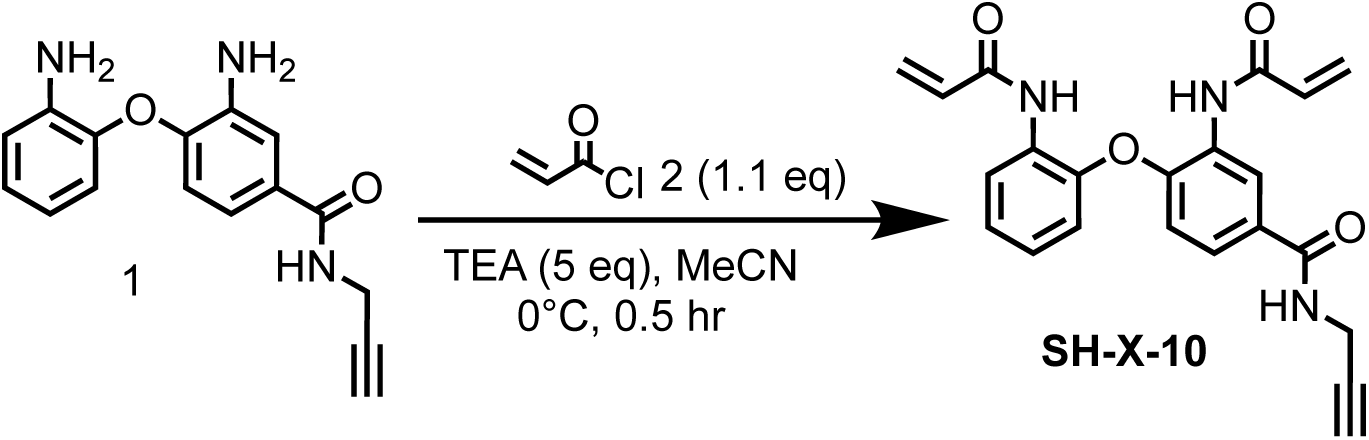

To a solution of 3-amino-4-(2-aminophenoxy)-*N*-(prop-2-yn-1-yl)benzamide (500 mg, 1.78 mmol) in MeCN (20 mL) was added TEA (899 mg, 8.89 mmol, 1.24 mL), then acryloyl chloride (177 mg, 1.96 mmol, 160 μL) in MeCN (10 mL) was added into the reaction mixture at 0 °C, the reaction mixture was stirred at 0 °C for 0.5 hr. LCMS showed the reaction was completed. The reaction mixture was poured into H_2_O (20 mL), extracted with EtOAc (20 mL×3). The organic layers were collected and washed with brine (20 mL×3), dried over Na_2_SO_4_ and concentrated to give a residue. The residue was purified by column chromatography on silica gel (eluted with petroleum ether: ethyl acetate=1:2) to give 3-acrylamido-4-(2-acrylamidophenoxy)-*N*-(prop-2-yn-1-yl)benzamide (79.0 mg, 201 μmol, 11.3% yield, 99.0% purity) as a white solid.

LC-MS: 390.1 [M+H]^+^ / Ret time: 0.383 min / method: 5-95AB_0.8min.lcm

^1^H NMR (400 MHz, DMSO-*d*_6_): *δ* 10.19 (s, 1H), 10.00 (s, 1H), 8.90 (t, *J* = 5.6 Hz, 1H), 8.36 (s, 1H), 8.11 (d, *J* = 7.6 Hz, 1H), 7.64 (d, *J* = 8.8 Hz, 1H), 7.27-7.16 (m, 2H), 7.00 (d, *J* = 7.6 Hz, 1H), 6.81 (d, *J* = 8.8 Hz, 1H), 6.63-6.52 (m, 2H), 6.34-6.22 (m, 2H), 5.82 (d, *J* = 10.8 Hz, 1H), 5.77-5.73 (m, 1H), 4.04 (d, *J* = 3.2 Hz, 2H), 3.14-3.10 (m, 1H).

### Synthesis of 2-chloro-*N*-(2-(4-(2-chloroacetyl)-1-(prop-2-yn-1-yl)piperazine-2-carbonyl)-1,2,3,4-tetrahydroisoquinolin-6-yl)acetamide (SH-X-1A)

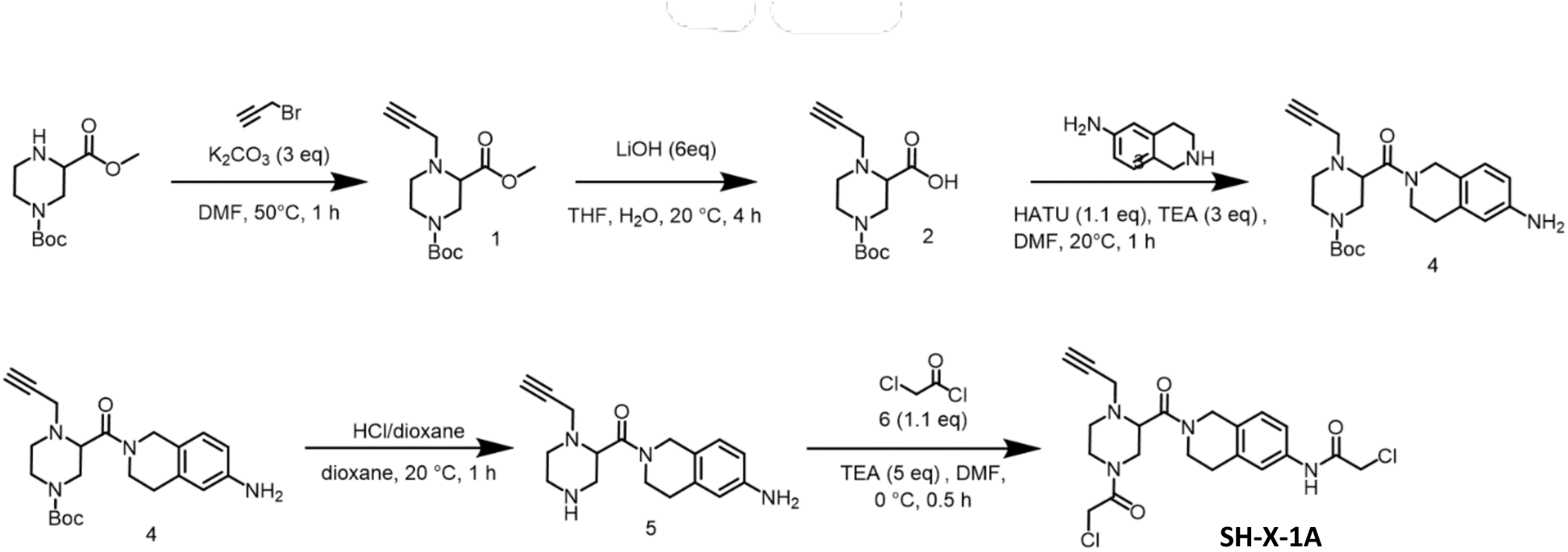

#### Step 1: Preparation of 1-(*tert*-butyl) 3-methyl 4-(prop-2-yn-1-yl)piperazine-1,3-dicarboxylate (1)

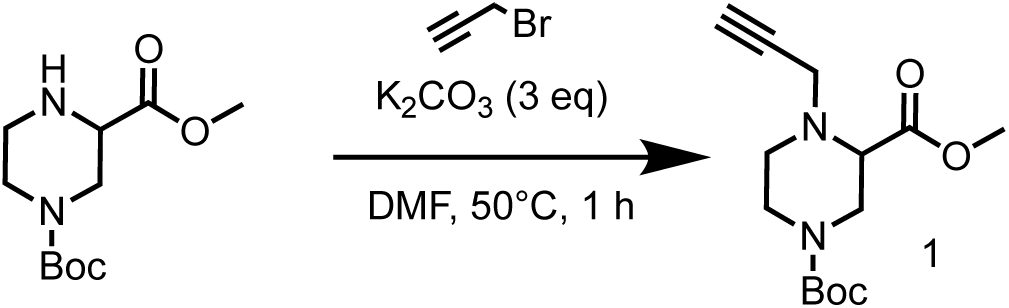

To a solution of 1-(*tert*-butyl) 3-methylpiperazine-1,3-dicarboxylate (4.00 g, 16.4 mmol) in DMF (40 mL) was added K_2_CO_3_ (6.79 g, 49.1 mmol), the reaction mixture was stirred at 20 °C for 0.5 hr. Then 3-bromoprop-1-yne (2.53 g, 21.3 mmol, 1.83 mL) was added into the reaction mixture, the reaction mixture was stirred at 50 °C for 1 hr. LCMS showed the reaction was completed. The reaction mixture was poured into H_2_O (100 mL), extracted with EtOAc (100 mL×3). The organic layers were collected and washed with brine (100 mL×3), dried over Na_2_SO_4_ and concentrated to give a residue. The residue was purified by column chromatography on silica gel (eluted with petroleum ether: ethyl acetate=2:1 to 3:1) to give 1-(*tert*-butyl) 3-methyl 4-(prop-2-yn-1-yl)piperazine-1,3-dicarboxylate (4.5 g, 15.9 mmol, 97.3% yield) as a yellow solid.

^1^H NMR (400 MHz, DMSO-*d*_6_): *δ* 3.64 (s, 3H), 3.60-3.48 (m, 2H), 3.43 (d, *J* = 2.4 Hz, 2H), 3.36 (s, 2H), 3.31 (s, 1H), 3.20 (t, *J* = 2.4 Hz, 1H), 2.89 (s, 1H), 2.73 (s, 1H), 1.38 (s, 9H).

#### Step 2: Preparation of 4-(*tert*-butoxycarbonyl)-1-(prop-2-yn-1-yl)piperazine-2-carboxylic acid (2)

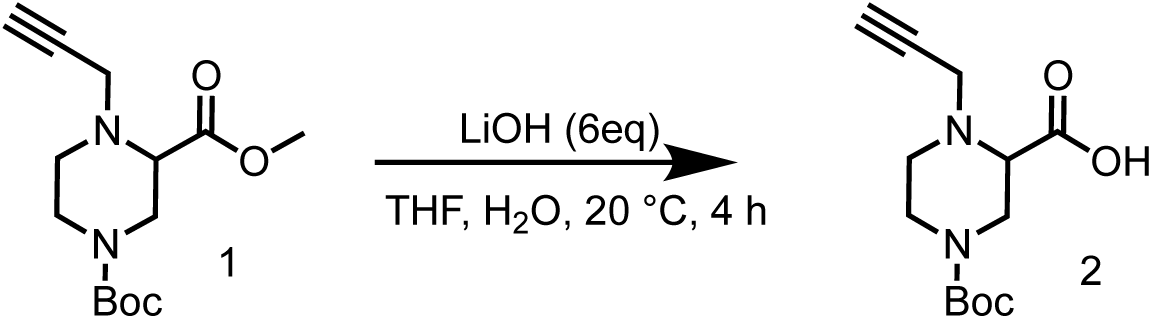

To a solution of 1-(*tert*-butyl) 3-methyl 4-(prop-2-yn-1-yl)piperazine-1,3-dicarboxylate (4.5 g, 15.94 mmol) in THF (50 mL) and H_2_O (15 mL)was added LiOH•H_2_O (4.02 g, 95.6 mmol). The mixture was stirred at 20 °C for 2 hrs. TLC (Petroleum ether: Ethyl acetate=3/1) indicated reaction was consumed completely and one new spot formed. The reaction mixture was concentrated on vacuum to remove excess solvent, and then acidified with 1M HCl to pH=4, the mixture was poured into H_2_O (50 mL), extracted with EtOAc (100 mL×3). The organic layers were collected and washed with brine (100 mL×3), dried over Na_2_SO_4_ and concentrated to give 4-(*tert*-butoxycarbonyl)-1-(prop-2-yn-1-yl)piperazine-2-carboxylic acid (3.30 g, 12.3 mmol, 77.2% yield) as a yellow solid.

^1^H NMR (400 MHz, DMSO-*d*_6_): *δ* 13.09-12.09 (m, 1H), 3.53 (d, *J* = 2.4 Hz, 2H), 3.43-3.20 (m, 4H), 3.20-3.16 (m, 1H), 2.78 (s, 1H), 2.49-2.38 (m, 2H), 1.41-1.36 (m, 9H).

#### Step 3: Preparation of *tert*-butyl 3-(6-amino-1,2,3,4-tetrahydroisoquinoline-2-carbonyl)-4-(prop-2-yn-1-yl)piperazine-1-carboxylate (4)

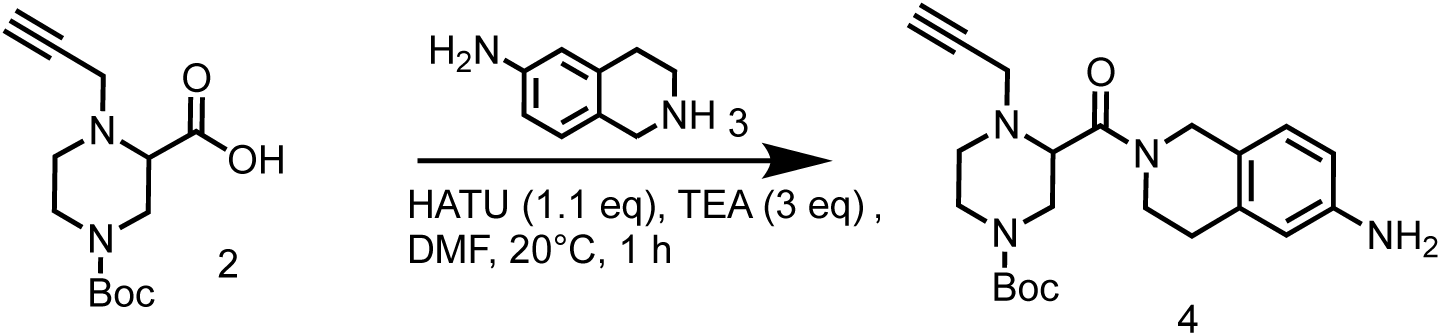

To a solution of 4-(*tert*-butoxycarbonyl)-1-(prop-2-yn-1-yl)piperazine-2-carboxylic acid (1.50 g, 5.59 mmol) in DMF (15 mL) was added HATU (2.34 g, 6.15 mmol) and TEA (1.70 g, 16.8 mmol, 2.33 mL) and 1,2,3,4-tetrahydroisoquinolin-6-amine (1.14 g, 6.15 mmol). The mixture was stirred at 20 °C for 1 hr. LCMS showed the reaction was completed. The reaction mixture was poured into H_2_O (100 mL), extracted with EtOAc (100 mL×3). The organic layers were collected and washed with brine (100 mL×3), dried over Na_2_SO_4_ and concentrated to give a residue. The residue was purified by column chromatography on silica gel (eluted with petroleum ether: ethyl acetate=1:3 to 0:1) to give *tert*-butyl 3-(6-amino-1,2,3,4-tetrahydroisoquinoline-2-carbonyl)-4-(prop-2-yn-1-yl)piperazine-1-carboxylate (1.60 g, 4.02 mmol, 71.8% yield) as a yellow solid.

LC-MS: 399.1 [M+H]^+^ / Ret time: 0.258 min / method: 5-95AB_0.8min.lcm

^1^H NMR (400 MHz, DMSO-*d*_6_): *δ* 6.80 (d, *J* = 8.0 Hz, 1H), 6.41 (dd, *J* = 2.0, 8.0 Hz, 1H), 6.34 (s, 1H), 4.91 (s, 2H), 4.76-4.64 (m, 1H), 4.57-4.35 (m, 1H), 3.87-3.58 (m, 4H), 3.56-3.43 (m, 1H), 3.24-3.18 (m, 1H), 3.17-3.04 (m, 2H), 2.97-2.83 (m, 2H), 2.78-2.66 (m, 2H), 2.62-2.55 (m, 1H), 2.45-2.36 (m, 1H), 1.38 (s, 9H).

#### Step 4: Preparation of (6-amino-3,4-dihydroisoquinolin-2(1H)-yl)(1-(prop-2-yn-1-yl) piperazin-2-yl)methanone (5)

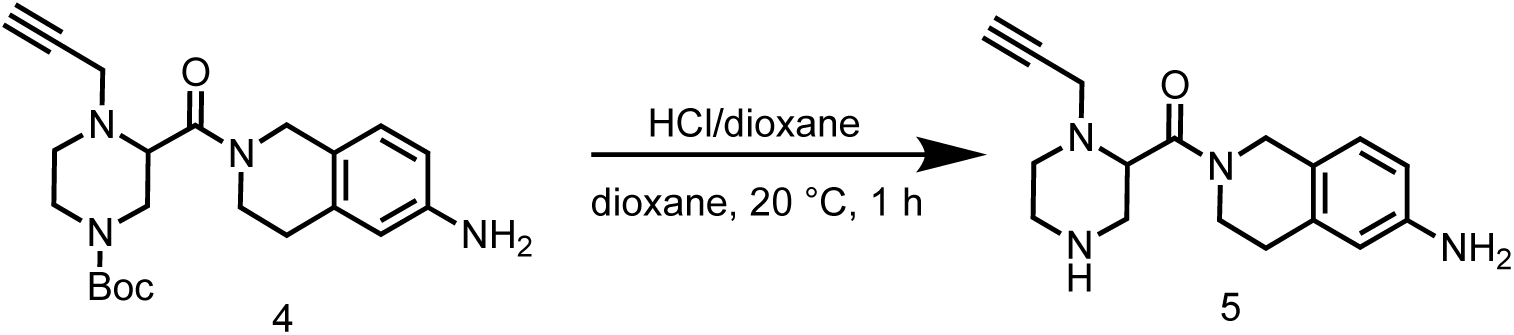

To a solution of *tert*-butyl 3-(6-amino-1,2,3,4-tetrahydroisoquinoline-2-carbonyl)-4-(prop-2-yn-1-yl)piperazine-1-carboxylate (1.60 g, 4.02 mmol) in dioxane (10 mL) was added HCl/dioxane (4 M, 10 mL). The mixture was stirred at 20 °C for 1 hr. LCMS showed the reaction was completed. The reaction mixture was concentrated in vacuum to remove excess solvent, then triturated with DCM and the mixture was filtered and the filter cake was concentrated to give a residue. The residue was purified by preparative-HPLC (column: Waters xbridge 150*25mm 10um; mobile phase: [water(NH_4_HCO_3_)-MeCN]; gradient: 1%-31% B over 10 min) to give (6-amino-3,4-dihydroisoquinolin-2(1H)-yl)(1-(prop-2-yn-1-yl)piperazin-2-yl)methanone (38.9 mg, 125 μmol, 3.13% yield, 96.3% purity) as a white solid.

LC-MS: 299.1 [M+H]^+^ / Ret time: 0.533 min / method: 0-60N_1min.lcm

^1^H NMR (400 MHz, DMSO-*d*_6_): *δ* 6.79 (d, *J* = 8.0 Hz, 1H), 6.40 (dd, *J* = 2.4, 8.0 Hz, 1H), 6.33 (s, 1H), 4.97-4.78 (m, 2H), 4.78-4.67 (m, 1H), 4.63-4.29 (m, 2H), 3.92-3.71 (m, 2H), 3.69-3.51 (m, 2H), 3.30-3.10 (m, 4H), 2.73 (s, 2H), 2.70-2.63 (m, 2H), 2.56 (t, *J* = 5.6 Hz, 1H), 2.42-2.33 (m,

1H).

#### Step 5: Preparation of 2-chloro-*N*-(2-(4-(2-chloroacetyl)-1-(prop-2-yn-1-yl)piperazine-2-carbonyl)-1,2,3,4-tetrahydroisoquinolin-6-yl)acetamide (SH-X-1A)

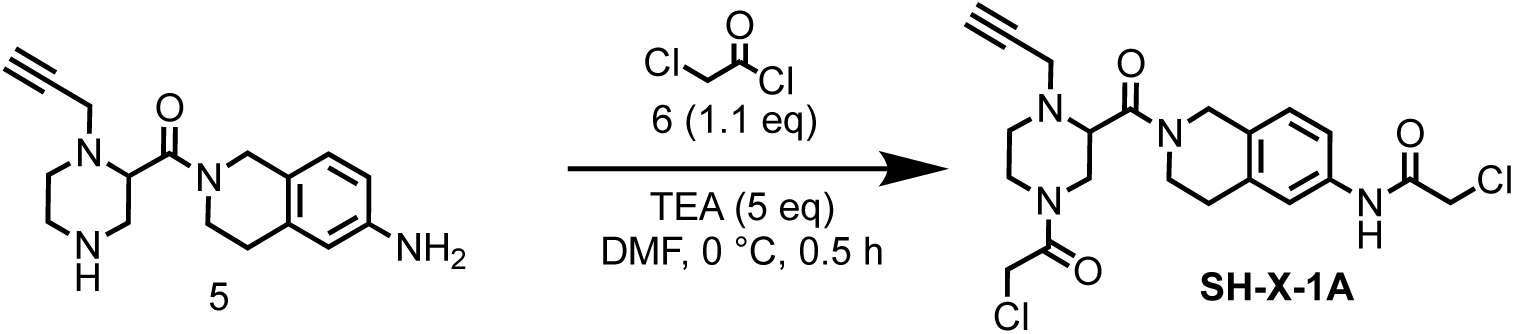

To a solution of (6-amino-3,4-dihydroisoquinolin-2(1H)-yl)(1-(prop-2-yn-1-yl)piperazin-2-yl)methanone (500 mg, 1.49 mmol) in MeCN (25 mL) was added TEA (755 mg, 7.47 mmol, 1.04 mL), then 2-chloroacetyl chloride (169 mg, 1.49 mmol, 119μL) in MeCN (10 mL) was added into the reaction mixture at 0 °C, the reaction mixture was stirred at 0 °C for 0.5 hr. LCMS showed the reaction was completed. The reaction mixture was filtered and the filtrate was concentrated to give a residue. The residue was purified by preparative-HPLC (column: Phenomenex luna C18 150*25mm* 10um; mobile phase: [water(FA)-MeCN]; gradient: 16%-46% B over 10 min) to give 2-chloro-*N*-(2-(4-(2-chloroacetyl)-1-(prop-2-yn-1-yl)piperazine-2-carbonyl)-1,2,3,4-tetrahydroisoquinolin-6-yl)acetamide (96.0 mg, 208 μmol, 13.9% yield, 97.8% purity) as a white solid.

LC-MS: 451.0 [M+H]^+^ / Ret time: 0.311 min / method: 5-95AB_0.8min.lcm

^1^H NMR (400 MHz, DMSO-*d*_6_): *δ* 10.25 (s, 1H), 7.54-7.27 (m, 2H), 7.14 (dd, *J* = 4.4, 8.0 Hz, 1H), 4.93-4.80 (m, 1H), 4.74-4.56 (m, 1H), 4.56-4.37 (m, 2H), 4.36-4.26 (m, 1H), 4.23 (s, 2H), 4.19-4.05 (m, 1H), 3.98-3.85 (m, 1H), 3.83-3.64 (m, 3H), 3.58-3.46 (m, 1H), 3.38 (s, 1H), 3.25 (d, *J* = 7.6 Hz, 1H), 3.05-2.81 (m, 4H), 2.73 (s, 1H)

### Synthesis of 4-(2-chloroacetyl)-1-(2-(2-chloroacetyl)-1,2,3,4-tetrahydroisoquinoline-6-carbonyl)-*N*-(1-(2-(2-(6-((4*R*,5*S*)-5-methylimidazolidin-4-yl)hexanamido)ethoxy)ethyl)-1*H*-1,2,3-triazol-4-yl)piperazine-2-carboxamide (SH-X-13)

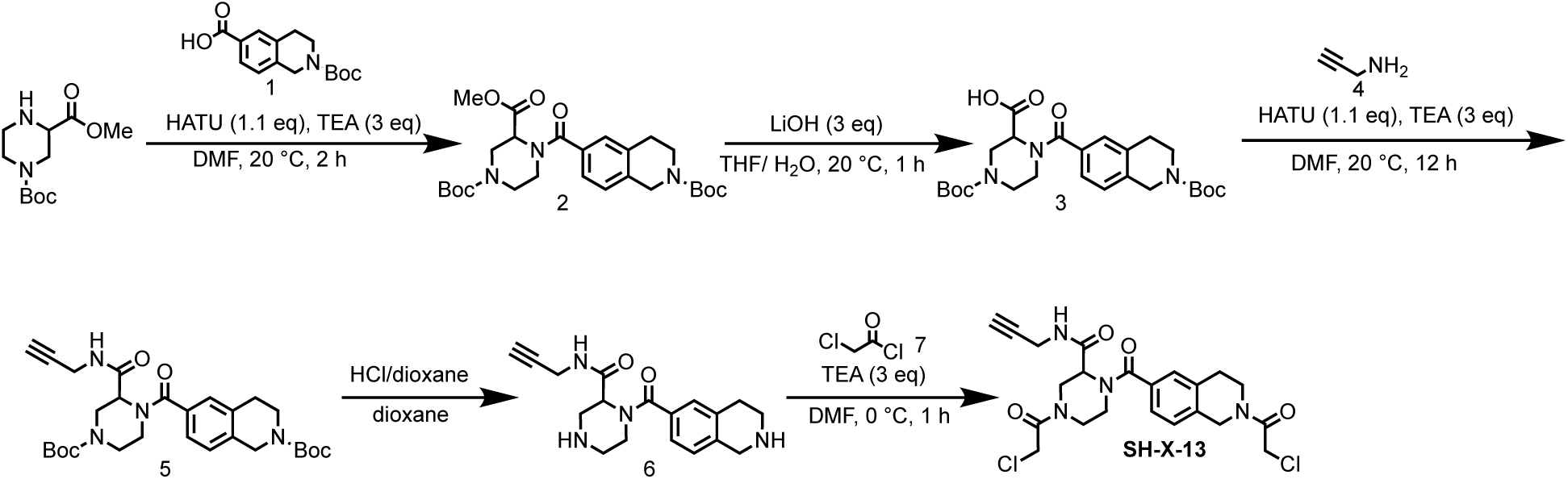

#### Step 1: Preparation of 1-(*tert*-butyl) 3-methyl 4-(2-(*tert*-butoxycarbonyl)-1,2,3,4-tetrahydroisoquinoline-6-carbonyl)piperazine-1,3-dicarboxylate (2)

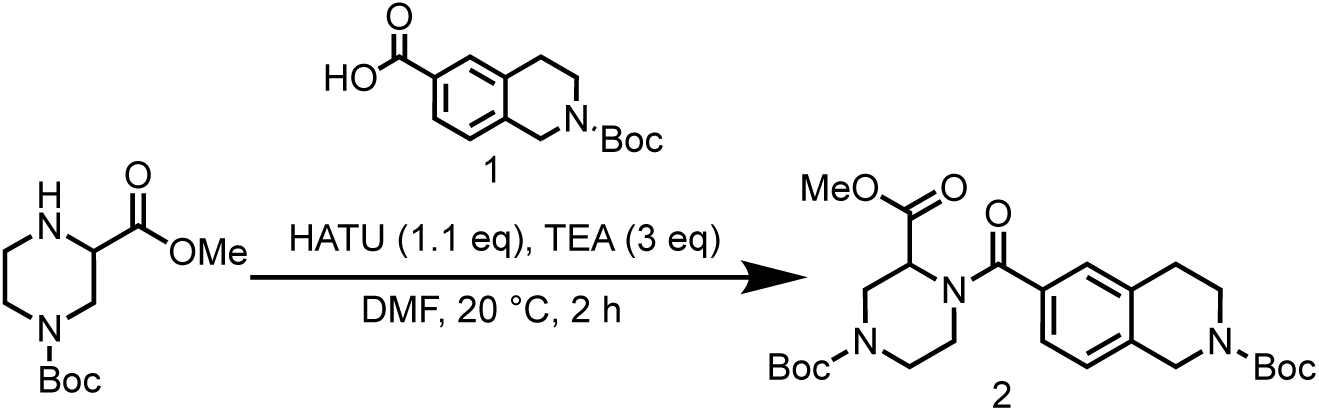

To a solution of 1-(*tert*-butyl) 3-methyl piperazine-1,3-dicarboxylate (1.00 g, 3.61 mmol) in DMF (10 mL) was added HATU (1.51 g, 3.97 mmol), TEA (1.09 g, 10.8 mmol, 1.51 mL) and 2-(*tert*-butoxycarbonyl)-1,2,3,4-tetrahydroisoquinoline-6-carboxylic acid (969 mg, 3.97 mmol). The mixture was stirred at 20 °C for 2 hrs. The reaction mixture was poured into H_2_O (150 mL), extracted with EtOAc (100 mL×3).

The organic layers were collected and washed with brine (200 mL×3), dried over Na_2_SO_4_ and concentrated to give a residue. The residue was purified by column chromatography on silica gel (eluted with petroleum ether: ethyl acetate=10:0 to 1:1) to give 1-(*tert*-butyl) 3-methyl 4-(2-(*tert*-butoxycarbonyl)-1,2,3,4-tetrahydroisoquinoline-6-carbonyl)piperazine-1,3-dicarboxylate (1.80 g, 3.57 mmol, 99.1% yield) as a light yellow oil.

#### Step 2: Preparation of 4-(*tert*-butoxycarbonyl)-1-(2-(*tert*-butoxycarbonyl)-1,2,3,4-tetrahydroisoquinoline-6-carbonyl)piperazine-2-carboxylic acid (3)

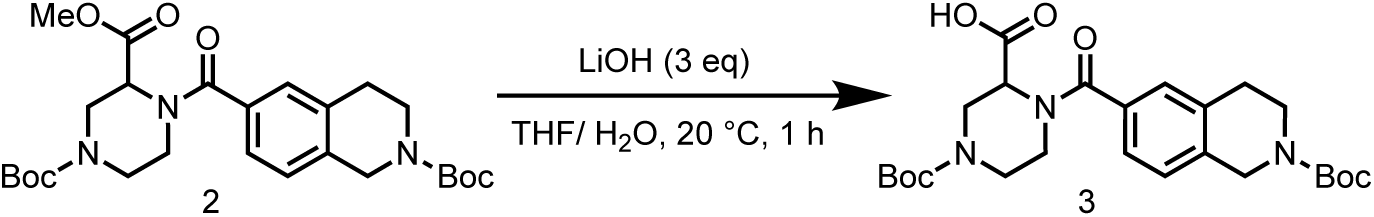

To a solution of 1-(*tert*-butyl) 3-methyl 4-(2-(*tert*-butoxycarbonyl)-1,2,3,4-tetrahydroisoquinoline-6-carbonyl)piperazine-1,3-dicarboxylate (1.80 g, 3.57 mmol) in THF (8 mL) and H_2_O (8 mL) was added LiOH (449 mg, 10.7 mmol). The mixture was stirred at 20 °C for 1 hr. LCMS showed the reaction was completed. The reaction mixture was acidified with HCl (1 M) to pH=3. Then concentrated to give 4-(*tert*-butoxycarbonyl)-1-(2-(*tert*-butoxycarbonyl)-1,2,3,4-tetrahydroisoquinoline-6-carbonyl)piperazine-2-carboxylic acid (1.64 g, 3.35 mmol, 93.7% yield) as a white solid.

LC-MS: 488.2 [M-H]^+^ / Ret time: 0.466 min / method: 5-95CD_1min_NEG.lcm

^1^H NMR (400 MHz, CDCl3) *δ* 7.35 (d, *J* = 5.2 Hz, 1H), 7.28-7.21 (m, 2H), 5.38 (s, 1H), 4.78 (d, *J* = 14.0 Hz, 1H), 4.67 (s, 2H), 4.23-3.93 (m, 1H), 3.73 (s, 2H), 3.23 (s, 1H), 2.97-2.87 (m, 3H), 1.58 (s, 9H), 1.52 (s, 9H).

#### Step 3: Preparation of *tert*-butyl 6-[4-*tert*-butoxycarbonyl-2-(prop-2-ynyl carbamoyl)piperazine-1-carbonyl]-3,4-dihydro-1*H*-isoquinoline-2-carboxylate (5)

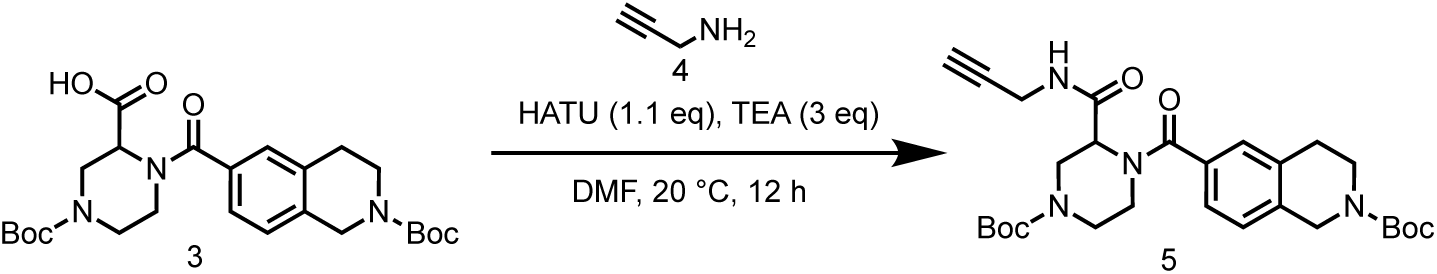

To a mixture of 4-(*tert*-butoxycarbonyl)-1-(2-(*tert*-butoxycarbonyl)-1,2,3,4-tetrahydroisoquinoline-6-carbonyl)piperazine-2-carboxylic acid (8.60 g, 17.57 mmol) and prop-2-yn-1-amine (1.16 g, 21.1 mmol, 1.35 mL) in DMF (15 mL) was added HATU (7.35 g, 19.3 mmol) and TEA (5.33 g, 52.7 mmol, 7.34 mL). The mixture was stirred at 20 °C for 12 hrs. LCMS showed the reaction was completed. The reaction mixture was poured into H_2_O (100 mL), extracted with EtOAc (200 mL×3). The organic layers were collected and washed with brine (200 mL×3), dried over Na_2_SO_4_ and concentrated to give a residue. The residue was purified by column chromatography on silica gel (eluted with petroleum ether: ethyl acetate=10:0 to 1:1) to give *tert*-butyl 6-(4-(*tert*-butoxycarbonyl)-2-(prop-2-yn-1-ylcarbamoyl)piperazine-1-carbonyl)-3,4-dihydroisoquinoline-2(1*H*)-carboxylate (6.00 g, 11.4 mmol, 64.8% yield) as a yellow oil.

LC-MS: 527.3 [M+H]^+^ / Ret time: 0.688 min / method: 5-95CD_1min.lcm

#### Step 4: Preparation of *N*-(prop-2-yn-1-yl)-1-(1,2,3,4-tetrahydroisoquinoline-6-carbonyl)piperazine-2-carboxamide (6)

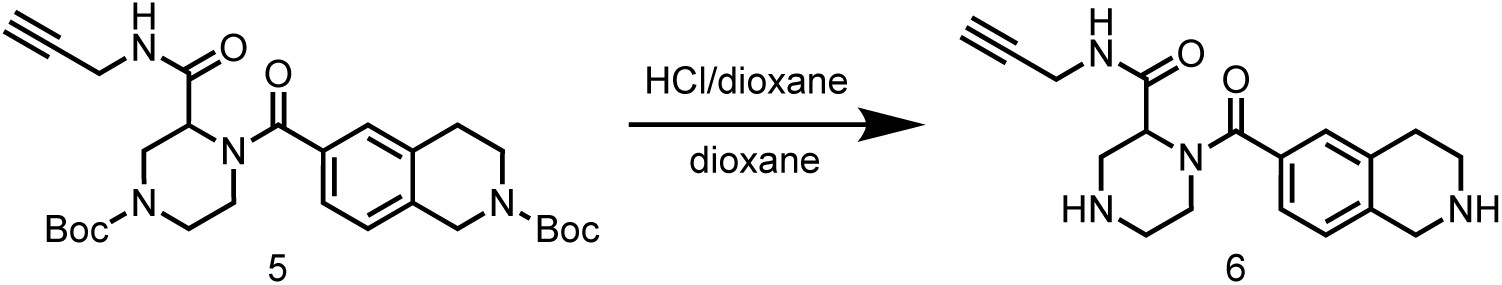

To a solution of *tert*-butyl 6-(4-(*tert*-butoxycarbonyl)-2-(prop-2-yn-1-ylcarbamoyl) piperazine-1-carbonyl)-3,4-dihydroisoquinoline-2(1*H*)-carboxylate (3.00 g, 5.70 mmol) in dioxane (4 mL) was added HCl/dioxane (4 M, 8 mL). The mixture was stirred at 20 °C for 1 hr. The reaction mixture was evaporated in vacuum to give *N*-(prop-2-yn-1-yl)-1-(1,2,3,4-tetrahydroisoquinoline-6-carbonyl)piperazine-2-carboxamide (2.00 g, 5.51 mmol, 96.7% yield, HCl salt) as a light yellow solid.

LC-MS: 327.1 [M+H]^+^ / Ret time: 0.323 min / 5-95CD_1min.lcm

#### Step 5: Preparation of 4-(2-chloroacetyl)-1-(2-(2-chloroacetyl)-1,2,3,4-tetrahydroisoquinoline-6-carbonyl)-*N*-(prop-2-yn-1-yl)piperazine-2-carboxamide (SH-X-13)

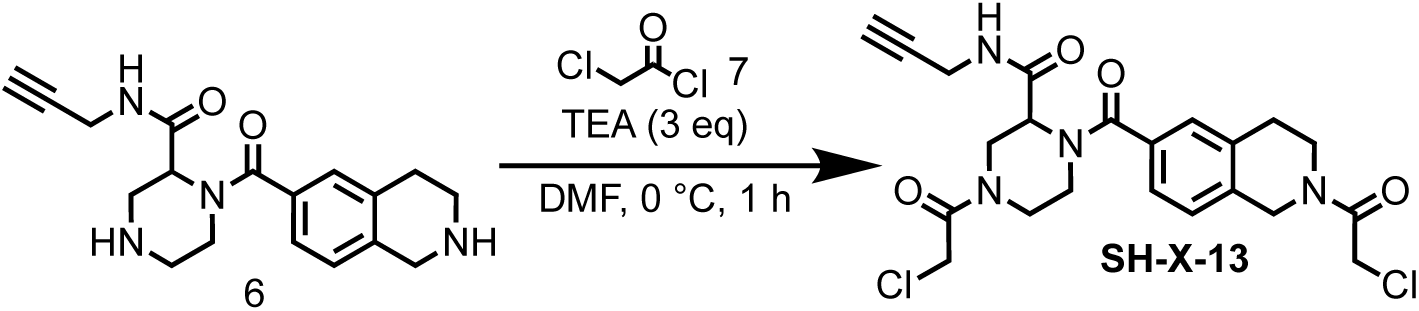

To a solution of *N*-prop-2-ynyl-1-(1,2,3,4-tetrahydroisoquinoline-6-carbonyl) piperazine-2-carboxamide (350 mg, 1.07 mmol) in DMF (3 mL) was added TEA (542 mg, 5.36 mmol, 746 μL) and 2-chloroacetyl chloride (68.5 mg, 606 μmol, 48.3 μL) in DMF (1 mL) at 0 °C. The mixture was stirred at 0 °C for 1 hr. The reaction mixture was filtered and the filtrate was concentrated to give a residue. The residue was purified by preparative-HPLC (0.1% FA; mobile phase: [water (FA)-MeCN]; gradient: 25%-45% B over 15 min condition). to give 4-(2-chloroacetyl)-1-(2-(2-chloroacetyl)-1,2,3,4-tetrahydroisoquinoline-6-carbonyl)-*N*-(prop-2-yn-1-yl)piperazine-2-carboxamide (25 mg, 50.1 μmol, 18.2% yield, 96% purity) as a white solid.

LC-MS: 479.1 [M+H]^+^ / Ret time: 0.325 min / 5-95AB_0.8min.lcm

^1^H NMR (400 MHz, DMSO-*d*_6_) *δ* 8.81 - 8.47 (m, 1H), 7.39 - 7.17 (m, 3H), 5.07 (s, 1H), 4.72 (s, 1H), 4.66 (s, 1H), 4.61 - 4.53 (m, 1H), 4.42 - 4.34 (m, 1H), 4.33 - 4.20 (m, 2H), 4.15 - 3.98 (m, 1H), 3.97 - 3.82 (m, 2H), 3.71 (s, 2H), 3.63 - 3.54 (m, 1H), 3.53 - 3.40 (m, 1H), 3.29 (s, 2H), 3.13 (s, 1H), 2.97 - 2.77 (m, 3H).

### Synthesis of 4-acryloyl-1-(2-acryloyl-1,2,3,4-tetrahydroisoquinoline-6-carbonyl)-*N*-(prop-2-yn-1-yl)piperazine-2-carboxamide (SH-X-14)

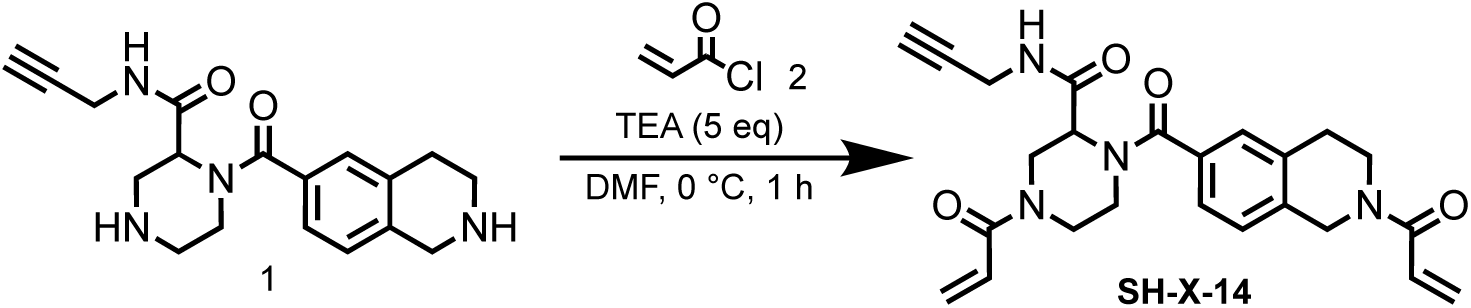

To a solution of *N*-prop-2-ynyl-1-(1,2,3,4-tetrahydroisoquinoline-6-carbonyl)piperazine-2-carboxamide (350 mg, 964 μmol, HCl salt) in DMF (2 mL) was added TEA (488 mg, 4.82 mmol, 671 μL) and prop-2-enoyl chloride (192 mg, 2.12 mmol, 172 μL) in DMF (0.5 mL) at 0 °C. The mixture was stirred at 0 °C for 1 hr. The reaction mixture was filtered and the filtrate was concentrated to give a residue. The residue was purified by preparative-HPLC (0.1% FA; mobile phase: [water(FA)-MeCN]; gradient: 13%-43% B over 15 min condition) to give 4-acryloyl-1-(2-acryloyl-1,2,3,4-tetrahydroisoquinoline-6-carbonyl)-*N*-(prop-2-yn-1-yl)piperazine-2-carboxamide (30.0 mg, 67.0 μmol, 6.94% yield, 97% purity) as a white solid.

LC-MS: 435.2 [M+H]^+^ / Ret time: 0.303 min / method /5-95AB_0.8min.lcm

^1^H NMR (400 MHz, DMSO-*d*_6_): *δ* 8.74-8.45 (m, 1H), 7.39-7.09 (m, 3H), 6.89 (m, *J* = 15.6 Hz, 1H), 6.81-6.58 (m, 1H), 6.20-6.02 (m, 2H), 5.80-5.62 (m, 2H), 5.09-4.87 (m, 1H), 4.82 (s, 1H), 4.71 (s, 1H), 4.64-4.26 (m, 2H), 4.24-4.08 (m, 1H), 3.93-3.73 (m, 4H), 3.65-3.44 (m, 2H), 3.13 (s, 1H), 2.91-2.81 (m, 2H), 2.63-2.52 (m, 1H).

### Synthesis of *N*,*N*’-(5-((4-ethynylbenzyl)oxy)-1,3-phenylene)bis(2-chloroacetamide) (SH-X-005)

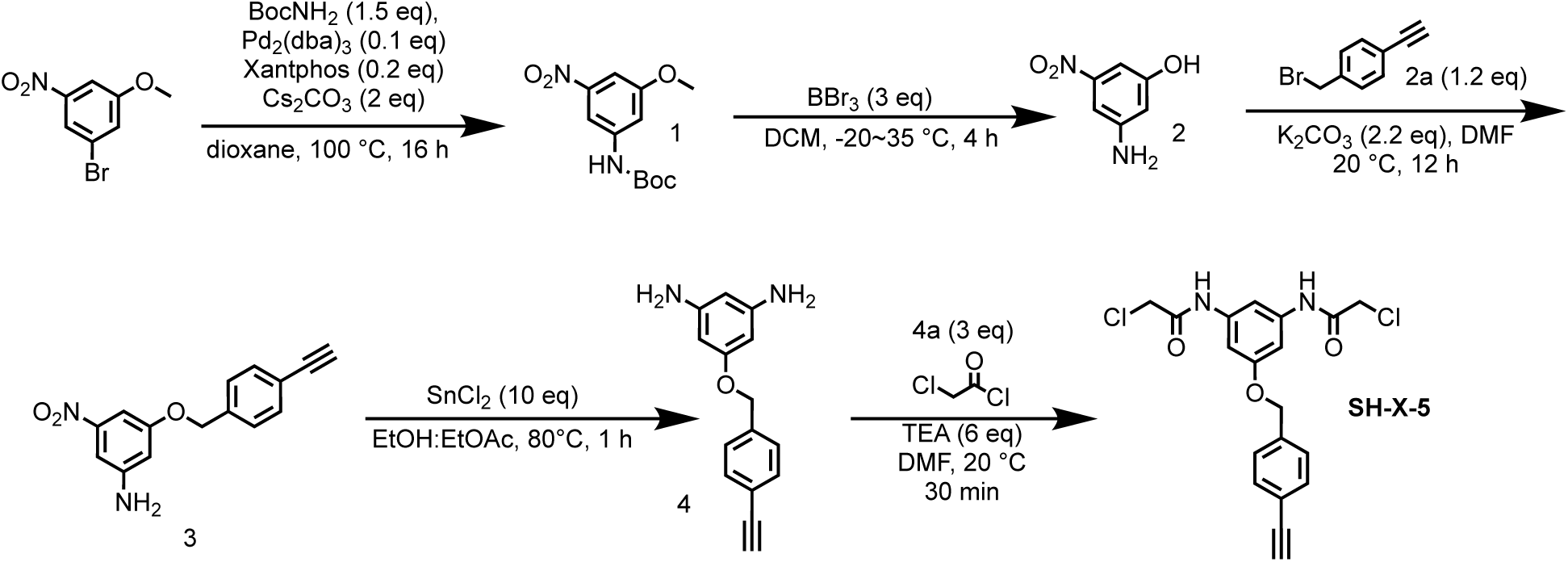

#### Step 1: Preparation of *tert*-butyl (3-methoxy-5-nitrophenyl)carbamate (1)

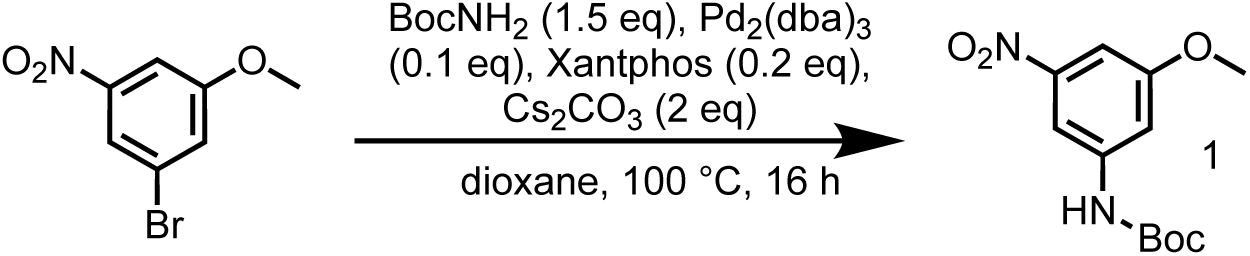

To a solution of 1-bromo-3-methoxy-5-nitrobenzene (3.79 g, 32.3 mmol), 1-bromo-3-methoxy-5-nitro-benzene (5.00 g, 21.5 mmol) and Cs_2_CO_3_ (14.0 g, 43.1 mmol) in dioxane (50 mL) was added Pd_2_(dba)_3_ (1.97 g, 2.15 mmol) and Xantphos (2.49 g, 4.31 mmol), the mixture was stirred at 100 °C for 12 hrs under N_2_. LCMS showed the reaction was completed. After the reaction was completed and cooled to room temperature, the mixture was filtered, the filtrate was concentrated to give a residue. The residue was purified by column chromatography on silica gel (eluted with petroleum ether: ethyl acetate=10:0 to 5:1) to give *tert*-butyl (3-methoxy-5-nitrophenyl)carbamate (6.30 g, crude) as yellow solid.

LC-MS: 213.2 [M+H-56]^+^ / Ret time: 0.440 min / method: 5-95AB_0.8min.lcm.

#### Step 2: Preparation of 3-amino-5-nitrophenol (2)

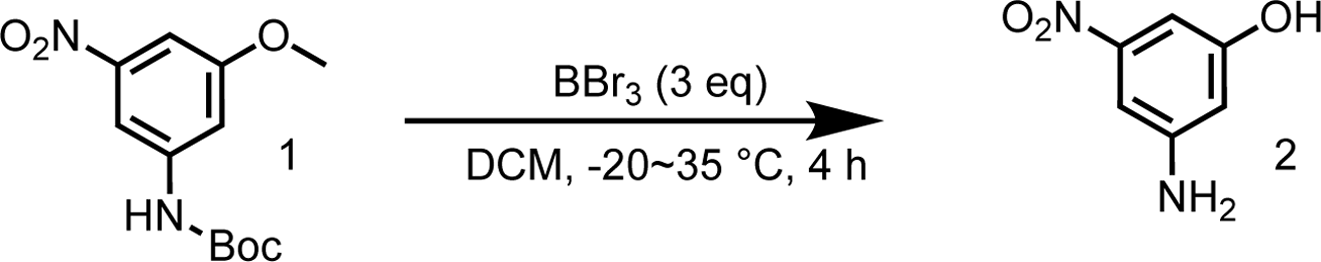

To a solution of *tert*-butyl (3-methoxy-5-nitrophenyl) carbamate (6.30 g, 23.4 mmol) in DCM (60 mL) in a 500 ml three neck round bottle was added BBr_3_ (17.6 g, 70.4 mmol, 6.79 mL) at −20 °C under N_2_, the mixture was stirred at 35 °C for 4 hrs under N_2_. LCMS showed the reaction was completed. MeOH (200 mL) was added slowly to the mixture at 20 °C. Then the mixture was stirred at 20 °C for 0.5 hr, and concentrated to give a residue. The residue was purified by column chromatography on silica gel (eluted with petroleum ether: ethyl acetate=5:1 to 3:1) to give 3-amino-5-nitrophenol (3.28 g, 21.2 mmol, 90.6% yield) as yellow solid.

LC-MS: 155.1 [M+H]^+^ / method: 5-95AB_0.8min.lcm

^1^H NMR (400 MHz, DMSO-*d*_6_): *δ* 9.83 (s, 1H), 6.88 (t, *J* = 2.0 Hz, 1H), 6.71 (t, *J* = 2.0 Hz, 1H), 6.37 (t, *J* = 2.0 Hz, 1H), 5.68 (s, 2H).

#### Step 3: Preparation of 3-((4-ethynylbenzyl)oxy)-5-nitroaniline (3)

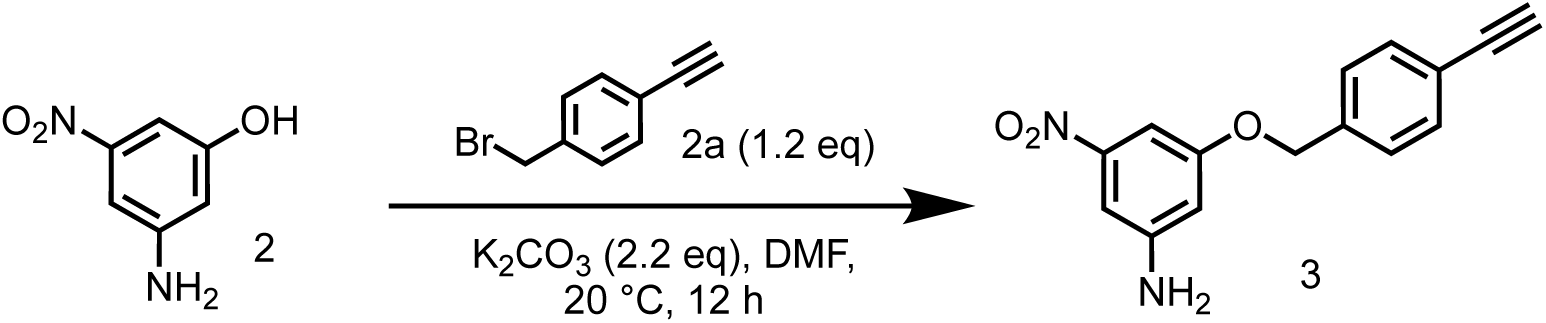

A solution of 1-(bromomethyl)-4-ethynylbenzene (4.99 g, 25.5 mmol) in DMF (10 mL) was added into a mixture of and 3-amino-5-nitrophenol (3.28 g, 21.3 mmol) and K2CO3 (6.48 g, 46.8 mmol) in DMF (50 mL), the mixture was stirred at 20 °C for 12 hrs under N2. LCMS showed the reaction was completed. The mixture was poured into water (200 mL), extracted with EtOAc (200 mL×3). The organic layer was collected and washed with brine (200 mL×3), dried with Na_2_SO_4_ and concentrated to give a residue. The residue was purified by column chromatography on silica gel (eluted with petroleum ether: ethyl acetate=10:1 to 5:1) to give 3-((4-ethynylbenzyl)oxy)-5-nitroaniline (1.80 g, 6.71 mmol, 31.5% yield) as a yellow solid.

LC-MS: 269.0 [M+H]^+^ / Ret time: 0.633 min / method: 5-95CD_1min.lcm

#### Step 4: Preparation of 5-((4-ethynylbenzyl)oxy)benzene-1,3-diamine (4)

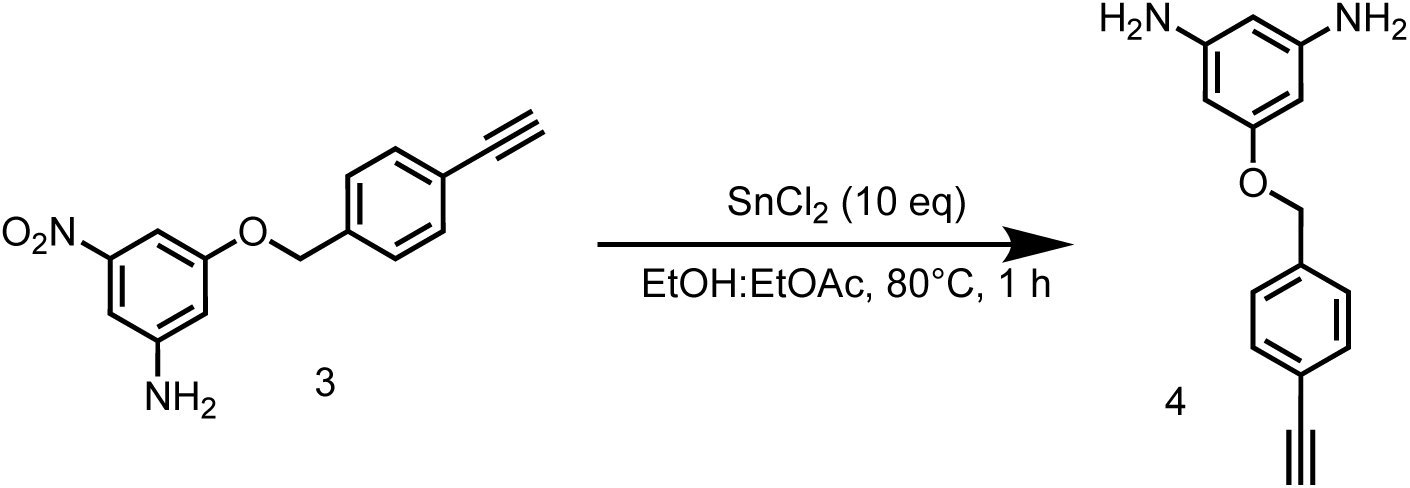

To a solution of 3-((4-ethynylbenzyl)oxy)-5-nitroaniline (1.80 g, 6.71 mmol) in EtOH (18 mL) and EtOAc (6 mL) was added SnCl_2_•2H_2_O (15.1 g, 67.1 mmol), the mixture was stirred at 80 °C for 1 hr. LCMS showed the reaction was completed. The reaction mixture was cooled and quenched with saturated NaHCO_3_ (aq), extracted with EtOAc (200 mL×3). The organic layer was collected and washed with brine (200 mL×3), dried over Na_2_SO_4_ and concentrated to give 5-((4-ethynylbenzyl)oxy)benzene-1,3-diamine (1.60 g, crude) as a yellow solid.

LC-MS: 239.1 [M+H]^+^ / Ret time: 0.520 min / method: 5-95CD_1min.lcm

^1^H NMR (400 MHz, DMSO-*d*_6_): *δ* 7.51-7.45 (m, 2H), 7.43-7.36 (m, 2H), 5.53-5.40 (m, 3H), 4.93 (s, 2H), 4.68 (s, 3H), 4.17 (s, 1H).

#### Step 5: Preparation of *N*,*N*’-(5-((4-ethynylbenzyl)oxy)-1,3-phenylene)bis(2-chloroacetamide) (SH-X-5)

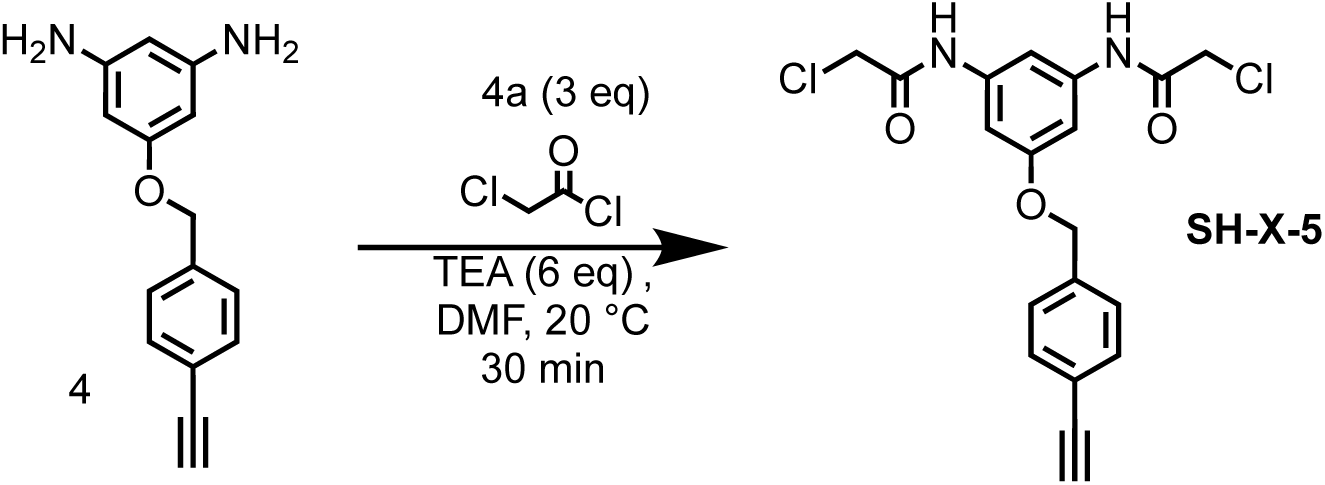

To a solution of 5-((4-ethynylbenzyl)oxy)benzene-1,3-diamine (300 mg, 1.26 mmol) in MeCN (8 mL) was added TEA (764 mg, 7.55 mmol), then 2-chloroacetyl chloride (312 mg, 2.77 mmol, 221 μL) was added into the reaction mixture was stirred at 20 °C for 0.5 hr. LCMS showed the reaction was completed. The reaction mixture was extracted with H_2_O (5 mL) and ethyl acetate (5 mL×3), washed with brine (5 mL), the organic phase was dried over Na_2_SO_4_, concentrated in vacuum to give a residue. The residue was purified by preparative-TLC (petroleum ether: ethyl acetate=1:1) to give *N*,*N*’-(5-((4-ethynylbenzyl)oxy)-1,3-phenylene)bis(2-chloroacetamide) (180 mg, 451 μmol, 35.8% yield, 98.0% purity) as a brown solid.

LC-MS: 391.2 [M+H]^+^ / Ret time: 0.421 min / method: 5-95AB_0.8min.lcm

^1^H NMR (400 MHz, DMSO-*d*_6_): *δ* 10.32 (s, 2H), 7.60-7.38 (m, 5H), 7.09 (d, *J* = 0.8 Hz, 2H), 5.10 (s, 2H), 4.24 (s, 4H), 4.20 (s, 1H).

### Synthesis of *N*,*N*’-(5-((4-ethynylbenzyl)oxy)-1,3-phenylene)diacrylamide (SH-X-6)

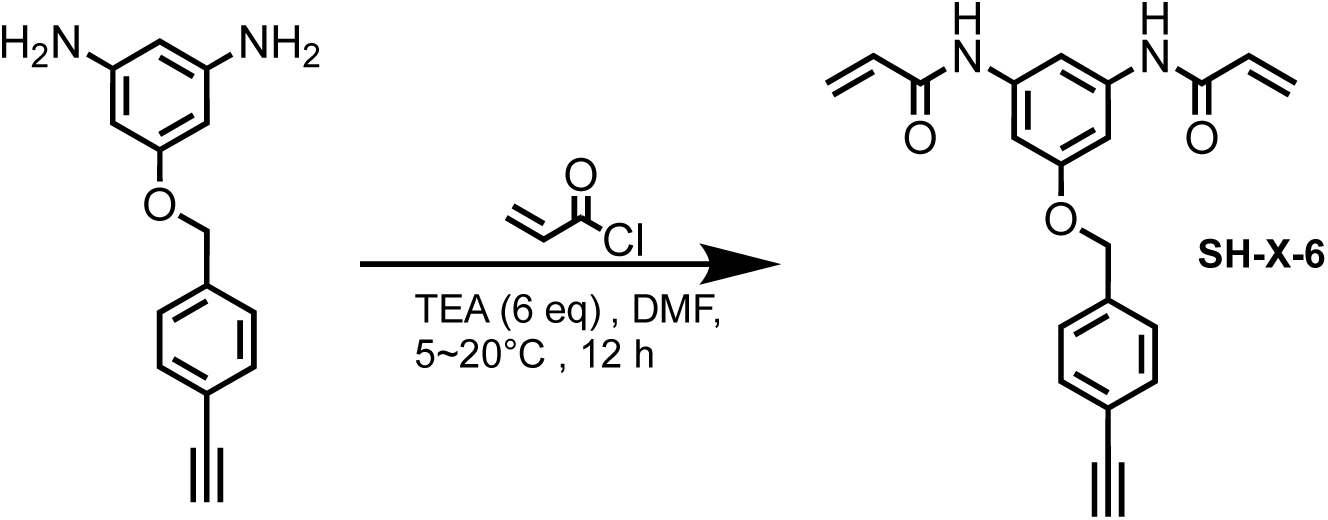

To a solution of 5-((4-ethynylbenzyl)oxy)benzene-1,3-diamine (300 mg, 1.26 mmol) in MeCN (8 mL) was added TEA (764 mg, 7.55 mmol), then 2-chloroacetyl chloride (312 mg, 2.77 mmol, 221 μL) was added into the reaction mixture was stirred at 20 °C for 0.5 hr. LCMS showed the reaction was completed. The reaction mixture was extracted with H_2_O (5 mL) and ethyl acetate (5 mL×3), washed with brine (5 mL), the organic phase was dried over Na_2_SO_4_, concentrated on vacuum to give a residue. The residue was purified by preparative-TLC (petroleum ether: ethyl acetate=1:1) to give *N,N*’-(5-((4-ethynylbenzyl) oxy)-1,3-phenylene)diacrylamide (170 mg, 481 μmol, 38.2% yield, 98.0% purity) as a brown solid.

LC-MS: 347.2 [M+H]^+^ / Ret time: 0.421 min / method: 5-95AB_0.8min.lcm

^1^H NMR (400 MHz, DMSO-*d*_6_): *δ* 10.16 (s, 2H), 7.63 (s, 1H), 7.59-7.40 (m, 4H), 7.19 (s, 2H), 6.51-6.38 (m, 2H), 6.32-6.20 (m, 2H), 5.82-5.69 (m, 2H), 5.09 (s, 2H), 4.21 (s, 1H).

### Synthesis of methyl 1-(3-(2-chloroacetamido)-4-(prop-2-yn-1-yloxy) benzoyl)-4-(2-chloroacetyl)piperazine-2-carboxylate (SH-X-026)

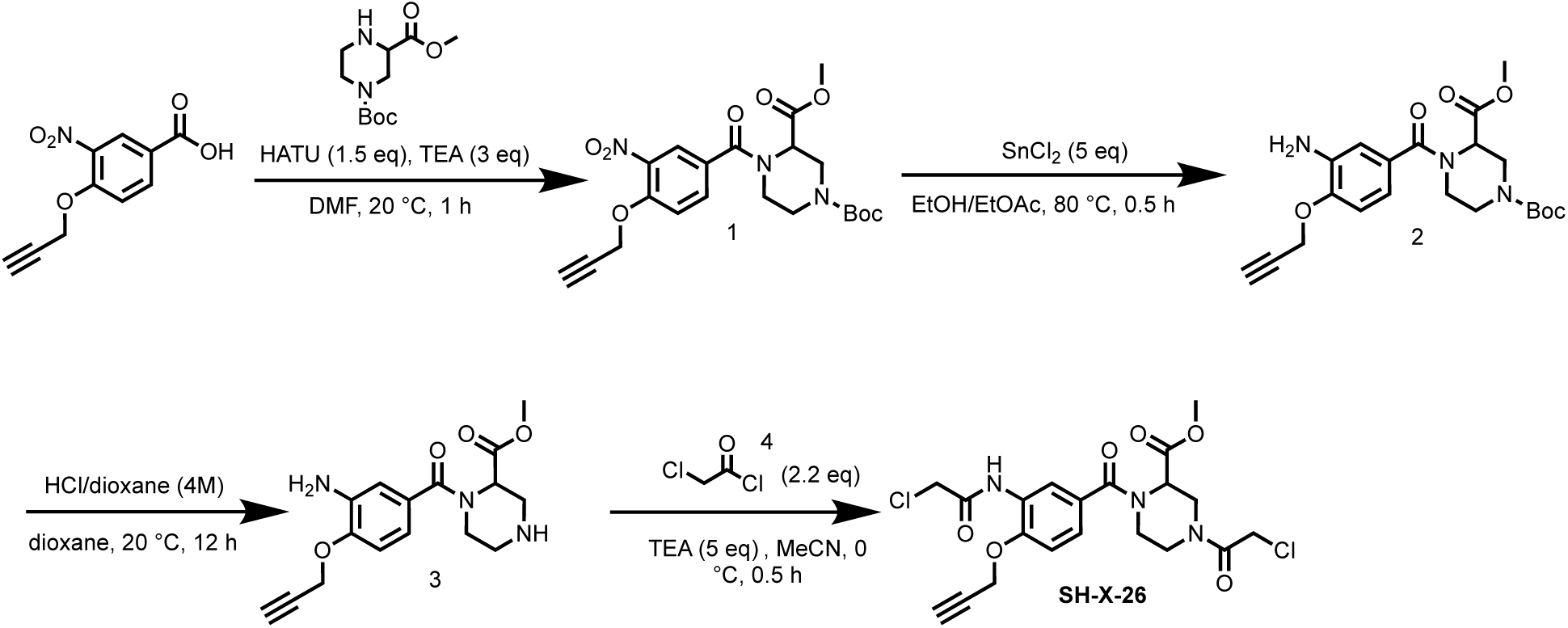

#### Step 1: Preparation of 1-(*tert*-butyl) 3-methyl 4-(3-nitro-4-(prop-2-yn-1-yloxy)benzoyl)piperazine-1,3-dicarboxylate (1)

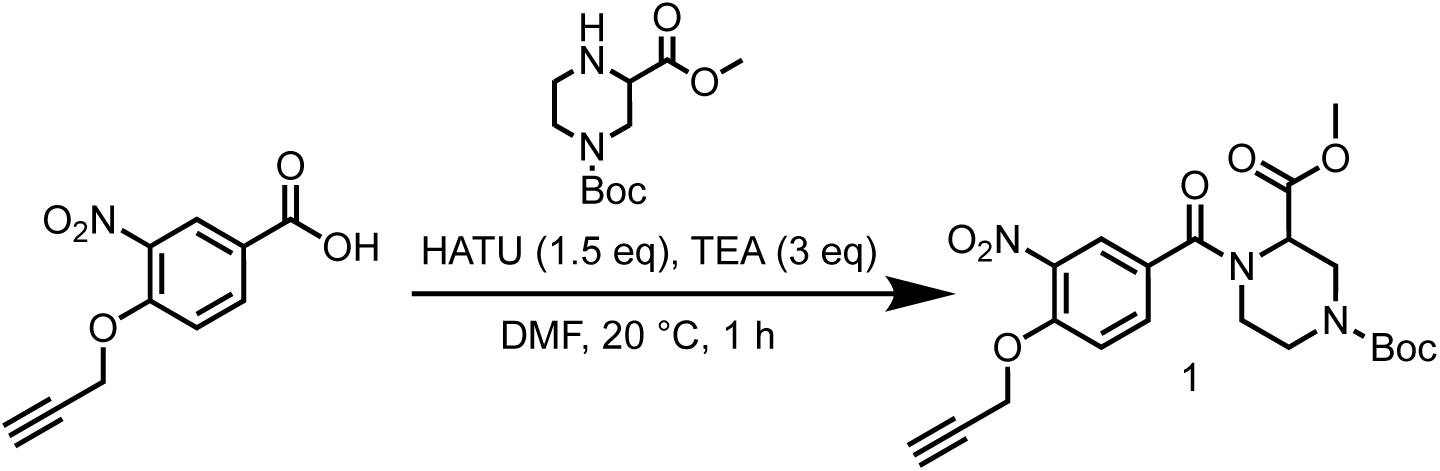

To a solution of 3-nitro-4-(prop-2-yn-1-yloxy)benzoic acid (2.00 g, 9.04 mmol) in DMF (20 mL) was added HATU (3.78 g, 9.95 mmol) and TEA (2.75 g, 27.13 mmol, 3.78 mL) and 1-(*tert*-butyl) 3-methyl piperazine-1,3-dicarboxylate (2.43 g, 9.95 mmol). The mixture was stirred at 20 °C for 1 hr. LCMS showed the reaction was completed. The reaction mixture was poured into H_2_O (100 mL), extracted with EtOAc (100 mL×3). The organic layers were collected and washed with brine (100 mL×3), dried over Na_2_SO_4_ and concentrated to give a residue. The residue was purified by column chromatography on silica gel (eluted with petroleum ether: ethyl acetate=1:1) to give 1-(*tert*-butyl) 3-methyl 4-(3-nitro-4-(prop-2-yn-1-yloxy)benzoyl)piperazine-1,3-dicarboxylate (4.00 g, 8.94 mmol, 98.9% yield) as a yellow solid.

LC-MS: 392.2 [M+H-56]^+^ / Ret time: 0.446 min / method: 5-95AB_0.8min.lcm

^1^H NMR (400 MHz, DMSO-*d*_6_): *δ* 7.96 (s, 1H), 7.80-7.63 (m, 1H), 7.54-7.43 (m, 1H), 5.09 (s, 2H), 4.39 (d, *J* = 13.6 Hz, 1H), 3.76-3.63 (m, 4H), 3.34-3.29 (m, 2H), 3.08-2.80 (m, 1H), 2.69 (s, 3H), 1.38 (s, 9H).

#### Step 2: Preparation of 1-(*tert*-butyl) 3-methyl 4-(3-amino-4-(prop-2-yn-1-yloxy) benzoyl) piperazine-1,3-dicarboxylate (2)

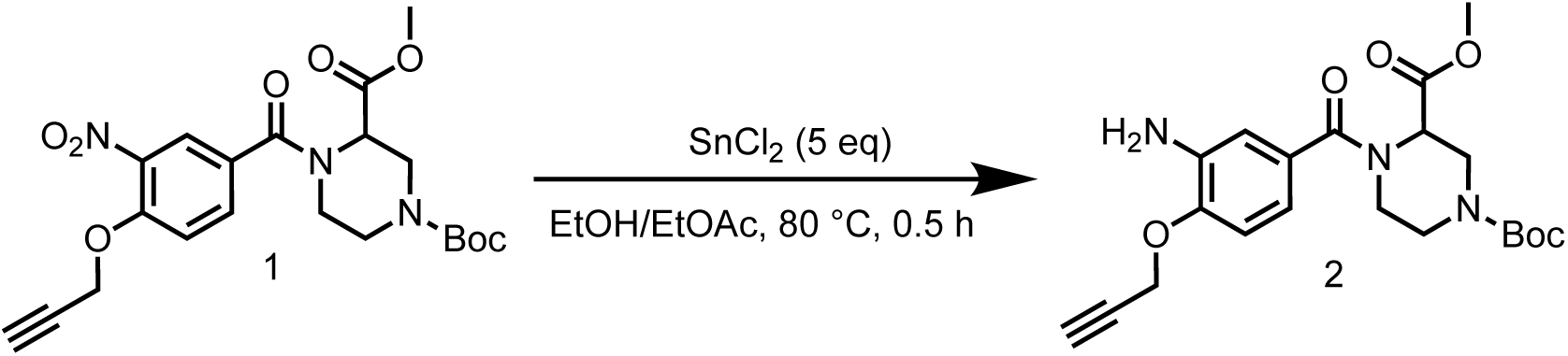

To a solution of 1-(*tert*-butyl) 3-methyl 4-(3-nitro-4-(prop-2-yn-1-yloxy)benzoyl)piperazine-1,3-dicarboxylate (3.00 g, 6.70 mmol) in EtOH (30 mL) and EtOAc (15 mL) was added SnCl_2_•2H_2_O (7.56 g, 33.5 mmol), the mixture was stirred at 80 °C for 0.5 hr. LCMS showed the reaction was completed. The reaction mixture was cooled and quenched with saturated NaHCO_3_(aq). The mixture was filtered and the filtrate was washed with EtOAc (100 mL×3). The organic layer of the filtrate was collected, dried and concentrated to give a residue. The residue was purified by column chromatography on silica gel (eluted with petroleum ether: ethyl acetate=1:2) to give 1-(*tert*-butyl) 3-methyl 4-(3-amino-4-(prop-2-yn-1-yloxy)benzoyl)piperazine-1,3-dicarboxylate (1.10 g, 2.64 mmol, 39.3% yield) was obtained as a yellow oil.

LC-MS: 362.1 [M+H-56]^+^ / Ret time: 0.401 min / method: 5-95AB_0.8min.lcm

#### Step 3: Preparation of methyl 1-(3-amino-4-(prop-2-yn-1-yloxy)benzoyl)piperazine-2-carboxylate (3)

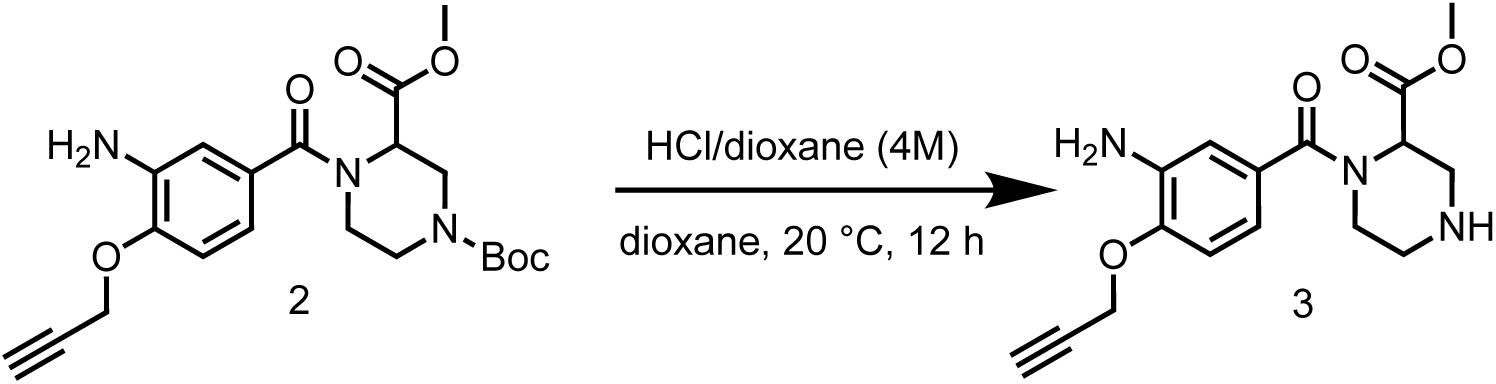

To a solution of 1-(*tert*-butyl) 3-methyl 4-(3-amino-4-(prop-2-yn-1-yloxy)benzoyl)piperazine-1,3-dicarboxylate (1.78 g, 4.26 mmol) in dioxane (10 mL) was added HCl/dioxane (4 M, 10 mL). The mixture was stirred at 20 °C for 12 hrs. LCMS showed the reaction was completed. The reaction mixture was filtered and the filter cake was concentrated to give a residue. The residue was purified by preparative-HPLC (column: Phenomenex luna C18 150*25mm* 10um;mobile phase: [water(FA)-MeCN]; gradient: 10%-40% B over 10 min) to give methyl 1-(3-amino-4-(prop-2-yn-1-yloxy) benzoyl)piperazine-2-carboxylate (30.9 mg, 95.4 μmol, 2.24% yield, 98.0% purity) was obtained as an off-white solid.

LC-MS: 318.1 [M+H]^+^ / Ret time: 0.143 min / method: 5-95AB_0.8min.lcm

^1^H NMR (400 MHz, DMSO-*d*_6_): *δ* 10.27-9.90 (m, 1H), 9.76-9.38 (m, 1H), 7.40 (s, 1H), 7.33-7.14 (m, 2H), 5.57-5.22 (m, 1H), 4.97 (d, *J* = 1.6 Hz, 2H), 4.58-4.20 (m, 1H), 3.82-3.73 (m, 3H), 3.65 (s, 2H), 3.56-2.97 (m, 4H).

#### Step 4: Preparation of methyl 1-(3-(2-chloroacetamido)-4-(prop-2-yn-1-yloxy) benzoyl)-4-(2-chloroacetyl)piperazine-2-carboxylate (SH-X-026)

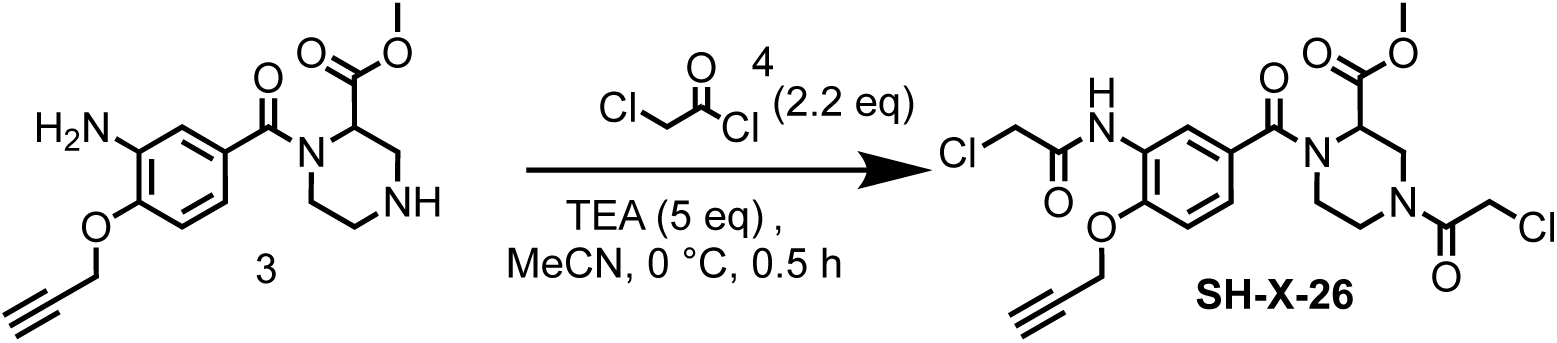

To a solution of methyl 1-(3-amino-4-(prop-2-yn-1-yloxy)benzoyl)piperazine-2-carboxylate (500 mg, 1.45 mmol) in MeCN (20 mL)was added TEA (733 mg, 7.25 mmol, 1.01 mL), then 2-chloroacetyl chloride (360 mg, 3.19 mmol, 254 μL) in MeCN (10 mL) was added into the reaction mixture at 0 °C, the reaction mixture was stirred at 0 °C for 0.5 hr. LCMS showed the reaction was completed. The reaction mixture was filtered and the filtrate was concentrated to give a residue. The residue was purified by preparative-HPLC (column: Phenomenex luna C18 150*25mm* 10um; mobile phase: [water(FA)-MeCN]; gradient: 25%-55% B over 10 min) to give methyl 1-(3-(2-chloroacetamido)-4-(prop-2-yn-1-yloxy)benzoyl)-4-(2-chloroacetyl)piperazine-2-carboxylate (85.3 mg, 182 μmol, 12.5% yield, 99.9% purity) as a white solid.

LC-MS: 470.1 [M+H]^+^ / Ret time: 0.381 min / method: 5-95AB_0.8min.lcm

^1^H NMR (400 MHz, DMSO-*d*_6_): *δ* 9.68 (s, 1H), 8.14 (s, 1H), 7.24 (s, 2H), 5.30-5.07 (m, 1H), 5.02-4.93 (m, 2H), 4.75-4.02 (m, 7H), 3.83-3.51 (m, 6H), 3.28-3.16 (m, 1H).

### Synthesis of methyl 1-(3-acrylamido-4-(prop-2-yn-1-yloxy)benzoyl)-4-acryloylpiperazine-2-carboxylate (SH-X-028)

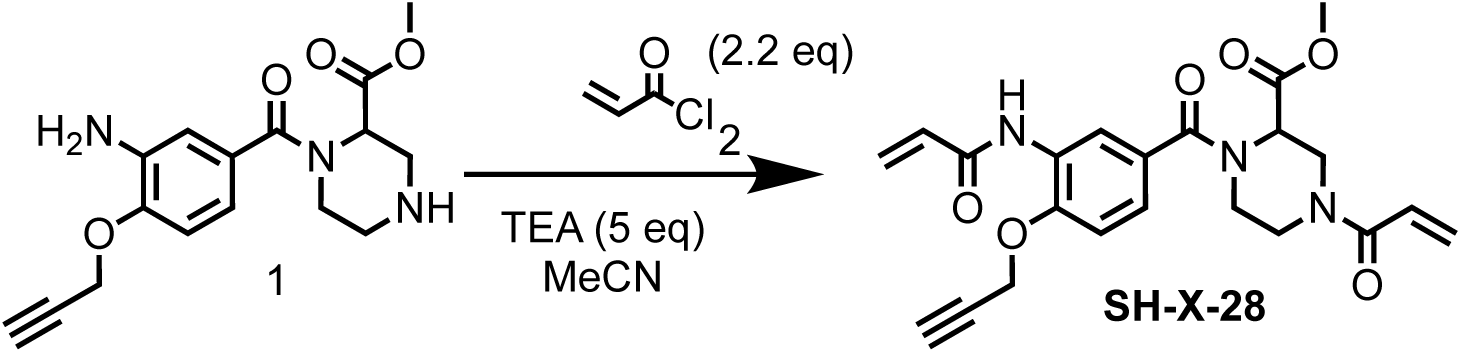

To a solution of methyl 1-(3-amino-4-(prop-2-yn-1-yloxy)benzoyl)piperazine-2-carboxylate (500 mg, 1.45 mmol) in MeCN (20 mL) was added TEA (734 mg, 7.25 mmol, 1.01 mL), then acryloyl chloride (289 mg, 3.19 mmol, 259 μL) in MeCN (10 mL) was added into the reaction mixture at 0 °C, the reaction mixture was stirred at 0 °C for 0.5 hr. LCMS showed the reaction was completed. The reaction mixture was filtered and the filtrate was concentrated to give a residue. The residue was purified by preparative-HPLC (column: Phenomenex luna C18 150*25mm* 10um; mobile phase: [water(FA)-MeCN]; gradient: 18%-48% B over 10 min) to give methyl 1-(3-acrylamido-4-(prop-2-yn-1-yloxy)benzoyl)-4-acryloylpiperazine-2-carboxylate (87.7 mg, 200 μmol, 13.8% yield, 97.1% purity) as a white solid.

LC-MS: 426.2 [M+H]^+^ / Ret time: 0.348 min / method: 5-95AB_0.8min.lcm

^1^H NMR (400 MHz, DMSO-*d*_6_): *δ* 9.56 (s, 1H), 8.22 (s, 1H), 7.23 (s, 2H), 6.72 (dd, *J* = 10.0, 16.4 Hz, 2H), 6.26 (d, *J* = 16.8 Hz, 1H), 6.12 (d, *J* = 16.8 Hz, 1H), 5.78-5.68 (m, 2H), 5.28-5.06 (m, 1H), 4.97 (s, 2H), 4.86-4.60 (m, 1H), 4.56-3.91 (m, 2H), 3.78-3.52 (m, 5H), 3.30-3.15 (m, 1H), 2.99-2.72 (m, 1H).

### Synthesis of 2-chloro-1-(5-(4-(2-chloroacetyl)-1-(prop-2-yn-1-yl)piperazine-2-carbonyl)-2,5-diazabicyclo[2.2.1]heptan-2-yl)ethan-1-one (SH-X-34A)

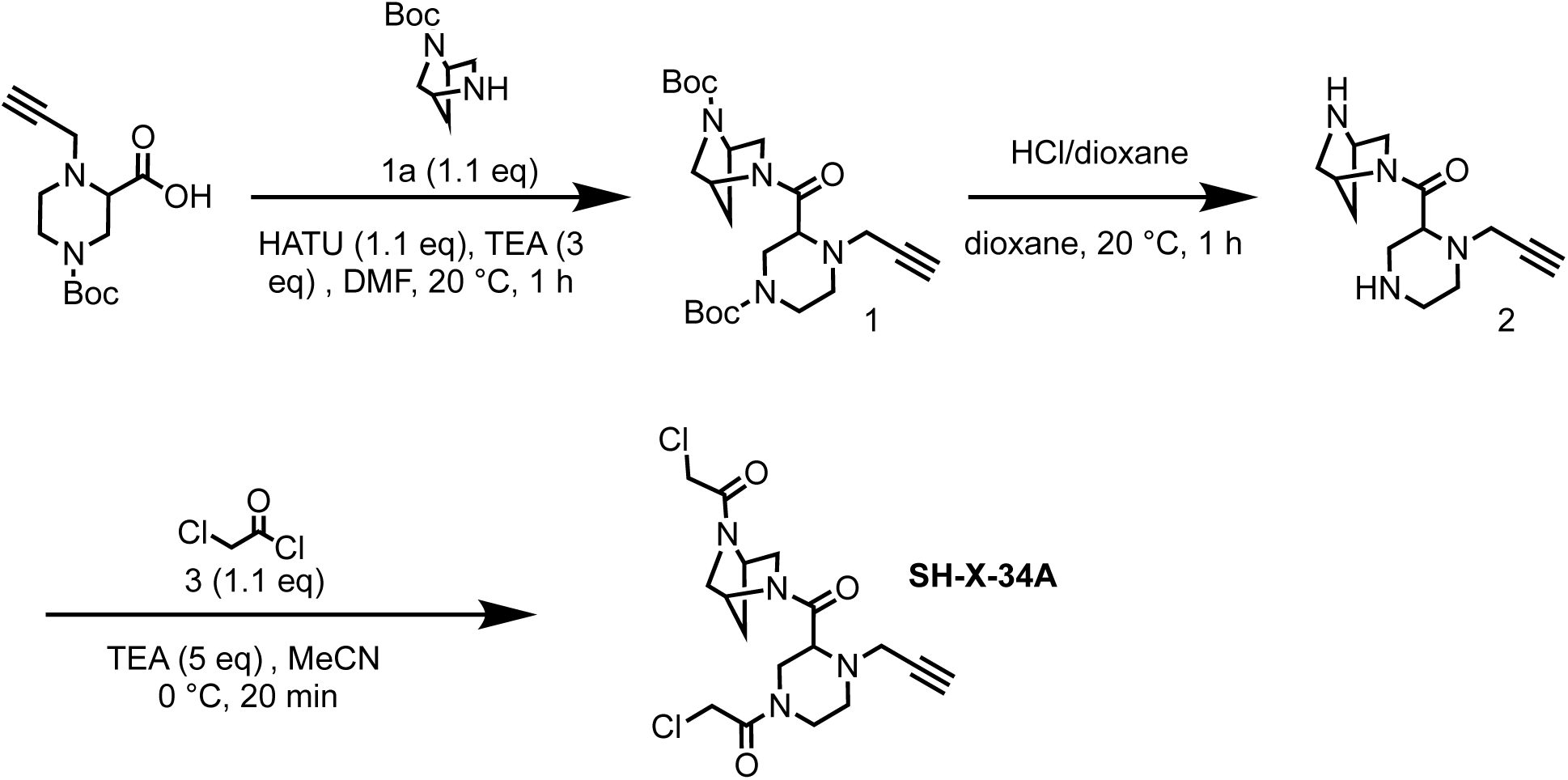

#### Step 1: Preparation of *tert*-butyl-5-(4-(*tert*-butoxycarbonyl)-1-(prop-2-yn-1-yl)piperazine-2-carbonyl)-2,5-diazabicyclo[2.2.1]heptane-2-carboxylate (1)

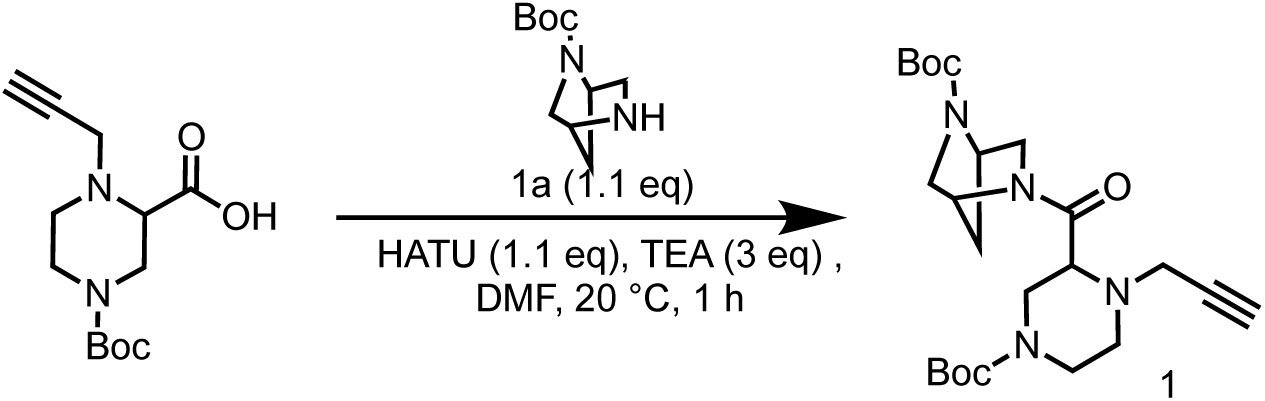

To a solution of 4-(*tert*-butoxycarbonyl)-1-(prop-2-yn-1-yl)piperazine-2-carboxylic acid (1.20 g, 4.47 mmol) in DMF (10 mL) was added HATU (1.87 g, 4.92 mmol) and TEA (1.36 g, 13.4 mmol, 1.87 mL) and *tert*-butyl 2,5-diazabicyclo[2.2.1]heptane-2-carboxylate (975 mg, 4.92 mmol). The mixture was stirred at 20 °C for 1 hr. LCMS showed the reaction was completed. The reaction mixture was poured into H_2_O (30 mL), extracted with EtOAc (30 mL×3). The organic layers were collected and washed with brine (30 mL×3), dried over Na_2_SO_4_ and concentrated to give a residue. The residue was purified by column chromatography on silica gel (eluted with petroleum ether: ethyl acetate=1:3) to give *tert*-butyl-5-(4-(*tert*-butoxycarbonyl)-1-(prop-2-yn-1-yl)piperazine-2-carbonyl)-2,5-diazabicyclo[2.2.1]heptane-2-carboxylate (2.00 g, 3.92 mmol, 87.7% yield, 88.0% purity) as a yellow oil.

LC-MS: 449.2 [M+H]^+^ / Ret time: 0.436 min / method: 5-95AB_0.8min.lcm

^1^H NMR (400 MHz, DMSO-*d*_6_): *δ* 4.99-4.83 (m, 1H), 4.78-4.64 (m, 1H), 4.39 (d, *J* = 13.6 Hz, 1H), 3.77-3.63 (m, 4H), 3.37 (s, 2H), 3.31-3.29 (m, 2H), 3.26 (d, *J* = 2.4 Hz, 2H), 3.19 (t, *J* = 2.0 Hz, 1H), 3.15-3.09 (m, 2H), 1.86-1.78 (m, 2H), 1.39 (s, 18H).

#### Step 2: Preparation of (2,5-diazabicyclo[2.2.1]heptan-2-yl)(1-(prop-2-yn-1-yl)piperazin-2-yl)methanone (2)

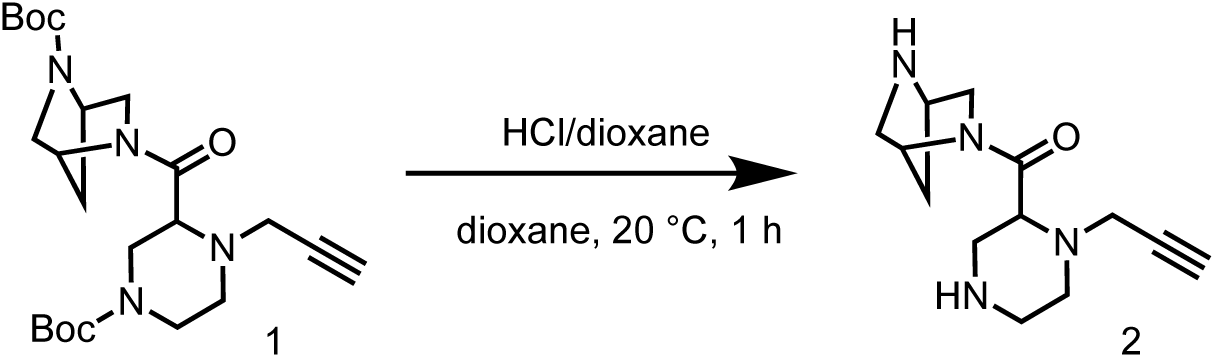

To a solution of *tert*-butyl 5-(4-(*tert*-butoxycarbonyl)-1-(prop-2-yn-1-yl)piperazine-2-carbonyl)-2,5-diazabicyclo[2.2.1]heptane-2-carboxylate (2.00 g, 3.92 mmol) in dioxane (5 mL) was added HCl/dioxane (4 M, 13.2 mL). The mixture was stirred at 20 °C or 1 hr. LCMS showed the reaction was completed. The reaction mixture was concentrated in vacuum to remove solvent, then triturated with DCM. The reaction mixture was filtered and the filter cake was concentrated to give (2,5-diazabicyclo[2.2.1]heptan-2-yl)(1-(prop-2-yn-1-yl)piperazin-2-yl)methanone (1.20 g, 3.86 mmol, 98.3% yield, 91.5% purity, HCl salt) as a white solid.

LC-MS: 249.2 [M+H]^+^ / Ret time: 0.432 min / method: 0-60N_1min.lcm

^1^H NMR (400 MHz, MeOD-*d*_4_): *δ* 5.18-4.80 (m, 1H), 4.69 (d, *J* = 16.4 Hz, 1H), 4.30 (d, *J* = 12.0 Hz, 1H), 4.23 (s, 1H), 3.85 (d, *J* = 18.4 Hz, 1H), 3.76 (s, 1H), 3.61 (d, *J* = 10.8 Hz, 1H), 3.44-3.34 (m, 2H), 3.28-3.22 (m, 1H), 3.21-3.12 (m, 2H), 3.11-3.00 (m, 2H), 2.96 (s, 1H), 2.68-2.52 (m, 1H), 2.01-1.70 (m, 2H).

#### Step 3: Preparation of 2-chloro-1-(5-(4-(2-chloroacetyl)-1-(prop-2-yn-1-yl)piperazine-2-carbonyl)-2,5-diazabicyclo[2.2.1]heptan-2-yl)ethan-1-one (SH-X-34A)

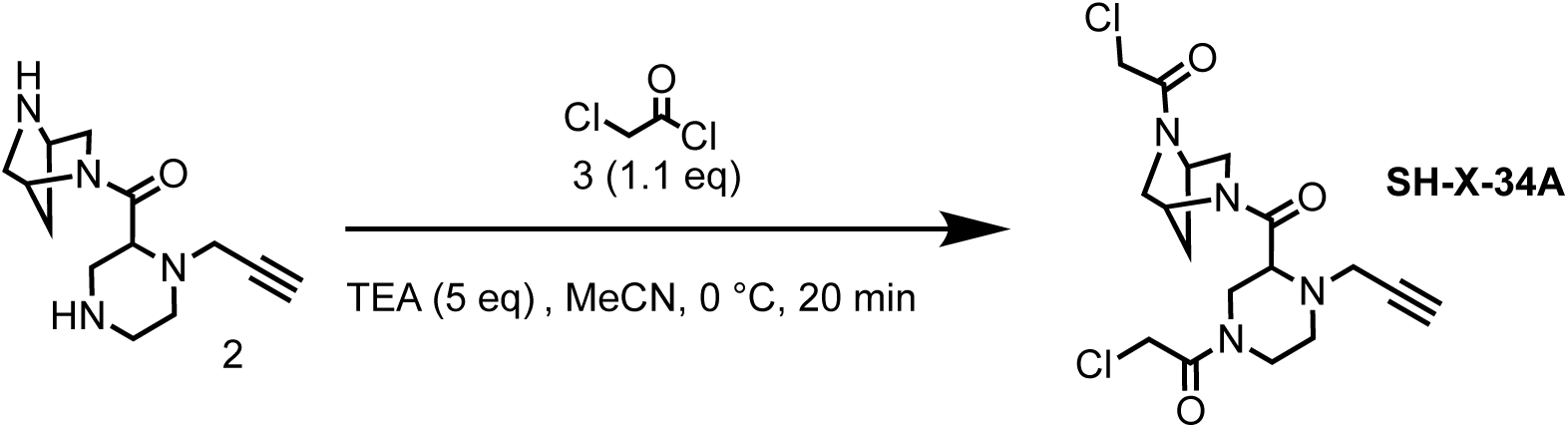

To a solution of (2,5-diazabicyclo[2.2.1]heptan-2-yl)(1-(prop-2-yn-1-yl)piperazin-2-yl) methanone (500 mg, 1.61 mmol, HCl salt) in MeCN (20 mL) was added TEA (812.80 mg, 8.03 mmol, 1.12 mL), then 2-chloroacetyl chloride (199 mg, 1.77 mmol, 140 μL) in MeCN (10 mL) was added into the reaction mixture at 0 °C, the reaction mixture was stirred at 0 °C for 20 min. LCMS showed the reaction was completed. The reaction mixture was filtered and the filtrate was concentrated to give a residue. The residue was purified by preparative-HPLC (column: Phenomenex Luna C18 150*25mm*10um; mobile phase: [water(FA)-MeCN]; gradient: 2%-32% B over 10 min) to give 2-chloro-1-(5-(4-(2-chloroacetyl)-1-(prop-2-yn-1-yl)piperazine-2-carbonyl)-2,5-diazabicyclo[2.2.1]heptan-2-yl)ethan-1-one (190 mg, 474 μmol, 29.5% yield, 99.9% purity) as an off-white solid.

LC-MS: 400.9 [M+H]^+^ / Ret time: 0.267 min / method: 5-95AB_0.8min.lcm

^1^H NMR (400 MHz, DMSO-*d*_6_): *δ* 5.15-4.68 (m, 2H), 4.57-4.36 (m, 2H), 4.35-4.02 (m, 2H), 3.87-3.62 (m, 2H), 3.61-3.49 (m, 1H), 3.48-3.11 (m, 8H), 3.06-2.76 (m, 2H), 2.50-2.30 (m, 1H), 2.01-1.68 (m, 2H)

### Synthesis of 1,1’-((2-((prop-2-yn-1-yloxy)methyl)piperazine-1,4-dicarbonyl)bis(indoline-5,1-diyl))bis(2-chloroethan-1-one) (SH-X-38)

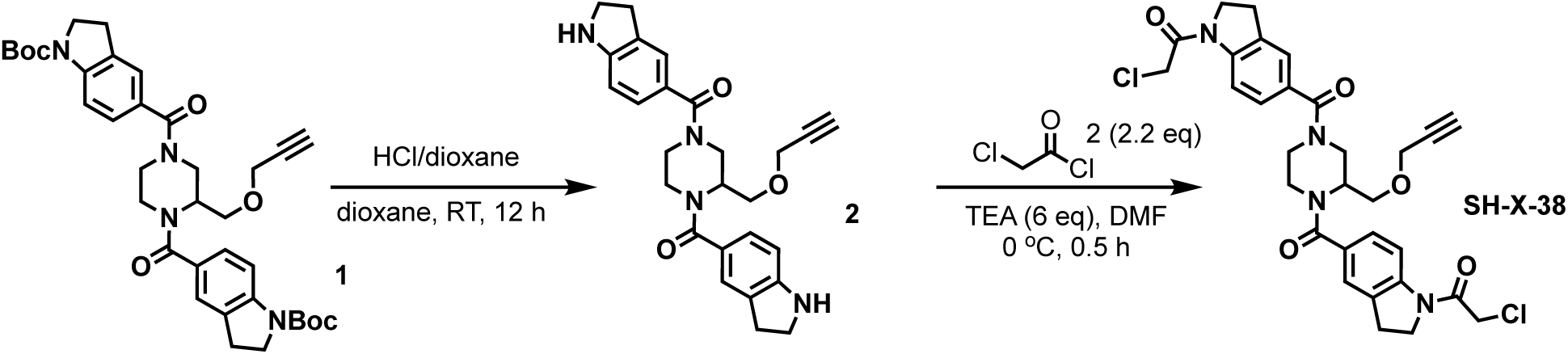

#### Step 1: Synthesis of (2-((prop-2-yn-1-yloxy)methyl)piperazine-1,4-diyl)bis(indolin-5-ylmethanone) (2)

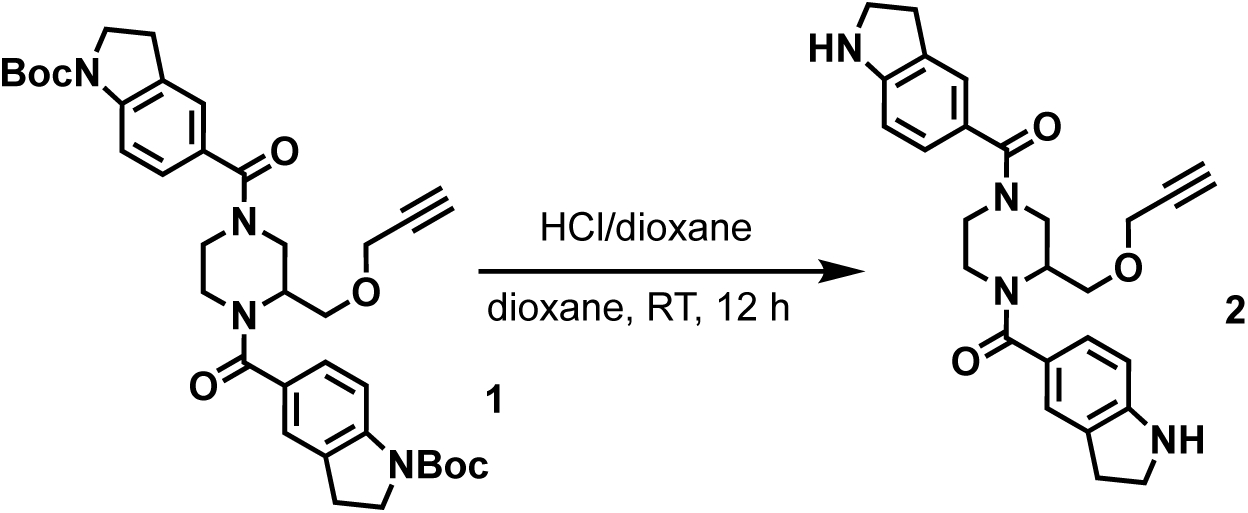

To a solution of di-*tert*-butyl 5,5’-(2-((prop-2-yn-1-yloxy)methyl)piperazine-1,4-dicarbonyl) bis(indoline-1-carboxylate) (1.13 g, 1.75 mmol) in dioxane (10 mL) was added HCl/dioxane (4 M, 10 mL), the mixture was stirred at 20 °C for 12 hrs. LCMS showed the reaction was completed. The reaction mixture was concentrated to give (2-((prop-2-yn-1-yloxy)methyl) piperazine-1,4-diyl)bis(indolin-5-ylmethanone) (100 mg, 209 μmol, 11.9% yield, 93.0% purity, HCl salt) as a yellow solid.

LC-MS: 445.4 [M+H]+/ Ret time: 0.200 min / method: 5-95AB_0.8min.lcm.

^1^H NMR (400 MHz, DMSO-*d*6) *δ* 7.44 (s, 2H), 7.37 - 7.32 (m, 2H), 7.32 - 7.27 (m, 2H), 4.11 (s, 2H), 4.06 - 3.99 (m, 1H), 3.69 (t, *J* = 8.0 Hz, 4H), 3.56 (s, 4H), 3.53 - 3.46 (m, 2H), 3.21 - 3.14 (m, 5H), 3.01 (s, 2H)

#### Step 2: Synthesis of 1,1’-((2-((prop-2-yn-1-yloxy)methyl)piperazine-1,4-dicarbonyl)bis(indoline-5,1-diyl))bis(2-chloroethan-1-one) (SH-X-38)

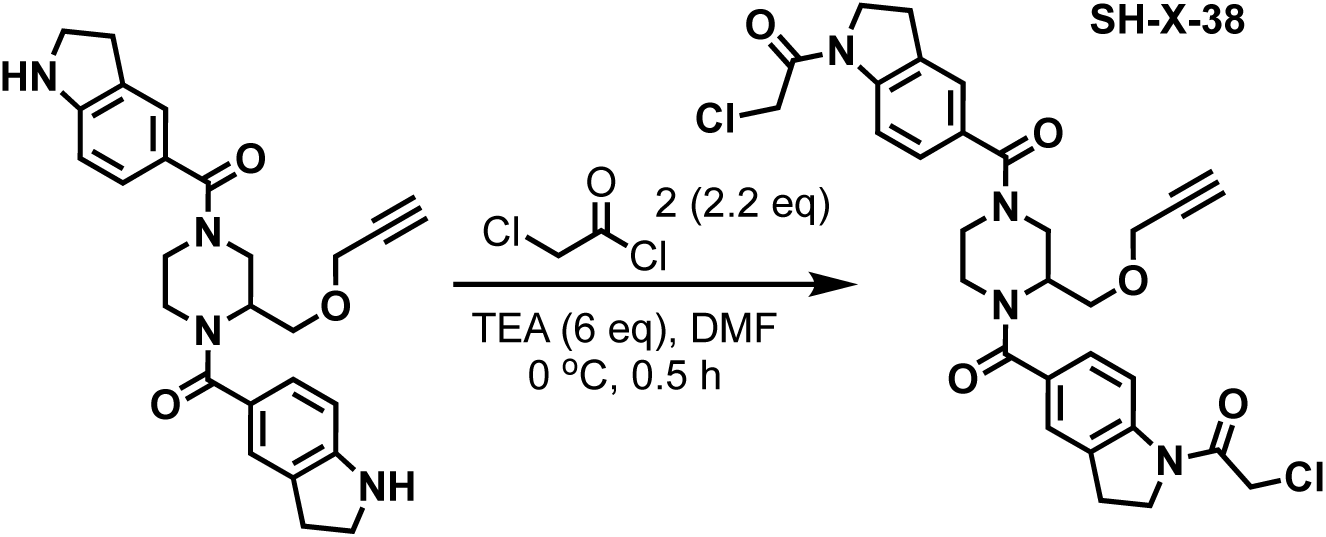

To a solution of (2-((prop-2-yn-1-yloxy)methyl)piperazine-1,4-diyl)bis(indolin-5-ylmethanone) (600 mg, 1.35 mmol) in DMF (20 mL) was added TEA (819 mg, 8.10 mmol, 1.13 mL), then 2-chloroacetyl chloride (335 mg, 2.97 mmol, 236 μL) was added into the reaction mixture at 0 °C, the mixture was stirred at 0 °C for 0.5 hr. LCMS showed the reaction was completed. The reaction mixture was quenched with H2O (1 mL) and the reaction mixture was filtered and the filtrate was concentrated to give a residue. The residue was purified by preparative-HPLC (column: Phenomenex luna C18 150*25mm* 10um;mobile phase: [water(FA)-MeCN];gradient:27%-57% B over 8 min) to give 1,1’-((2-((prop-2-yn-1-yloxy)methyl)piperazine-1,4-dicarbonyl)bis(indoline-5,1-diyl))bis(2-chloroethan-1-one) (60.0 mg, 96.9 μmol, 7.16% yield, 96.2% purity) as an yellow solid.

LC-MS: 597.4 [M+H]+/ Ret time: 0.390 min / method: 5-95AB_0.8min.lcm.

^1^H NMR (400 MHz, DMSO-*d*6) *δ* 8.06 (t, *J* = 7.2 Hz, 2H), 7.33 (d, *J* = 3.6 Hz, 2H), 7.27 (d, *J* = 8.0 Hz, 2H), 4.56 (s, 4H), 4.42 - 4.34 (m, 1H), 4.24 - 4.00 (m, 8H), 3.70 - 3.45 (m, 4H), 3.21 (s, 7H).

### Synthesis of 3-(2-chloroacetamido)-*N*-(1-(2-chloroacetyl)-1,2,3,4-tetrahydroquinolin-4-yl)-4-(prop-2-yn-1-yloxy)benzamide (SH-X-22A)

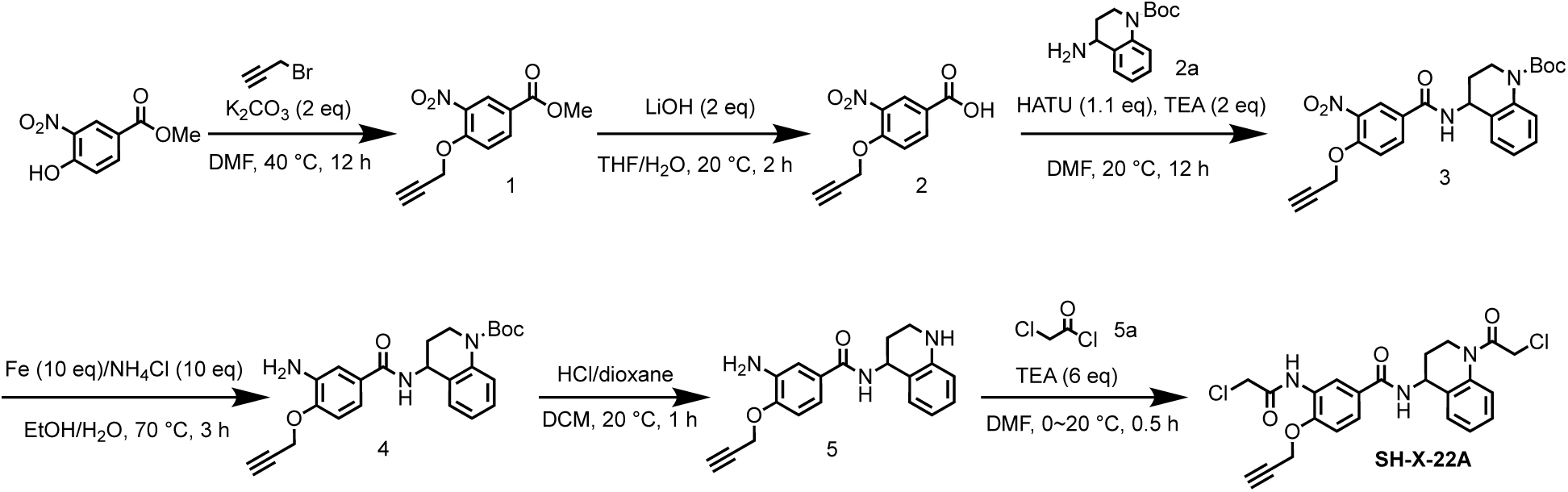

#### Step 1: Preparation of methyl 3-nitro-4-(prop-2-yn-1-yloxy)benzoate (1)

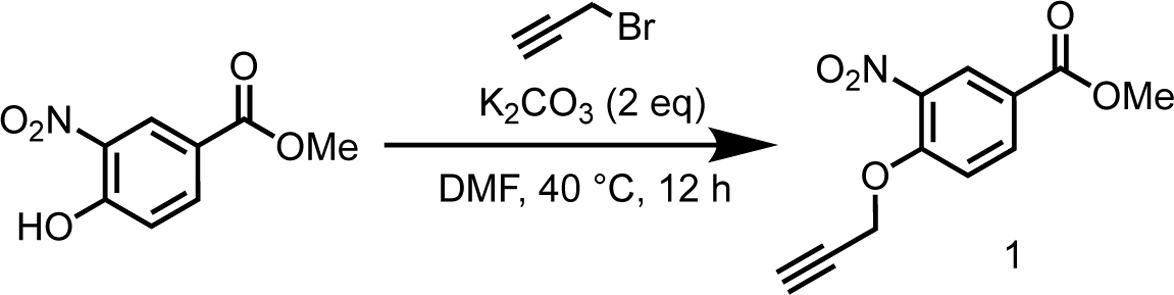

To a solution of methyl 4-hydroxy-3-nitrobenzoate (5.00 g, 25.4 mmol) in DMF (50 mL) was added K_2_CO_3_ (4.21 g, 30.4 mmol), the mixture was stirred at 20 °C for 0.5 hr, then 3-bromoprop-1-yne (5.66 g, 38.0 mmol, 4.10 mL) was added to the mixture, the reaction mixture was stirred at 40 °C for 12 hrs. TLC (petroleum ether: ethyl acetate=5:1) showed the reaction was completed. The reaction mixture was poured into H_2_O (250 mL), extracted with EtOAc (250 mL×3). The organic layers were collected and washed with brine (250 mL×3), dried over Na_2_SO_4_ and concentrated to give methyl 3-nitro-4-(prop-2-yn-1-yloxy)benzoate (5.60 g, 23.8 mmol, 93.9% yield) as a yellow solid.

#### Step 2: Preparation of 3-nitro-4-(prop-2-yn-1-yloxy)benzoic acid (2)

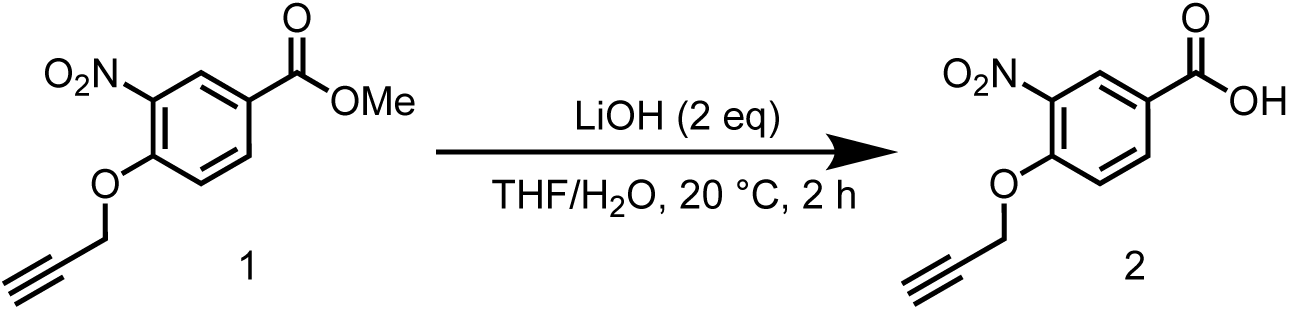

To a solution of 3-nitro-4-(prop-2-yn-1-yloxy)benzoate (5.60 g, 23.8 mmol) in THF (25 mL) and H_2_O (25 mL) was added LiOH-H_2_O (3.00 g, 71.4 mmol), the reaction mixture was stirred at 20 °C for 1.5 hrs. TLC (petroleum ether: ethyl acetate=5:1) showed the reaction was completed. The reaction mixture was concentrated in vacuum to remove solvent, and then acidified with 1M HCl to pH=4, filtered, the filter cake was concentrated in vacuum to give 3-nitro-4-(prop-2-yn-1-yloxy)benzoic acid (4.90 g, 22.2 mmol, 93.1% yield) as a yellow solid.

^1^H NMR (400 MHz, DMSO-*d*_6_): *δ* 13.35 (s, 1H), 8.37 (d, *J* = 2.0 Hz, 1H), 8.21 (m, 8.8 Hz, 1H), 7.52 (d, *J* = 8.8 Hz, 1H), 5.13 (d, *J* = 2.0 Hz, 2H), 3.76 (t, *J* = 2.0 Hz, 1H).

#### Step 3: Preparation of *tert*-butyl 4-(3-nitro-4-(prop-2-yn-1-yloxy)benzamido)-3,4-dihydroquinoline-1(2*H*)-carboxylate (3)

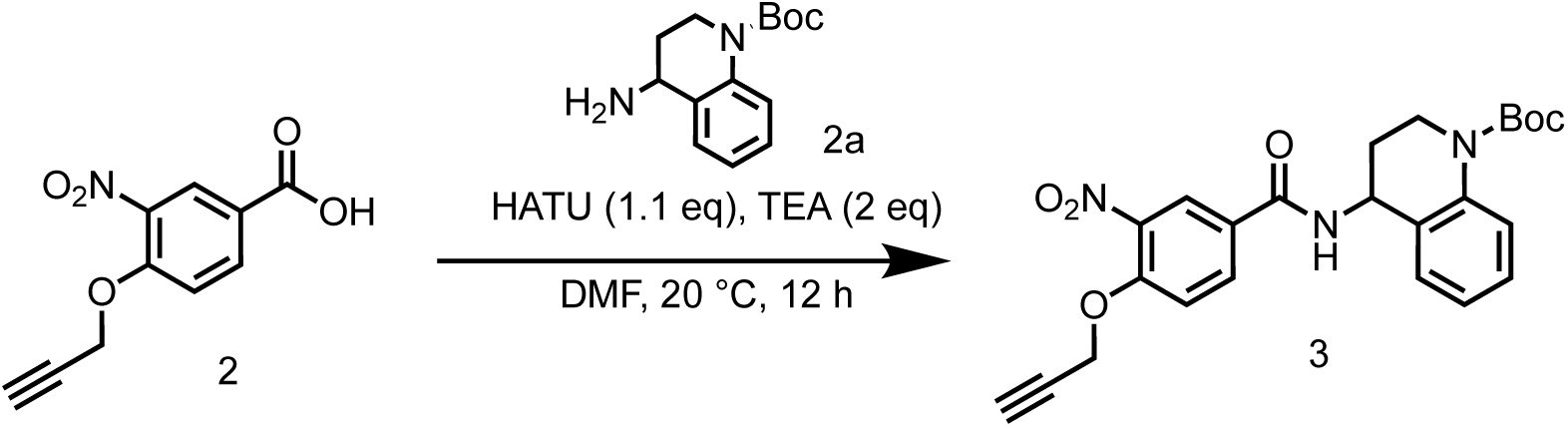

To a solution of 3-nitro-4-(prop-2-yn-1-yloxy)benzoic acid (500 mg, 2.26 mmol) and *tert*-butyl 4-amino-3,4-dihydroquinoline-1(2H)-carboxylate (505 mg, 2.03 mmol) in DCM (10 mL) was added EDCI (650 mg, 3.39 mmol) and DMAP (27.6 mg, 226 μmol). The reaction mixture was stirred at 20 °C for 12 hrs. LCMS showed the reaction was completed. The reaction mixture was concentrated in vacuum to give a residue. The residue was purified by column chromatography on silica gel (eluted with petroleum ether: ethyl acetate=5:1 to 1:1) to give *tert*-butyl 4-(3-nitro-4-(prop-2-yn-1-yloxy)benzamido)-3,4-dihydroquinoline-1(2*H*)-carboxylate (880 mg, 1.95 mmol, 86.2% yield) as a yellow oil.

LC-MS: 497.4 [M+Na]^+^ / Ret time: 0.480 min / method: 5-95AB_0.8min.

#### Step 4: Preparation of *tert*-butyl 4-(3-amino-4-(prop-2-yn-1-yloxy)benzamido)-3,4-dihydroquinoline-1(2*H*)-carboxylate (4)

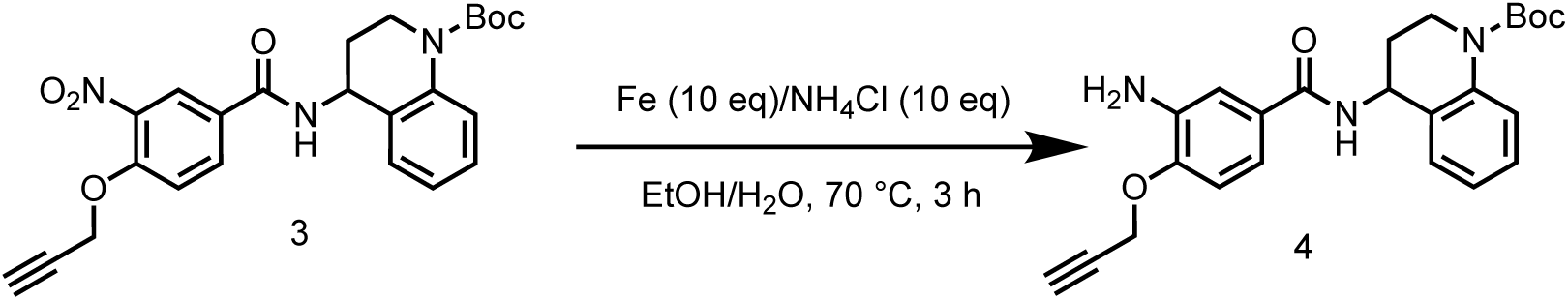

To a solution of *tert*-butyl 4-(3-nitro-4-(prop-2-yn-1-yloxy)benzamido)-3,4-dihydroquinoline-1(2*H*)-carboxylate (880 mg, 1.95 mmol) in EtOH (20 mL) and H_2_O (6 mL) was added Fe (1.09 g, 19.5 mmol) and NH_4_Cl (1.04 g, 19.5 mmol), the reaction mixture was stirred at 70 °C for 2 hrs. LCMS showed the reaction was completed. The mixture was cooled to 20 °C and filtered through celite pad, then the filtrate was concentrated in vacuum to give *tert*-butyl 4-(3-amino-4-(prop-2-yn-1-yloxy)benzamido)-3,4-dihydroquinoline-1(2*H*)-carboxylate (500 mg, crude) as a brown solid.

LC-MS: 366.3 [M-56+H]^+^ / Ret time: 0.404 min / method: 5-95AB_0.8min.lcm.

#### Step 5: Preparation of 3-amino-4-(prop-2-yn-1-yloxy)*-N-*(1,2,3,4-tetrahydroquinolin-4-yl)benzamide (5)

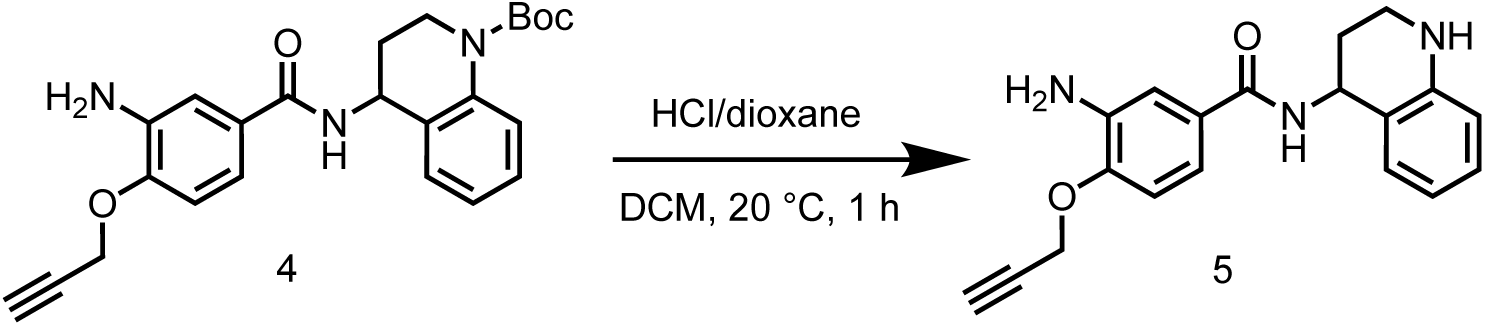

To a solution of *tert*-butyl 4-(3-amino-4-(prop-2-yn-1-yloxy)benzamido)-3,4-dihydroquinoline-1(2*H*)-carboxylate (500 mg, 1.19 mmol) in DCM (5 mL) was added HCl/dioxane (4 M, 5 mL), the mixture was stirred at 20 °C for 0.5 hr. LCMS showed the reaction was completed. The reaction mixture was concentrated in vacuum to give 3-amino-4-(prop-2-yn-1-yloxy)-*N*-(1,2,3,4-tetrahydroquinolin-4-yl)benzamide (300 mg, crude, HCl salt) was obtained as a yellow solid.

LC-MS: 322.2 [M+H]^+^ / Ret time: 0.231 min / method: 5-95AB_0.8min.lcm.

#### Step 6: Preparation of 3-(2-chloroacetamido)-*N*-(1-(2-chloroacetyl)-1,2,3,4-tetrahydroquinolin-4-yl)-4-(prop-2-yn-1-yloxy)benzamide (6)

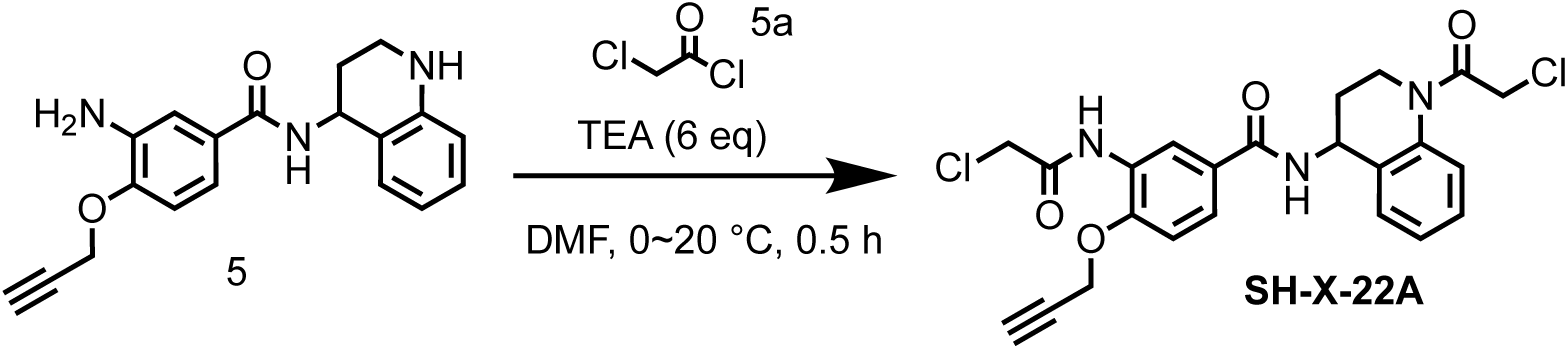

To a mixture of 3-amino-4-(prop-2-yn-1-yloxy)*-N-*(1,2,3,4-tetrahydroquinolin-4-yl) benzamide (300 mg, crude, HCl salt) and TEA (509 mg, 5.03 mmol, 700 μL) in MeCN (20 mL) was added 2-chloroacetyl chloride (208 mg, 1.84 mmol, 147μL), the mixture was stirred at 0 °C for 0.5 hr under N2. LCMS showed the reaction was completed. The reaction mixture was extracted with H_2_O (10 mL) and ethyl acetate (10 mL × 3), washed with brine (10 mL), the organic phase was dried over Na_2_SO_4_, concentrated in vacuum to give 3-(2-chloroacetamido)*-N-*(1-(2-chloroacetyl)-1,2,3,4-tetrahydroquinolin-4-yl)-4-(prop-2-yn-1-yloxy)benzamide (300 mg, 632 μmol, 75.4% yield) as a brown oil.

LC-MS: 474.2 [M+H]^+^ / Ret time: 0.395 min / method: 5-95AB_0.8min.

^1^H NMR (400 MHz, DMSO-*d*_6_) *δ* 9.64 (s, 1H), 8.82 (d, *J* = 8.4 Hz, 1H), 8.48 (s, 1H), 7.74 (m, *J* = 2.0, 8.4 Hz, 1H), 7.69 - 7.57 (m, 1H), 7.31 - 7.16 (m, 4H), 5.35 - 5.16 (m, 1H), 4.99 (d, *J* = 2.0 Hz, 2H), 4.68 - 4.52 (m, 2H), 4.40 (s, 2H), 3.99 - 3.76 (m, 2H), 3.66 (t, *J* = 2.0 Hz, 1H), 2.29 - 2.17 (m, 1H), 1.98 (m, *J* = 4.4, 8.7 Hz, 1H).

### Synthesis of 5-(2-chloroacetamido)*-N-*(1-(2-chloroacetyl)azepan-3-yl)-2-(prop-2-yn-1-yloxy)benzamide (SH-X-17)

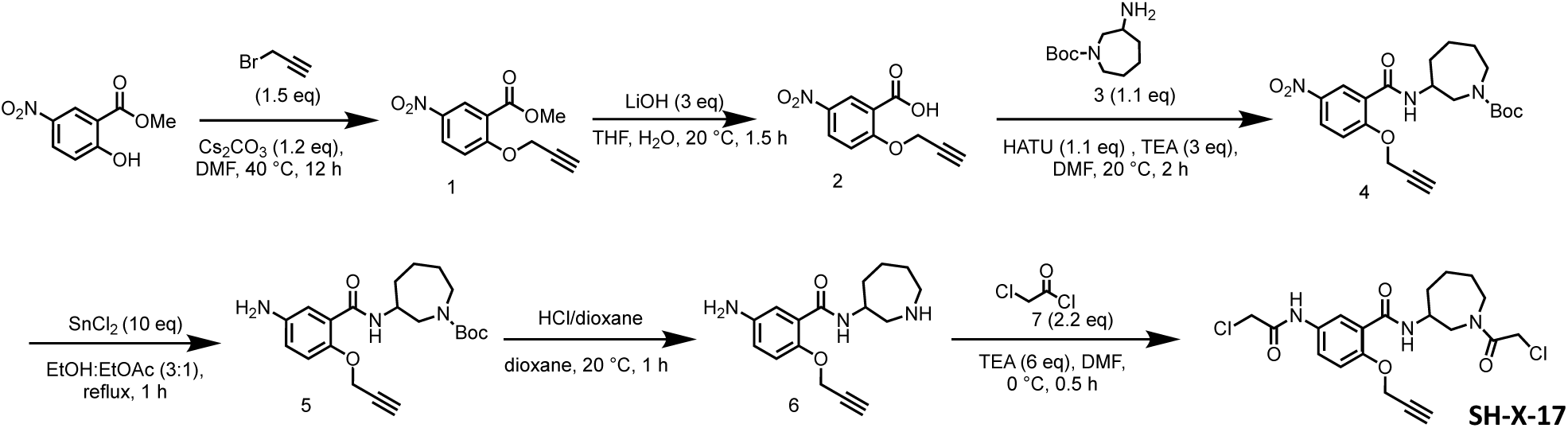

#### Step 1: Preparation of methyl 5-nitro-2-(prop-2-yn-1-yloxy)benzoate (1)

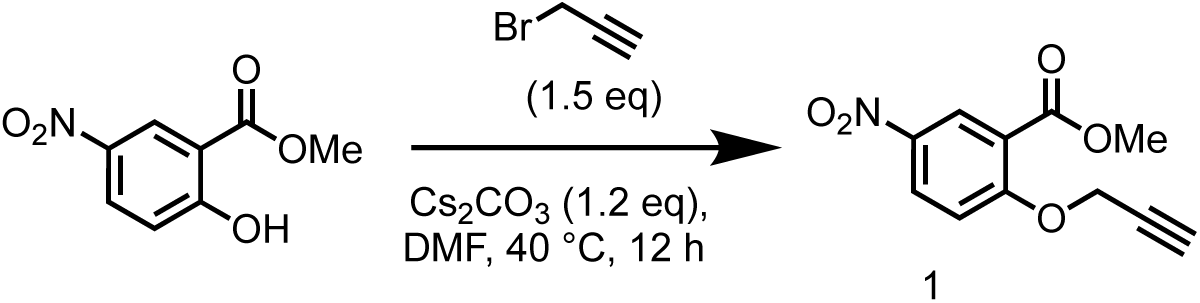

To a solution of methyl 2-hydroxy-5-nitrobenzoate (200 mg, 1.01 mmol) in DMF (5 mL) was added Cs_2_CO_3_ (397 mg, 1.22 mmol), the reaction mixture was stirred at 20 °C for 0.5 hr under N2 atmosphere, then 3-bromoprop-1-yne (226 mg, 1.52 mmol, 164 μL) was added into the reaction mixture, the mixture was stirred at 40 °C for 12 hrs. TLC (Petroleum ether: Ethyl acetate=5:1) showed the reaction was completed. The reaction mixture was poured into H_2_O (30 mL), extracted with EtOAc (30 mL×3). The organic layers were collected and washed with brine (30 mL×3), dried over Na_2_SO_4_ and concentrated to give methyl 5-nitro-2-(prop-2-yn-1-yloxy)benzoate (170 mg, 723 μmol, 71.3% yield) as a brown oil.

^1^H NMR (400 MHz, CDCl_3_) *δ* 8.76 (d, *J* = 2.8 Hz, 1H), 8.41 (dd, *J* = 2.8, 9.2 Hz, 1H), 7.30-7.29 (m, 1H), 4.95 (d, *J* = 2.4 Hz, 2H), 3.98 (s, 3H), 2.64 (t, *J* = 2.4 Hz, 1H).

#### Step 2: Preparation of 5-nitro-2-(prop-2-yn-1-yloxy)benzoic acid (2)

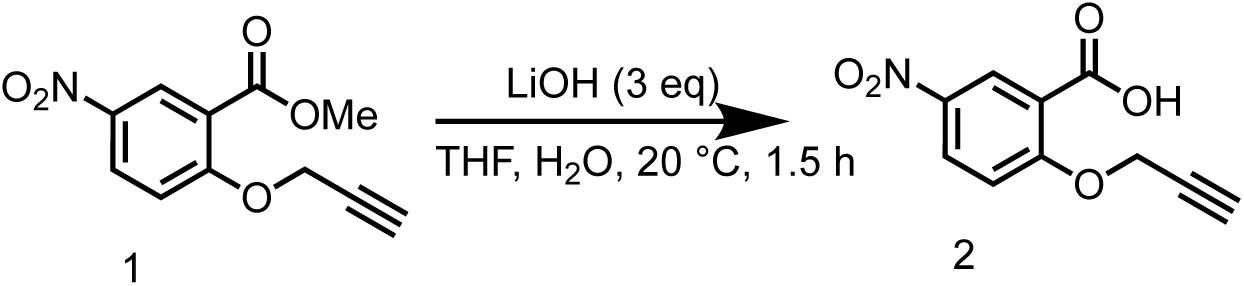

To a solution of methyl 5-nitro-2-(prop-2-yn-1-yloxy)benzoate (5.2 g, 22.1 mmol) in THF (26 mL) and H_2_O (26 mL) was added LiOH•H_2_O (2.78 g, 66.3 mmol), the reaction mixture was stirred at 20 °C for 1.5 hrs. TLC (Petroleum ether: Ethyl acetate=5:1) showed the reaction was completed. The reaction mixture was concentrated in vacuum to remove solvent, and then acidified with 1M HCl to pH=4. The reaction mixture was filtered and the filter cake was concentrated in vacuum to give 5-nitro-2-(prop-2-yn-1-yloxy)benzoic acid (4.3 g, crude) as a brown solid.

#### Step 3: Preparation of *tert*-butyl 3-(5-nitro-2-(prop-2-yn-1-yloxy)benzamido)azepane-1-carboxylate (4)

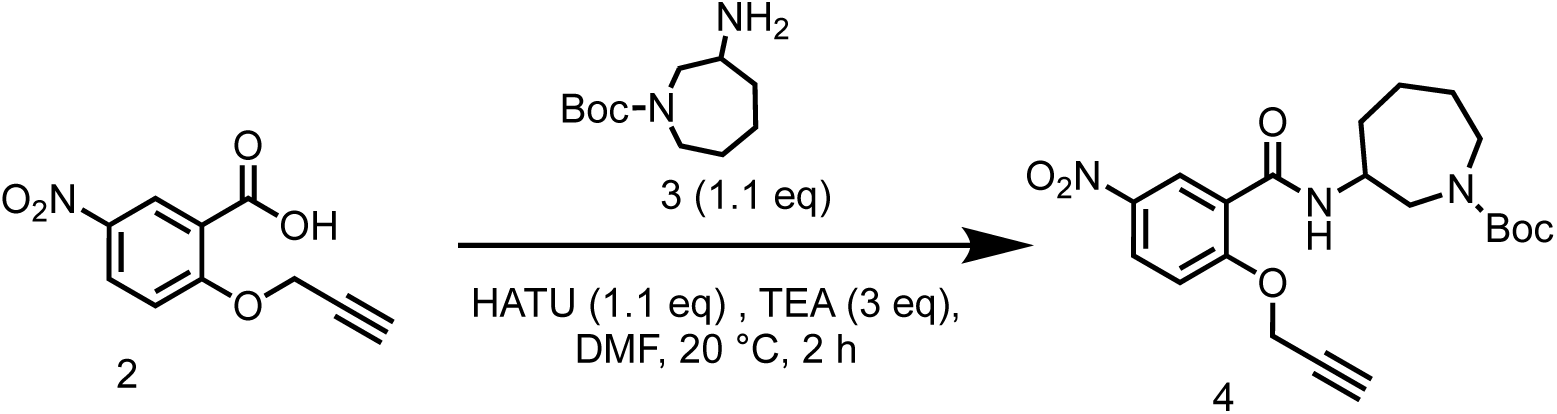

To a solution of 5-nitro-2-(prop-2-yn-1-yloxy)benzoic acid (4.3 g, crude) in DMF (43 mL) was added HATU (8.13 g, 21.4 mmol) and TEA (5.90 g, 58.3 mmol, 8.12 mL), then *tert*-butyl 3-aminoazepane-1-carboxylate (4.58 g, 21.4 mmol) was added into the reaction mixture, the reaction mixture was stirred at 20 °C for 12 hrs. LCMS showed the reaction was completed. The reaction mixture was poured into H_2_O (200 mL), extracted with EtOAc (200 mL×3). The organic layers were collected and washed with brine (200 mL×3), dried over Na_2_SO_4_ and concentrated in vacuum to give a residue. The residue was purified by column chromatography on silica gel (eluted with petroleum ether: ethyl acetate=3:1 to 1:1) to give *tert*-butyl 3-(5-nitro-2-(prop-2-yn-1-yloxy)benzamido)azepane-1-carboxylate (7.00 g, 16.8 mmol, 86.3% yield) as a yellow oil.

LC-MS: 418.4 [M+H]^+^ / Ret time: 0.439 min / method: 5-95AB_0.8min.lcm.

^1^H NMR (400 MHz, CDCl_3_) *δ* 9.11-8.99 (m, 2H), 8.31 (dd, *J* = 2.8, 8.8 Hz, 1H), 5.25-5.01 (m, 2H), 4.47-4.34 (m, 1H), 4.06-3.92 (m, 2H), 3.25 (dd, *J* = 6.8, 15.2 Hz, 1H), 2.33-2.21 (m, 1H), 1.75 (s, 4H), 1.50 (s, 9H), 1.36 (s, 4H).

#### Step 4: Preparation of *tert*-butyl 3-(5-amino-2-(prop-2-yn-1-yloxy)benzamido)azepane-1-carboxylate (5)

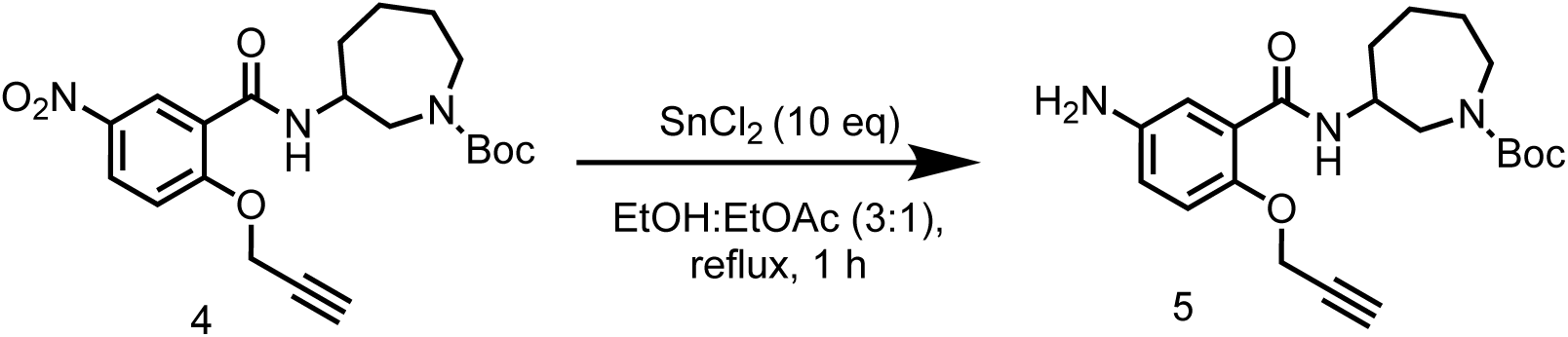

To a solution of *tert*-butyl 3-[(5-nitro-2-prop-2-ynoxy-benzoyl)amino]azepane-1-carboxylate (3.00 g, 7.19 mmol) in EtOH (30 mL) and EtOAc (10 mL) was added SnCl2•2 H_2_O (16.2 g, 71.9 mmol), the reaction mixture was stirred at 80 °C for 1 hr. The reaction mixture was cooled and quenched with saturated NaHCO_3_ (aq), then extracted with EtOAc (200 mL×3). The organic layers were collected and washed with brine (200 mL×3), dried over Na_2_SO_4_ and concentrated to give *tert*-butyl 3-(5-amino-2-(prop-2-yn-1-yloxy)benzamido)azepane-1-carboxylate (1.06 g, 2.74 mmol, 38.1% yield) as a yellow oil.

LC-MS: 288.2 [M+H-100]+ / method: 5-95AB_0.8min.lcm

#### Step 5: Preparation of 5-amino*-N-*(azepan-3-yl)-2-(prop-2-yn-1-yloxy)benzamide (5)

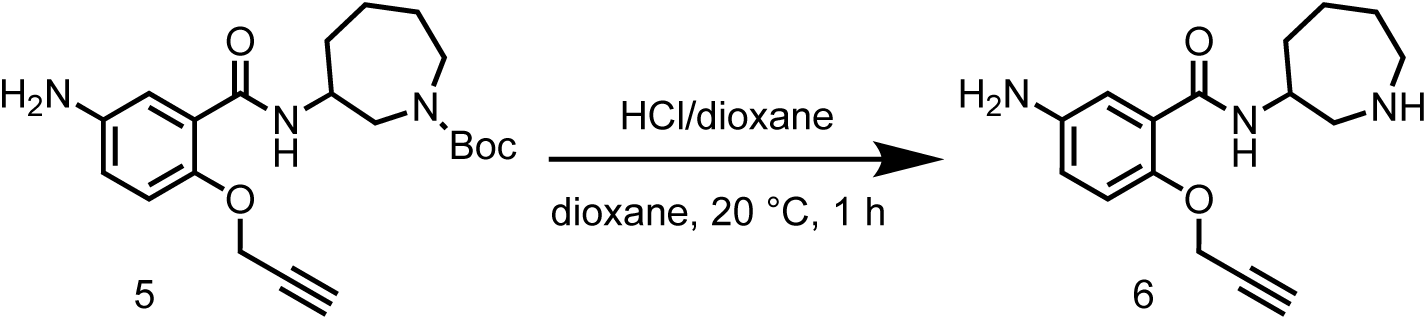

To a solution of *tert*-butyl 3-(5-amino-2-(prop-2-yn-1-yloxy)benzamido)azepane-1-carboxylate (400 mg, 1.03 mmol) in dioxane (1 mL) was added HCl/dioxane (4 M, 3 mL), the reaction mixture was stirred at 20 °C for 1 hr. LCMS showed the reaction was completed. The reaction mixture was concentrated in vacuum to give 5-amino*-N-*(azepan-3-yl)-2-(prop-2-yn-1-yloxy)benzamide (770 mg, crude, HCl salt) as a yellow solid.

LC-MS: 288.3 [M+H]^+^ / method: 0-60AB_1min.lcm.

#### Step 6: Preparation of 5-(2-chloroacetamido)*-N-*(1-(2-chloroacetyl)azepan-3-yl)-2-(prop-2-yn-1-yloxy)benzamide (SH-X-17)

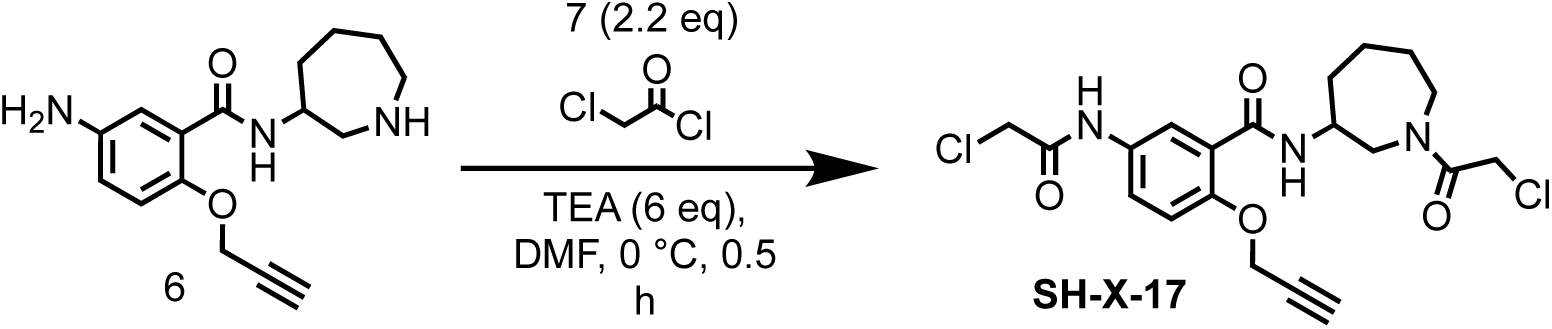

To a mixture of 5-amino*-N-*(azepan-3-yl)-2-(prop-2-yn-1-yloxy)benzamide (300 mg, 926 μmol, HCl salt) and TEA (562 mg, 5.56 mmol, 774 μL) in DMF (2 mL) was added 2-chloroacetyl chloride (230 mg, 2.04 mmol, 162 μL) at 0 °C, the reaction mixture was stirred at 0 °C for 0.5 hr. LCMS showed the reaction was completed. The reaction mixture was quenched with H_2_O (0.1 mL) and concentrated in vacuum to give a residue. The residue was purified by reversed-phase HPLC (0.1% FA condition; 80 g Sepa Flash Silica Flash Column, Eluent of 40∼44% MeCN / H_2_O gradient 60 mL/min) to give 5-(2-chloroacetamido)*-N-*(1-(2-chloroacetyl)azepan-3-yl)-2-(prop-2-yn-1-yloxy)benzamide (60.0 mg, 136 μmol, 14.7% yield) as a white solid.

LC-MS: 440.3 [M+H]^+^ / Ret time: 0.348 min / method: 5-95AB_0.8min.lcm.

^1^H NMR (400 MHz, DMSO-*d*_6_) *δ* 10.34 (s, 1H), 8.54 - 8.12 (m, 1H), 8.01 - 7.88 (m, 1H), 7.80 - 7.69 (m, 1H), 7.19 (d, *J* = 8.8 Hz, 1H), 5.00 - 4.87 (m, 2H), 4.69 - 4.38 (m, 2H), 4.23 (s, 2H), 4.21 - 3.94 (m, 1H), 3.81 - 3.70 (m, 1H), 3.69 - 3.61 (m, 2H), 3.46 - 3.38 (m, 1H), 3.31 (s, 1H), 1.94 - 1.37 (m, 6H).

### Synthesis of 3-(2-chloroacetamido)-*N*-(3-(4-(2-chloroacetyl)piperazin-1-yl)propyl)-4-(prop-2-yn-1-yloxy)benzamide (SH-X-18)

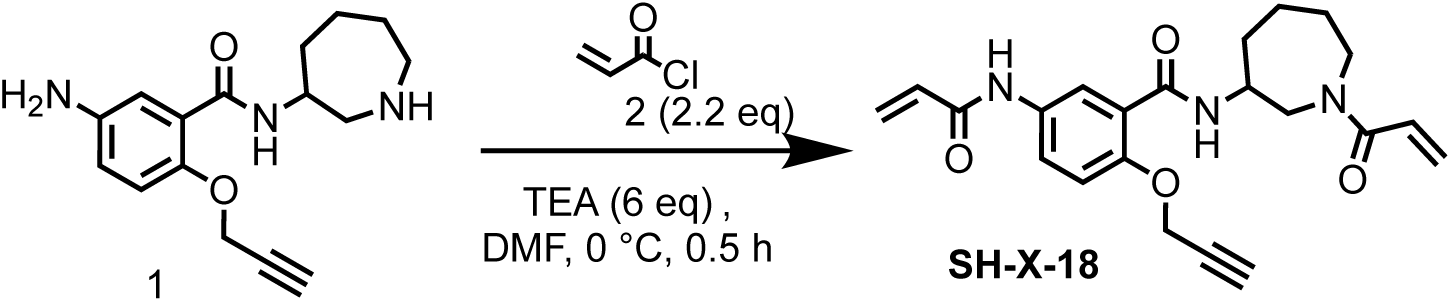

To a mixture of 5-amino-*N*-(azepan-3-yl)-2-(prop-2-yn-1-yloxy)benzamide (400 mg, 1.24 mmol, HCl salt) and TEA (750 mg, 7.41 mmol, 1.03 mL) in DMF (3 mL) was added acryloyl chloride (246 mg, 2.72 mmol, 221 μL) at 0°C, the reaction mixture was stirred at 0 °C for 0.5 hr. LCMS showed the reaction was completed. The reaction mixture was quenched with H_2_O (0.1 mL) and concentrated in vacuum to give a residue. The residue was purified by reversed-phase MPLC (0.1% FA condition; 80 g Sepa Flash Silica Flash Column, Eluent of 25∼30% MeCN/H_2_O gradient 60 mL/min) to give 5-acrylamido-*N*-(1-acryloylazepan-3-yl)-2-(prop-2-yn-1-yloxy)benzamide (38.7 mg, 97.8 μmol, 7.92% yield, 100% purity) as a white solid.

LC-MS: 396.3 [M+H]^+^ / Ret time: 0.361 min / method: 5-95AB_0.8min.lcm.

^1^H NMR (400 MHz, DMSO-*d*_6_): *δ* 10.19 (s, 1H), 8.66-8.09 (m, 1H), 8.02 (dd, *J* = 2.4, 10.8 Hz, 1H), 7.92-7.81 (m, 1H), 7.18 (dd, *J* = 5.6, 8.8 Hz, 1H), 7.08-6.77 (m, 1H), 6.46-6.35 (m, 1H), 6.29-6.13 (m, 2H), 5.78-5.65 (m, 2H), 5.02-4.84 (m, 2H), 4.27-3.94 (m, 1H), 3.92-3.76 (m, 1H), 3.76-3.58 (m, 2H), 3.42 (m, *J* = 4.8, 10.0 Hz, 1H), 3.24-3.14 (m, 1H), 1.94-1.33 (m, 6H).

### Synthesis of 3-(2-chloroacetamido)-*N*-(3-(4-(2-chloroacetyl)piperazin-1-yl)propyl)-4-((1-(2-(2-(6-((4*R*,5*S*)-5-methyl-2-oxoimidazolidin-4-yl)hexanamido)ethoxy)ethyl)-1*H*-1,2,3-triazol-4-yl)methoxy)benzamide (SH-X-30)

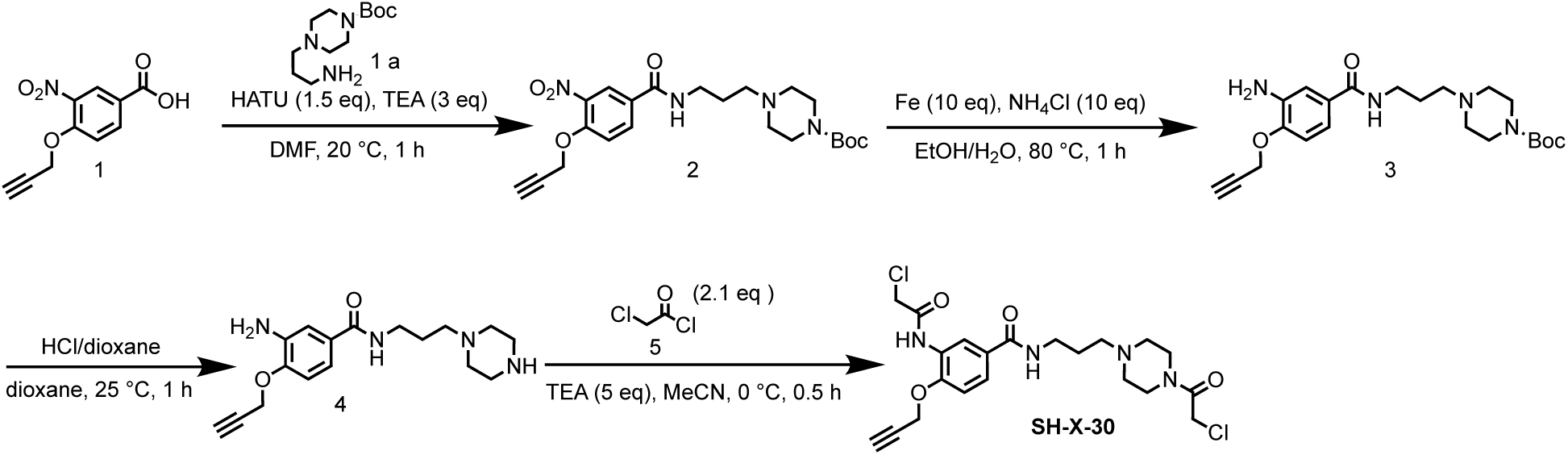

#### Step 1: Preparation of *tert*-butyl 4-(3-(3-nitro-4-(prop-2-yn-1-yloxy)benzamido)propyl)piperazine-1-carboxylate (2)

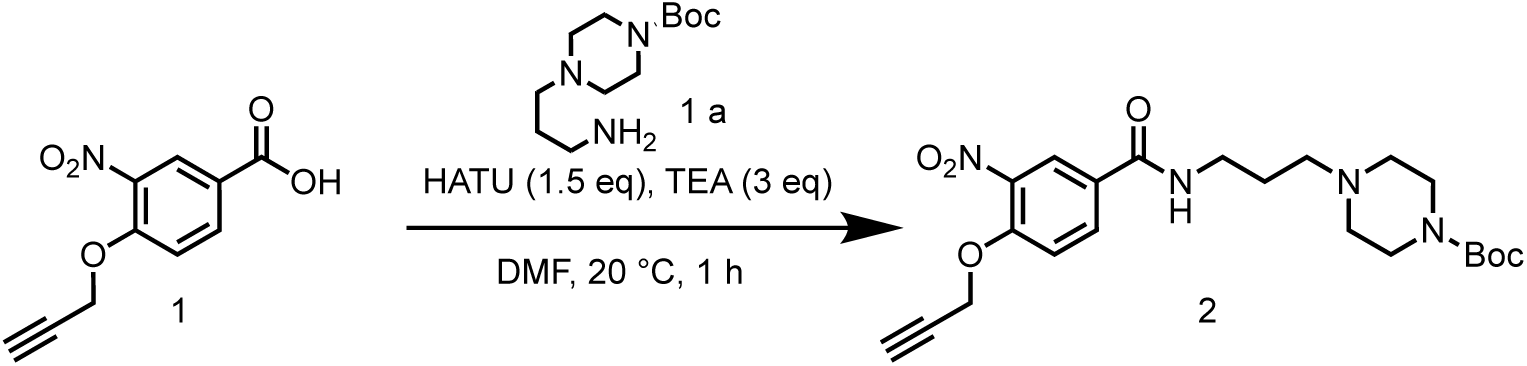

To a solution of 3-nitro-4-(prop-2-yn-1-yloxy)benzoic acid (3.00 g, 13.5 mmol) and *tert*-butyl 4-(3-aminopropyl)piperazine-1-carboxylate (3.63 g, 14.9 mmol) in DMF (30 mL) was added HATU (7.74 g, 20.3 mmol) and TEA (4.12 g, 40.6 mmol, 5.66 mL). The mixture was stirred at 20 °C for 1 hr. LCMS showed the reaction was completed. The reaction mixture was poured into H_2_O (100 mL), extracted with EtOAc (100 mL×3). The organic layers were collected and washed with brine (100 mL×3), dried over Na_2_SO_4_ and concentrated to give a residue. The residue was purified by column chromatography on silica gel (eluted with petroleum ether: ethyl acetate=10:0 to 0:1) to give *tert*-butyl 4-(3-(3-nitro-4-(prop-2-yn-1-yloxy)benzamido)propyl)piperazine-1-carboxylate (6.00 g, 13.4 mmol, 99.1% yield) as a yellow oil.

LC-MS: 445.2 [M-H]^+^ / Ret time: 0.590 min / method: \5-95CD_1min_NEG.lcm

#### Step 2: Preparation of *tert*-butyl 4-(3-(3-amino-4-(prop-2-yn-1-yloxy)benzamido)propyl)piperazine-1-carboxylate (3)

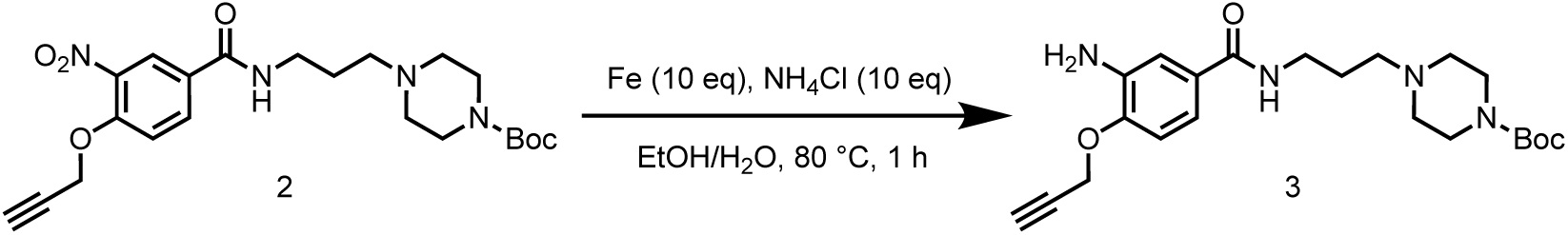

To a solution of *tert*-butyl 4-(3-(3-nitro-4-(prop-2-yn-1-yloxy)benzamido)propyl)piperazine-1-carboxylate (4.00 g, 8.96 mmol) in ethyl alcohol (9 mL) and H_2_O (3 mL) was added NH_4_Cl (2.40 g, 44.8 mmol) and Fe (2.50 g, 44.8 mmol). The mixture was stirred at 80 °C for 1 hr. LCMS showed the reaction was completed. The suspension was filtered through a pad of silica gel to give *tert*-butyl 4-(3-(3-amino-4-(prop-2-yn-1-yloxy)benzamido)propyl)piperazine-1-carboxylate (3.50 g, 8.40 mmol, 93.8% yield) as a yellow oil.

LC-MS: 417.3 [M+H]^+^ / Ret time: 0.300 min / method: \ 5-95AB_0.8min.lcm

#### Step 3: Preparation of 3-amino-*N*-(3-(piperazin-1-yl)propyl)-4-(prop-2-yn-1-yloxy)benzamide (4)

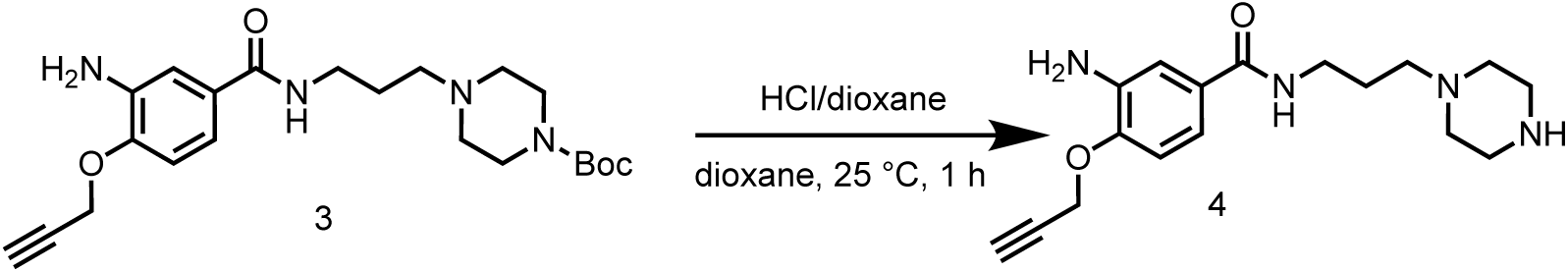

To a solution of *tert*-butyl 4-(3-(3-amino-4-(prop-2-yn-1-yloxy)benzamido)propyl)piperazine-1-carboxylate (3.50 g, 8.40 mmol) in dioxane (17 mL) was added HCl/dioxane (4 M, 17 mL). The mixture was stirred at 25 °C for 1 hr. LCMS showed the reaction was completed. The reaction mixture was concentrated in vacuum to give 3-amino-*N*-(3-(piperazin-1-yl)propyl)-4-(prop-2-yn-1-yloxy)benzamide (4.00 g, crude, HCl salt) as a yellow solid.

LC-MS: 317.1 [M+H]^+^ / Ret time: 0.245 min / method: \0-60AB_0.8min.lcm

#### Step 4: Preparation of 3-(2-chloroacetamido)-*N*-(3-(4-(2-chloroacetyl)piperazin-1-yl)propyl)-4-(prop-2-yn-1-yloxy)benzamide (SH-X-30)

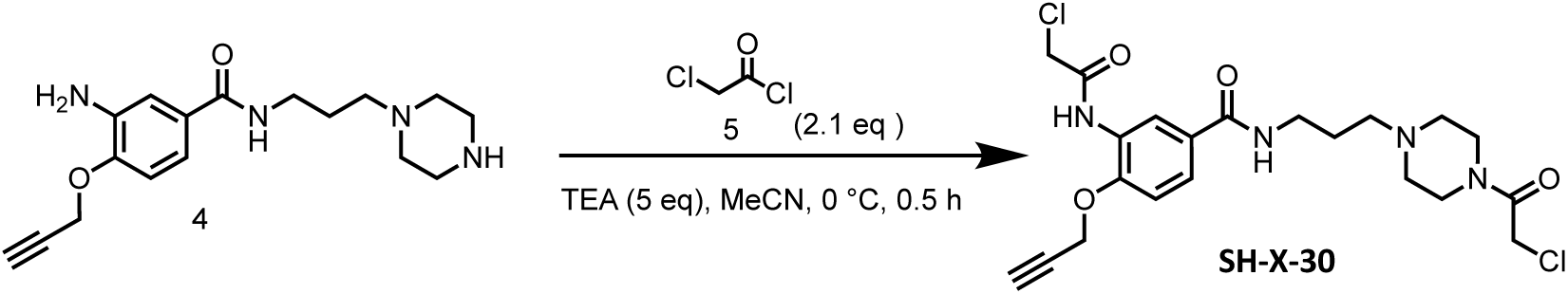

To a solution of 3-amino-*N*-(3-piperazin-1-ylpropyl)-4-prop-2-ynoxy-benzamide (500 mg, 1.42 mmol, HCl salt) in acetonitrile (5 mL) was added triethylamine (716 mg, 7.09 mmol, 986 μL) and 2-chloroacetyl chloride (336 mg, 2.98 mmol, 237 μL) in MeCN (5 mL). The mixture was stirred at 0 °C for 0.5 hr. The reaction mixture was filtered and the filtrate was concentrated to give a residue. The residue was purified by preparative-HPLC (column: Welch Xtimate C18 150*25mm*5um; mobile phase: [water (HCl)-MeCN]; gradient: 5%-35% B over 10 min) to give 3-(2-chloroacetamido)-*N*-(3-(4-(2-chloroacetyl)piperazin-1-yl)propyl)-4-(prop-2-yn-1-yloxy)benzamide (200 mg, 426 μmol, 30.1% yield) as a yellow solid.

LC-MS: 469.1 [M+H]^+^ / Ret time: 0.282 min/ method\5-95AB_0.8min.lcm

^1^H NMR (400 MHz, MeOD-*d*_4_) *δ* 8.59 (s, 1H), 7.69 (d, *J* = 8.8 Hz, 1H), 7.26 (d, *J* = 8.8 Hz, 1H), 4.95 (s, 2H), 4.69 - 4.60 (m, 1H), 4.38 - 4.30 (m, 4H), 4.23 - 4.15 (m, 1H), 3.66 (d, *J* = 12.4 Hz, 3H), 3.52 (t, *J* = 6.4 Hz, 2H), 3.29 - 3.22 (m, 3H), 3.22 - 3.14 (m, 2H), 3.08 (s, 2H), 2.15 - 2.06 (m, 2H).

### Synthesis of 3-acrylamido-*N*-(3-(4-acryloylpiperazin-1-yl)propyl)-4-(prop-2-yn-1-yloxy)benzamide (SH-X-32)

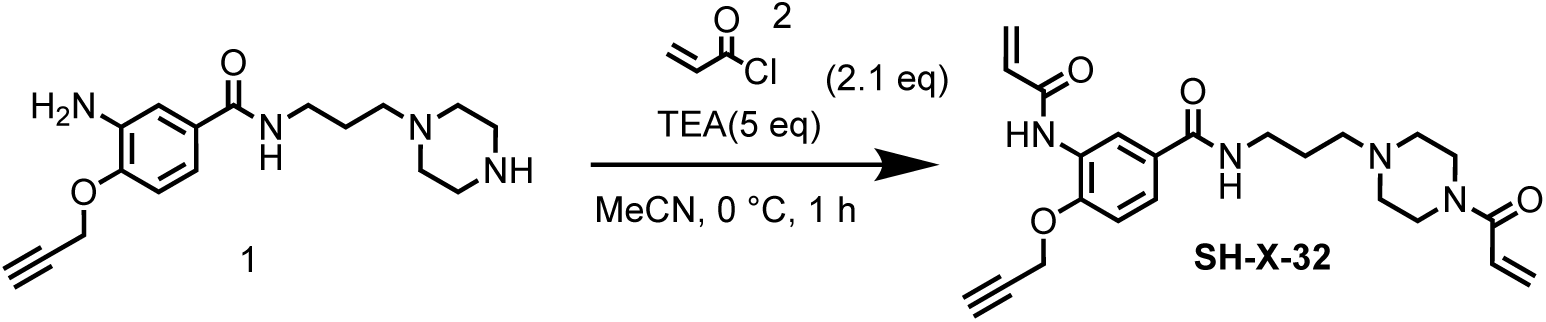

To a solution of 3-amino-*N*-(3-(piperazin-1-yl)propyl)-4-(prop-2-yn-1-yloxy)benzamide (600 mg, 1.70 mmol, HCl salt) in MeCN (6 mL) was added TEA (860 mg, 8.50 mmol, 1.18 mL) and acryloyl chloride (323 mg, 3.57 mmol, 290 μL) in at 0 °C. The mixture was stirred at 0 °C for 0.5 hr. LCMS showed the reaction was completed. The reaction mixture was filtered and the filtrate was concentrated to give a residue. The residue was purified by preparative-HPLC (column: Phenomenex luna C18 150*25mm* 10um; mobile phase: [water(FA)-MeCN]; gradient: 0%-28% B over 10 min) to give 3-acrylamido-*N*-(3-(4-acryloylpiperazin-1-yl)propyl)-4-(prop-2-yn-1-yloxy)benzamide (63.5 mg, 143 μmol, 8.44% yield, 95.9% purity) as a white solid.

LC-MS: 425.2 [M+H]^+^ / Ret time: 0.266 min / method: 5-95CD_1min.lcm.

^1^H NMR (400 MHz, MeOD-*d*_4_): *δ* 8.53 (d, *J* = 2.0 Hz, 1H), 8.21 (s, 1H), 7.65 (dd, *J* = 2.0, 8.4 Hz, 1H), 7.24 (d, *J* = 8.8 Hz, 1H), 6.75 (dd, *J* = 10.4, 16.8 Hz, 1H), 6.64-6.54 (m, 1H), 6.39 (dd, *J* = 1.6, 17.2 Hz, 1H), 6.22 (dd, *J* = 2.0, 16.8 Hz, 1H), 5.82-5.74 (m, 2H), 4.93 (d, *J* = 2.4 Hz, 2H), 3.79-3.72 (m, 4H), 3.47 (t, *J* = 6.8 Hz, 2H), 3.05 (t, *J* = 2.4 Hz, 1H), 2.82-2.71 (m, 6H), 1.97-1.87 (m, 2H)

### Synthesis of 5-acrylamido-*N*-(1-acryloylazepan-3-yl)-2-((1-(2-(2-(6-((4*R*,5*S*)-5-methyl-2-oxoimidazolidin-4-yl)hexanamido)ethoxy)ethyl)-1*H*-1,2,3-triazol-4-yl)methoxy)benzamide (SH-X-42)

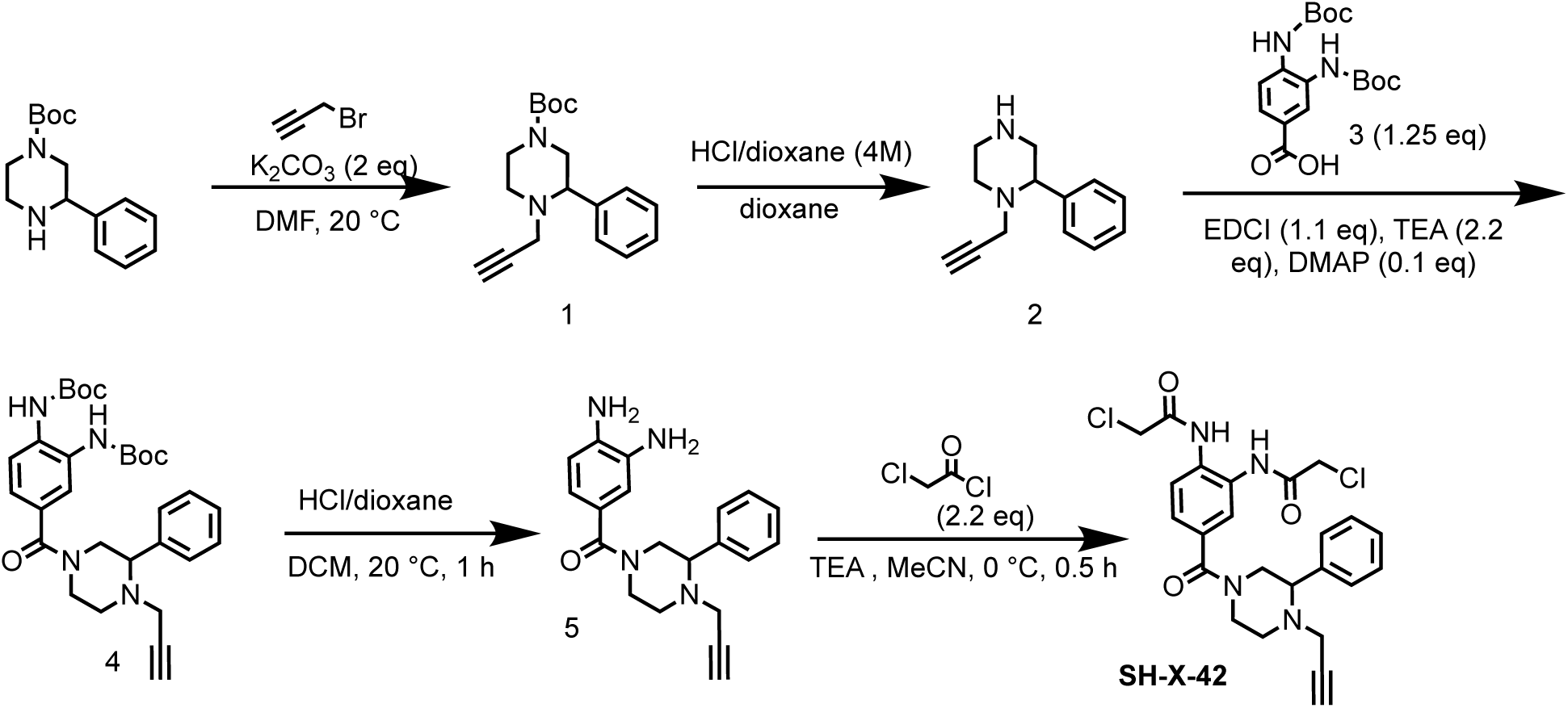

#### Step 1: Preparation of *tert*-butyl 3-phenyl-4-(prop-2-yn-1-yl)piperazine-1-carboxylate (1)

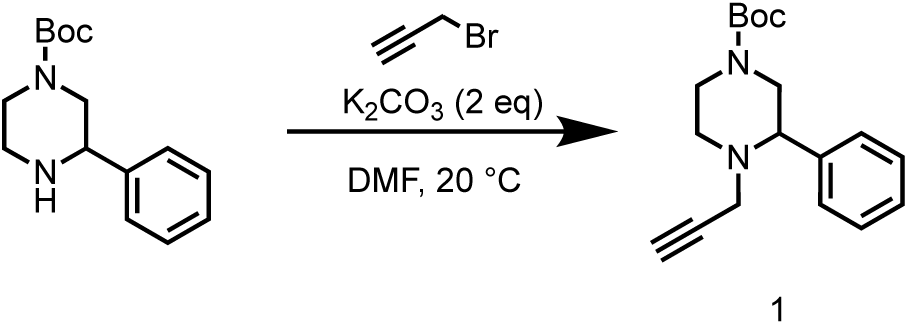

To a solution of *tert*-butyl 3-phenylpiperazine-1-carboxylate (3.00 g, 11.4 mmol) in DMF (30 mL) was added K_2_CO_3_ (4.74 g, 34.3 mmol), the reaction mixture was stirred at 20 °C for 0.5 hr. Then 3-bromoprop-1-yne (2.21 g, 14.8 mmol, 1.60 mL) was added into the reaction mixture, the mixture was stirred at 50 °C for 2 hrs. LCMS showed the reaction was completed. The reaction mixture was poured into H_2_O (200 mL), extracted with EtOAc (200 mL×3). The organic layers were collected and washed with brine (100 mL×3), dried over Na_2_SO_4_ and concentrated to give a residue. The residue was purified by column chromatography on silica gel (eluted with petroleum ether: ethyl acetate=10:0 to 10:1) to give *tert*-butyl 3-phenyl-4-(prop-2-yn-1-yl)piperazine-1-carboxylate (2.10 g, 6.99 mmol, 61.1% yield) as a yellow oil.

LC-MS: 301.2 [M+H]^+^ / Ret time: 0.443 min / method: 5-95AB_0.8min.lcm

#### Step 2: Preparation of 2-phenyl-1-(prop-2-yn-1-yl)piperazine (2)

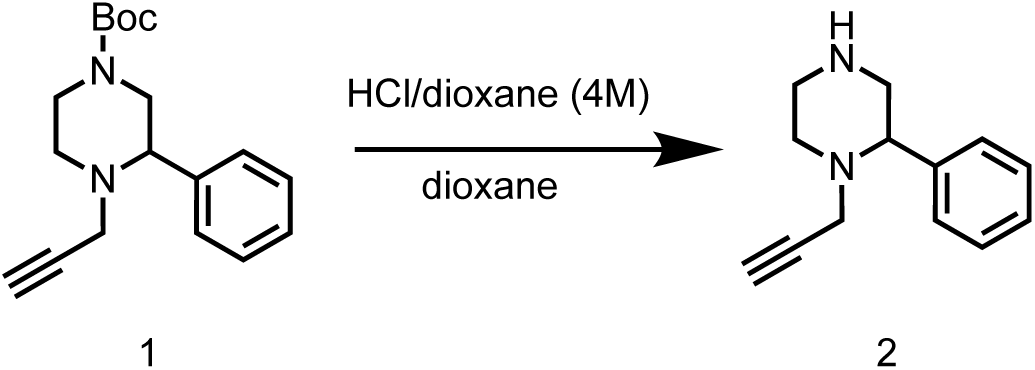

To a solution of *tert*-butyl 3-phenyl-4-(prop-2-yn-1-yl)piperazine-1-carboxylate (2.10 g, 6.99 mmol) in dioxane (1 mL) was added HCl/dioxane (4 M, 20 mL), the reaction mixture was stirred at 20 °C for 1 hr. LCMS showed the reaction was completed. The reaction mixture was concentrated in vacuum to give 2-phenyl-1-(prop-2-yn-1-yl)piperazine (3.10 g, crude, HCl salt) as a white solid.

LC-MS: 201.2 [M+H]^+^ / Ret time: 0.607 min / method: 0-60CD_1min.lcm

^1^H NMR (400 MHz, DMSO-*d*_6_): *δ* 9.66-9.06 (m, 1H), 7.42 (s, 4H), 4.73-4.50 (m, 2H), 4.05-3.78 (m, 1H), 3.45-3.35 (m, 2H), 3.26 (s, 2H), 3.09 (s, 3H).

#### Step 3: Preparation of di-tert-butyl (4-(3-phenyl-4-(prop-2-yn-1-yl)piperazine-1-carbonyl)-1,2-phenylene)dicarbamate (4)

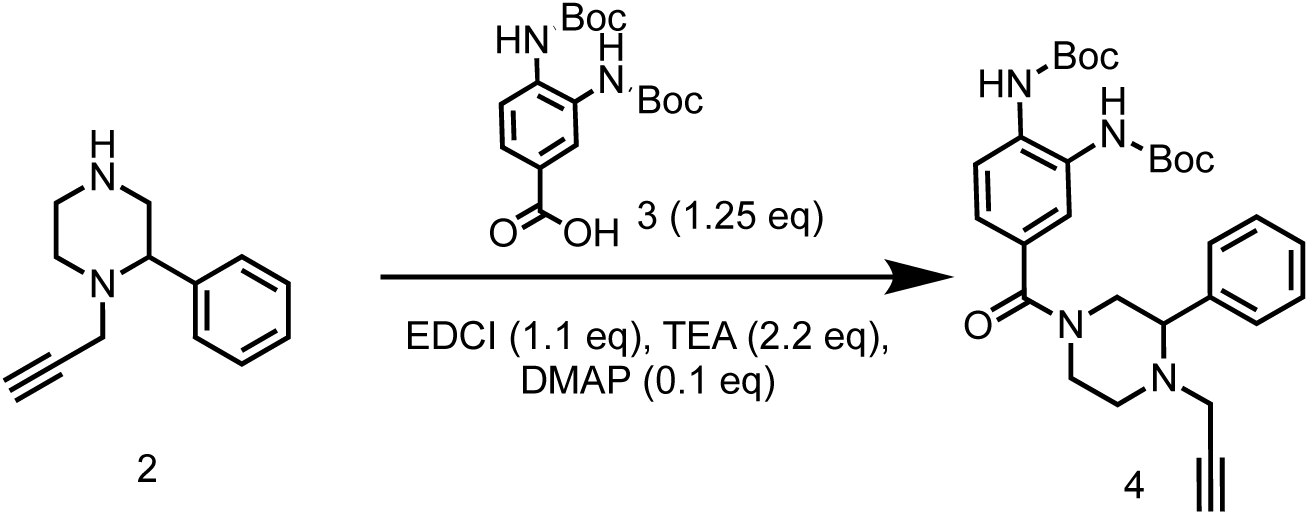

To a solution of 3,4-bis((*tert*-butoxycarbonyl)amino)benzoic acid (1.7 g, 4.82 mmol) in DCM (20 mL) was added EDCI (1.39 g, 7.24 mmol) and DMAP (58.9 mg, 482 μmol). Then a mixture of 2-phenyl-1-(prop-2-yn-1-yl)piperazine (914 mg, 3.86 mmol, HCl salt) and TEA (1.46 g, 14.5 mmol, 2.01 mL) in DCM (10 mL) was added into the reaction mixture, then stirred at 20 °C for 1 hr. LCMS showed the reaction was completed. The reaction mixture was concentrated in vacuum to give a residue. The residue was purified by column chromatography on silica gel (eluted with petroleum ether: ethyl acetate=5:1 to 1:1) to give di-*tert*-butyl (4-(3-phenyl-4-(prop-2-yn-1-yl)piperazine-1-carbonyl)-1,2-phenylene) dicarbamate (1.57 g, 2.94 mmol, 60.9% yield) as a yellow oil.

LC-MS: 535.3 [M+H]^+^ / Ret time: 0.422 min / method: 5-95AB_0.8min.lcm

^1^H NMR (400 MHz, DMSO-*d*_6_): *δ* 8.66 (s, 2H), 7.60 (s, 2H), 7.36 (s, 5H), 7.13 (d, *J* = 8.0 Hz, 1H), 4.58-4.15 (m, 1H), 3.81-3.48 (m, 1H), 3.39 (dd, *J* = 2.8, 10.8 Hz, 2H), 3.25 (d, *J* = 18.0 Hz, 1H), 3.18-3.15 (m, 1H), 3.00 (d, *J* = 2.4 Hz, 2H), 2.92-2.77 (m, 1H), 2.60-2.51 (m, 1H), 1.51-1.45 (m, 18H).

#### Step 4: Preparation of (3,4-diaminophenyl)(3-phenyl-4-(prop-2-yn-1-yl)piperazin-1-yl)methanone (5)

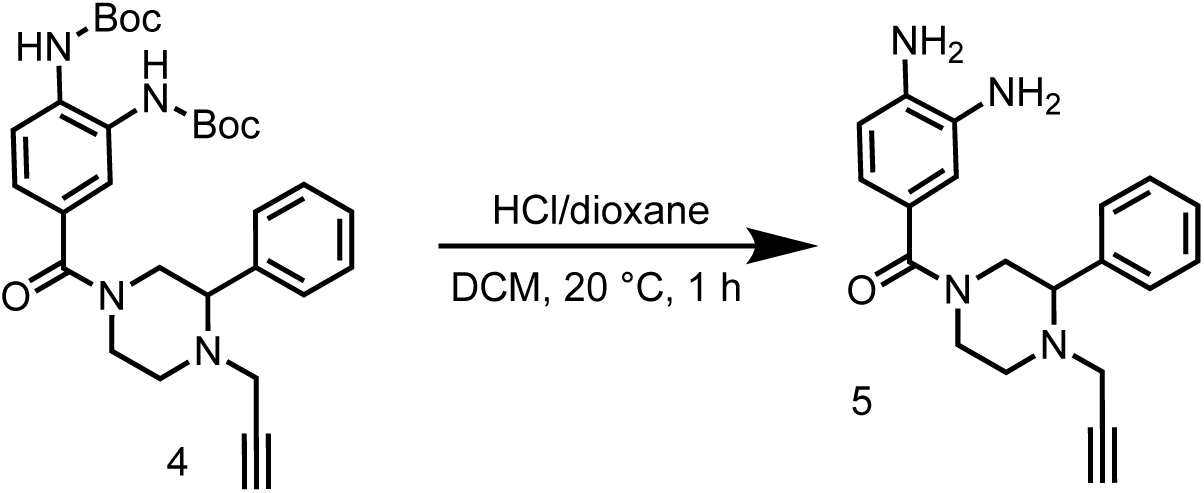

A mixture of di-*tert*-butyl (4-(3-phenyl-4-(prop-2-yn-1-yl)piperazine-1-carbonyl)-1,2-phenylene)dicarbamate (700 mg, 1.31 mmol) in HCl/dioxane (4 M, 7.00 mL) was stirred at 20 °C for 1 hr. LCMS showed the reaction was completed. The mixture was concentrated in vacuum to give a residue. The residue was triturated with DCM (20 mL), then filtered and the filter cake was concentrated in vacuum to give (3,4-diaminophenyl)(3-phenyl-4-(prop-2-yn-1-yl)piperazin-1-yl)methanone (528 mg, crude, HCl salt) as a yellow solid.

LC-MS: 335.3 [M+H]^+^ / Ret time: 0.305 min / method: 0-60AB_0.8min.lcm

#### Step 5: Preparation of *N,N*’-(4-(3-phenyl-4-(prop-2-yn-1-yl)piperazine-1-carbonyl)-1,2-phenylene)bis(2-chloroacetamide) (SH-X-042)

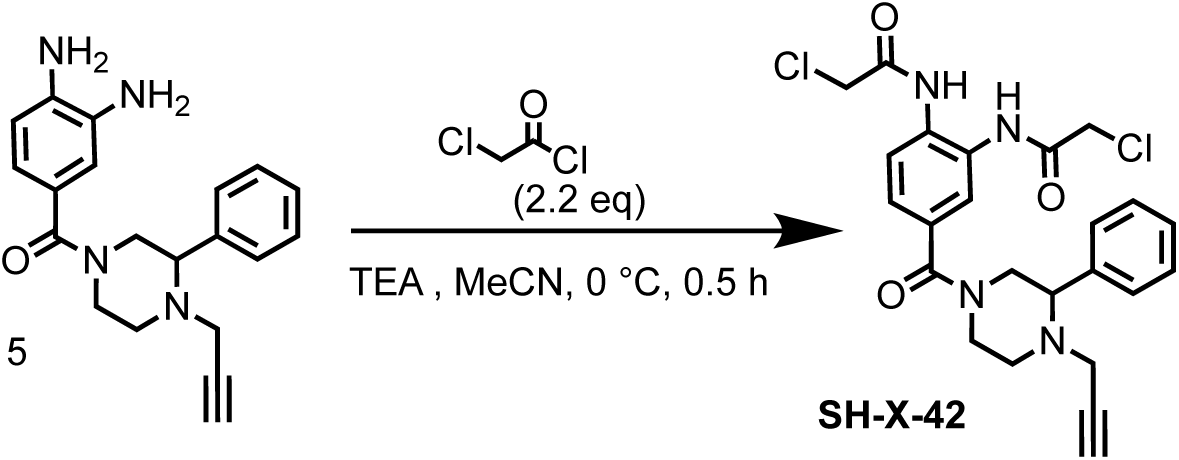

To a solution of 2-chloroacetyl chloride (335 mg, 2.97 mmol, 236 μL) in MeCN (10 mL) was added into a mixture of (3,4-diaminophenyl)(3-phenyl-4-(prop-2-yn-1-yl)piperazin-1-yl)methanone (500 mg, 1.35 mmol, HCl salt) and TEA (682 mg, 6.74 mmol, 938 μL) in MeCN (20 mL), then the reaction mixture was stirred at 0 °C for 0.5 hr. LCMS showed the reaction was completed. The reaction mixture was quenched with H_2_O (0.1 mL) and concentrated in vacuum to give a residue. The residue was purified by preparative-HPLC (column: Phenomenex luna C18 150*25mm* 10um; mobile phase: [water (FA)-MeCN]; gradient: 29%-59% B over 10 min). The purified solution was lyophilized to give *N,N’*-(4-(3-phenyl-4-(prop-2-yn-1-yl)piperazine-1-carbonyl)-1,2-phenylene)bis(2-chloroacetamide) (410 mg, 841 μmol, 62.4% yield) as a yellow oil.

LC-MS: 487.1 [M+H]^+^ / Ret time: 0.346 min / method: 5-95AB_0.8min.lcm

^1^H NMR (400 MHz, DMSO-*d*_6_) *δ* 9.81 (s, 2H), 7.65 (d, J = 0.8 Hz, 2H), 7.47 - 7.21 (m, 6H), 4.63 - 4.38 (m, 1H), 4.34 (s, 4H), 3.86 - 3.61 (m, 1H), 3.37 (d, J = 10.8 Hz, 2H), 3.26 (d, J = 18.0 Hz, 1H), 3.17 (t, J = 2.0 Hz, 1H), 3.09 - 2.96 (m, 2H), 2.95 - 2.79 (m, 1H), 2.53 (d, J = 2.4 Hz, 1H).

### Synthesis of *N,N*’-(4-(5-(prop-2-yn-1-ylcarbamoyl)indoline-1-carbonyl)-1,2-phenylene)bis(2-chloroacetamide) (SH-X-46)

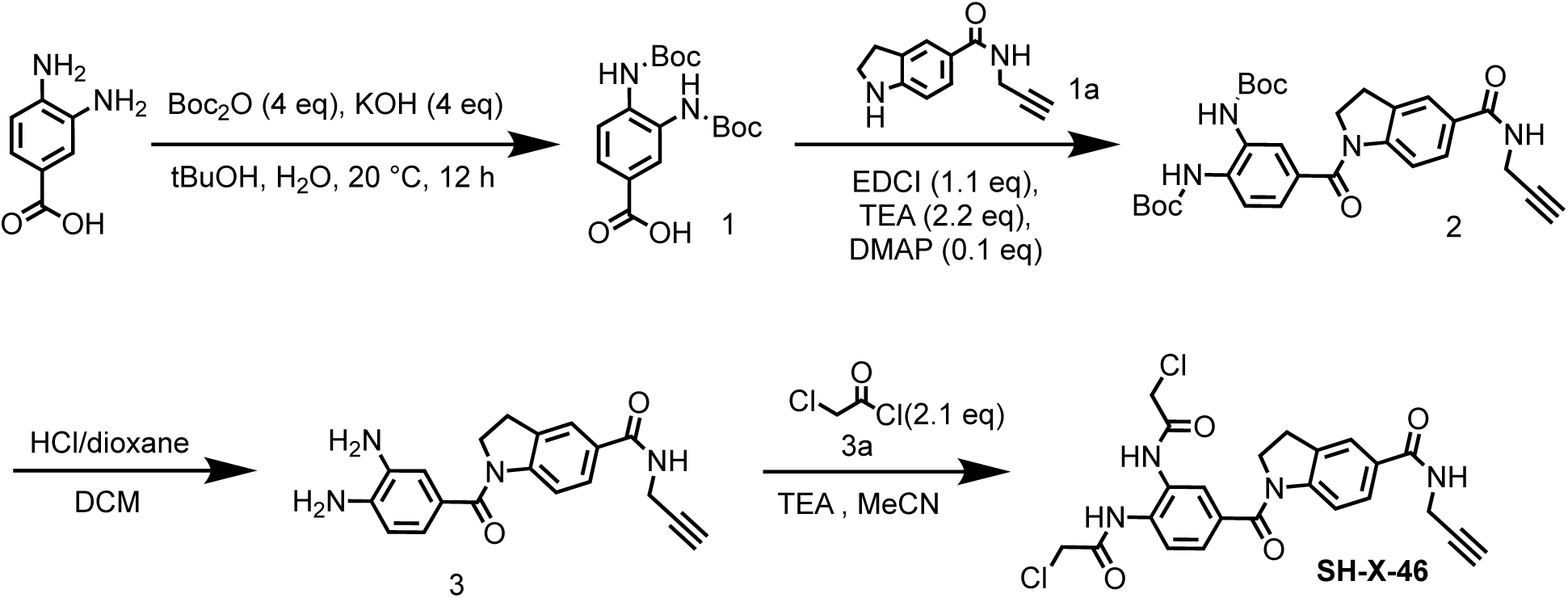

#### Step 1: Preparation of 3, 4-bis((*tert*-butoxycarbonyl)amino)benzoic acid (1)

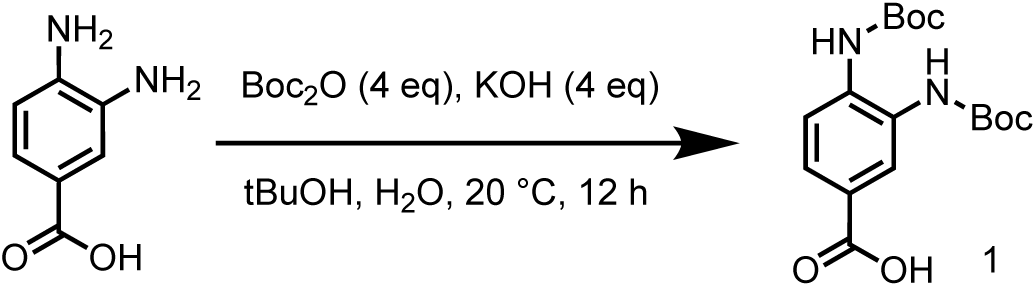

To a solution of 3,4-diaminobenzoic acid (3.00 g, 19.7 mmol) and KOH (4.43 g, 78.9 mmol) in H_2_O (60 mL) and *tert*-butyl alcohol (50 mL) was added Boc_2_O (17.2 g, 78.9 mmol, 18.1 mL), the mixture was stirred at 35 °C for 12 hrs. LCMS showed the reaction was completed. After the reaction was completed, the mixture was adjusted to pH=4 with citric acid, the mixture was filtered, and the filter cake was concentrated to give 3,4-bis((*tert*-butoxycarbonyl)amino)benzoic acid (7.00 g, crude) as brown solid.

LC-MS: 375.4 [M+Na]^+^ / Ret time: 0.433 min / method: 5-95AB_0.8min.

^1^H NMR (400 MHz, DMSO-*d*_6_): *δ* 13.00-12.59 (m, 1H), 8.85-8.62 (m, 1H), 8.09 (s, 1H), 7.76-7.68 (m, 1H), 7.67-7.61 (m, 1H), 1.50-1.47 (m, 18H).

#### Step 2: Preparation of di-tert-butyl (4-(5-(prop-2-yn-1-ylcarbamoyl)indoline-1-carbonyl)-1,2-phenylene)dicarbamate (2)

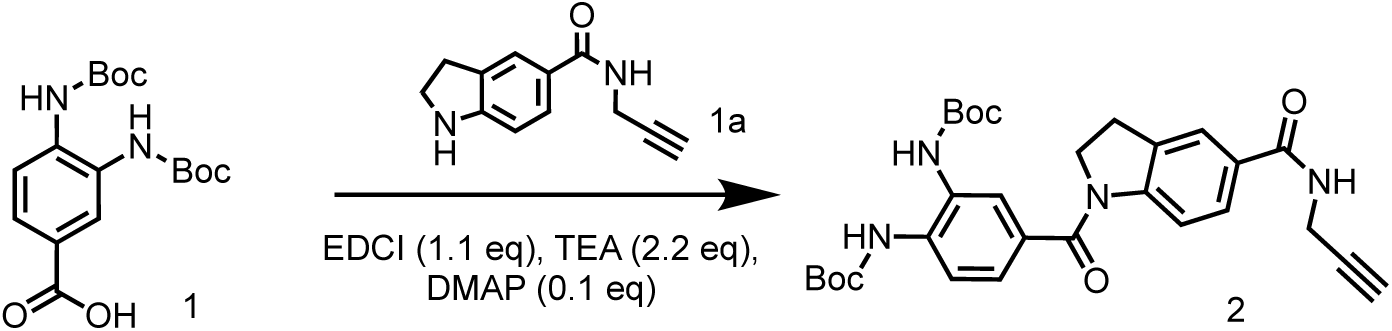

To a solution of 3,4-bis((*tert*-butoxycarbonyl)amino)benzoic acid (3.65 g, 10.4 mmol) and *N*-(prop-2-yn-1-yl)indoline-5-carboxamide (2.49 g, 12.4 mmol) in DCM (40 mL) was added EDCI (2.98 g, 15.5 mmol) and DMAP (127 mg, 1.04 mmol), the mixture was stirred at 20 °C for 2 hrs. LCMS showed the reaction was completed. The reaction mixture was concentrated in vacuum to give a residue. The residue was purified by column chromatography on silica gel (eluted with petroleum ether: ethyl acetate=1:1 to 1:1) to give di-*tert*-butyl(4-(5-(prop-2-yn-1-ylcarbamoyl)indoline-1-carbonyl)-1,2-phenylene)dicarbamate (3.20 g, 5.99 mmol, 57.8% yield) as a white solid.

LC-MS: 379.3 [M-156+H]^+^ / Ret time: 0.468 min / method: 5-95AB_0.8min.

^1^H NMR (400 MHz, DMSO-*d*_6_): *δ* 8.90-8.63 (m, 2H), 7.79-7.76 (m, 1H), 7.76-7.64 (m, 3H), 7.34 (d, *J* = 8.4 Hz, 1H), 4.12-4.06 (m, 2H), 4.05-4.02 (m, 2H), 3.19-3.07 (m, 3H), 1.49 (d, *J* = 8.4 Hz, 18H).

#### Step 3: Preparation of 1-(3,4-diaminobenzoyl)-N-(prop-2-yn-1-yl)indoline-5-carboxamide (3)

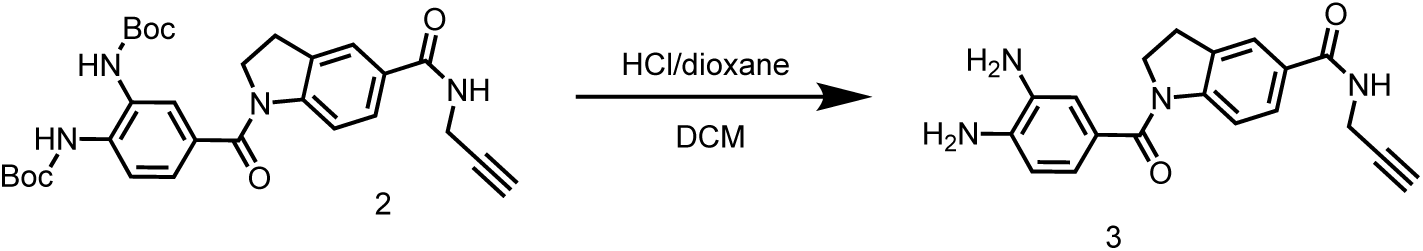

To a solution of di-*tert*-butyl(4-(5-(prop-2-yn-1-ylcarbamoyl)indoline-1-carbonyl)-1,2-phenylene)dicarbamate (3.20 g, 5.99 mmol) in dioxane (32 mL) was added HCl/dioxane (4 M, 32 mL). The mixture was stirred at 20 °C for 1 hr. LCMS showed the reaction was completed. The reaction mixture was concentrated in vacuum to give a residue. The residue was purified by preparative-HPLC (column: Phenomenex luna C18 150*25mm* 10um; mobile phase: [water(FA)-MeCN]; gradient: 0%-30% B over 15 min) to give 1-(3,4-diaminobenzoyl)-*N*-(prop-2-yn-1-yl)indoline-5-carboxamide (62.3 mg, 175 μmol, 2.93% yield, 94.0% purity) as a white solid.

LC-MS: 355.3 [M+H]^+^ / Ret time: 0.237 min / method: 5-95AB_0.8min.

^1^H NMR (400 MHz, DMSO-*d*_6_): *δ* 8.76 (m, *J* = 5.4 Hz, 1H), 8.20 (s, 1H), 7.74 (s, 1H), 7.65 (d, *J* = 8.8 Hz, 1H), 7.51 (d, *J* = 8.4 Hz, 1H), 6.81 (d, *J* = 2.0 Hz, 1H), 6.74 (m, *J* = 2.0, 7.9 Hz, 1H), 6.53 (d, *J* = 8.0 Hz, 1H), 5.40-4.46 (m, 2H), 4.11 (t, *J* = 8.4 Hz, 2H), 4.03 (m, *J* = 2.4, 2H), 3.14-3.05 (m, 3H).

#### Step 4: Preparation of *N,N*’-(4-(5-(prop-2-yn-1-ylcarbamoyl)indoline-1-carbonyl)-1,2-phenylene)bis(2-chloroacetamide) (SH-X-46)

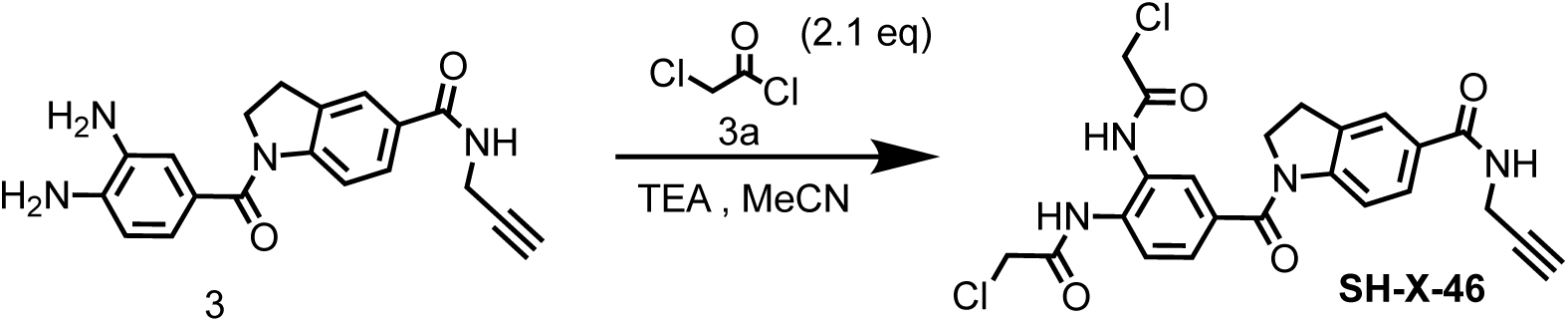

To a mixture of 1-(3,4-diaminobenzoyl)-*N*-(prop-2-yn-1-yl)indoline-5-carboxamide (500 mg, crude, HCl salt) and TEA (819 mg, 8.09 mmol, 1.13 mL) in MeCN (10 mL) was added 2-chloroacetyl chloride (335 mg, 2.97 mmol, 236 μL) and stirred at 20 °C for 0.5 hr. LCMS showed the reaction was completed. The reaction mixture was extracted with H_2_O (10 mL) and ethyl acetate (10 mL×3), washed with brine (10 mL), the organic phase was dried over Na_2_SO_4_, concentrated in vacuum to give *N,N’*-(4-(5-(prop-2-yn-1-ylcarbamoyl)indoline-1-carbonyl)-1,2-phenylene)bis(2-chloroacetamide) (134 mg, 264 μmol, 24.5% yield, 95.6% purity) as a brown oil.

LC-MS: 487.1 [M+H]^+^ / Ret time: 0.385 min / method: 5-95AB_0.8min.

^1^H NMR (400 MHz, DMSO-*d*_6_): *δ* 9.85 (s, 2H), 8.83 (s, 1H), 7.91-7.63 (m, 5H), 7.51 (d, *J* = 1.2 Hz, 1H), 4.38 (s, 2H), 4.35 (s, 2H), 4.14-4.08 (m, 2H), 4.04 (, *J* = 2.4, 2H), 3.18-3.10 (m, 3H).

### Synthesis of N,N’-(4-(5-(prop-2-yn-1-ylcarbamoyl)indoline-1-carbonyl)-1,2-phenylene) diacrylamide (SH-X-48)

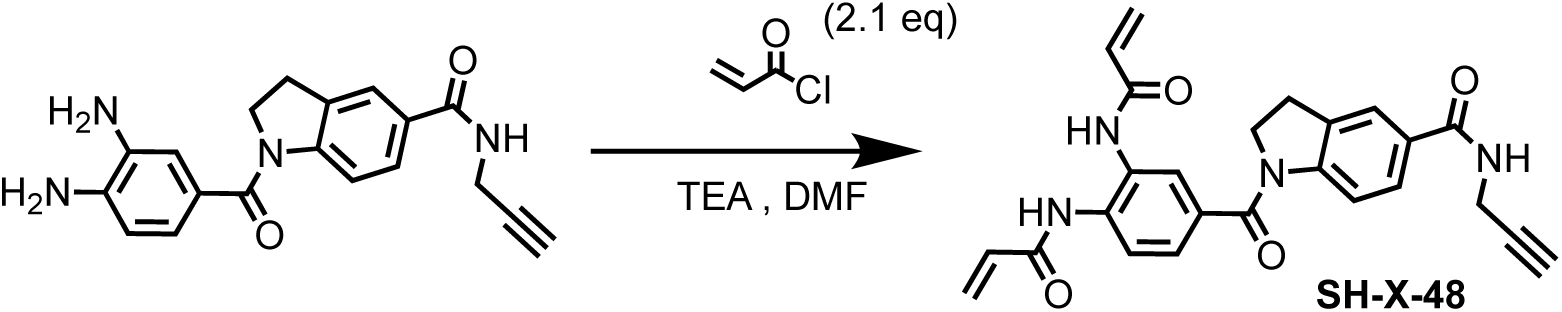

To a solution of 1-(3,4-diaminobenzoyl)-*N*-(prop-2-yn-1-yl)indoline-5-carboxamide (400 mg, 1.08 mmol, HCl salt) in MeCN (40 mL) was added TEA (655 mg, 6.47 mmol, 901 μL), then acryloyl chloride (215 mg, 2.37 mmol, 193 μL) was added into the reaction mixture and stirred at 20 °C for 0.5 hr. LCMS showed the reaction was completed. The reaction mixture was extracted with H_2_O (5 mL) and ethyl acetate (5 mL×3), washed with brine (5 mL), and the organic phase was dried over Na_2_SO_4_, concentrated in vacuum to give a residue. The residue was purified by preparative-HPLC (column: Phenomenex luna C18 150*25mm* 10um; mobile phase: [water(FA)-MeCN]; gradient: 30%-60% B over 10 min) to give *N,N’*-(4-(5-(prop-2-yn-1-ylcarbamoyl)indoline-1-carbonyl)-1,2-phenylene)diacrylamide (63.4 mg, 142 μmol, 13.2% yield, 99.4% purity) as a white solid.

LC-MS: 443.1 [M+H]^+^ / Ret time: 0.370 min / method: 5-95AB_0.8min.lcm.

^1^H NMR (400 MHz, DMSO-*d*_6_): *δ* 9.78 (s, 2H), 8.83 (t, *J* = 5.2 Hz, 1H), 8.00-7.67 (m, 5H), 7.45 (m, *J* = 8.4 Hz, 1H), 6.53 (m, *J* = 10.4 Hz, 2H), 6.33-6.25 (m, 2H), 5.81 (m, *J* = 10.0 Hz, 2H), 4.13 (t, *J* = 8.4 Hz, 2H), 4.04 (m, *J* = 2.4 Hz, 2H), 3.20-3.10 (m, 3H).

### Synthesis of *N,N*’-((2-((prop-2-yn-1-yloxy)methyl)piperazine-1,4-dicarbonyl)bis(pyridine-6,3-diyl))bis(2-chloroacetamide) (SH-X-59)

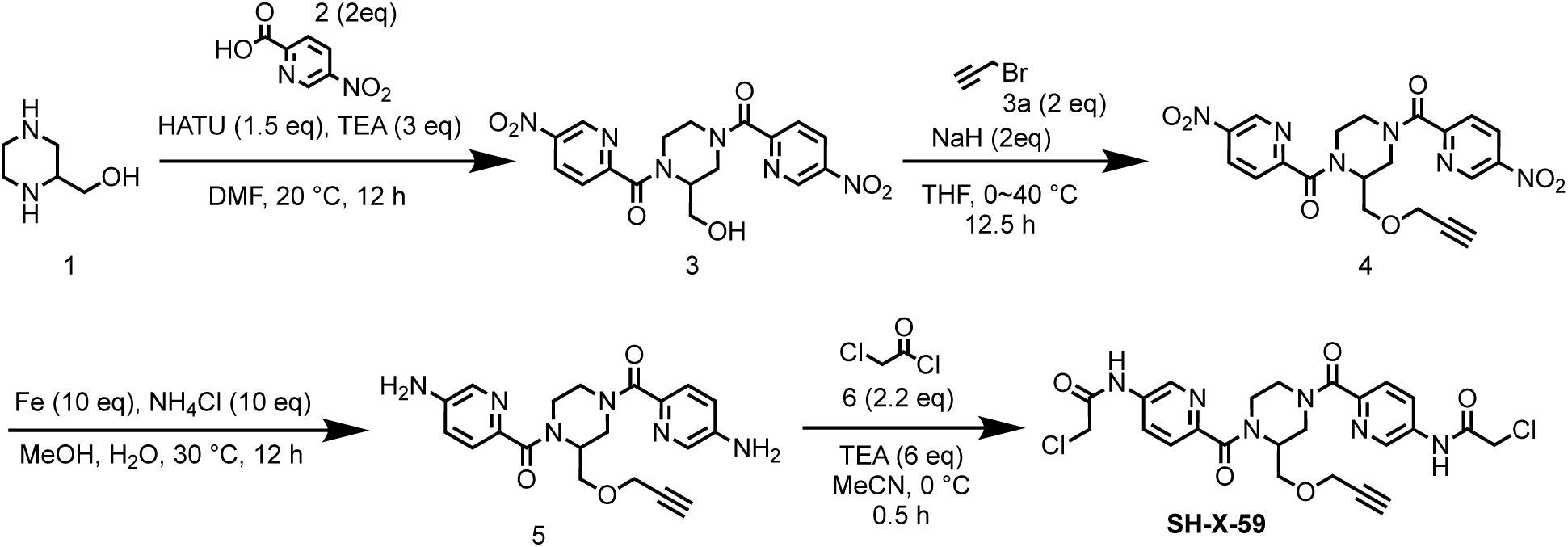

#### Step 1: Preparation of (2-(hydroxymethyl)piperazine-1,4-diyl)bis((5-nitropyridin-2-yl)methanone) (3)

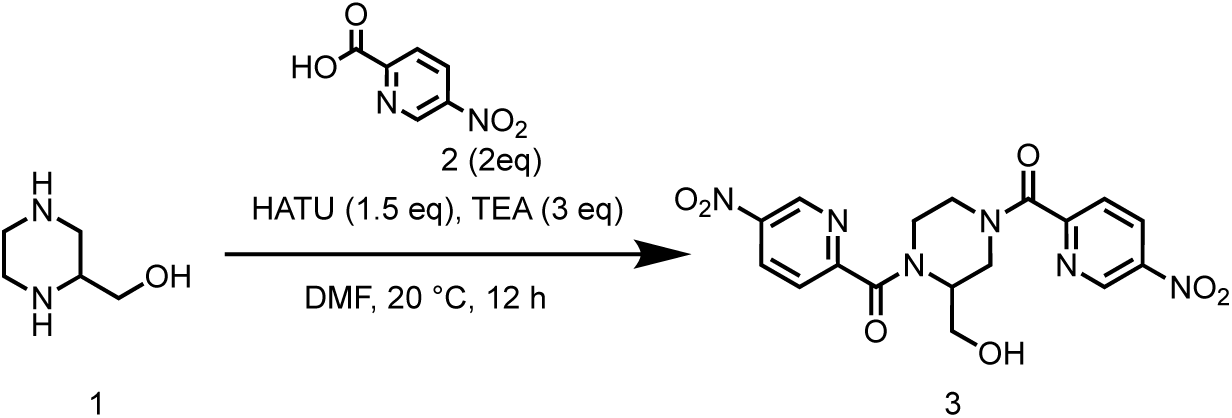

To a solution of 5-nitropicolinic acid (4.34 g, 25.8 mmol) in DMF (50 mL) was added HATU (7.36 g, 19.4 mmol) and TEA (3.92 g, 38.7 mmol, 5.39 mL), then piperazin-2-ylmethanol (1.5 g, 12.9 mmol) was added into the reaction mixture. The reaction mixture was stirred at 20 °C for 12 hrs. LCMS showed the reaction was completed. The reaction mixture was poured into H_2_O (300 mL) and extracted with EtOAc (200 mL×3). The organic layers were collected and washed with brine (300 mL×3), dried over Na_2_SO_4_ and concentrated to give a residue. The residue was purified by column chromatography on silica gel (eluted with petroleum ether: ethyl acetate=1:1 to 0:1) to give (2-(hydroxymethyl)piperazine-1,4-diyl)bis((5-nitropyridin-2-yl)methanone) (2.81 g, 6.75 mmol, 52.3% yield) as a white solid.

LC-MS: 417.1 [M+H]^+^ / Ret time: 0.373 min / method: 0-60AB_0.8min.lcm

#### Step 2: Preparation of (2-((prop-2-yn-1-yloxy)methyl)piperazine-1,4-diyl)bis((5-nitropyridin-2-yl)methanone) (4)

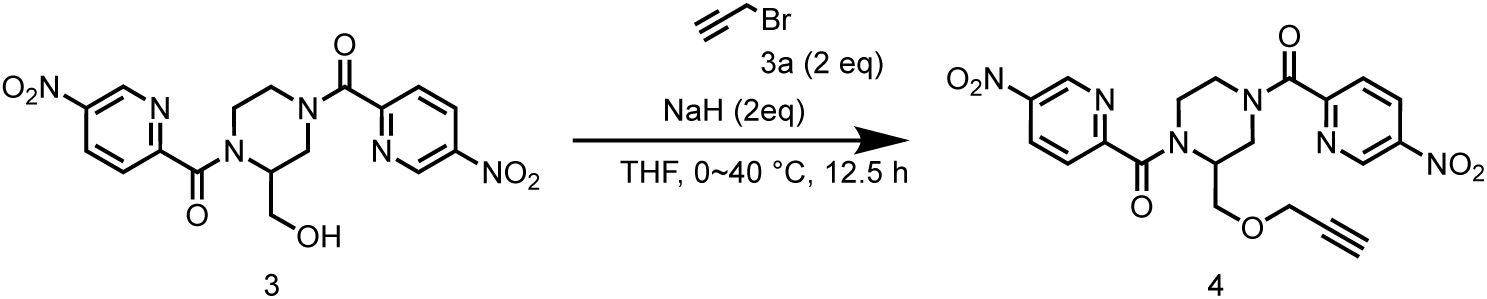

To a solution of (2-(hydroxymethyl)piperazine-1,4-diyl)bis((5-nitropyridin-2-yl)methanone) (2.80 g, 6.73 mmol) in THF (50 mL) was added NaH (538 mg, 13.5 mmol, 60% purity) at 0 °C. The reaction mixture was stirred at 0 °C for 0.5 hr. Then 3-bromoprop-1-yne (10.0 g, 67.3 mmol, 7.25 mL) was added into the reaction mixture at 0 °C, and the mixture was stirred at 40 °C for 12 hrs. LCMS showed the reaction was completed. The reaction mixture was quenched with saturated NH_4_Cl (aq), then extracted with EtOAc (100 mL×3). The organic layers were collected and washed with brine (100 mL×3), dried over Na_2_SO_4_ and concentrated to give a residue. The residue was purified by column chromatography on silica gel (eluted with petroleum ether: ethyl acetate=1:1 to 0:1) to give (2-((prop-2-yn-1-yloxy)methyl)piperazine-1,4-diyl)bis((5-nitropyridin-2-yl)methanone) (800 mg, 1.76 mmol, 26.28% yield) as a yellow solid.

LC-MS: 455.3 [M+H]^+^ / Ret time: 0.348 min / method: 5-95AB_0.8min.lcm.

#### Step 3: Preparation of (2-((prop-2-yn-1-yloxy)methyl)piperazine-1,4-diyl)bis((5-aminopyridin-2-yl)methanone) (5)

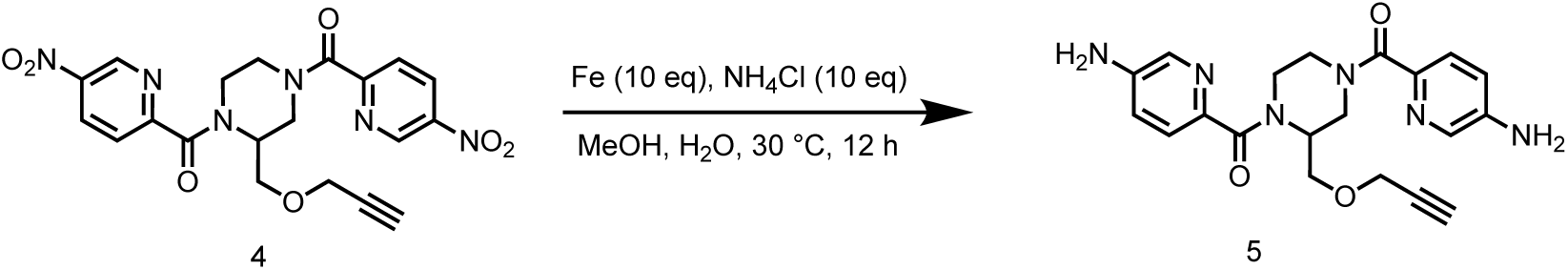

To a solution of (2-((prop-2-yn-1-yloxy)methyl)piperazine-1,4-diyl)bis((5-nitropyridin-2-yl)methanone) (800 mg, 1.76 mmol) in MeOH (20 mL) and H_2_O (0.2 mL) was added NH_4_Cl (942 mg, 17.6 mmol) and Fe (983 mg, 17.6 mmol). The reaction mixture was stirred at 30 °C for 12 hrs. LCMS showed the reaction was completed. The reaction mixture was filtered through celite pad, and the filtrate was concentrated to give (2-((prop-2-yn-1-yloxy)methyl)piperazine-1,4-diyl)bis((5-aminopyridin-2-yl)methanone) (690 mg, 1.75 mmol, 99.4% yield) as yellow solid.

LC-MS: 395.3 [M+H]^+^ / Ret time: 0.195 min / method: 5-95AB_0.8min.lcm

#### Step 4: Preparation of *N,N*’-((2-((prop-2-yn-1-yloxy)methyl)piperazine-1,4-dicarbonyl)bis(pyridine-6,3-diyl))bis(2-chloroacetamide) (SH-X-59)

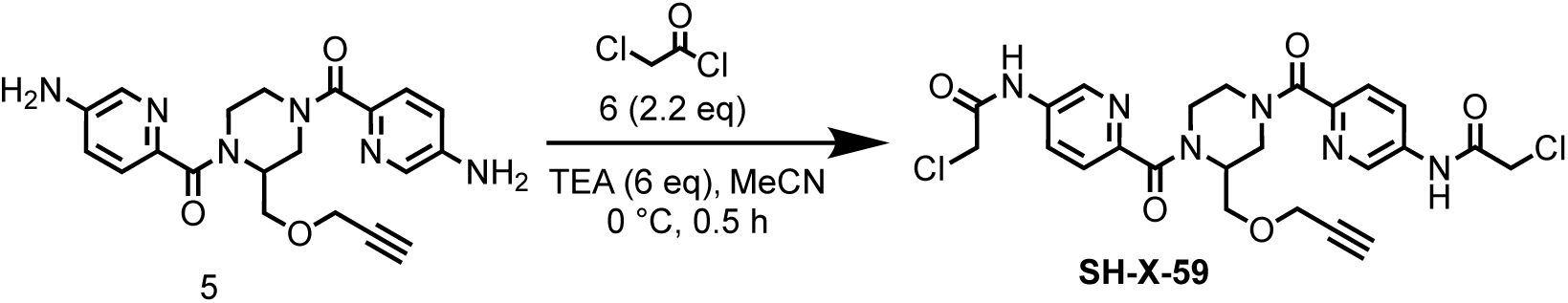

To a solution of (2-((prop-2-yn-1-yloxy)methyl)piperazine-1,4-diyl)bis((5-aminopyridin-2-yl)methanone) (480 mg, 1.22 mmol) in MeCN (10 mL) was added TEA (739 mg, 7.30 mmol, 1.02 mL), then 2-chloroacetyl chloride (302 mg, 2.68 mmol, 213 μL) was added into the reaction mixture at 0 °C, the mixture was stirred at 0 °C for 0.5 hr. LCMS showed the reaction was completed. The reaction mixture was quenched with H_2_O (1 mL) to give a residue. The residue was purified by preparative-HPLC (column: Phenomenex Luna C18 200*40mm*10um; mobile phase: [water (TFA)-MeCN]; gradient: 10%-40% B over 13 min). The purified solution was lyophilized to give *N*,*N*’-((2-((prop-2-yn-1-yloxy)methyl)piperazine-1,4-dicarbonyl)bis(pyridine-6,3-diyl))bis(2-chloroacetamide) (70 mg, 128 μmol, 10.5% yield) as a yellow solid.

LC-MS: 546.9 [M+H]^+^ / Ret time: 0.309 min / method: 5-95AB_0.8min.lcm

^1^H NMR (400 MHz, DMSO-*d*_6_) *δ* 10.81 - 10.71 (m, 2H), 8.74 (d, *J* = 19.6 Hz, 2H), 8.15 (d, *J* = 4.8 Hz, 2H), 7.71 - 7.58 (m, 2H), 4.55 (s, 2H), 4.34 (s, 4H), 4.28 - 4.19 (m, 1H), 4.07 - 3.96 (m, 2H), 3.92 - 3.76 (m, 2H), 3.40 - 3.31 (m, 2H), 3.30 - 3.18 (m, 1H), 3.16 - 2.92 (m, 2H).

### Synthesis of 5-(2-chloroacetamido)-*N*-((3*R*,4*S*)-1-(2-chloroacetyl)-3-(prop-2-yn-1-ylcarbamoyl)piperidin-4-yl)picolinamide (SH-X-61A) and 5-(2-chloroacetamido)-*N*-((3*R*,4*R*)-1-(2-chloroacetyl)-3-(prop-2-yn-1-ylcarbamoyl)piperidin-4-yl)picolinamide (SH-X-61B)

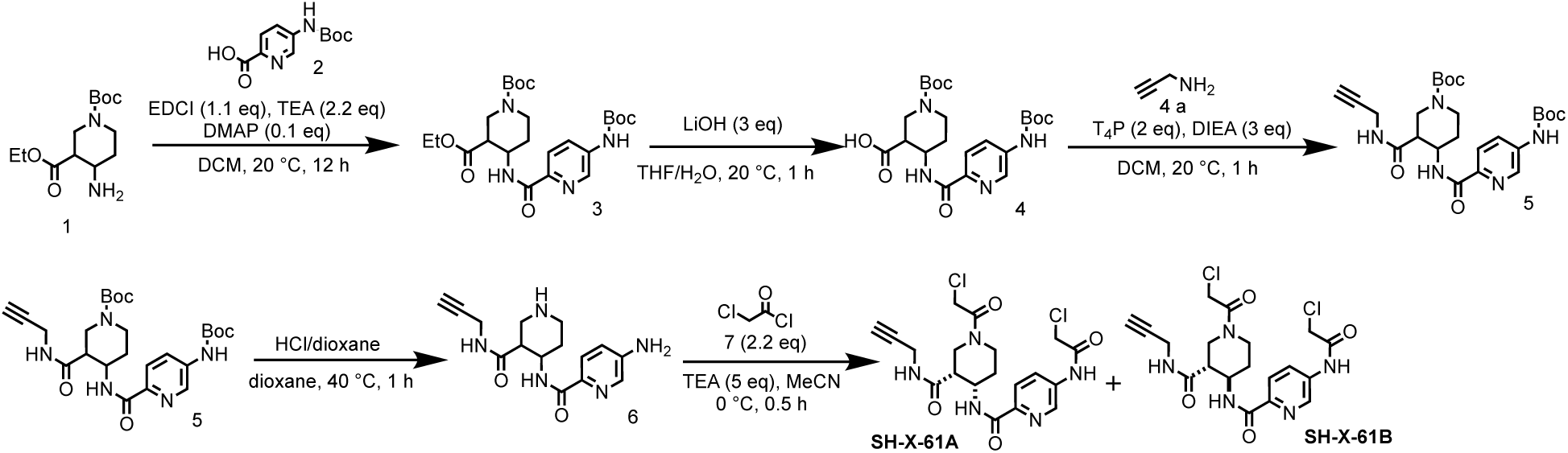

#### Step 1: Preparation of 1-(*tert*-butyl) 3-ethyl 4-(5-((*tert*-butoxycarbonyl) amino) picolinamido)piperidine-1,3-dicarboxylate (3)

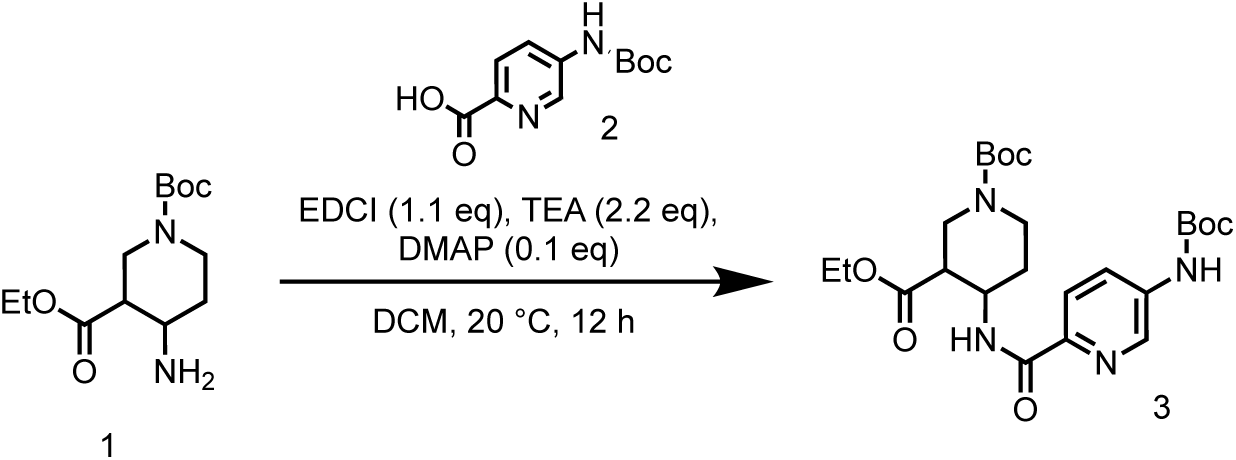

To a solution of 1-(*tert*-butyl) 3-ethyl 4-(5-((*tert*-butoxycarbonyl) amino) picolinamido) piperidine-1,3-dicarboxylate (1.20 g, 2.44 mmol) in THF (5 mL) and H_2_O (5 mL) was added LiOH•H_2_O (306 mg, 7.31 mmol). The mixture was stirred at 20 °C for 1 hr. LCMS showed the reaction was completed. The reaction mixture was concentrated in vacuum. Then the mixture was acidified with 1 M HCl to pH=4. The suspension was filtered and the filter cake was washed with methanol (10 mL) and concentrated in vacuum to give 1-(*tert*-butoxycarbonyl)-4-(5-((*tert*-butoxycarbonyl)amino) picolinamido)piperidine-3-carboxylic acid (1.15 g, crude) as a white solid.

LC-MS: 439.2 [M+H]^+^ / Ret time: 0.516 min / method: 5-95AB_0.8min.lcm

#### Step 2: Preparation of 1-(*tert*-butoxycarbonyl)-4-(5-((*tert*-butoxycarbonyl)amino) picolinamido)piperidine-3-carboxylic acid (4)

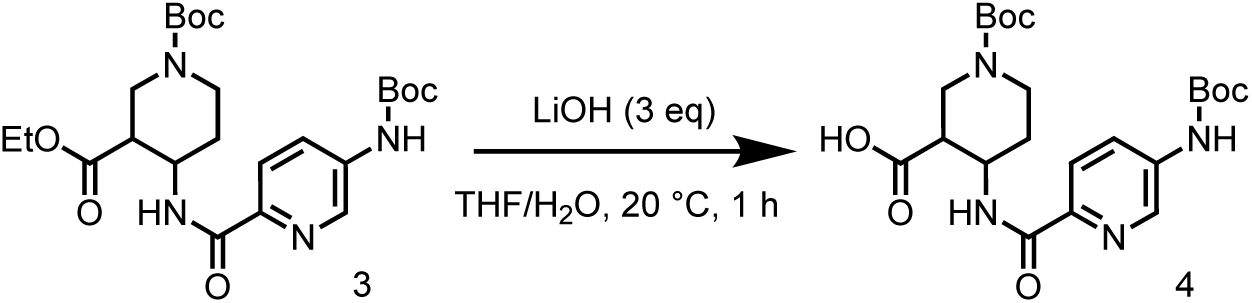

To a solution of 1-(*tert*-butyl) 3-ethyl 4-(5-((*tert*-butoxycarbonyl)amino)picolinamido)piperidine-1,3-dicarboxylate (1.20 g, 2.44 mmol) in THF (5 mL) and H_2_O (5 mL) was added LiOH•H_2_O (306 mg, 7.31 mmol). The mixture was stirred at 20 °C for 1 hr. LCMS showed the reaction was completed. The reaction mixture was concentrated in vacuum. Then the mixture was acidified with 1 M HCl to pH=4. The suspension was filtered and the filter cake was washed with methanol (10 mL) and concentrated in vacuum to give 1-(*tert*-butoxycarbonyl)-4-(5-((*tert*-butoxycarbonyl)amino)picolinamido)piperidine-3-carboxylic acid (1.15 g, crude) as a white solid.

LC-MS: 465.2 [M+H]^+^ / Ret time: 0.456 min / method: 5-95AB_0.8min.lcm.

#### Step 3: Preparation of *tert*-butyl 4-(5-((*tert*-butoxycarbonyl)amino)picolinamido)-3-(prop-2-yn-1-ylcarbamoyl)piperidine-1-carboxylate (5)

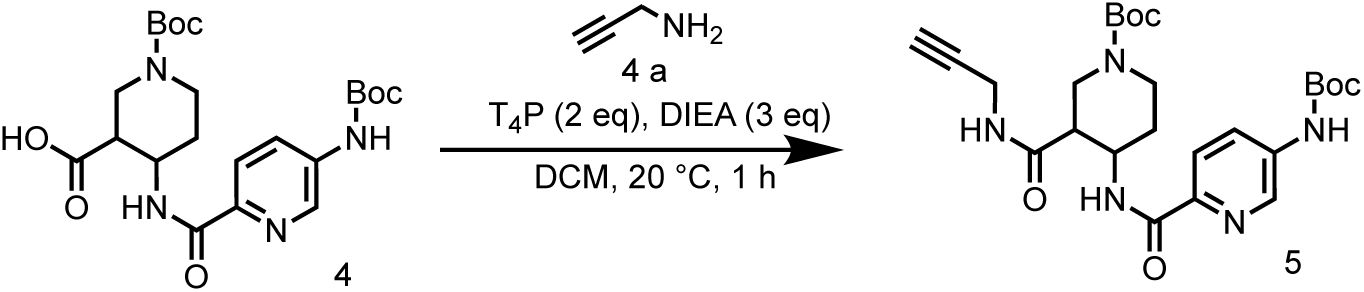

To a solution of 1-(*tert*-butoxycarbonyl)-4-(5-((tert-butoxycarbonyl)amino)picolinamido)piperidine-3-carboxylic acid (1.10 g, 2.37 mmol) in DCM (10 mL) was added T_4_P (3.41 g, 4.74 mmol, 50% purity), DIEA (918 mg, 7.10 mmol, 1.24 mL) and prop-2-yn-1-amine (156 mg, 2.84 mmol, 182 μL). The mixture was stirred at 20 °C for 12 hrs. LCMS showed the reaction was completed. The reaction mixture was concentrated in vacuum to give a residue. The residue was purified by column chromatography on silica gel (eluted with petroleum ether: ethyl acetate=10:0 to 1:1) to give *tert*-butyl 4-(5-((*tert*-butoxycarbonyl)amino)picolinamido)-3-(prop-2-yn-1-ylcarbamoyl)piperidine-1-carboxylate (610 mg, 1.22 mmol, 51.3% yield) as a white solid.

#### Step 4: Preparation of 5-amino-*N*-(3-(prop-2-yn-1-ylcarbamoyl)piperidin-4-yl)picolinamide (6)

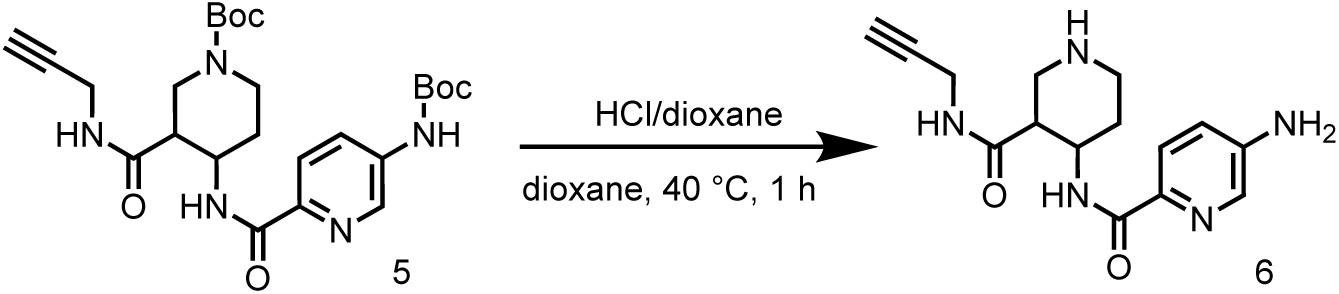

To a solution of *tert*-butyl 4-(5-((tert-butoxycarbonyl)amino)picolinamido)-3-(prop-2-yn-1-ylcarbamoyl)piperidine-1-carboxylate (600 mg, 1.20 mmol) in dioxane (5 mL) was added HCl/dioxane (4 M, 5 mL). The mixture was stirred at 40 °C for 1 hr. LCMS showed the reaction was completed. The mixture was concentrated in vacuum to give 5-amino-*N*-(3-(prop-2-yn-1-ylcarbamoyl)piperidin-4-yl)picolinamide (300 mg, 888 μmol, 74.2% yield, HCl salt) as a white solid.

LC-MS: 302.1 [M+H]^+^ / Ret time: 0.334 min / method: 5-95CD_1min.lcm.

#### Step 5: Preparation of 5-(2-chloroacetamido)-*N*-((3*R*,4*S*)-1-(2-chloroacetyl)-3-(prop-2-yn-1-ylcarbamoyl)piperidin-4-yl)picolinamide (SH-X-61A) and 5-(2-chloroacetamido)-*N*-((3*R*,4*R*)-1-(2-chloroacetyl)-3-(prop-2-yn-1-ylcarbamoyl)piperidin-4-yl)picolinamide (SH-X-61B)

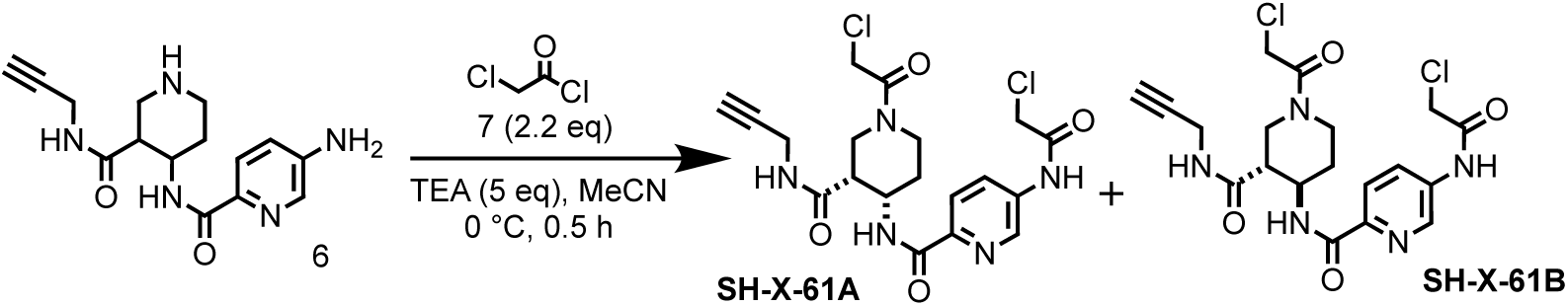

To a solution of 5-amino-*N*-(3-(prop-2-yn-1-ylcarbamoyl)piperidin-4-yl)picolinamide (300 mg, 888 μmol, HCl salt) in MeCN (1 mL) was added TEA (449.0 mg, 4.44 mmol, 618.0 μL) and 2-chloroacetyl chloride (220 mg, 1.95 mmol, 155 μL). The mixture was stirred at 0 °C for 0.5 hr. LCMS showed the reaction was completed. The reaction mixture was concentrated in vacuum to give a residue. Then the residue was poured into H_2_O (50 mL), extracted with EtOAc (100 mL×3). The organic layers were collected and washed with brine (100 mL×3), dried over Na_2_SO_4_ and concentrated to give 5-(2-chloroacetamido)-*N*-(1-(2-chloroacetyl)-3-(prop-2-yn-1-ylcarbamoyl)piperidin-4-yl)picolinamide (100 mg, 220 μmol, 24.8% yield) as a white solid.

LC-MS: 454.0 [M+H]^+^ / Ret time: 0.303 min / method: 5-95AB_0.8min.lcm.

SH-X-61B: ^1^H NMR (400 MHz, DMSO-*d*_6_) *δ* 10.77 (s, 1H), 8.83 (d, *J* = 2.0 Hz, 1H), 8.50 (m, 1H), 8.35 - 8.23 (m, 1H), 8.17 (m, 1H), 8.01 (d, *J* = 8.4 Hz, 1H), 4.51 - 4.33 (m, 5H), 4.30 - 4.21 (m, 1H), 3.92 - 3.83 (m, 1H), 3.79 (d, *J* = 4.4 Hz, 2H), 3.25 - 3.15 (m, 1H), 2.91 (d, *J* = 17.6 Hz, 1H), 2.76 (t, *J* = 12.0 Hz, 1H), 2.64 - 2.53 (m, 1H), 1.97 - 1.83 (m, 1H), 1.64 - 1.33 (m, 1H)

SH-X-61B: ^1^H NMR (400 MHz, DMSO-*d*_6_) *δ* 10.81 (s, 1H), 8.74 (s, 1H), 8.66 - 8.39 (m, 2H), 8.26 (m, 1H), 8.03 (d, *J* = 8.4 Hz, 1H), 4.50 - 4.40 (m, 1H), 4.39 - 4.17 (m, 4H), 4.05 - 3.71 (m, 4H), 3.62 - 3.53 (m, 1H), 3.23 (t, *J* = 9.6 Hz, 1H), 3.08 - 3.03 (m, 1H), 2.87 - 2.69 (m, 1H), 2.29 - 2.07 (m, 1H), 1.74 - 1.54 (m, 1H).

### Synthesis of *N,N*’-(2-(7-(prop-2-yn-1-yloxy)-2,3-dihydrobenzo[b][1,4] dioxine-6-carboxamido)-[1,1’-biphenyl]-4,4’-diyl)bis(2-chloroacetamide) (SH-X-54)

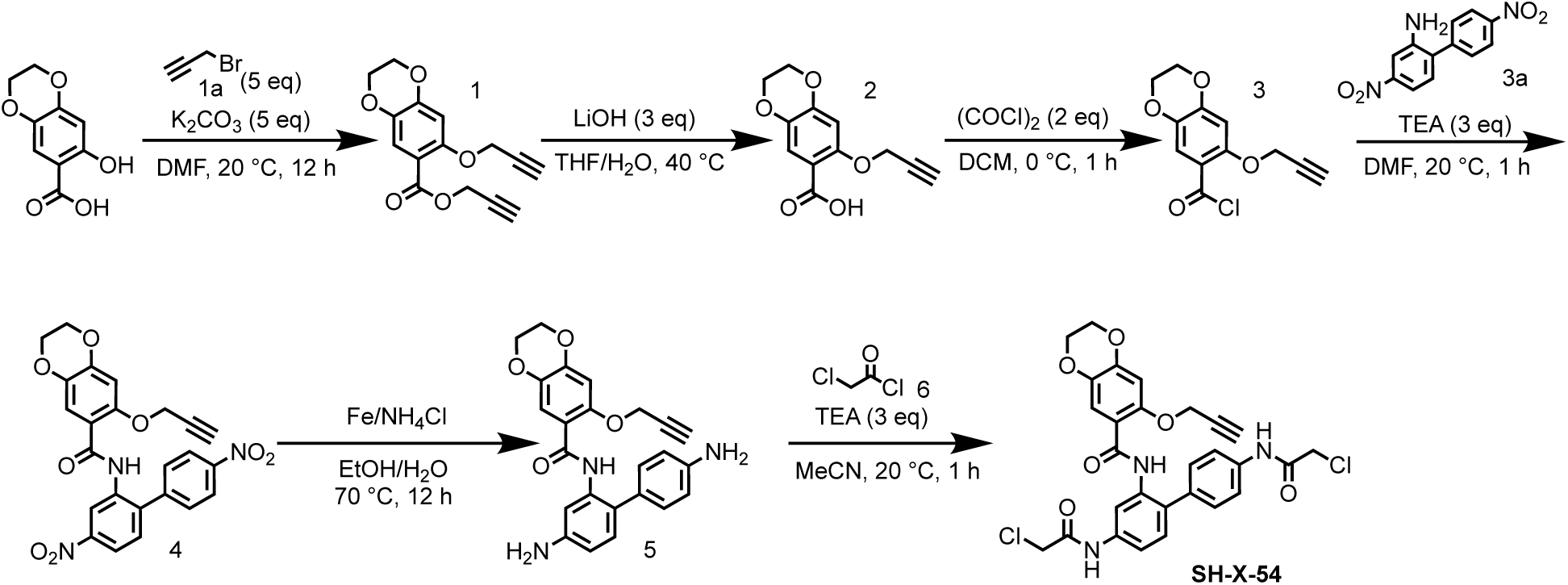

#### Step 1: Preparation of prop-2-yn-1-yl 7-(prop-2-yn-1-yloxy)-2,3-dihydrobenzo[b] [1,4]dioxine-6-carboxylate (1)

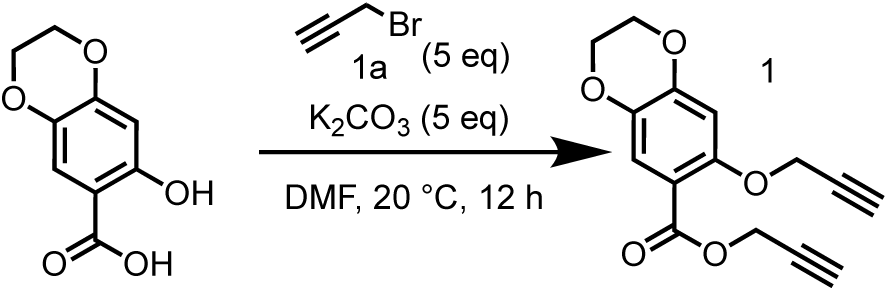

To a solution of 7-hydroxy-2,3-dihydrobenzo[b][1,4]dioxine-6-carboxylic acid (3.00 g, 15.3 mmol) in DMF (30 mL) was added K_2_CO_3_ (10.6 g, 76.5 mmol) and 3-bromoprop-1-yne (11.4 g, 76.5 mmol, 8.24 mL). The mixture was stirred at 20 °C for 12 hrs. LCMS showed the reaction was completed. The reaction mixture was poured into H_2_O (150 mL), extracted with EtOAc (200 mL×3). The organic layers were collected and washed with brine (200 mL×3), dried over Na_2_SO_4_ and concentrated to give prop-2-yn-1-yl 7-(prop-2-yn-1-yloxy)-2,3-dihydrobenzo[b][1,4]dioxine-6-carboxylate (4 g, crude) as a white solid.

LC-MS: 273.2 [M+H]^+^ / Ret time: 0.383 min / method: 5-95AB_0.8min.lcm

#### Step 2: Preparation of 7-(prop-2-yn-1-yloxy)-2,3-dihydrobenzo[b][1,4]dioxine-6-carboxylic acid (2)

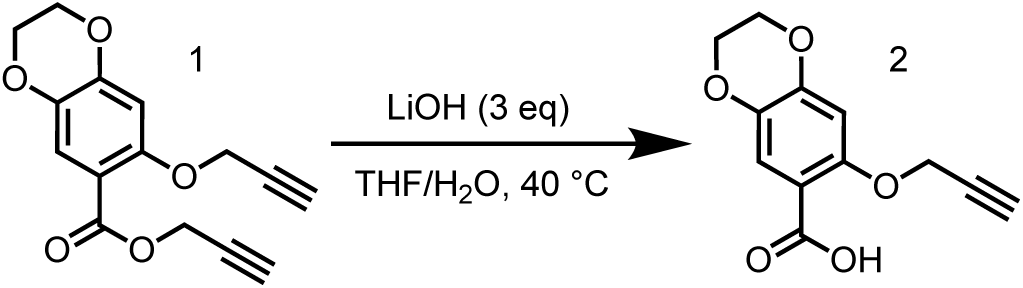

To a solution of prop-2-yn-1-yl 7-(prop-2-yn-1-yloxy)-2,3-dihydrobenzo[b][1,4] dioxine-6-carboxylate (4.00 g, 14.7 mmol) in THF (20 mL) and H2O (20 mL) was added LiOH•H2O (1.85 g, 44.1 mmol). The mixture was stirred at 20 °C for 1 hr. LCMS showed the reaction was completed. The reaction mixture was evaporated in vacuum. The mixture was acidified with 1 mol HCl to pH 4. The suspension was filtered and the filter cake was washed with MeOH (30 mL). The solid was concentrated under vacuum to give 7-(prop-2-yn-1-yloxy)-2,3-dihydrobenzo[b][1,4]dioxine-6-carboxylic acid (3.30 g, 14.1 mmol, 95.9% yield) as a white solid.

LC-MS: 235.0 [M+H]^+^ / Ret time: 0.257 min / method: 5-95AB_0.8min.lcm

#### Step 3: Preparation of 7-(prop-2-yn-1-yloxy)-2,3-dihydrobenzo[b][1,4]dioxine-6-carbonyl chloride (3)

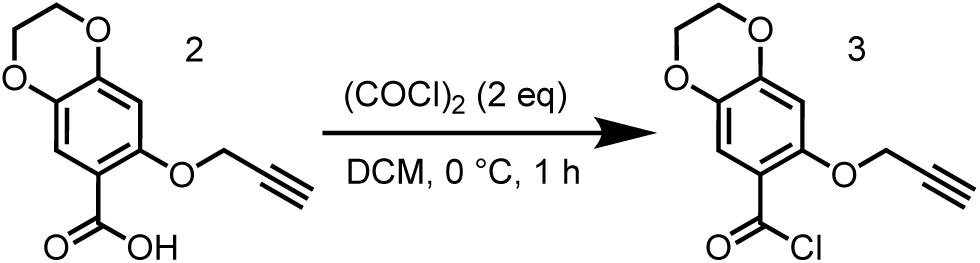

To a solution of 7-(prop-2-yn-1-yloxy)-2,3-dihydrobenzo[b][1,4]dioxine-6-carboxylic acid (1.50 g, 6.40 mmol) in DCM (2 mL) was added oxalyl dichloride (1.63 g, 12.8 mmol, 1.12 mL) and DMF (46.8 mg, 640 μmol, 49.3 μL) at 0 °C. The mixture was stirred at 20 °C for 1 hr. TLC indicated Reactant 1 was consumed completely and one new spot formed. The reaction was clean according to TLC. The reaction mixture was evaporated in vacuum to give 7-(prop-2-yn-1-yloxy)-2,3-dihydrobenzo[b][1,4]dioxine-6-carbonyl chloride (1.62 g, crude) as a light yellow solid.

#### Step 4: Preparation of *N*-(4,4’-dinitro-[1,1’-biphenyl]-2-yl)-7-(prop-2-yn-1-yloxy)-2,3-dihydrobenzo[b][1,4]dioxine-6-carboxamide (4)

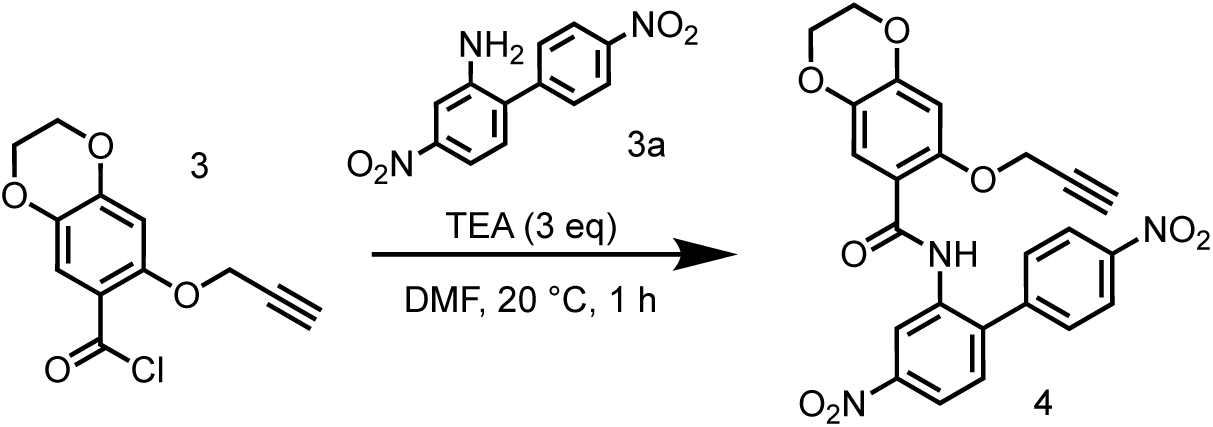

To a solution of 7-(prop-2-yn-1-yloxy)-2,3-dihydrobenzo[b][1,4] dioxine-6-carbonyl chloride (1.62 g, 6.41 mmol) in DCM (20 mL) was added TEA (1.95 g, 19.2 mmol, 2.68 mL) and 5-nitro-2-(4-nitrophenyl)aniline (1.66 g, 6.41 mmol). The mixture was stirred at 0 °C for 1 hr. LCMS showed the reaction was completed. The residue was triturated in MeCN (20 ml) and stirred for 30 min. The mixture was filtered and the filter cake was concentrated to give *N*-(4,4’-dinitro-[1,1’-biphenyl]-2-yl)-7-(prop-2-yn-1-yloxy)-2,3-dihydrobenzo[b][1,4]dioxine-6-carboxamide (600 mg, 1.26 mmol, 19.7% yield) as a white solid.

LC-MS: 476.0 [M+H]^+^ / Ret time: 0.524 min / method: 5-95AB_0.8min.lcm

#### Step 5: Preparation of *N*-(4,4’-diamino-[1,1’-biphenyl]-2-yl)-7-(prop-2-yn-1-yloxy)-2,3-dihydrobenzo[b][1,4]dioxine-6-carboxamide (5)

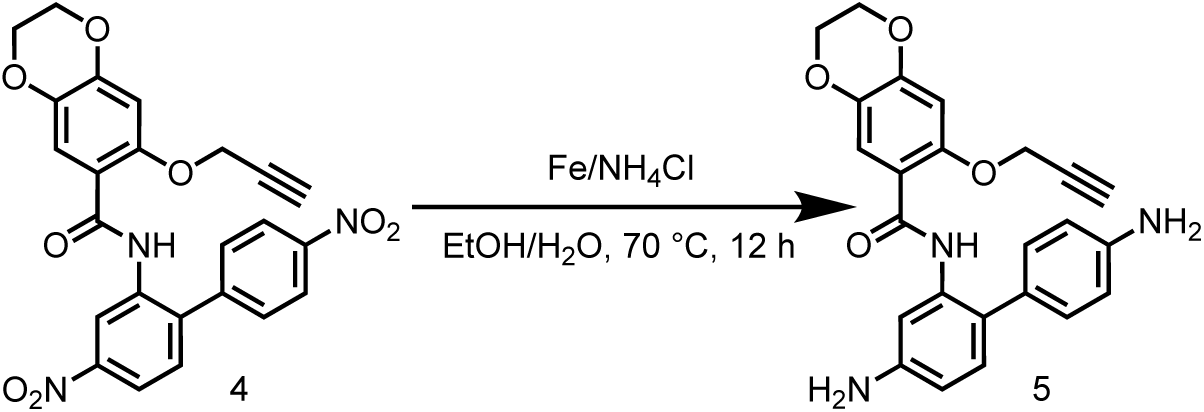

To a solution of *N*-(4,4’-dinitro-[1,1’-biphenyl]-2-yl)-7-(prop-2-yn-1-yloxy)-2,3-dihydrobenzo[b][1,4]dioxine-6-carboxamide (600 mg, 1.26 mmol) in EtOH (3 mL) and H_2_O (3 mL) was added Fe (704 mg, 12.6 mmol) and NH_4_Cl (675 mg, 12.6 mmol). The mixture was stirred at 70 °C for 12 hrs. LCMS showed the reaction was completed. The reaction mixture was filtered and the filtrate was concentrated to give *N*-(4,4’-dinitro-[1,1’-biphenyl]-2-yl)-7-(prop-2-yn-1-yloxy)-2,3-dihydrobenzo[b][1,4] dioxine-6-carboxamide (360 mg, crude) as an off-white solid.

LC-MS: 416.1 [M+H]^+^ / Ret time: 0.289 min / method: 5-95AB_0.8min.lcm

#### Step 6: Preparation of *N,N*’-(2-(7-(prop-2-yn-1-yloxy)-2,3-dihydrobenzo[b][1,4] dioxine-6-carboxamido)-[1,1’-biphenyl]-4,4’-diyl)bis(2-chloroacetamide) (SH-X-54)

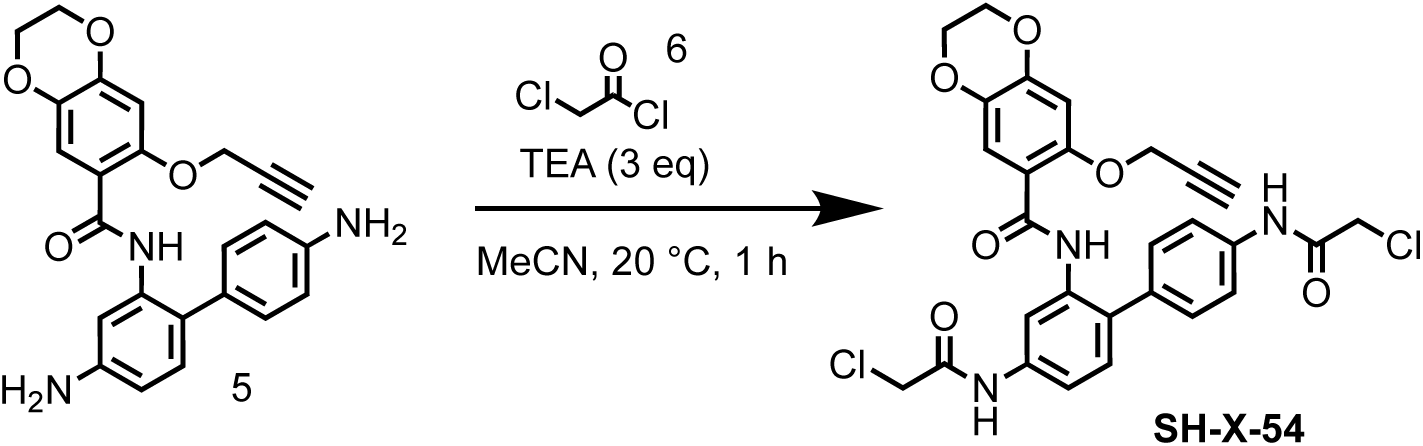

To a solution of *N*-(4,4’-dinitro-[1,1’-biphenyl]-2-yl)-7-(prop-2-yn-1-yloxy)-2,3-dihydrobenzo[b][1,4] dioxine-6-carboxamide (300 mg, 722 μmol) in MeCN (10 mL) was added TEA (219 mg, 2.17 mmol, 301 μL) and 2-chloroacetyl chloride (179 mg, 1.59 mmol, 126 μL). The mixture was stirred at 0 °C for 1 hr. LCMS showed the reaction was completed. The reaction mixture was evaporated in vacuum. The residue was purified by preparative-HPLC (column: Welch Ultimate C18 150*25mm*5um; mobile phase: [water(FA)-MeCN];gradient:34%-64% B over 10 min) to give *N*,*N*’-(2-(7-(prop-2-yn-1-yloxy)-2,3-dihydrobenzo[b][1,4]dioxine-6-carboxamido)-[1,1’-biphenyl]-4,4’-diyl)bis(2-chloroacetamide) (89.6 mg, 149 μmol, 20.8% yield, 95.1% purity) as a white solid.

LC-MS: 568.0 [M+H]^+^ / Ret time: 0.459 min / method: 5-95AB_0.8min.lcm

^1^H NMR (400 MHz, DMSO-*d*_6_) *δ* 10.47 (s, 1H), 10.42 (s, 1H), 9.65 (s, 1H), 8.51 (s, 1H), 7.71 (d, *J* = 8.4 Hz, 2H), 7.62 (m, *J* = 1.6, 8.4 Hz, 1H), 7.45 (s, 1H), 7.38 (d, *J* = 8.4 Hz, 2H), 7.20 (d, *J* = 8.4 Hz, 1H), 6.72 (s, 1H), 4.36 (d, *J* = 2.0 Hz, 2H), 4.28 (d, *J* = 5.6 Hz, 6H), 4.22 (d, *J* = 3.2 Hz, 2H), 3.52 - 3.47 (m, 1H).

### Synthesis of *N,N*’-(5-(prop-2-yn-1-ylcarbamoyl)-1,3-phenylene)bis(1-(2-chloroacetyl)-4-methylpiperidine-4-carboxamide) (SH-X-67)

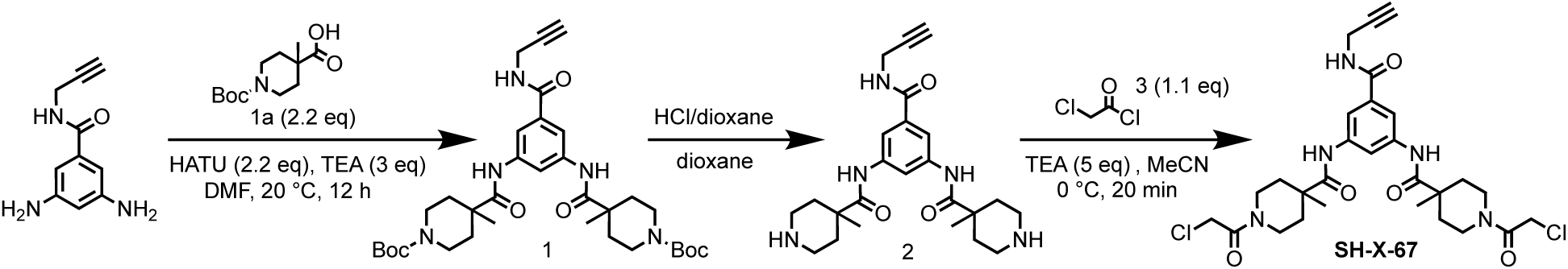

#### Step 1: Preparation of di-*tert*-butyl 4,4’-(((5-(prop-2-yn-1-ylcarbamoyl)-1,3-phenylene)bis(azanediyl))bis(carbonyl))bis(4-methylpiperidine-1-carboxylate) (1)

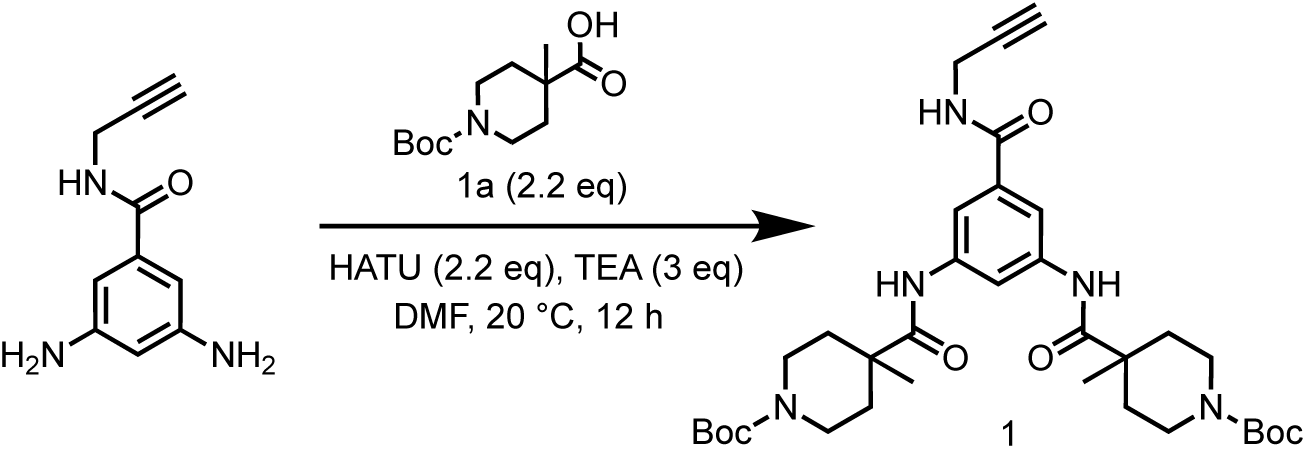

To a solution of 3,5-diamino-*N*-(prop-2-yn-1-yl)benzamide in DMF (6 mL) was added HATU (1.33 g, 3.49 mmol) and TEA (481 mg, 4.76 mmol, 662 μL) and 1-(tert-butoxycarbonyl)-4-methylpiperidine-4-carboxylic acid (849 mg, 3.49 mmol). The mixture was stirred at 20 °C for 12 hrs. LCMS showed the reaction was completed. The reaction mixture was poured into H_2_O (20 mL), extracted with EtOAc (20 mL×3). The organic layers were collected and washed with brine (20 mL×3), dried over Na_2_SO_4_ and concentrated to give a residue. The residue was purified by column chromatography on silica gel (eluted with petroleum ether: ethyl acetate=1:2) to give di-*tert*-butyl 4,4’-(((5-(prop-2-yn-1-ylcarbamoyl)-1,3-phenylene)bis(azanediyl))bis(carbonyl))bis(4-methylpiperidine-1-carboxylate) (1.00 g, 1.56 mmol, 98.6% yield) as an off-white solid.

LC-MS: 484.2 [M-156+H]^+^ / Ret time: 0.524 min / method: 5-95AB_0.8min.lcm

#### Step 2: Preparation of *N,N*’-(5-(prop-2-yn-1-ylcarbamoyl)-1,3-phenylene)bis(4-methylpiperidine-4-carboxamide) (2)

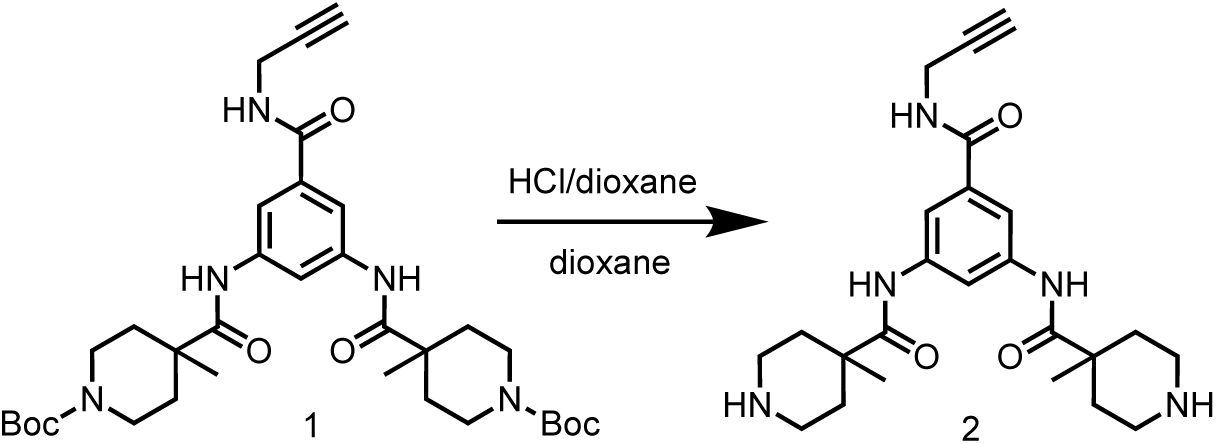

To a solution of di-*tert*-butyl 4,4’-(((5-(prop-2-yn-1-ylcarbamoyl)-1,3-phenylene)bis (azanediyl))bis(carbonyl))bis(4-methylpiperidine-1-carboxylate) (60.0 mg, 93.8 μmol) in dioxane (0.5 mL) was added HCl/dioxane (4 M, 0.5 mL). The mixture was stirred at 20 °C for 1 hr. LCMS showed the reaction was completed. The reaction mixture was concentrated on vacuum to remove solvent, the solid was triturated with DCM. The reaction mixture was filtered and the filter cake was concentrated to give *N*,*N*’-(5-(prop-2-yn-1-ylcarbamoyl)-1,3-phenylene)bis(4-methylpiperidine-4-carboxamide) (31.8 mg, 64.4 μmol, HCl salt) as a yellow solid.

LC-MS: 440.0 [M+H]^+^ / Ret time: 0.294 min / method: 0-60AB_0.8min.lcm

^1^H NMR (400 MHz, DMSO-*d*_6_): *δ* 9.73 (s, 2H), 8.88 (t, *J* = 5.6 Hz, 1H), 8.81-8.74 (m, 2H), 8.23 (s, 1H), 7.77 (d, *J* = 1.2 Hz, 2H), 4.04-4.00 (m, 2H), 3.23-3.17 (m, 4H), 3.12 (t, *J* = 2.0 Hz, 1H), 2.97-2.85 (m, 4H), 2.30 (d, *J* = 15.2 Hz, 4H), 1.75-1.64 (m, 4H), 1.31 (s, 6H).

#### Step 3: Preparation of *N,N*’-(5-(prop-2-yn-1-ylcarbamoyl)-1,3-phenylene)bis(1-(2-chloroacetyl)-4-methylpiperidine-4-carboxamide) (SH-X-067)

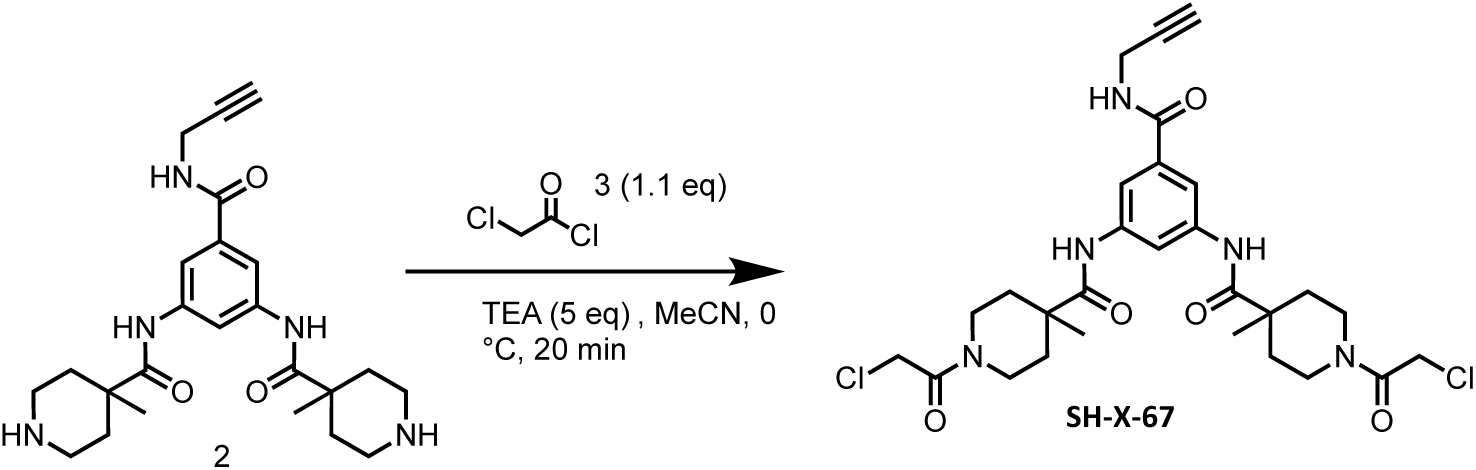

To a solution of *N*,*N*’-(5-(prop-2-yn-1-ylcarbamoyl)-1,3-phenylene)bis(4-methylpiperidine-4-carboxamide) (300 mg, 630 μmol, HCl salt) in MeCN (12 mL) was added TEA (319 mg, 3.15 mmol, 439 μL), then 2-chloroacetyl chloride (78.3 mg, 693 μmol, 55.2 μL) in MeCN (6 mL) was added into the reaction mixture at 0 °C, the reaction mixture was stirred at 0 °C for 20 min. LCMS showed the reaction was completed. The reaction mixture was filtered and the filtrate was concentrated to give a residue. The residue was purified by preparative-HPLC (column: Phenomenex luna C18 150*25mm* 10um;mobile phase: [water(FA)-MeCN];gradient:20%-50% B over 10 min) to give *N*,*N*’-(5-(prop-2-yn-1-ylcarbamoyl)-1,3-phenylene)bis(1-(2-chloroacetyl)-4-methylpiperidine-4-carboxamide) (58.3 mg, 97.0 μmol, 15.4% yield, 98.6% purity) as a white solid.

LC-MS: 592.0 [M+H]^+^ / Ret time: 0.393 min / method: 5-95AB_0.8min.lcm

^1^H NMR (400 MHz, DMSO-*d*_6_): *δ* 9.58 (s, 2H), 8.84 (s, 1H), 8.19-8.17 (m, 1H), 7.76 (d, *J* = 1.6 Hz, 2H), 4.37 (d, *J* = 1.6 Hz, 4H), 4.02 (dd, *J* = 2.4, 5.6 Hz, 2H), 3.91-3.82 (m, 2H), 3.67-3.57 (m, 2H), 3.27 (t, *J* = 10.8 Hz, 2H), 3.13-3.03 (m, 3H), 2.16 (t, *J* = 16.4 Hz, 4H), 1.55-1.46 (m, 2H), 1.44-1.35 (m, 2H), 1.27 (s, 6H).

### Synthesis of *N,N*’-(5-(prop-2-yn-1-ylcarbamoyl)-1,3-phenylene)bis(2-(2-chloroacetyl)-2-azaspiro[3.3]heptane-6-carboxamide) (SH-X-79)

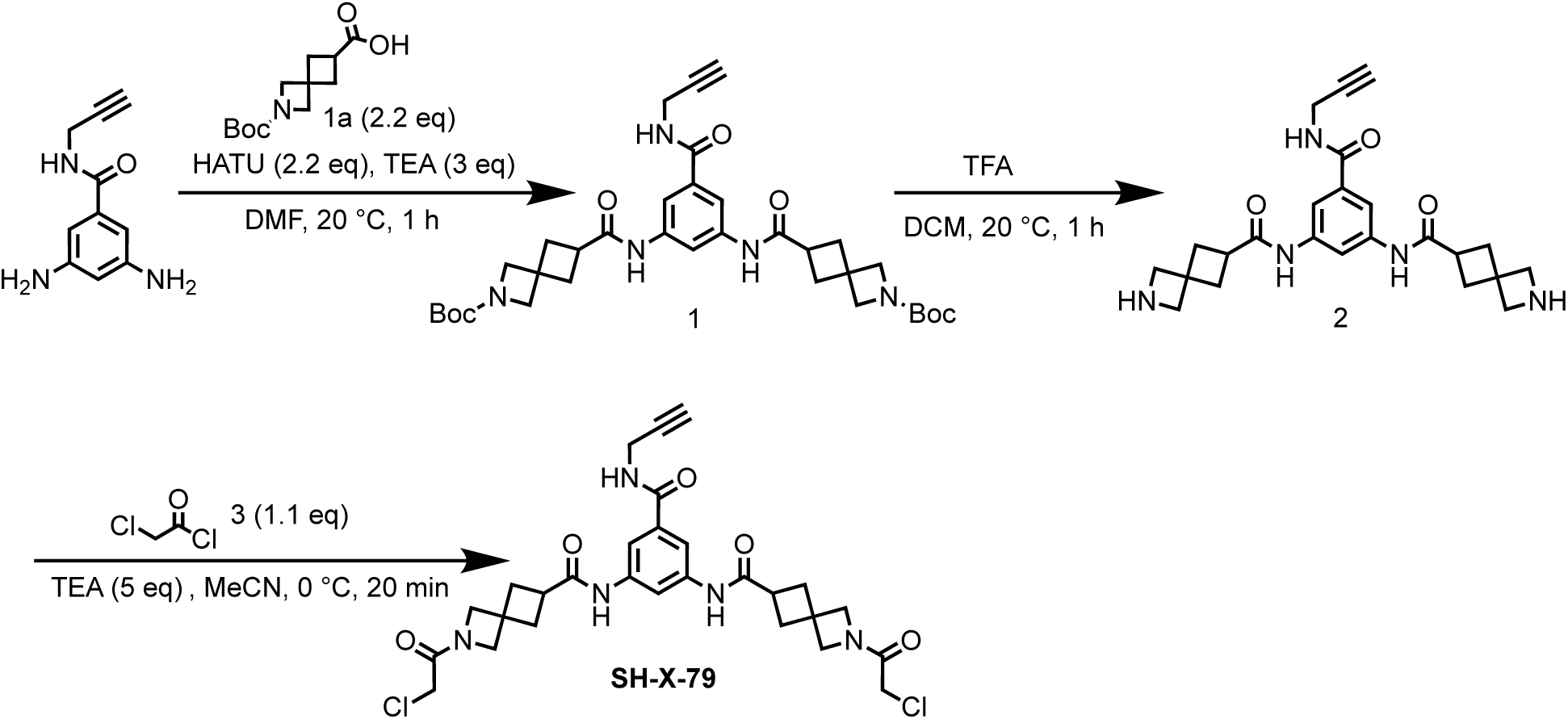

#### Step 1: Preparation of di-*tert*-butyl 6,6’-(((5-(prop-2-yn-1-ylcarbamoyl)-1,3-phenylene)bis(azanediyl))bis(carbonyl))bis(2-azaspiro[3.3]heptane-2-carboxylate) (1)

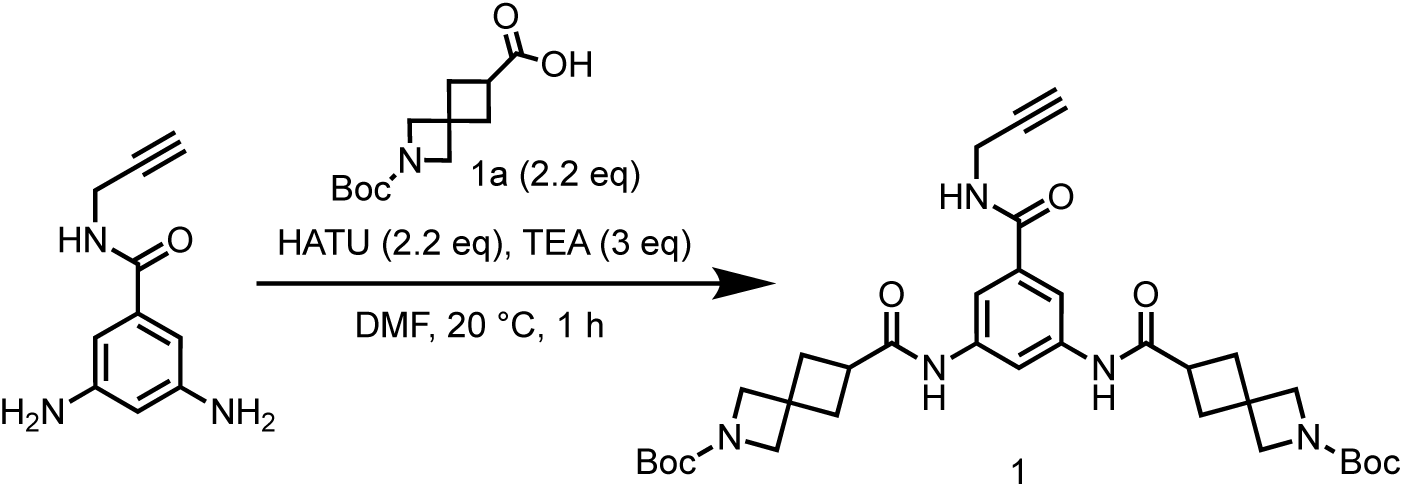

To a solution of 2-(*tert*-butoxycarbonyl)-2-azaspiro[3.3]heptane-6-carboxylic acid (841 mg, 3.49 mmol) in DMF (3 mL) was added HATU (1.33 g, 3.49 mmol) and TEA (481 mg, 4.76 mmol, 662 μL), the mixture was stirred at 20°C for 10 min, the mixture was added 3,5-diamino-*N*-(prop-2-yn-1-yl)benzamide (300 mg, 1.59 mmol). The mixture was stirred at 20 °C for 1 hr. LCMS showed the reaction was completed. The reaction mixture was poured into H_2_O (30 mL), extracted with EtOAc (30 mL×3). The organic layers were collected and washed with brine (30 mL×3), dried over Na_2_SO_4_ and concentrated to give a residue. The residue was purified by column chromatography on silica gel (eluted with petroleum ether: ethyl acetate=0:1) to give di-*tert*-butyl 6,6’-(((5-(prop-2-yn-1-ylcarbamoyl)-1,3-phenylene)bis(azanediyl))bis(carbonyl))bis(2-azaspiro[3.3]heptane-2-carboxylate) (1.00 g, 1.57 mmol, 99.2% yield) as a yellow oil.

LC-MS: 480.1 [M-156+H]^+^ / Ret time: 0.499 min / method: 5-95AB_0.8min.lcm

#### Step 2: Preparation of *N,N*’-(5-(prop-2-yn-1-ylcarbamoyl)-1,3-phenylene)bis(2-azaspiro[3.3]heptane-6-carboxamide) (2)

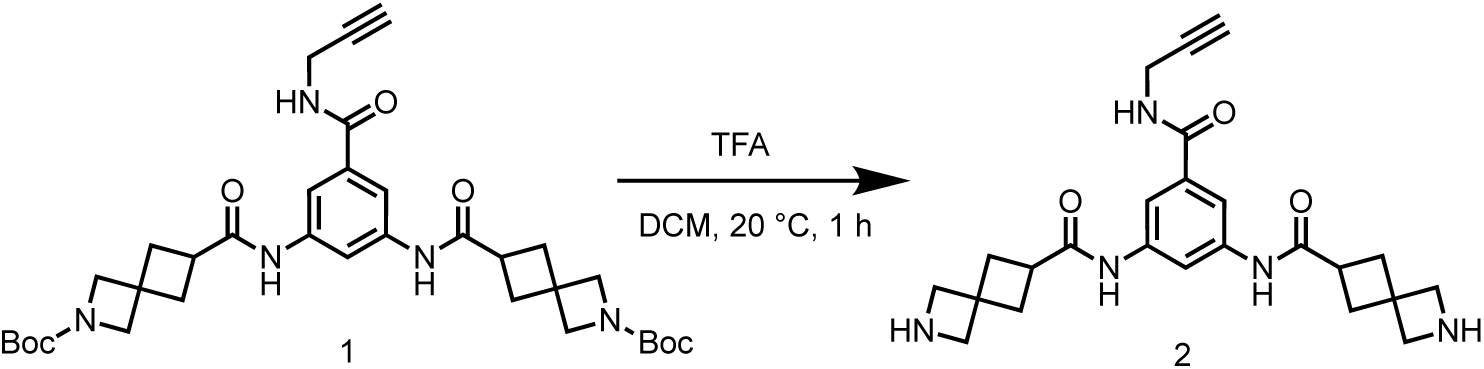

To a solution of di-*tert*-butyl 6,6’-(((5-(prop-2-yn-1-ylcarbamoyl)-1,3-phenylene)bis (azanediyl))bis(carbonyl))bis(2-azaspiro[3.3]heptane-2-carboxylate) (1.00 g, 1.57 mmol) in DCM (5 mL) was added TFA (7.68 g, 67.3 mmol, 5.00 mL). The mixture was stirred at 20 °C for 1 hr. LCMS showed the reaction was completed. The reaction mixture was concentrated in vacuum to give *N*,*N*’-(5-(prop-2-yn-1-ylcarbamoyl)-1,3-phenylene)bis(2-azaspiro [3.3]heptane-6-carboxamide) (2.00 g, crude, TFA salt) as a white solid.

LC-MS: 436.1 [M+H]^+^ / Ret time: 0.210 min / method: 5-95AB_0.8min.lcm

^1^H NMR (400 MHz, DMSO-*d*_6_): *δ* 10.02 (s, 2H), 8.85 (t, *J* = 5.6 Hz, 1H), 8.42 (s, 2H), 8.10 (s, 1H), 7.69 (s, 2H), 3.99 (dd, *J* = 2.4, 5.2 Hz, 2H), 3.85 (s, 4H), 3.79 (s, 4H), 3.10 (t, *J* = 2.4 Hz, 1H), 3.08-3.01 (m, 3H), 2.43-2.29 (m, 9H).

#### Step 3: Preparation of *N,N*’-(5-(prop-2-yn-1-ylcarbamoyl)-1,3-phenylene)bis(2-(2-chloroacetyl)-2-azaspiro[3.3]heptane-6-carboxamide) (SH-X-79)

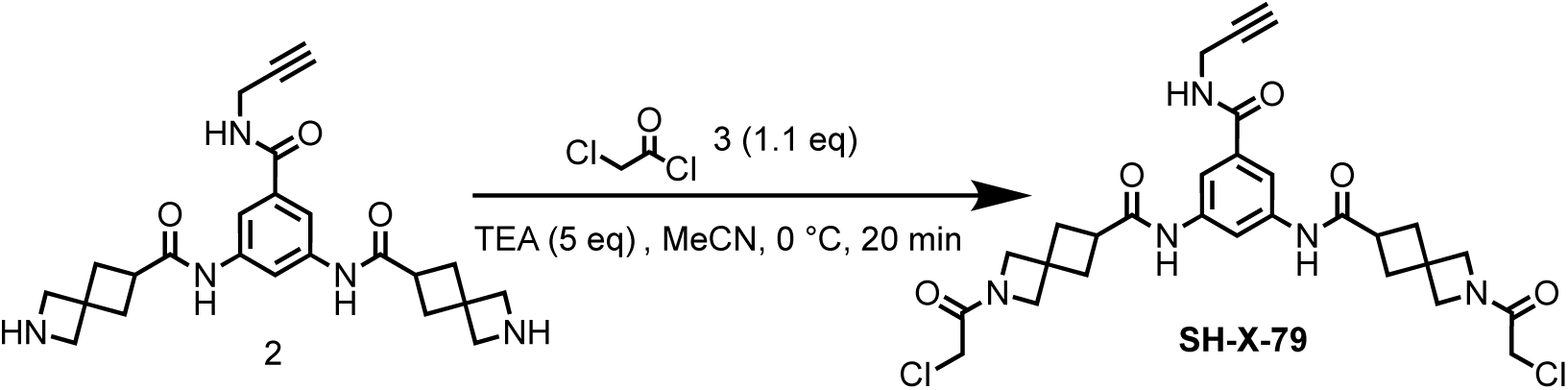

To a solution of *N*,*N*’-(5-(prop-2-yn-1-ylcarbamoyl)-1,3-phenylene)bis(2-azaspiro[3.3]heptane-6-carboxamide) (500 mg, 910 μmol, TFA salt) in MeCN (20 mL) was added TEA (460 mg, 4.55 mmol, 633 μL), then 2-chloroacetyl chloride (113 mg, 1.00 mmol, 79.7 μL) in MeCN (10 mL) was added into the reaction mixture at 0 °C, the reaction mixture was stirred at 0 °C for 20 min. LCMS showed the reaction was completed. The reaction mixture was filtered and the filtrate was concentrated to give a residue. The residue was purified by preparative-HPLC (column: Phenomenex luna C18 150*25mm* 10um;mobile phase: [water(FA)-MeCN]; gradient:14%-34% B over 10 min) to give *N*,*N*’-(5-(prop-2-yn-1-ylcarbamoyl)-1,3-phenylene)bis(2-(2-chloroacetyl)-2-azaspiro[3.3]heptane-6-carboxamide) (11.5 mg, 18.4 μmol, 2.02% yield, 94.1% purity) as a white solid.

LC-MS: 588.1 [M+H]^+^ / Ret time: 0.356 min / method: 5-95AB_0.8min.lcm

^1^H NMR (400 MHz, DMSO-*d*_6_): *δ* 9.98 (d, *J* = 7.2 Hz, 2H), 8.81 (t, *J* = 5.6 Hz, 1H), 8.10 (s, 1H), 7.68 (s, 2H), 4.24 (s, 2H), 4.17 (s, 2H), 4.10 (d, *J* = 3.6 Hz, 4H), 4.00 (dd, *J* = 2.4, 5.6 Hz, 2H), 3.94 (s, 2H), 3.86 (s, 2H), 3.12-3.07 (m, 2H), 2.44-2.31 (m, 9H).

### Synthesis of *N,N*’-(5-(prop-2-yn-1-ylcarbamoyl)-1,3-phenylene)bis(3-(2-chloroacetyl)-3-azabicyclo[3.1.0]hexane-1-carboxamide) (SH-X-85)

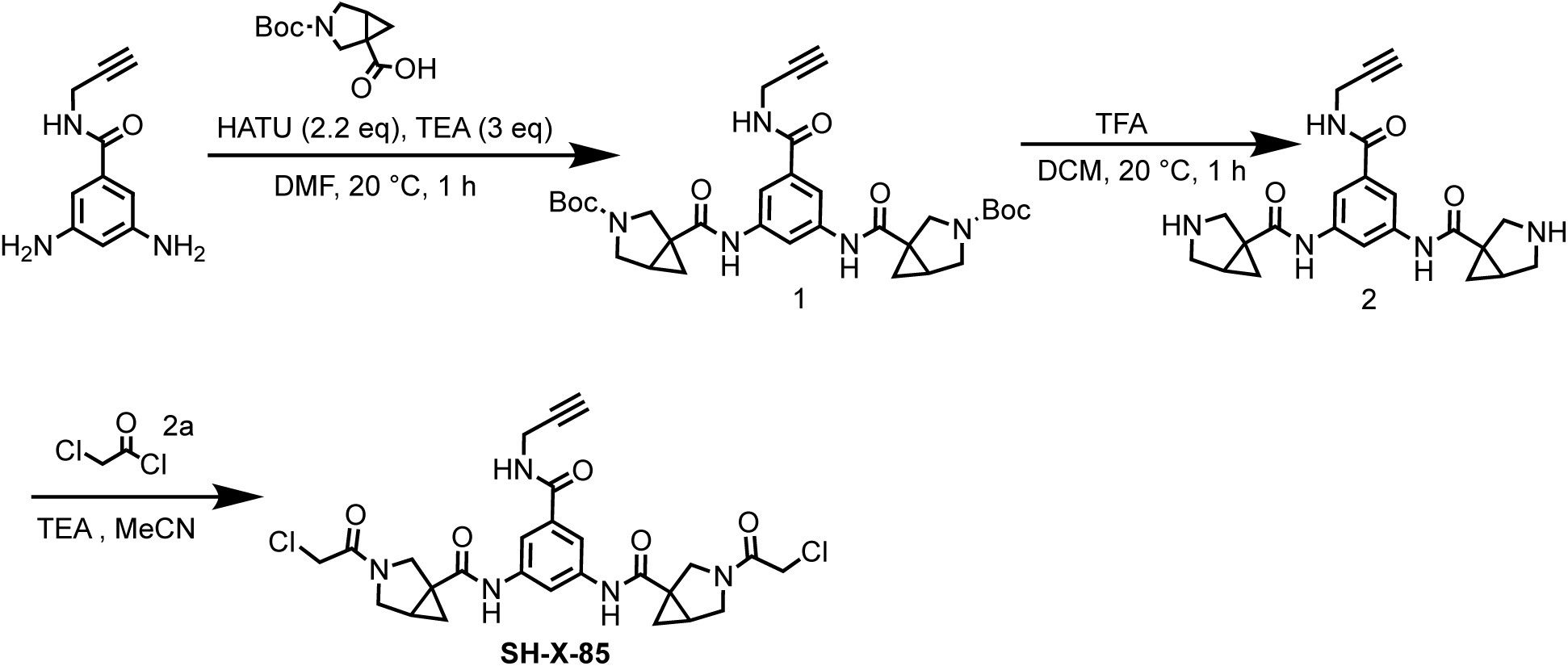

#### Step 1: Preparation di-*tert*-butyl 1,1’-(((5-(prop-2-yn-1-ylcarbamoyl)-1,3-phenylene)bis(azanediyl))bis(carbonyl))bis(3-azabicyclo[3.1.0]hexane-3-carboxylate) (1)

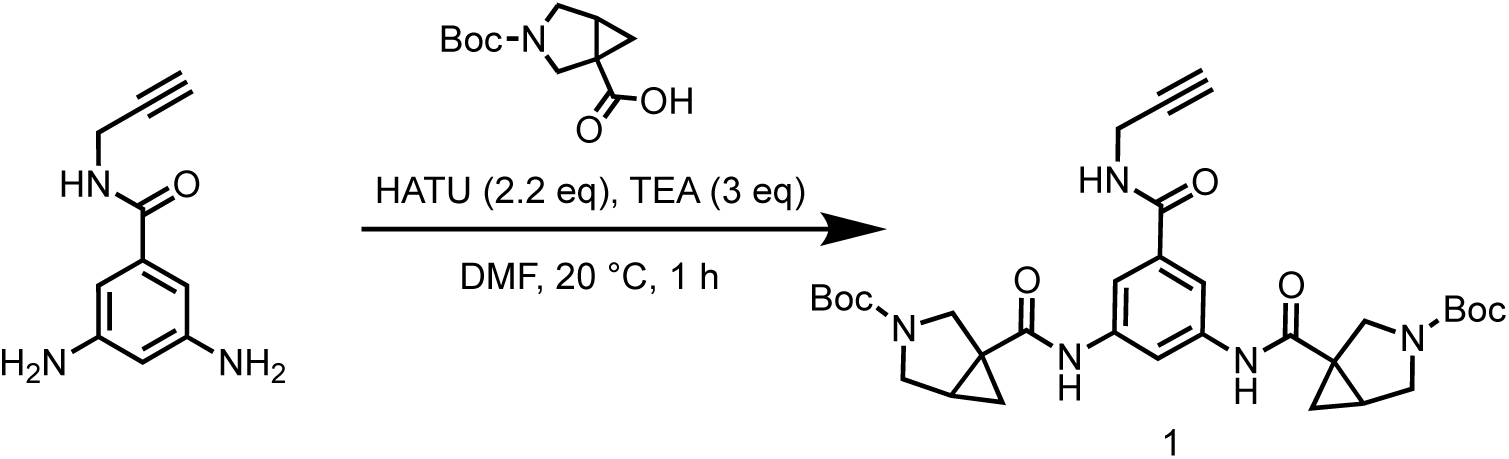

To a solution of 3-(*tert*-butoxycarbonyl)-3-azabicyclo[3.1.0]hexane-1-carboxylic acid (740 mg, 3.26 mmol) in DMF (10 mL) was added HATU (1.35 g, 3.55 mmol) and TEA (449 mg, 4.44 mmol, 618 μL), the mixture was stirred at 20 °C for 0.5 hr. Then 3,5-diamino-N-(prop-2-yn-1-yl)benzamide (280 mg, 1.48 mmol) was added into the reaction mixture and stirred at 20 °C for 1 hr. LCMS showed the reaction was completed. The reaction mixture was extracted with H_2_O (50 mL) and ethyl acetate (50 mL×3), washed with brine (50×3 mL), the organic phase was dried over Na_2_SO_4_, concentrated in vacuum to give a residue. The residue was purified by column chromatography on silica gel (eluted with petroleum ether: ethyl acetate=1:1 to 1:1) to give di-*tert*-butyl 1,1’-(((5-(prop-2-yn-1-ylcarbamoyl)-1,3-phenylene)bis(azanediyl))bis(carbonyl))bis(3-azabicyclo[3.1.0]hexane-3-carboxylate) (1.2 g, crude) as a yellow solid.

LC-MS: 452.2 [M-156+H]^+^ / Ret time: 0.455 min / method: 5-95AB_0.8min.lcm.

#### Step 2: Preparation of *N,N*’-(5-(prop-2-yn-1-ylcarbamoyl)-1,3-phenylene)bis(3-azabicyclo[3.1.0]hexane-1-carboxamide) (2)

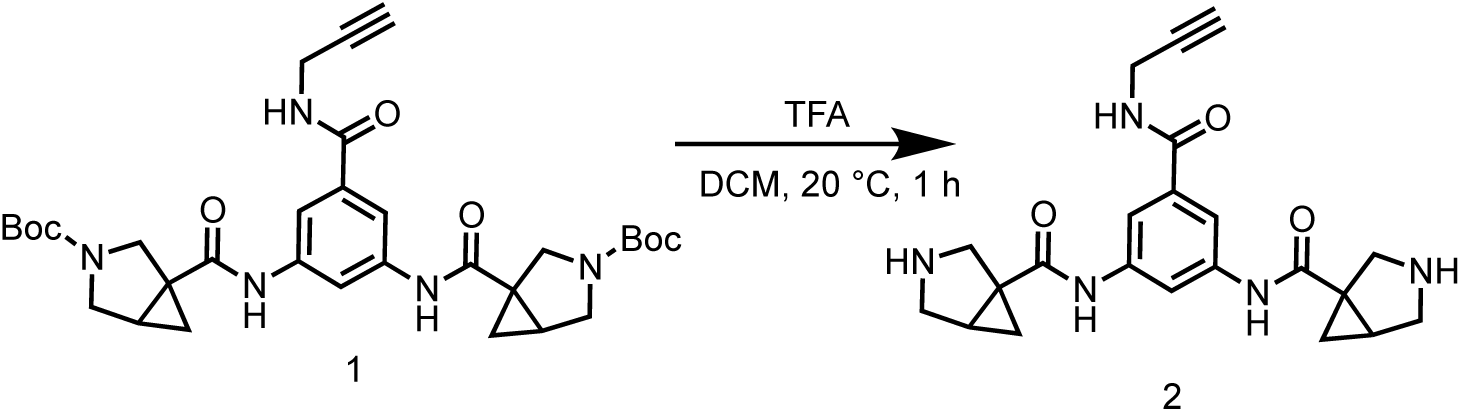

To a solution of di-*tert*-butyl 1,1’-(((5-(prop-2-yn-1-ylcarbamoyl)-1,3-phenylene)bis (azanediyl))bis(carbonyl))bis(3-azabicyclo[3.1.0]hexane-3-carboxylate) (1.20 g, 1.97 mmol) in DCM (20 mL) was added TFA (3.07 g, 26.9 mmol, 2 mL) at 0 °C, the mixture was stirred 20 °C for 1 hr. LCMS showed the reaction was completed. The mixture was concentrated in vacuum to give a residue. The residue was purified by preparative-HPLC (column: Waters Xbridge C18 150*50mm* 10um; mobile phase: [water(NH_3_H_2_O+NH_4_HCO_3_)-MeOH]; gradient:1%-29% B over 11 min) to give *N*,*N*’-(5-(prop-2-yn-1-ylcarbamoyl)-1,3-phenylene)bis(3-azabicyclo[3.1.0]hexane-1-carboxamide) (165 mg, 404 μmol, 20.5% yield, 99.5% purity) as a white solid.

LC-MS: 408.2 [M+H]^+^ / Ret time: 0.419 min / method: 5-95N_1min.lcm.

^1^H NMR (400 MHz, DMSO-*d*_6_): *δ* 9.60-9.21 (m, 2H), 8.80 (s, 1H), 8.13 (s, 1H), 7.72 (s, 2H), 4.01 (m, *J* = 5.4 Hz, 2H), 3.86 (s, 5H), 3.12-3.09 (m, 1H), 2.79 (s, 3H), 2.16-1.85 (m, 2H), 1.23 (d, *J* = 3.6 Hz, 2H), 1.00-0.72 (m, 2H).

#### Step 3: Preparation of *N,N*’-(5-(prop-2-yn-1-ylcarbamoyl)-1,3-phenylene)bis(3-(2-chloroacetyl)-3-azabicyclo[3.1.0]hexane-1-carboxamide) (SH-X-85)

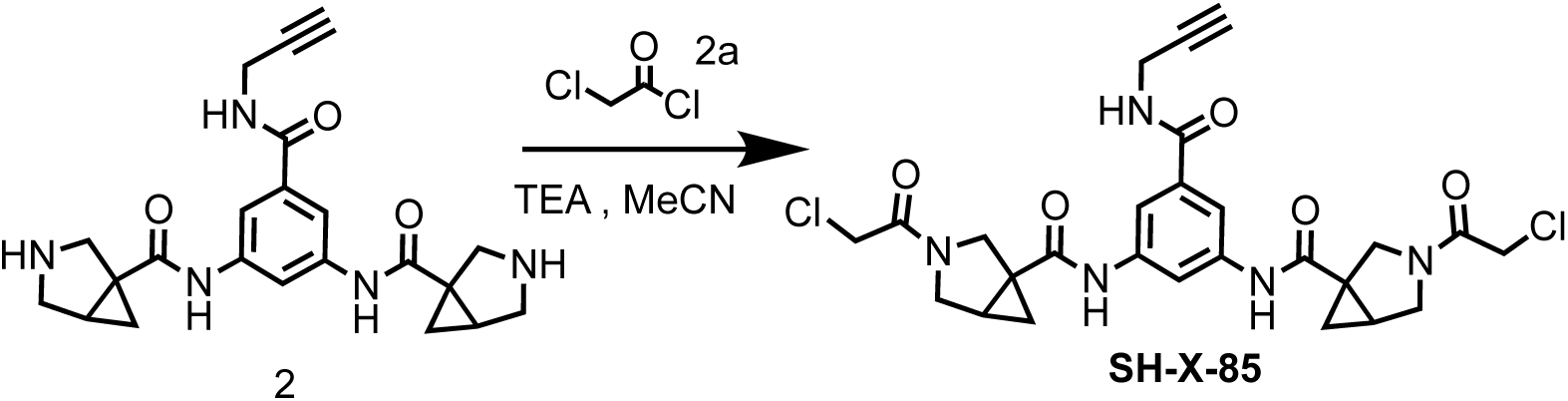

To a mixture of *N,N’*-(5-(prop-2-yn-1-ylcarbamoyl)-1,3-phenylene)bis(3-azabicyclo[3.1.0] hexane-1-carboxamide) (700 mg, 1.72 mmol) and TEA (1.04 g, 10.3 mmol, 1.43 mL) in MeCN (30 mL) was added 2-chloroacetyl chloride (427 mg, 3.80 mmol, 301 μL), the mixture was stirred at 0 °C for 0.5 hr under N_2_. LCMS showed the reaction was completed. The reaction mixture was extracted with H_2_O (5 mL) and ethyl acetate (5 mL×3), washed with brine (5 mL), the organic phase was dried over Na_2_SO_4_, concentrated in vacuum to give a residue. The residue was purified by preparative-HPLC (column: Waters Xbridge C18 150*50mm* 10um; mobile phase: [column: Phenomenex luna C18 150*25mm* 10um;mobile phase: [water(FA)-MeCN]; gradient:16%-46% B over 10 min) to give *N,N’*-(5-(prop-2-yn-1-ylcarbamoyl)-1,3-phenylene)bis(3-(2-chloroacetyl)-3-azabicyclo[3.1.0]hexane-1-carboxamide) (94.7 mg, 157 μmol, 9.13% yield, 92.9% purity) as a white solid.

LC-MS: 559.9 [M+H]^+^ / Ret time: 0.328 min / method: 5-95AB_0.8min.lcm.

^1^H NMR (400 MHz, DMSO-*d*_6_): *δ* 9.62 (d, *J* = 3.6 Hz, 1H), 9.50 (d, *J* = 3.6 Hz, 1H), 8.82 (d, *J* = 6.0 Hz, 1H), 8.25-8.13 (m, 1H), 7.74 (m, *J* = 1.6 Hz, 2H), 4.36-4.25 (m, 4H), 4.04-3.86 (m, 5H), 3.78-3.67 (m, 4H), 3.46-3.42 (m, 1H), 3.15-3.08 (m, 1H), 2.30-2.18 (m, 2H), 1.56-1.46 (m, 2H), 0.86-0.75 (m, 2H).

### Synthesis of N,N’-((4-(benzyloxy)-3-methoxyphenyl)methylene)bis(2-chloroacetamide) (SH-X-1C9)

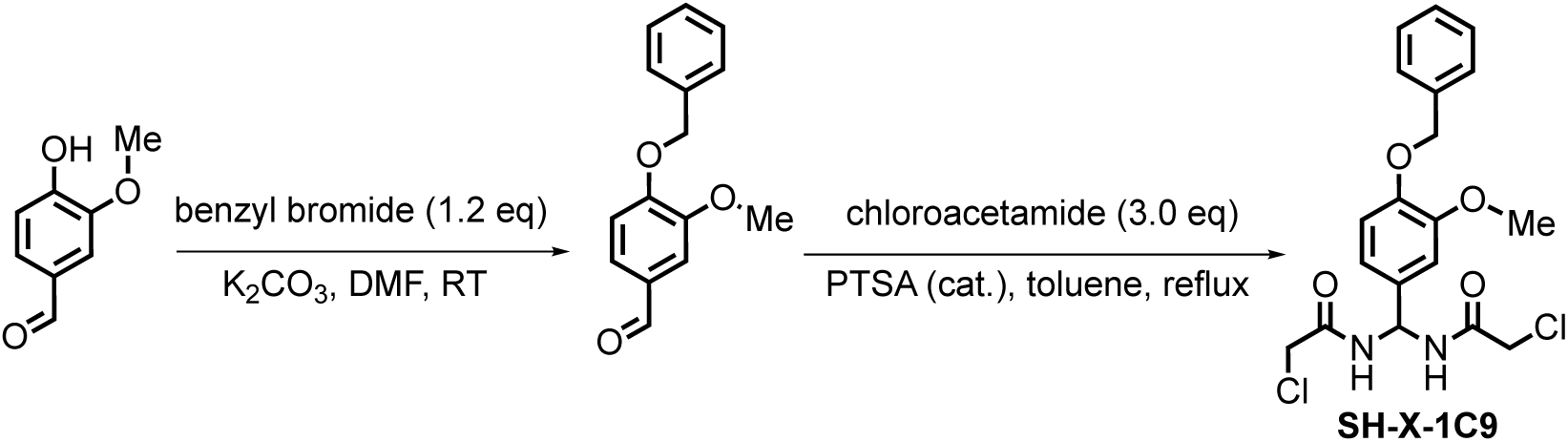

In a round bottom flask, 4-hydroxy-3-methoxybenzaldehyde (1.00 g, 6.57 mmol) was combined with benzyl bromide (1.20 g, 7.00 mmol), K_2_CO_3_ (3.00 g), and DMF (15 mL), along with a stir bar. This mixture was stirred at room temperature overnight. Subsequently, water (50 mL) and diethyl ether (50 mL) were added, and the contents were transferred to a separatory funnel. The organic layer was isolated, dried over magnesium sulfate, and concentrated to obtain a crude mixture, which was used directly without further purification.

This crude mixture was then redissolved in toluene (20 mL), and chloroacetamide (1.88 g, 19.7 mmol) along with PTSA (50 mg) were added. The mixture was refluxed overnight, cooled to room temperature, and the solid precipitate (comprising the product and excess chloroacetamide) was filtered off and rinsed with diethyl ethyl ether and ethyl acetate. The remaining chloroacetamide was dissolved by sonicating the crude solid in water, yielding N,N’-((4-(benzyloxy)-3-methoxyphenyl)methylene)bis(2-chloroacetamide). The product was isolated by filtration and dried under vacuum.

^1^H NMR (400 MHz, DMSO-*D*_6_) δ 8.88 (d, *J* = 7.9 Hz, 2H), 7.40-7.26 (m, 5H), 6.98 (d, *J* = 8.3 Hz, 1H), 6.94 (d, *J* = 2.2 Hz, 1H), 6.81 (dd, *J* = 8.2, 2.1, 1H), 6.45 (t, *J* = 7.9 Hz, 1H), 5.06 (s, 2H), 4.16 – 4.03 (m, 4H), 3.74 (s, 3H).

### Synthesis of *N,N*’-(5-(prop-2-yn-1-ylcarbamoyl)-1,3-phenylene)bis(1-(2-chloroacetyl)pyrrolidine-3-carboxamide) (SH-X-69)

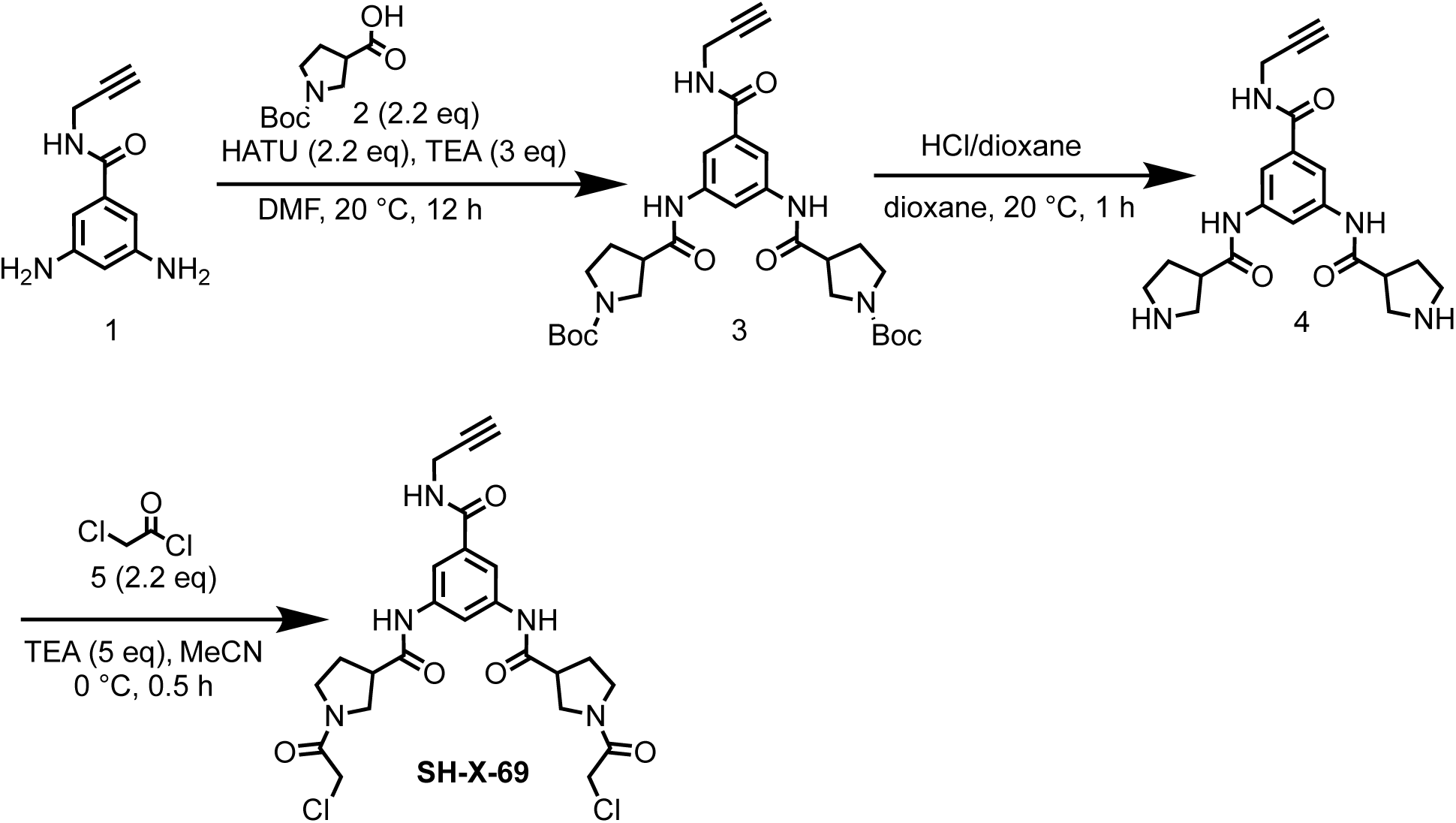

#### Step 1: Preparation of di-*tert*-butyl 3,3’-(((5-(prop-2-yn-1-ylcarbamoyl)-1,3-phenylene)bis(azanediyl))bis(carbonyl))bis(pyrrolidine-1-carboxylate) (3)

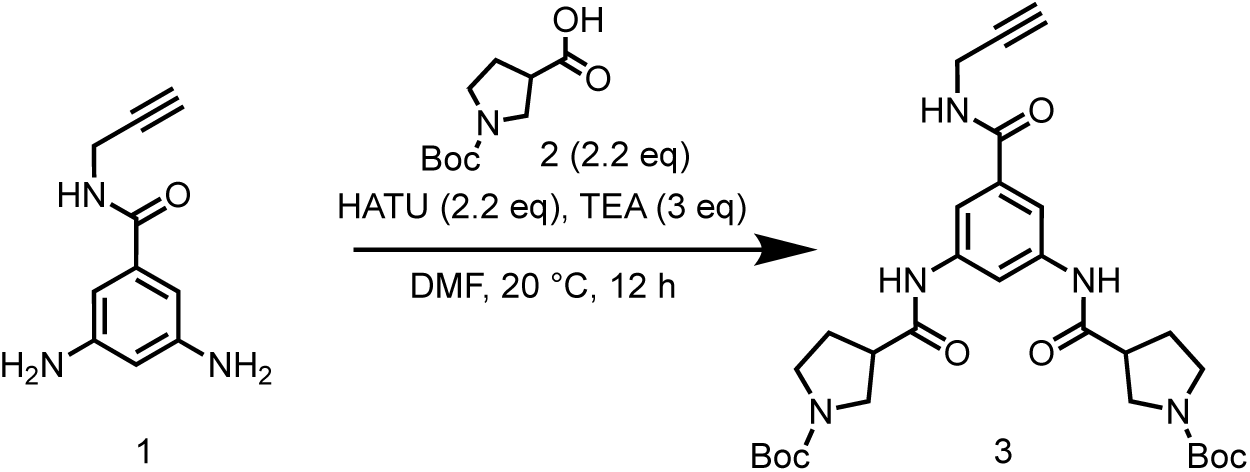

To a solution of 1-(*ter*t-butoxycarbonyl)pyrrolidine-3-carboxylic acid (1.00 g, 4.65 mmol) in DMF (10 mL) was added HATU (3.89 g, 10.2 mmol) and TEA (1.41 g, 14.0 mmol, 1.94 mL), then 3,5-diamino-*N*-(prop-2-yn-1-yl)benzamide (396 mg, 2.09 mmol) was added into the reaction mixture, the reaction mixture was stirred at 20 °C for 12 hrs. LCMS showed the reaction was completed. The reaction mixture was poured into H_2_O (50 mL), extracted with EtOAc (50 mL×3). The organic layers were collected and washed with brine (50 mL×3), dried over Na_2_SO_4_ and concentrated to give a residue. The residue was purified by reversed-phase HPLC (0.1% NH_3_•H_2_O) to give di-*tert*-butyl 3,3’-(((5-(prop-2-yn-1-ylcarbamoyl)-1,3-phenylene)bis(azanediyl))bis(carbonyl))bis(pyrrolidine-1-carboxylate) (380 mg, 651. μmol, 14.0% yield) as a brown solid.

LC-MS: 484.4 [M+H-100]^+^ / Ret time: 0.479 min / method: 5-95AB_0.8min.lcm.

^1^H NMR (400 MHz, DMSO-*d*_6_): *δ* 10.20 (s, 2H), 8.86 (s, 1H), 8.12 (s, 1H), 7.70 (s, 2H), 4.00 (dd, *J* = 2.4, 5.2 Hz, 2H), 3.55-3.48 (m, 2H), 3.39 (d, *J* = 10.4 Hz, 2H), 3.36 (d, *J* = 7.2 Hz, 2H), 3.29-3.21 (m, 2H), 3.19-3.11 (m, 2H), 3.10 (t, *J* = 2.4 Hz, 1H), 2.10 (d, *J* = 6.0 Hz, 2H), 2.01 (d, *J* = 7.2 Hz, 2H), 1.40 (s, 18H).

#### Step 2: Preparation of *N,N*’-(5-(prop-2-yn-1-ylcarbamoyl)-1,3-phenylene)bis(pyrrolidine-3-carboxamide) (4)

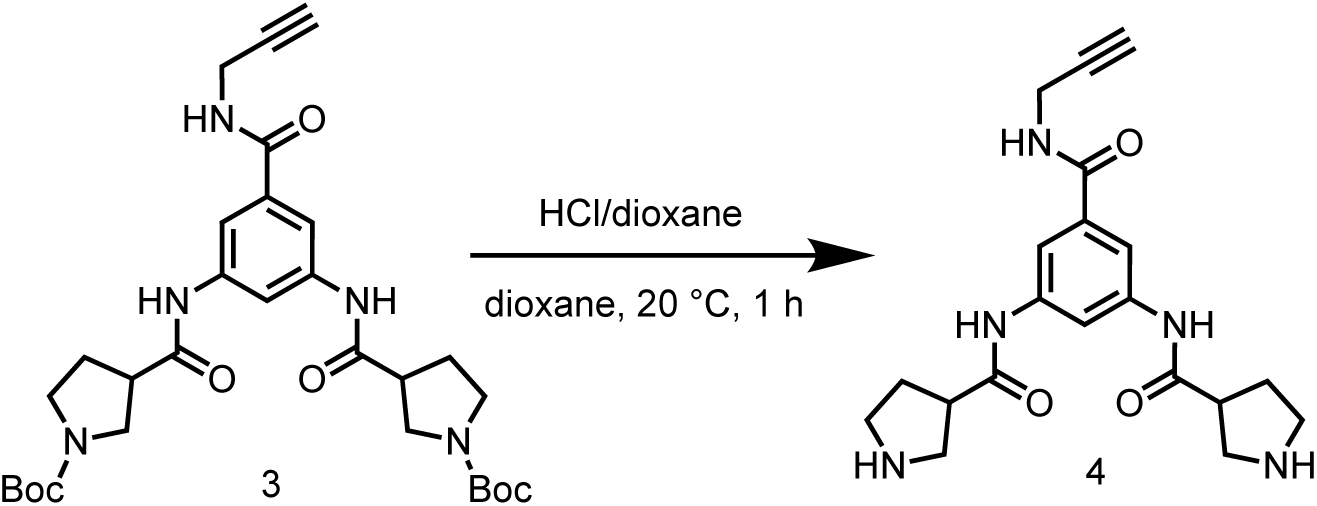

To a solution of di-*tert*-butyl 3,3’-(((5-(prop-2-yn-1-ylcarbamoyl)-1,3-phenylene)bis (azanediyl))bis(carbonyl))bis(pyrrolidine-1-carboxylate) (380 mg, 651μmol) in dioxane (2 mL) was added HCl/dioxane (4 M, 2 mL), the mixture was stirred at 20 °C for 1 hr. LCMS showed the reaction was completed. The reaction mixture was concentrated in vacuum to give *N*,*N*’-(5-(prop-2-yn-1-ylcarbamoyl)-1,3-phenylene)bis(pyrrolidine-3-carboxamide) (220 mg, crude, HCl salt) as an orange solid.

LC-MS: 384.2 [M+H]^+^ / Ret time: 0.257 min / method: 0-60AB_0.8min.lcm.

#### Step 3: Preparation of *N,N*’-(5-(prop-2-yn-1-ylcarbamoyl)-1,3-phenylene)bis(1-(2-chloroacetyl)pyrrolidine-3-carboxamide) (SH-X-69)

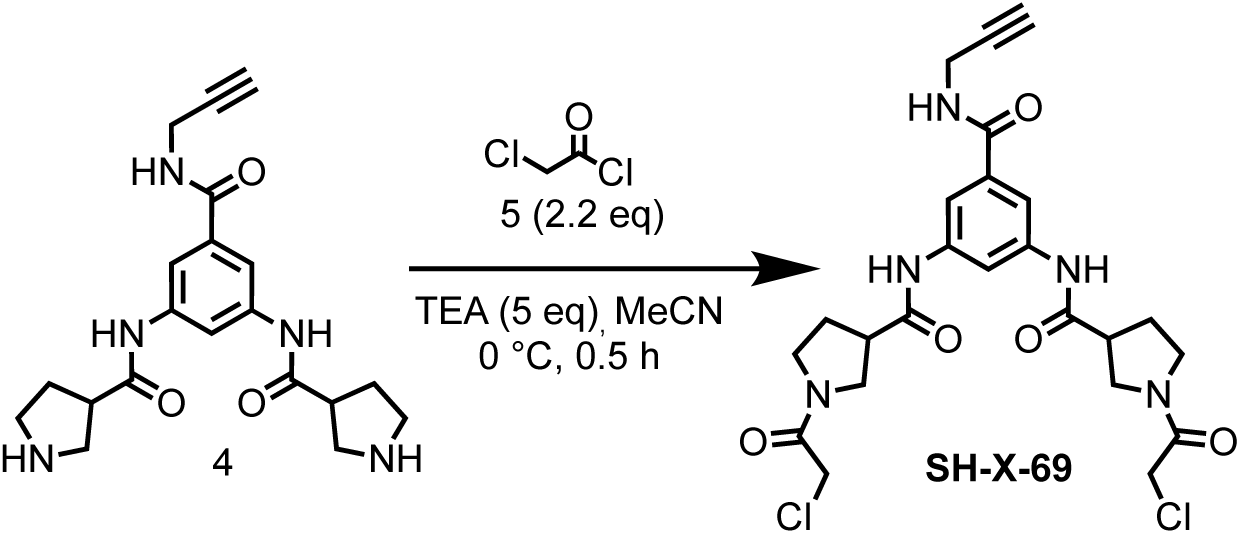

To a solution of *N*,*N*’-(5-(prop-2-yn-1-ylcarbamoyl)-1,3-phenylene)bis(pyrrolidine-3-carboxamide) (200 mg, 476 μmol, HCl) in MeCN (20 mL) was added TEA (241 mg, 2.38 mmol, 331 μL), then 2-chloroacetyl chloride (118 mg, 1.05 mmol, 83.5 μL) was droppwised into the mixture, the reaction mixture was stirred at 0 °C for 0.5 hr. LCMS showed the reaction was completed. The reaction mixture was quenched with H_2_O (0.1 mL) and concentrated in vacuum to give a residue. The residue was purified by preparative-HPLC (column: Phenomenex luna C18 150*25mm* 10um; mobile phase: [water (FA)-MeCN]; gradient: 12%-42% B over 10 min) to give *N,N’*-(5-(prop-2-yn-1-ylcarbamoyl)-1,3-phenylene)bis(1-(2-chloroacetyl)pyrrolidine-3-carboxamide) (50 mg, 93.2 μmol, 19.6% yield) as a white solid.

LC-MS: 536.3 [M+H]^+^ / Ret time: 0.317 min / method: 5-95AB_0.8min.lcm

^1^H NMR (400 MHz, DMSO-*d*_6_) *δ* 10.27 (d, *J* = 9.2 Hz, 2H), 8.86 (t, *J* = 5.2 Hz, 1H), 8.14 (s, 1H), 7.72 (s, 2H), 4.31 (s, 4H), 4.03 - 3.98 (m, 2H), 3.81 - 3.73 (m, 1H), 3.65 (m, *J* = 8.0, 11.2 Hz, 3H), 3.57 - 3.44 (m, 4H), 3.28 - 3.22 (m, 1H), 3.16 (t, *J* = 7.2 Hz, 1H), 3.10 (s, 1H), 2.27 - 2.18 (m, 1H), 2.17 - 2.07 (m, 2H), 2.05 - 1.94 (m, 1H).

### Synthesis of *N,N*’-(5-(prop-2-yn-1-ylcarbamoyl)-1,3-phenylene)bis(3-(2-chloroacetamido)bicyclo[1.1.1]pentane-1-carboxamide) (SH-X-81)

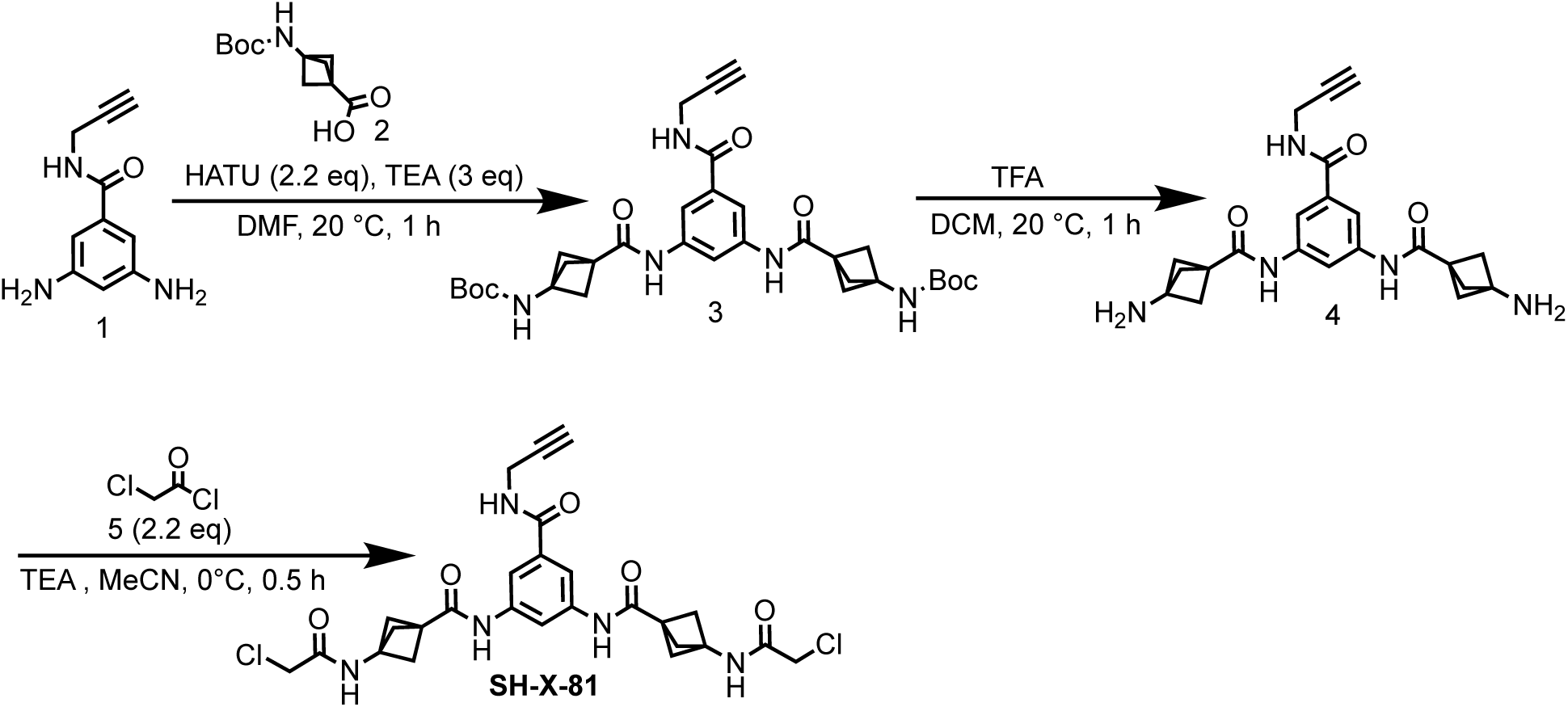

#### Step 1: Preparation of di-*tert*-butyl ((((5-(prop-2-yn-1-ylcarbamoyl)-1,3-phenylene)bis(azanediyl))bis(carbonyl))bis(bicyclo[1.1.1]pentane-3,1-diyl))dicarbamate (3)

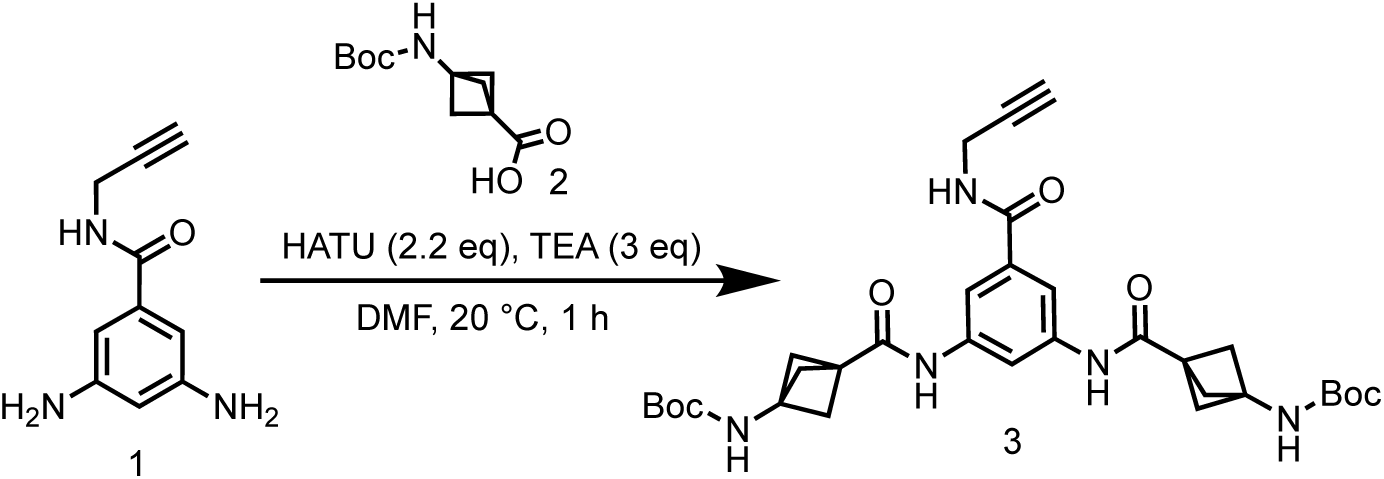

To a mixture of 3-((*tert*-butoxycarbonyl)amino)bicyclo[1.1.1]pentane-1-carboxylic acid (793 mg, 3.49 mmol) and 3,5-diamino-*N*-(prop-2-yn-1-yl)benzamide (300 mg, 1.59 mmol) in DMF (8 mL) was added HATU (1.33 g, 3.49 mmol) and TEA (481 mg, 4.76 mmol, 662 μL), the reaction mixture was stirred at 20 °C for 1 hr. LCMS showed the reaction was completed. The reaction mixture was poured into H_2_O (50 mL), extracted with EtOAc (50 mL×3). The organic layers were collected and washed with brine (50 mL×3), dried over Na_2_SO_4_ and concentrated in vacuum to give a residue. The residue was purified by reversed-phase HPLC (0.1% FA condition) to give di-*tert*-butyl ((((5-(prop-2-yn-1-ylcarbamoyl)-1,3-phenylene) bis(azanediyl))bis(carbonyl))bis(bicyclo[1.1.1]pentane-3,1-diyl))dicarbamate (800 mg, 1.32 mmol, 83.0% yield) as a white solid.

LC-MS: 496.2 [M+H-112]^+^ / Ret time: 0.461 min / method: 5-95AB_0.8min.lcm.

^1^H NMR (400 MHz, DMSO-*d*_6_): *δ* 9.70 (s, 2H), 8.81 (t, *J* = 5.6 Hz, 1H), 8.17 (s, 1H), 7.73 (d, *J* = 1.6 Hz, 2H), 7.57 (s, 1H), 4.00 (m, *J* = 2.4, 5.2 Hz, 2H), 3.08 (t, *J* = 2.4 Hz, 1H), 2.19 (s, 12H), 1.39 (s, 18H).

#### Step 2: Preparation of *N,N*’-(5-(prop-2-yn-1-ylcarbamoyl)-1,3-phenylene)bis(3-aminobicyclo[1.1.1]pentane-1-carboxamide) (4)

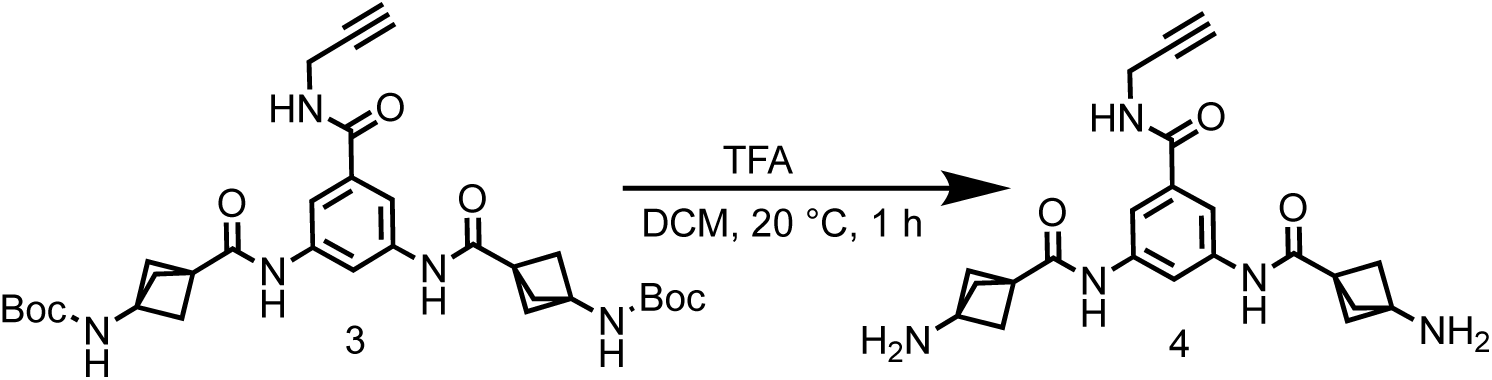

To a solution of di-*tert*-butyl ((((5-(prop-2-yn-1-ylcarbamoyl)-1,3-phenylene)bis(azanediyl)) bis(carbonyl))bis(bicyclo[1.1.1]pentane-3,1-diyl))dicarbamate (700 mg, 1.15 mmol) in DCM (7 mL) was added TFA (10.8 g, 94.2 mmol, 7 mL). The mixture was stirred at 20 °C for 1 hr. LCMS showed the reaction was completed. The mixture was concentrated in vacuum to give *N*,*N*’-(5-(prop-2-yn-1-ylcarbamoyl)-1,3-phenylene)bis(3-aminobicyclo[1.1.1]pentane-1-carboxamide) (1.1 g, crude, TFA salt) as a brown oil.

LC-MS: 408.2 [M+H]^+^ / Ret time: 0.229 min / method: 0-60AB_0.8min.lcm.

#### Step 3: Preparation of *N,N*’-(5-(prop-2-yn-1-ylcarbamoyl)-1,3-phenylene)bis(3-(2-chloroacetamido)bicyclo[1.1.1]pentane-1-carboxamide) (SH-X-81)

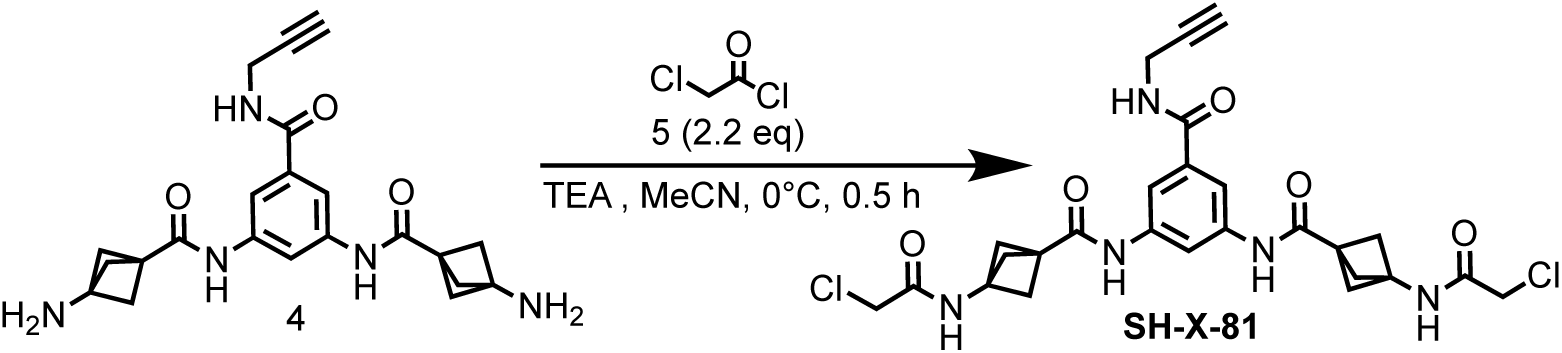

To a mixture of *N*,*N*’-(5-(prop-2-yn-1-ylcarbamoyl)-1,3-phenylene)bis(3-aminobicyclo[1.1.1] pentane-1-carboxamide) (1.1 g, 2.11 mmol, TFA salt) and TEA (1.07 g, 10.6 mmol, 1.47 mL) in MeCN (44 mL) was added 2-chloroacetyl chloride (524 mg, 4.64 mmol, 370 μL) at 0 °C. The reaction mixture was stirred at 0 °C for 0.5 hr. LCMS showed the reaction was completed. The reaction mixture was quenched with water (1 mL) and filtered, the filtrate was concentrated in vacuum to remove solvent, then extracted with EtOAc (20 mL×3) and H_2_O (20 mL). The organic layers were collected and dried over Na_2_SO_4_ then concentrated in vacuum to give *N*,*N*’-(5-(prop-2-yn-1-ylcarbamoyl)-1,3-phenylene)bis(3-(2-chloroacetamido) bicyclo[1.1.1]pentane-1-carboxamide) (340 mg, 607 μmol, 28.8% yield) as a brown oil.

^1^H NMR (400 MHz, DMSO-*d*_6_) *δ* 9.79 (s, 2H), 8.89 (s, 2H), 8.86 - 8.80 (m, 1H), 8.19 (s, 1H), 7.73 (d, *J* = 1.6 Hz, 2H), 4.03 - 3.98 (m, 6H), 3.10 (t, *J* = 2.4 Hz, 1H), 2.30 (s, 12H).

LC-MS: 560.2 [M+H]^+^ / Ret time: 0.346 min / method: 5-95AB_0.8min.lcm.

### Synthesis of *N,N*’-(5-(prop-2-yn-1-ylcarbamoyl)-1,3-phenylene)bis(1-(2-chloroacetyl)-5,5-difluoropiperidine-3-carboxamide) (SH-X-83)

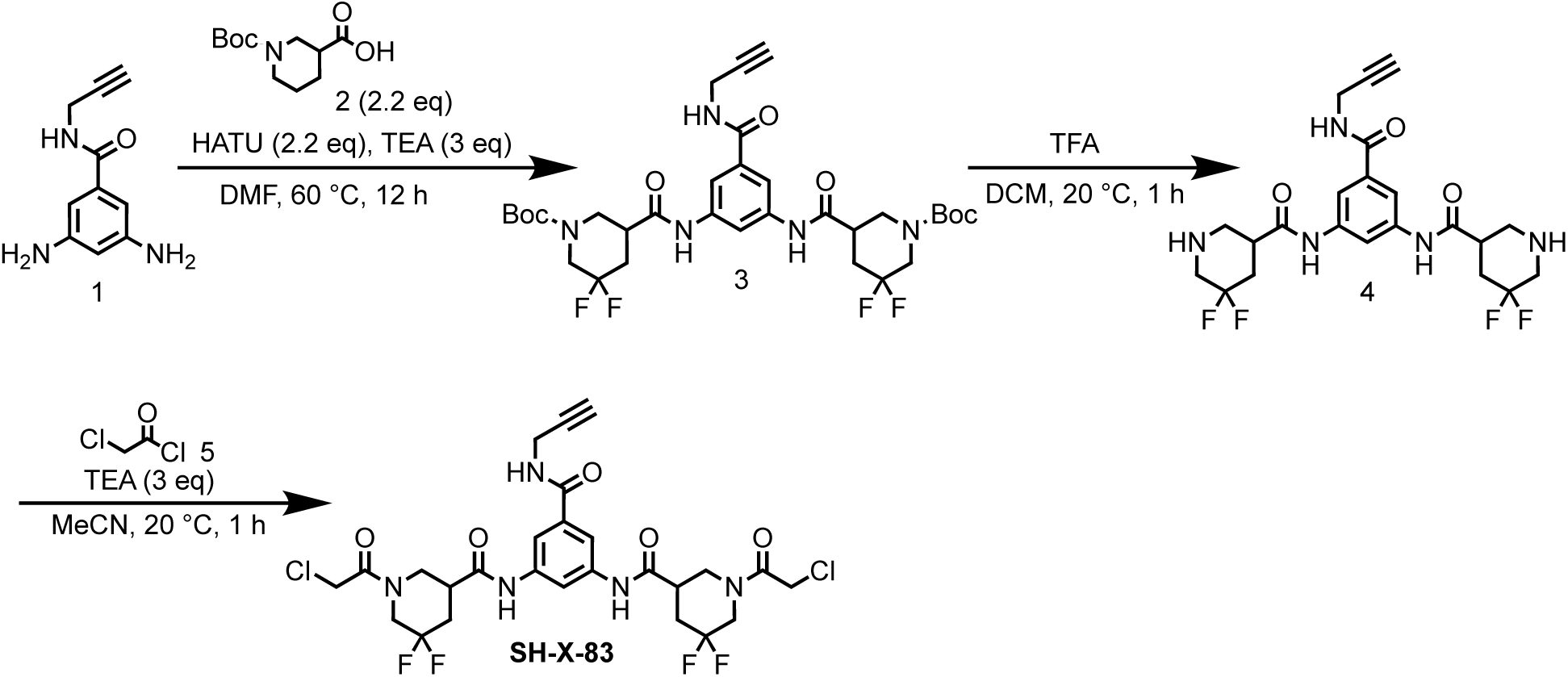

#### Step 1: Preparation of di-*tert*-butyl 5,5’-(((5-(prop-2-yn-1-ylcarbamoyl)-1,3-phenylene)bis(azanediyl))bis(carbonyl))bis(3,3-difluoropiperidine-1-carboxylate) (3)

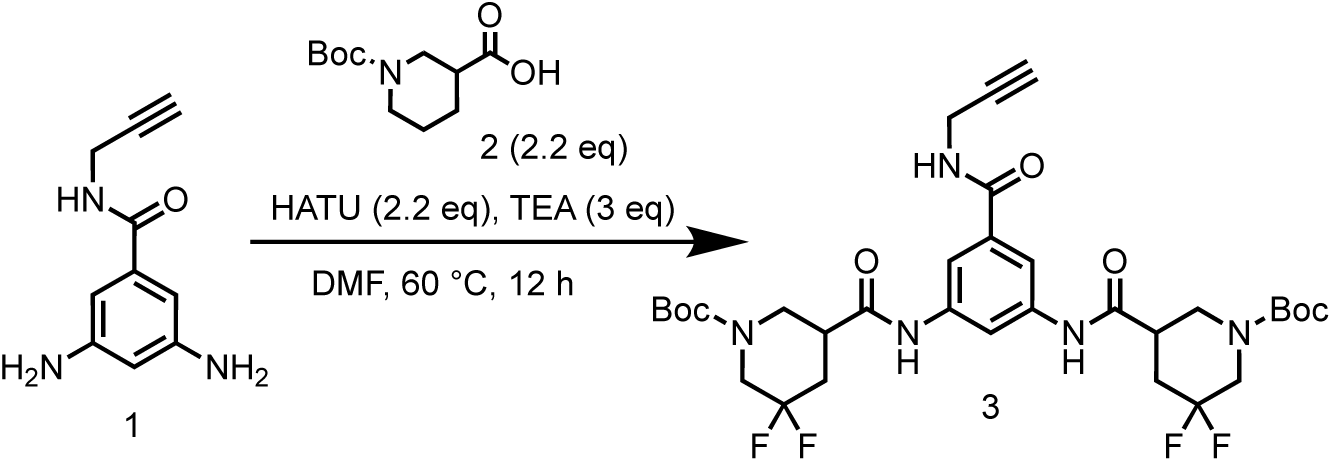

To a solution of 1-(*tert*-butoxycarbonyl)piperidine-3-carboxylic acid (280 mg, 1.48 mmol) in DMF (1 mL) was added HATU (1.24 g, 3.26 mmol) and TEA (449 mg, 4.44 mmol, 618 μL), 3,5-diamino-*N*-(prop-2-yn-1-yl)benzamide (863 mg, 3.26 mmol). The mixture was stirred at 60 °C for 12 hrs. LCMS showed the reaction was completed. The reaction mixture was poured into H_2_O (50 mL), extracted with EtOAc (50 mL×3). The organic layers were collected and washed with brine (50 mL×3), dried over Na_2_SO_4_ and concentrated to give a residue. The residue was purified by column chromatography on silica gel (eluted with petroleum ether: ethyl acetate=10:0 to 1:1) to give di-*tert*-butyl 5,5’-(((5-(prop-2-yn-1-ylcarbamoyl)-1,3-phenylene)bis(azanediyl))bis(carbonyl)) bis(3,3-difluoropiperidine-1-carboxylate) (200 mg, 292 μmol, 19.8% yield) as a white solid.

LC-MS: 528.1 [M-156+H]^+^ / Ret time: 0.529 min / method: 5-95AB_0.8min.lcm

#### Step 2: Preparation of *N,N*’-(5-(prop-2-yn-1-ylcarbamoyl)-1,3-phenylene)bis(5,5-difluoropiperidine-3-carboxamide) (4)

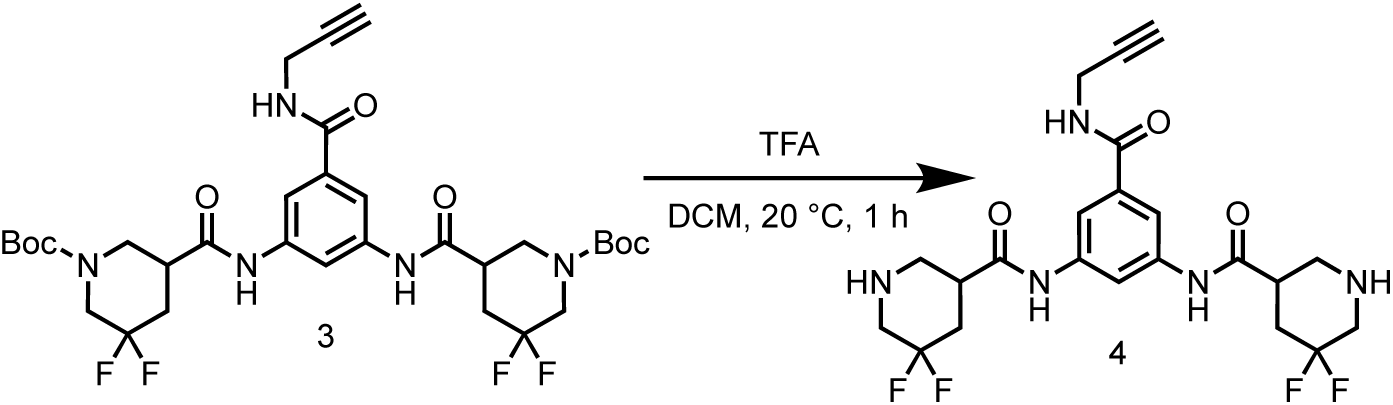

To a solution of di-*tert*-butyl 5,5’-(((5-(prop-2-yn-1-ylcarbamoyl)-1,3-phenylene) bis(azanediyl))bis(carbonyl))bis(3,3-difluoropiperidine-1-carboxylate) (200 mg, 292 μmol) in DCM (0.5 mL) was added TFA (0.5 mL). The mixture was stirred at 20 °C for 1 hr. LCMS showed the reaction was completed. The reaction mixture was evaporated in vacuum to give *N,N’*-(5-(prop-2-yn-1-ylcarbamoyl)-1,3-phenylene)bis (5,5-difluoropiperidine-3-carboxamide) (150 mg, crude) as a yellow solid.

LC-MS: 484.2 [M+H]^+^ / Ret time: 0.234 min / method: 5-95AB_0.8min.lcm

#### Step 3: Synthesis of *N,N*’-(5-(prop-2-yn-1-ylcarbamoyl)-1,3-phenylene)bis(1-(2-chloroacetyl)-5,5-difluoropiperidine-3-carboxamide) (SH-X-083)

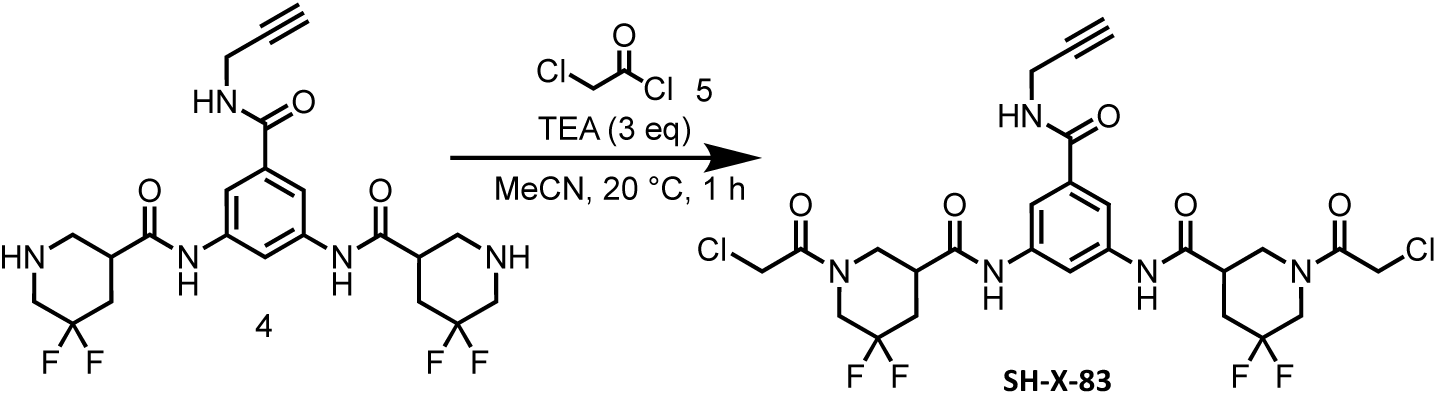

To a solution of *N,N’*-(5-(prop-2-yn-1-ylcarbamoyl)-1,3-phenylene)bis(5,5-difluoropiperidine-3-carboxamide) (150 mg, 310 μmol) in MeCN (4 mL) was added TEA (94.2 mg, 931 μmol, 129 μL) and 2-chloroacetyl chloride (70.1 mg, 620 μmol, 49.4 μL). The mixture was stirred at 20 °C for 1 hr. LCMS showed the reaction was completed. The reaction mixture was concentrated in vacuum to give a residue. The residue was purified by preparative-HPLC (column: Welch Ultimate C18 150*25mm*5um;mobile phase: [water(FA)-MeCN];gradient:24%-54% B over 10 min) to give *N*,*N*’-(5-(prop-2-yn-1-ylcarbamoyl)-1,3-phenylene)bis(1-(2-chloroacetyl)-5,5-difluoropiperidine-3-carboxamide) (8.60 mg, 13.5 μmol, 4.36% yield) as a white solid.

LC-MS: 636.1 [M+H]^+^ / Ret time: 0.410 min / method: 5-95AB_0.8min.lcm

^1^H NMR (400 MHz, DMSO-*d*_6_): *δ* 10.45-10.29 (m, 2H), 8.90 (s, 1H), 8.12 (d, *J* = 6.0 Hz, 1H), 7.72 (d, *J* = 1.6 Hz, 2H), 4.60 (d, *J* = 6.0 Hz, 2H), 4.57-4.38 (m, 4H), 4.25-4.12 (m, 1H), 4.03-3.97 (m, 3H), 3.74-3.55 (m, 1H), 3.39-3.36 (m, 1H), 3.29-3.24 (m, 1H), 3.11 (s, 1H), 2.99-2.75 (m, 3H), 2.47-2.37 (m, 2H), 2.36-2.29 (m, 1H), 2.28-2.18 (m, 1H).

### Synthesis of *N,N*’-(5-(((1-(2-(2-(6-((4*R*,5*S*)-5-methyl-2-oxoimidazolidin-4-yl)hexanamido)ethoxy)ethyl)-1*H*-1,2,3-triazol-4-yl)methyl)carbamoyl)-1,3-phenylene)bis(3-(2-chloroacetamido)bicyclo[1.1.1]pentane-1-carboxamide) (SH-X-65)

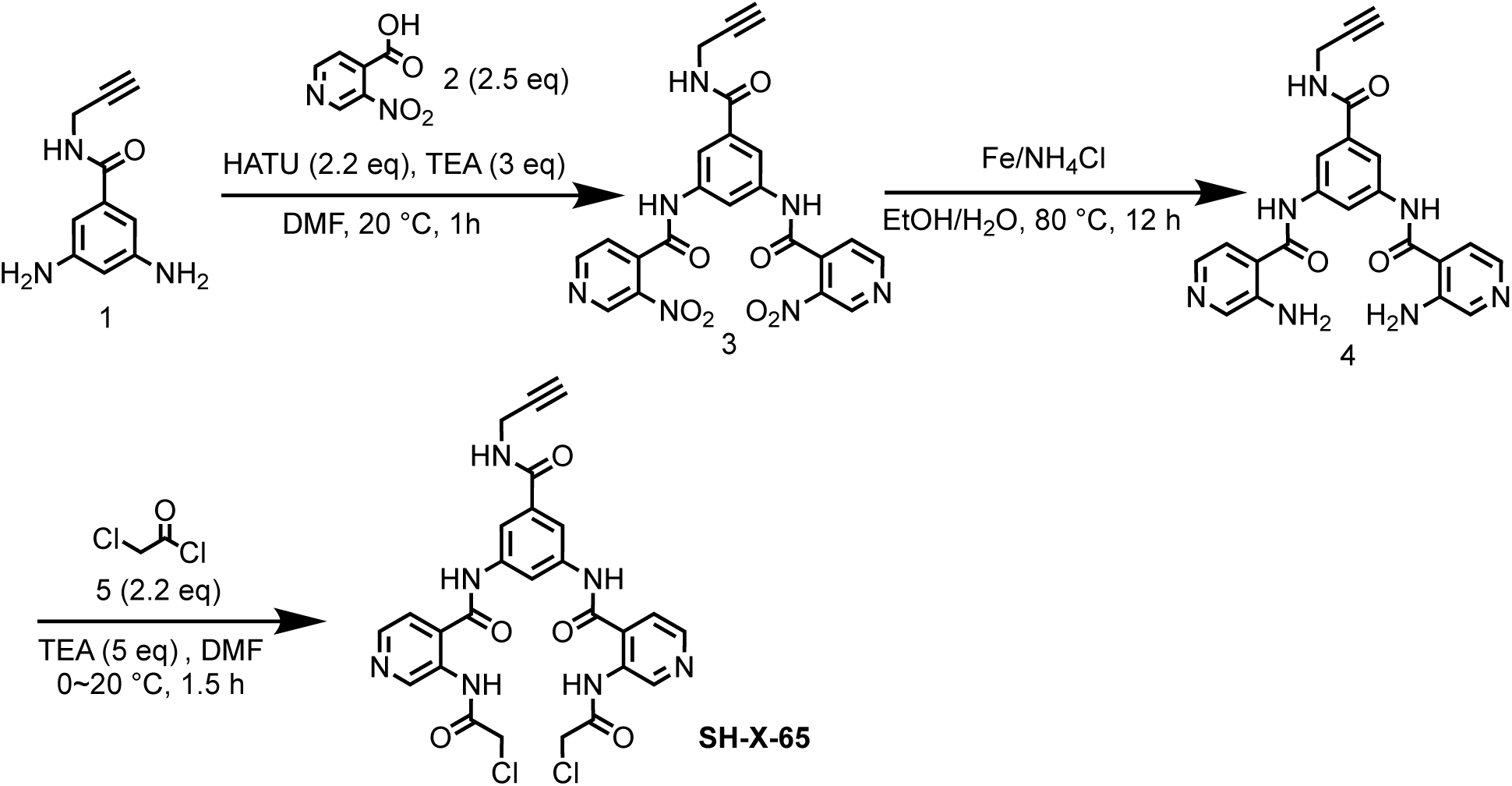

#### Step 1: Preparation of *N,N*’-(5-(prop-2-yn-1-ylcarbamoyl)-1,3-phenylene)bis(3-nitroisonicotinamide) (3)

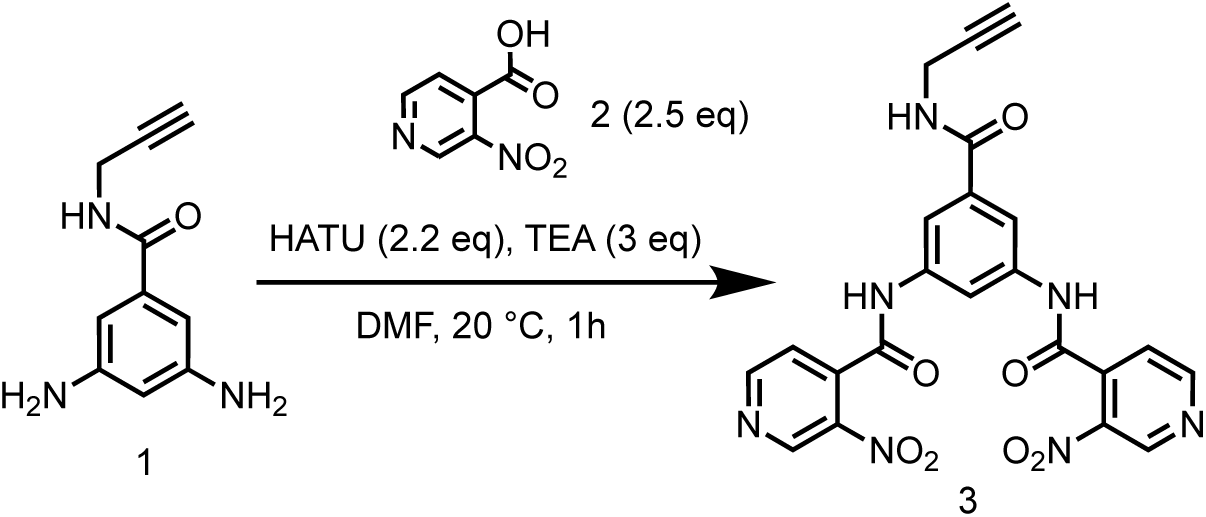

To a solution of 3-nitroisonicotinic acid (1 g, 5.95 mmol) and 3,5-diamino-*N*-(prop-2-yn-1-yl) benzamide (450 mg, 2.38 mmol) in DMF (10 mL) was added HATU (1.99 g, 5.23 mmol,) and TEA (722 mg, 7.13 mmol, 993 μL), the reaction mixture was stirred at 20 °C for 1 hr. LCMS showed the reaction was completed. The reaction mixture was poured into H_2_O (50 mL), extracted with EtOAc (50 mL×3). The organic layers were collected and washed with brine (50 mL×3), dried over Na_2_SO_4_ and concentrated to give a residue. The residue was dissolved in EtOAc (10 mL), then filtered, the filter cake was washed with MeOH (30 mL), then concentrated in vacuum to give *N*,*N*’-(5-(prop-2-yn-1-ylcarbamoyl)-1,3-phenylene)bis(3 nitroisonicotinamide) (590 mg, 1.21 mmol, 50.7% yield) as a yellow solid.

LC-MS: 490.2 [M+H]^+^ / Ret time: 0.353 min / method: 5-95AB_0.8min.lcm

^1^H NMR (400 MHz, DMSO-*d*_6_): *δ* 11.06 (s, 2H), 9.38 (s, 2H), 9.07 (d, *J* = 4.8 Hz, 3H), 8.24 (s, 1H), 7.89 (d, *J* = 4.8 Hz, 2H), 7.85 (d, *J* = 1.6 Hz, 2H), 4.04-4.01 (m, 2H), 3.12 (t, *J* = 2.4 Hz, 1H).

#### Step 2: Preparation of *N,N*’-(5-(prop-2-yn-1-ylcarbamoyl)-1,3-phenylene)bis(3-aminobicyclo[1.1.1]pentane-1-carboxamide) (4)

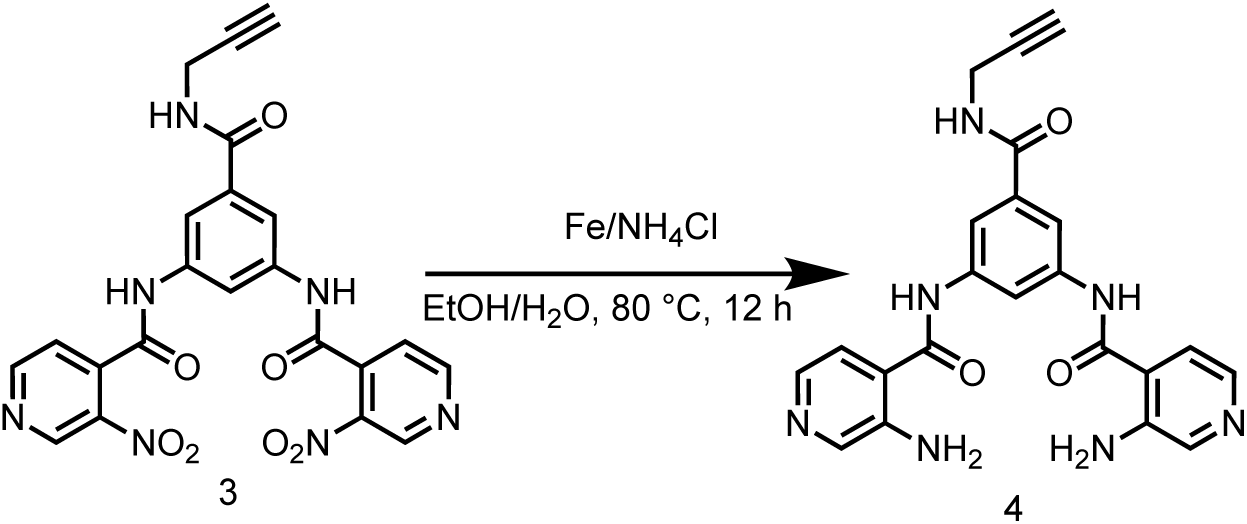

To a solution of *N*,*N*’-(5-(prop-2-yn-1-ylcarbamoyl)-1,3-phenylene)bis(3-nitroisonicotinamide) (490 mg, 1.00 mmol) in EtOH (10 mL) and H_2_O (2 mL) was added NH_4_Cl (536 mg, 10.0 mmol) and Fe (559 mg, 10.0 mmol), the mixture was stirred at 80 °C for 12 hrs. LCMS showed the reaction was completed. The reaction mixture was filtered, the filtrate was concentrated in vacuum to give *N,N*’-(5-(prop-2-yn-1-ylcarbamoyl)-1,3-phenylene)bis(3-aminoisonicotinamide) (430 mg, crude) as a yellow solid.

LC-MS: 430.2 [M+H]^+^ / Ret time: 0.282 min / method: 0-60AB_0.8min.lcm.

#### Step 3: Synthesis of *N,N*’-(5-(prop-2-yn-1-ylcarbamoyl)-1,3-phenylene)bis(3-(2-chloroacetamido)isonicotinamide) (SH-X-65)

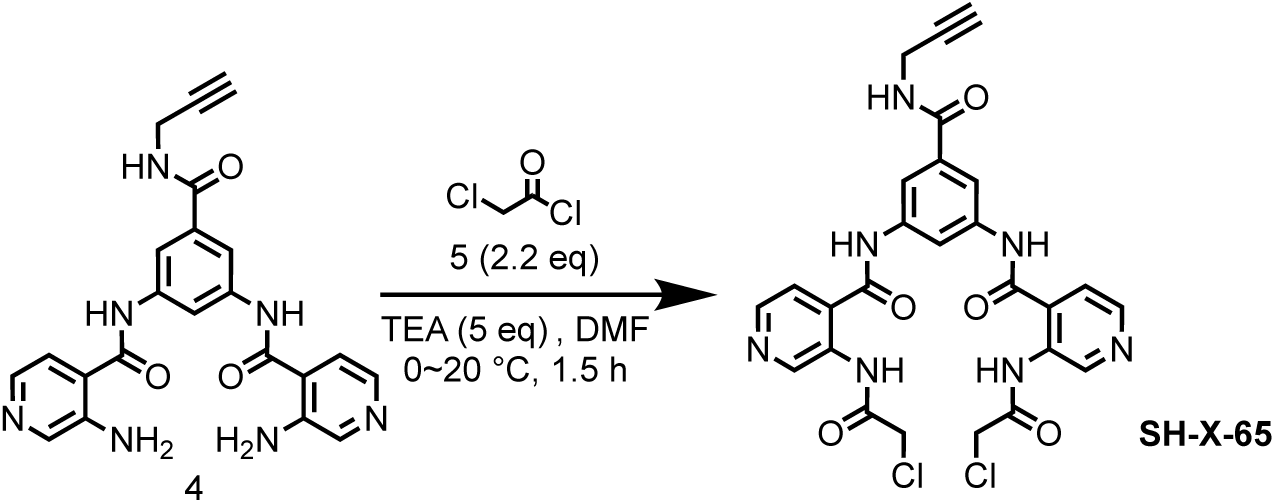

To a solution of *N,N*’-(5-(prop-2-yn-1-ylcarbamoyl)-1,3-phenylene)bis(3-aminoisonicotinamide) (200 mg, 466 μmol) in DMF (10 mL) was added TEA (236 mg, 2.33 mmol, 324 μL), then 2-chloroacetyl chloride (116 mg, 1.02 mmol, 81.6 μL) was added into the reaction mixture at 0 °C, the reaction mixture was stirred at 0 °C for 0.5 hr. LCMS showed 49% of *N,N*’-(5-(prop-2-yn-1-ylcarbamoyl)-1,3-phenylene)bis(3-aminoisonicotinamide) remained. 2-chloroacetyl chloride (116 mg, 1.02 mmol, 81.6 μL) was added into the reaction mixture, the reaction mixture was stirred at 20 °C for 1 hr. LCMS showed the reaction was completed. The reaction mixture was extracted with EtOAc (30 mL×3) and H_2_O (50 mL). The organic layers were collected and dried over Na_2_SO_4_ then concentrated to give a residue. The residue was purified by preparative-HPLC (column: Phenomenex luna C18 150*25mm* 10um; mobile phase: [water (FA)-MeCN]; gradient: 15%-45% B over 10 min). The purified solution was lyophilized to give *N,N*’-(5-(prop-2-yn-1-ylcarbamoyl)-1,3-phenylene)bis(3-(2-chloroacetamido)isonicotinamide) (6.7 mg, 10.3 μmol, 2.22% yield, 89.7% purity) as a yellow solid.

LC-MS: 582.1 [M+H]^+^ / Ret time: 0.337 min / method: 5-95AB_0.8min.lcm

^1^H NMR (400 MHz, DMSO-*d*_6_): *δ* 10.96-10.85 (m, 2H), 10.67 (s, 2H), 9.17 (s, 2H), 8.97-8.91 (m, 1H), 8.55 (d, *J* = 4.8 Hz, 2H), 8.26 (s, 1H), 7.93 (s, 2H), 7.72 (d, *J* = 5.2 Hz, 2H), 4.37 (s, 4H), 4.04 (d, *J* = 3.2 Hz, 2H), 3.13 (s, 1H).

### Synthesis of 1,1’-(((*S*)-2-((prop-2-yn-1-yloxy)methyl)piperazine-1,4-dicarbonyl)bis(isoindoline-1,2-diyl))bis(2-chloroethan-1-one) (SH-X-89A) and 1,1’-(((*R*)-2-((prop-2-yn-1-yloxy)methyl)piperazine-1,4-dicarbonyl)bis(isoindoline-1,2-diyl))bis(2-chloroethan-1-one) (SH-X-89B)

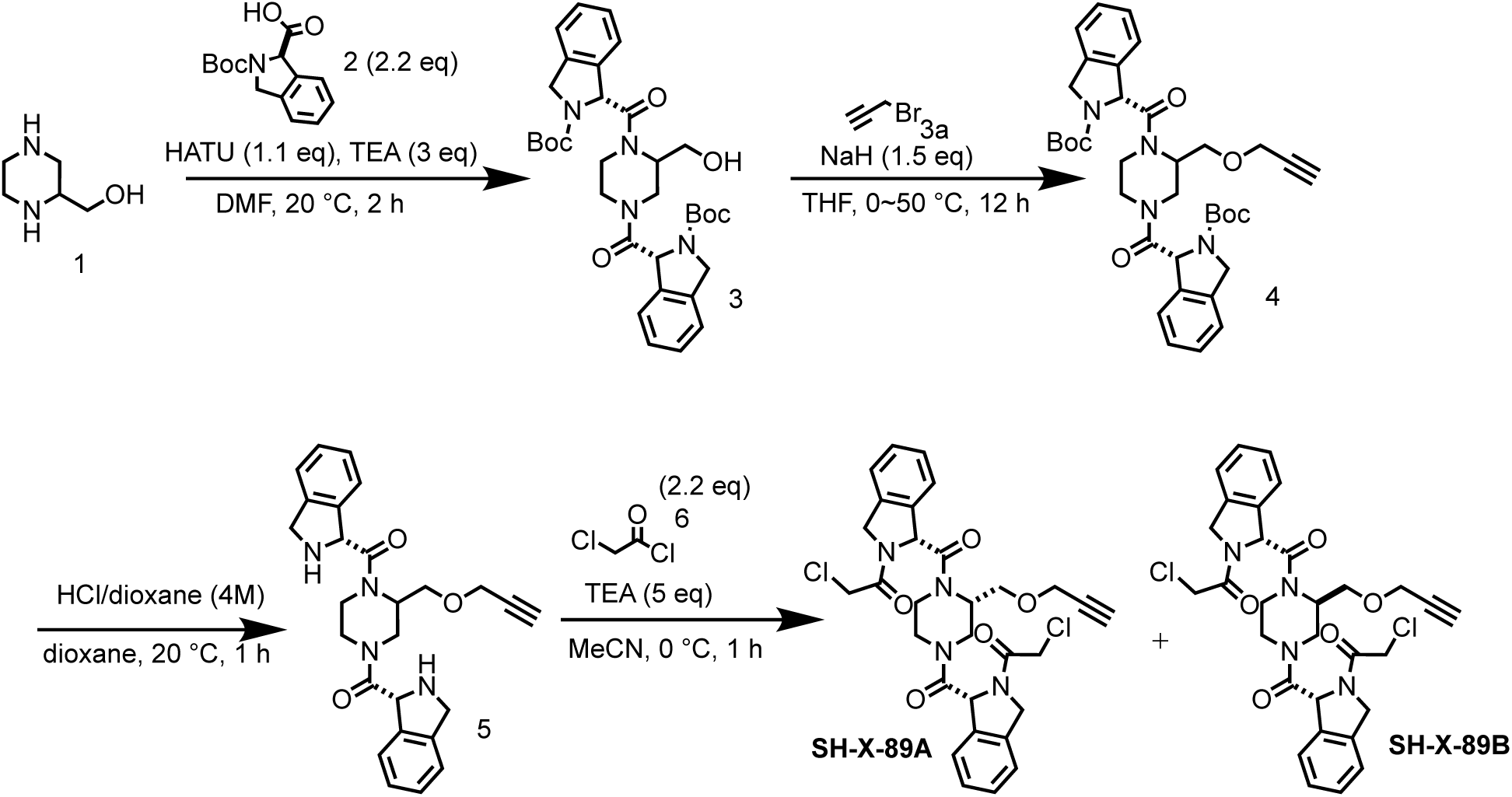

#### Step 1: Preparation of di-*tert*-butyl 1,1’-(2-(hydroxymethyl)piperazine-1,4-dicarbonyl)bis(isoindoline-2-carboxylate) (3)

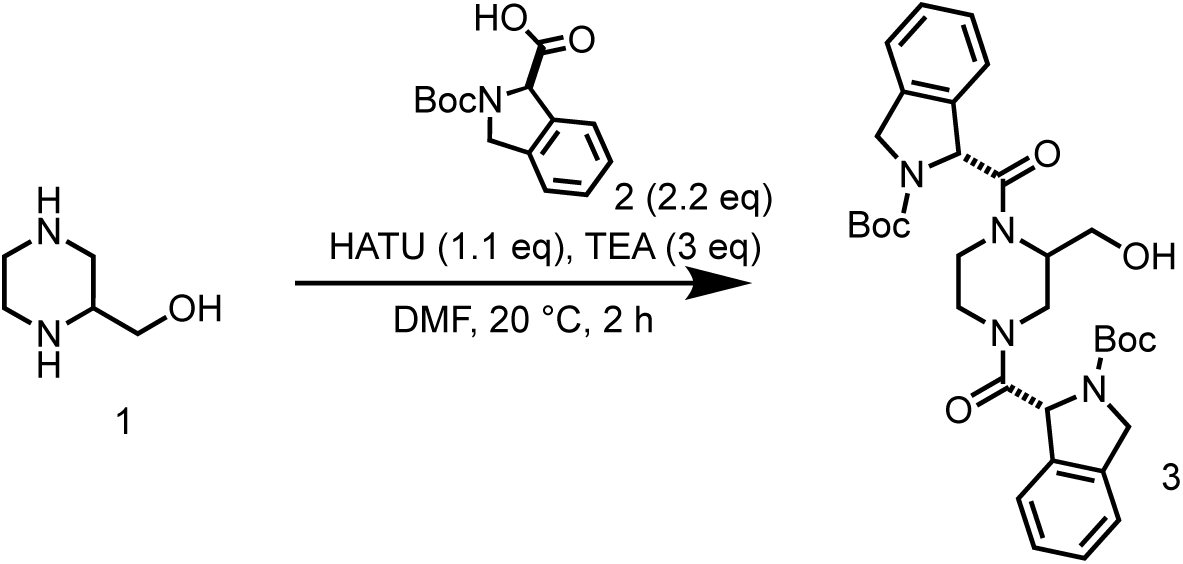

To a solution of piperazin-2-ylmethanol (250 mg, 2.15 mmol) and 2-(*tert*-butoxycarbonyl)isoindoline-1-carboxylic acid (1.19 g, 4.52 mmol) in DCM (10 mL) was added EDCI (1.24 g, 6.46 mmol) and DMAP (52.6 mg, 430 μmol). The mixture was stirred at 20 °C for 1 hr. LCMS showed the reaction was completed. The reaction mixture was concentrated in vacuum to afford a residue. The residue was purified by reversed-phase HPLC (0.1% TFA) to give di-*tert*-butyl 1,1’-(2-(hydroxymethyl) piperazine-1,4-dicarbonyl)bis(isoindoline-2-carboxylate) (700 mg, 1.15 mmol, 53.6% yield) as a yellow solid.

LC-MS: 607.3 [M+H]^+^ / Ret time: 0.426 min / 5-95AB_0.8min.lcm

#### Step 2: Preparation of di-*tert*-butyl 1,1’-(2-((prop-2-yn-1-yloxy)methyl)piperazine-1,4-dicarbonyl)bis(isoindoline-2-carboxylate) (4)

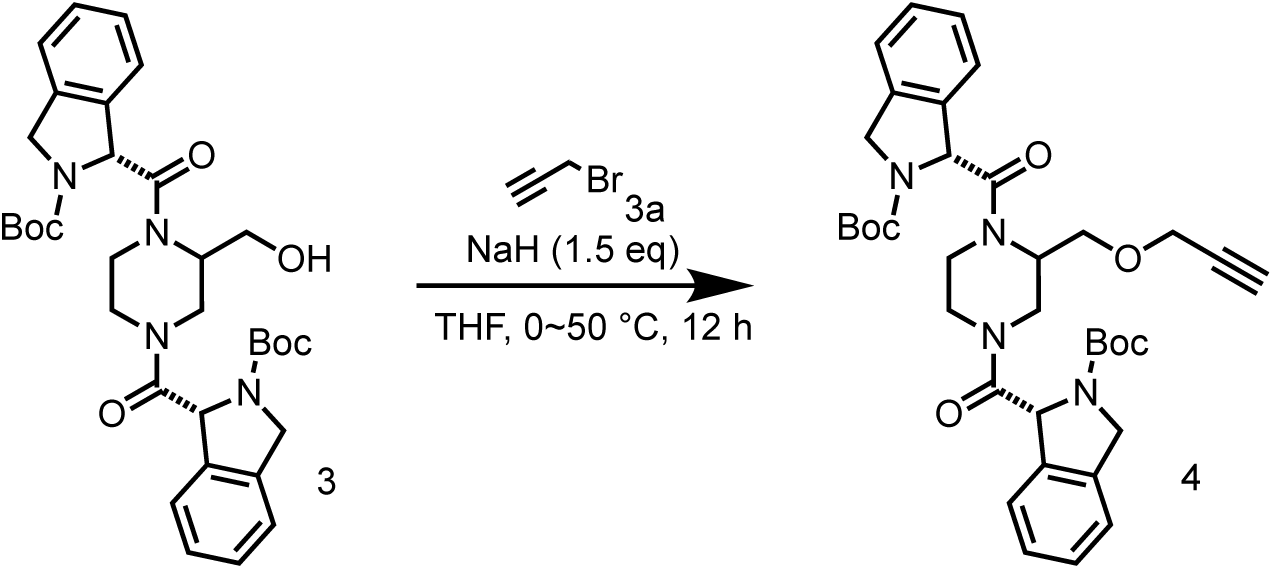

To a solution of di-*tert*-butyl 1,1’-(2-(hydroxymethyl) piperazine-1,4-dicarbonyl) bis(isoindoline-2-carboxylate) (700 mg, 1.15 mmol) in THF (7 mL) was added NaH (69.2 mg, 1.73 mmol, 60% purity) at 0 °C, The mixture was stirred at 0 °C for 0.5 hr, and then 3-bromoprop-1-yne (514 mg, 3.46 mmol, 373 μL) was added at 50 °C. The resulting mixture was stirred at 50 °C for 12 hrs. LCMS showed the reaction was completed. The reaction mixture was added aqueous NH_4_Cl (20 mL) and extracted with EtOAc (20 mL×3), the organic phase was wash with brine (20 mL), dried over with Na_2_SO_4_, then concentrated in vacuum to give a residue. The residue was purified by column chromatography on silica gel (eluted with petroleum ether: ethyl acetate=10:0 to 0:1) to give di-t*ert*-butyl 1,1’-(2-((prop-2-yn-1-yloxy)methyl)piperazine-1,4-dicarbonyl)bis(isoindoline-2-carboxylate) (110 mg, 170 μmol, 14.8% yield) as a yellow solid.

LC-MS: 645.3 [M+H]^+^ / Ret time: 0.533 min / 5-95AB_0.8min.lcm.

#### Step 3: Preparation of (2-((prop-2-yn-1-yloxy)methyl)piperazine-1,4-diyl) bis(isoindolin-1-ylmethanone) (5)

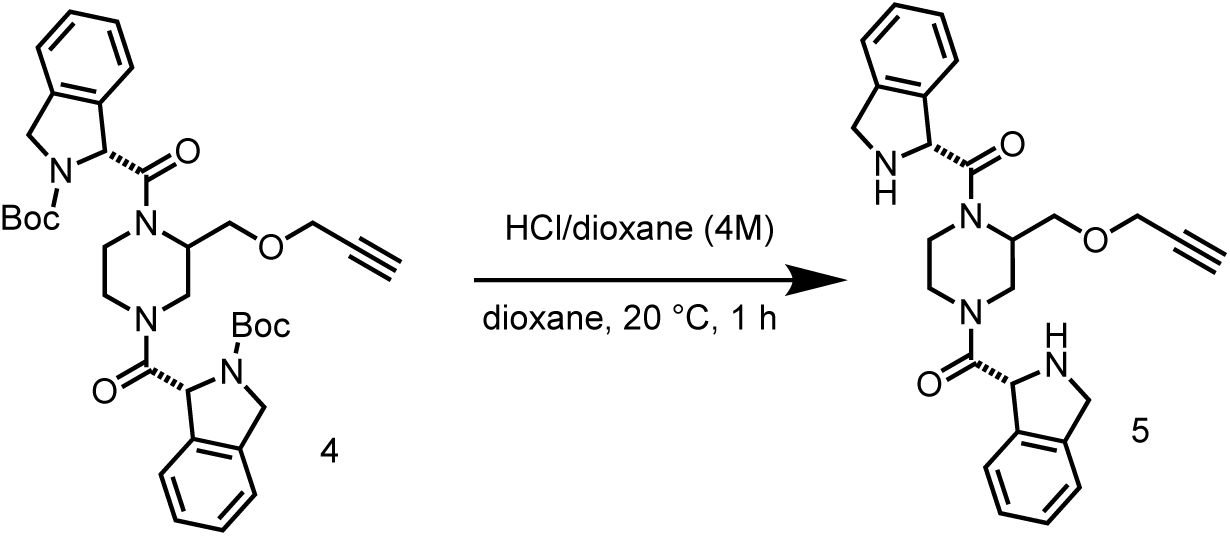

To a solution of di-*tert*-butyl 1,1’-(2-((prop-2-yn-1-yloxy)methyl)piperazine-1,4-dicarbonyl)bis(isoindoline-2-carboxylate) (110 mg, 170 μmol) in DCM (1 mL) was added TFA (1.54 g, 13.5 mmol, 1.00 mL). The mixture was stirred at 20 °C for 1 hr. LCMS showed the reaction was completed. The reaction mixture was concentrated in vacuum to give (2-((prop-2-yn-1-yloxy)methyl)piperazine-1,4-diyl)bis(isoindolin-1-ylmethanone) (70.0 mg, crude, HCl salt) as a yellow solid.

LC-MS: 445.1 [M+H]^+^ / Ret time: 0.250 min / 5-95AB_0.8min.lcm.

#### Step 4: Synthesis of 1,1’-(((*S*)-2-((prop-2-yn-1-yloxy)methyl)piperazine-1,4-dicarbonyl)bis(isoindoline-1,2-diyl))bis(2-chloroethan-1-one) (SH-X-89A) and 1,1’-(((*R*)-2-((prop-2-yn-1-yloxy)methyl)piperazine-1,4-dicarbonyl)bis(isoindoline-1,2-diyl))bis(2-chloroethan-1-one) (SH-X-89B)

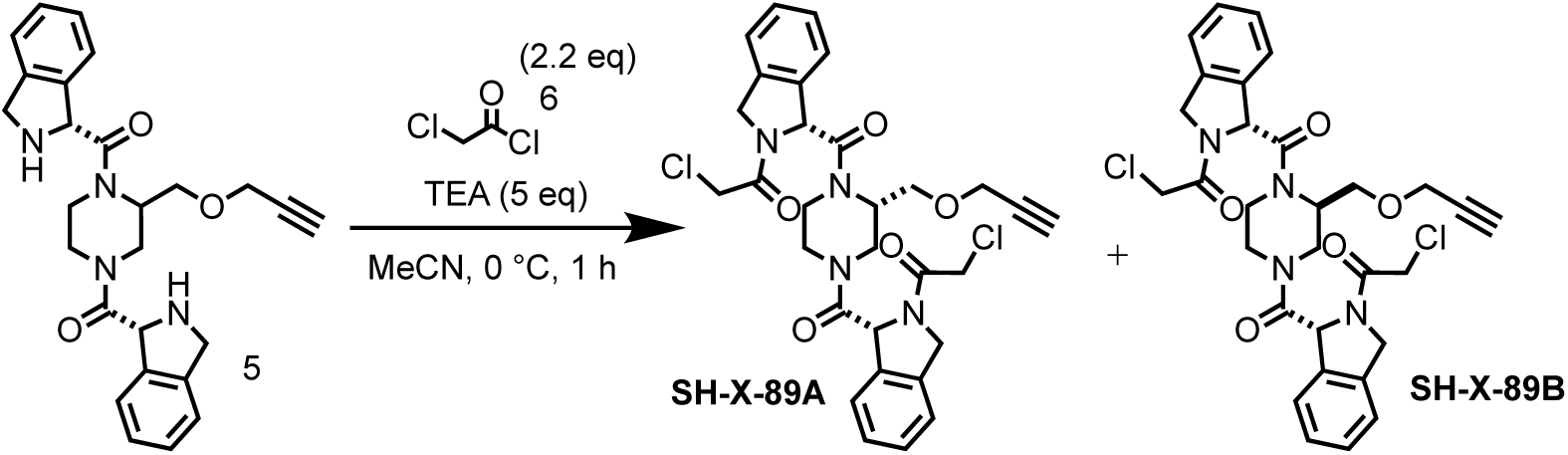

To a solution of (2-((prop-2-yn-1-yloxy)methyl)piperazine-1,4-diyl)bis(isoindolin-1-ylmethanone) (70.0 mg, 157 μmol) in MeCN (1 mL) was added TEA (79.7 mg, 787 μmol, 109 μL) and 2-chloroacetyl chloride (39.1 mg, 346 μmol, 27.6 μL). The mixture was stirred at 0 °C for 1 hr. LCMS showed the reaction was completed. The reaction mixture was filtered and the filtrate was concentrated to give a residue. The residue was purified by preparative-HPLC (column: Welch Ultimate C18 150*25mm*5um; mobile phase: [water(FA)-MeCN]; gradient:30%-60% B over 10 min) to give 1,1’-(((*S*)-2-((prop-2-yn-1-yloxy)methyl)piperazine-1,4-dicarbonyl)bis(isoindoline-1,2-diyl))bis(2-chloroethan-1-one) (3.40 mg, 4.81 μmol, 3.06% yield, 84.6% purity) as an off-white solid and 1,1’-(((*R*)-2-((prop-2-yn-1-yloxy)methyl)piperazine-1,4-dicarbonyl)bis(isoindoline-1,2-diyl))bis(2-chloroethan-1-one) (12.8 mg, 19.8 μmol, 12.6% yield, 92.4% purity) as a white solid.

LC-MS: 597.1 [M+H]^+^ / Ret time: 0.419 min / 5-95AB_0.8min.lcm.

SH-X-89A: ^1^H NMR (400 MHz, DMSO-*d*_6_): *δ* 7.44-7.31 (m, 7H), 7.26-7.15 (m, 1H), 6.26-5.85 (m, 1H), 5.70-5.56 (m, 1H), 5.01-4.91 (m, 4H), 4.58-4.48 (m, 4H), 4.47-4.35 (m, 2H), 4.33-4.21 (m, 2H), 3.60-3.45 (m, 3H), 3.27-3.20 (m, 1H), 3.03-2.81 (m, 2H), 2.79-2.65 (m, 2H). LC-MS: 597.1 [M+H]^+^ / Ret time: 0.416 min / 5-95AB_0.8min.lcm.

SH-X-89B: ^1^H NMR (400 MHz, MeOD-*d*_4_): *δ* 7.55-7.46 (m, 1H), 7.39 (d, *J* = 4.0 Hz, 4H), 7.36-7.30 (m, 2H), 7.26-7.18 (m, 1H), 6.31-5.98 (m, 1H), 5.76-5.66 (m, 1H), 5.08-4.98 (m, 4H), 4.48-4.38 (m, 4H), 4.38-4.29 (m, 3H), 4.27 (d, *J* = 6.0 Hz, 1H), 3.76-3.57 (m, 2H), 3.50-3.42 (m, 1H), 3.17-2.98 (m, 1H), 2.95-2.82 (m, 2H), 2.76-2.62 (m, 2H).

### Synthesis of 3,5-diacrylamidobenzoic acid (5) and 3-acetamido-5-acrylamidobenzoic acid (8)

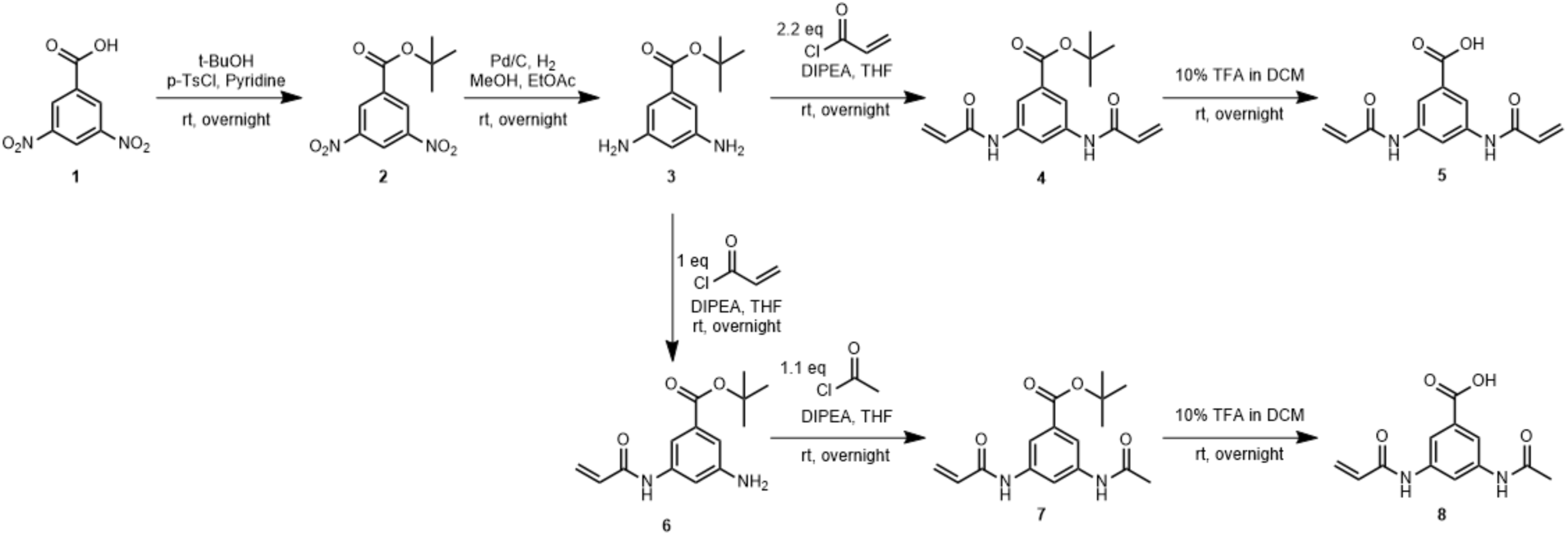

#### Step 1: Preparation of *tert*-butyl 3,5-dinitrobenzoate (2)

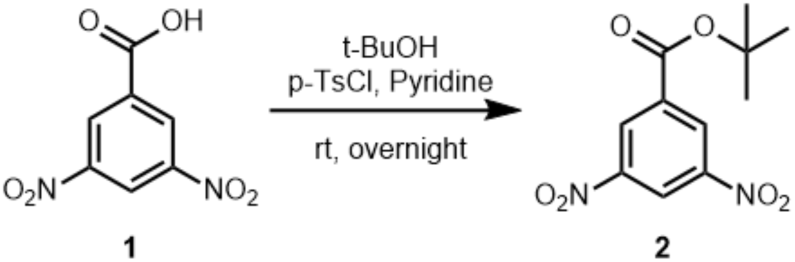

To a solution of 3,5-dinitrobenzoic acid (**1**) (2.0g, 0.009 mol) in pyridine (38.5mL), *p-* toluenesulphonyl chloride (8.1g, 0.042 mol) and 15 minutes later *tert-*butanol (9.7mL, 0.102 mol) were added. The reaction mixture was stirred at room temperature overnight until completion and then poured onto ice. The white solid precipitate was filtered and washed with water. The crude product was analyzed by NMR and carried to the next step. Yield: 2.2g (87%).

^1^H NMR (400 MHz, MeOD) *δ* 9.17 (dd, *J* = 2.1, 2.1 Hz, 1H), 9.02 (d, *J* = 2.1 Hz, 2H), 1.66 (s, 9H).

#### Step 2: Preparation of *tert*-butyl 3,5-diaminobenzoate (3)

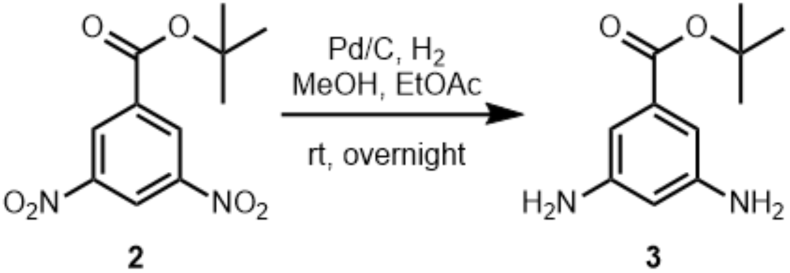

Under nitrogen, **2** (2.2g, 0.008 mol) was dissolved in a 1:1 mixture of methanol (40mL) and ethyl acetate (40mL). 10% Pd/C (174mg, 1.64 mmol) was slowly added, and a balloon of H_2_ gas was then inserted. The reaction mixture was stirred overnight until completion by chromatography. The mixture was filtered through celite, and the solvent was removed under reduced pressure, leaving pure product by NMR. Yield: 1.6g (93%).

^1^H NMR (400 MHz, MeOD) *δ* 6.66 (d, *J* = 2.1 Hz, 2H), 6.26 (dd, *J* = 2.1, 2.1 Hz, 1H), 1.52 (s, 9H).

#### Step 3a: Preparation of tert-butyl 3,5-diacrylamidobenzoate (4)

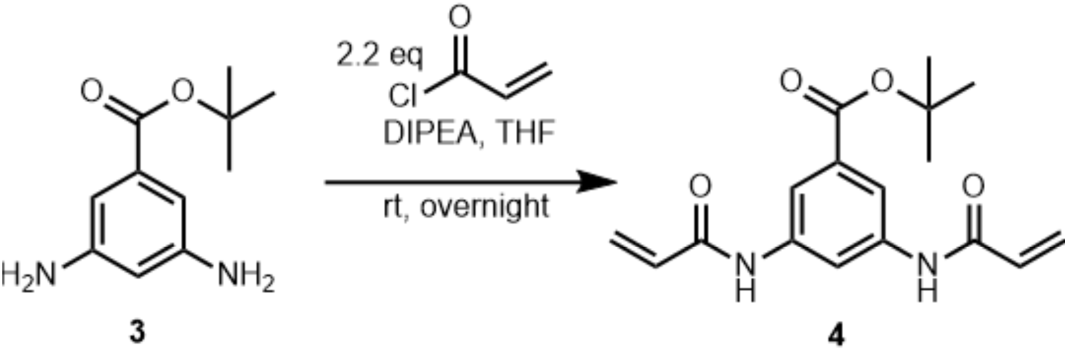

To a solution of **3** (2.0g, 0.010 mol) in THF, diisopropylethylamine (3.7mL, 0.021 mol) and acryloyl chloride (1.7mL, 0.021 mol) were added. The reaction was stirred overnight to completion and the solvent removed under reduced pressure. The crude product was purified under column chromatography (eluent: DCM/MeOH). Yield: 1.7g (54%).

^1^H NMR (400 MHz, CDCl_3_) *δ* 6.52 – 6.39 (m, 2H), 6.27 (dd, *J* = 16.9, 10.2 Hz, 2H), 5.83 (dd, *J* = 9.6, 9.6 Hz, 2H), 1.59 (s, 9H).

#### Step 4a: Preparation of 3,5-diacrylamidobenzoic acid (5)

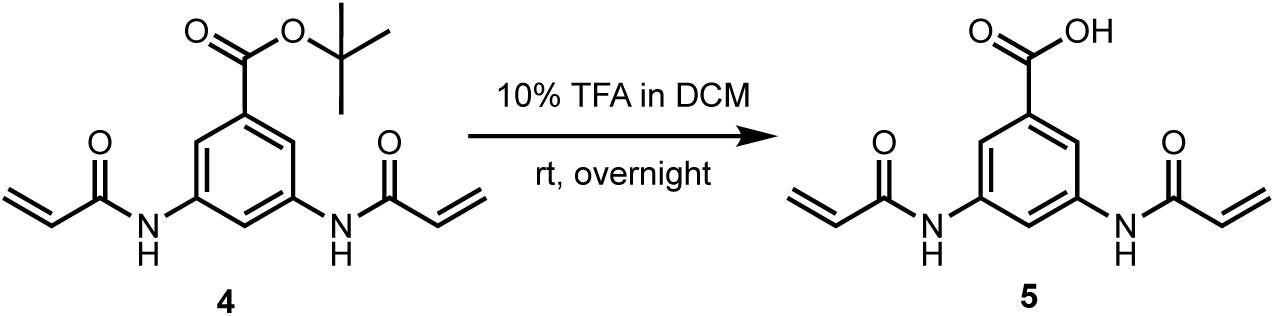

**4** (500mg, 1.6mmol) was dissolved in a solution of 10% trifluoroacetic acid in dichloromethane (8mL) and allowed to react overnight until reaction completion. The solvent was removed under reduced pressure. Yield: 410.3mg (99%)

^1^H NMR (400 MHz, MeOD) *δ* 8.33 (dd, *J* = 1.9, 1.8 Hz, 1H), 8.07 (d, *J* = 2.1 Hz, 2H), 6.48 – 6.37 (m, 4H), 5.79 (dd, *J* = 9.5, 2.4 Hz, 2H).

#### Step 3b: Preparation of tert-butyl 3-acrylamido-5-aminobenzoate (6)

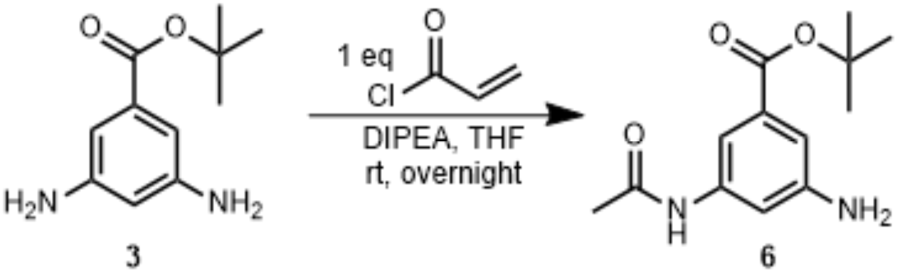

Acryloyl chloride (1.7mL, 0.021mol) was slowly added to a solution of **3** (4.4g, 0.021 mol) and diisopropylethylamine (7.4mL, 0.042mol) in THF (105mL) at −78℃. The mixture was stirred at - 78℃ for 10m, then allowed to warm to room temperature and stirred overnight. The solvent was removed under reduced pressure and the residual mixture purified by C18 column chromatography (eluent: H_2_O/ACN with 0.1% formic acid). Yield: 2.2g (40%).

^1^H NMR (400 MHz, MeOD) *δ* 7.42 (dd, *J* = 1.7, 1.7 Hz, 1H), 7.36 (dd, *J* = 2.1, 2.1 Hz, 1H), 7.06 (dd, *J* = 2.2, 1.5 Hz, 1H), 6.40 (d, *J* = 9.6 Hz, 1H), 6.36 (d, *J* = 2.3 Hz, 1H), 5.75 (dd, *J* = 9.6, 2.3 Hz, 1H), 1.58 (s, 9H).

#### Step 4b: Preparation of tert-butyl 3-acetamido-5-acrylamidobenzoate (7)

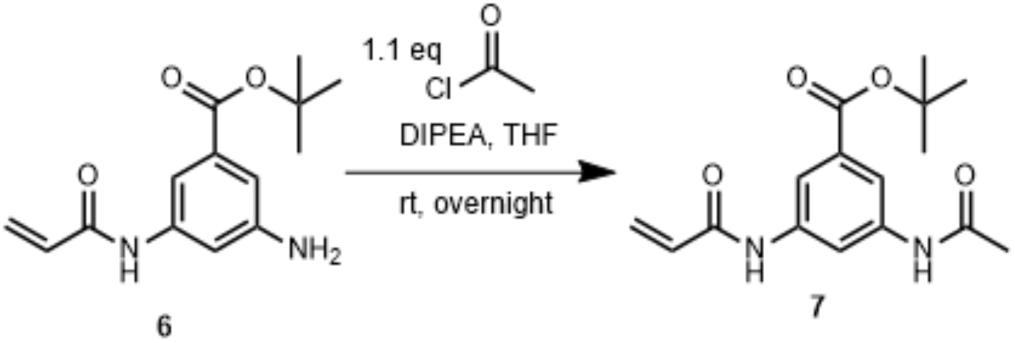

Acetyl chloride (0.12mL, 1.7 mmol) was added to a solution of **6** (227mg, 0.8mmol) in THF (9mL) and stirred overnight at room temperature. The solvent was removed under reduced pressure and the crude material purified by column chromatography (eluent: Hexanes/Ethyl Acetate). Yield: 103mg (39%).

^1^H NMR (400 MHz, MeOD) *δ* 8.24 (dd, *J* = 2.1, 2.1 Hz, 1H), 7.98 (dd, *J* = 1.8, 1.8 Hz, 1H), 7.90 (dd, *J* = 1.7, 1.7 Hz, 1H), 6.42 (d, *J* = 9.4 Hz, 1H), 6.39 (d, *J* = 2.4 Hz, 1H), 5.78 (dd, *J* = 9.4, 2.5 Hz, 1H), 2.14 (s, 3H), 1.60 (s, 9H).

#### Step 5: Preparation of 3-acetamido-5-acrylamidobenzoic acid (8)

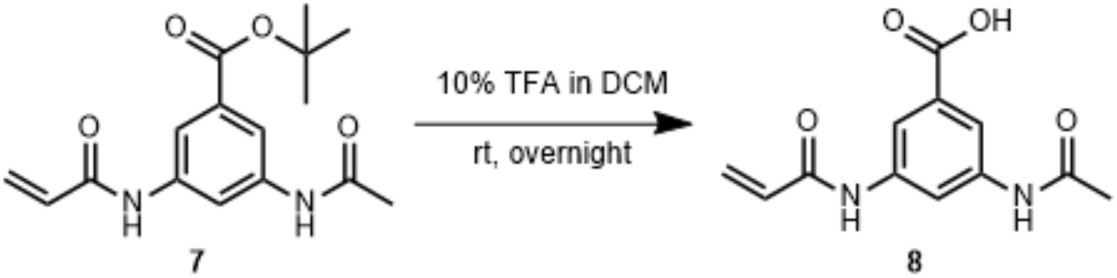

**7** (500mg, 1.6mmol) was dissolved in a solution of 10% trifluoroacetic acid in dichloromethane (8mL) and reacted overnight until the reaction was complete. The solvent was removed under reduced pressure. Yield: 357mg (88%)

^1^H NMR (400 MHz, MeOD) *δ* 8.24 (dd, *J* = 2.1, 2.1 Hz, 1H), 8.05 (dd, *J* = 1.8, 1.8 Hz, 1H), 7.97 (dd, *J* = 1.8, 1.8 Hz, 1H), 6.45 – 6.37 (m, 2H), 5.79 (dd, *J* = 9.4, 2.5 Hz, 1H), 2.14 (s, 3H).

### General procedure for amide couplings for mono-warhead and dual-warhead compounds

To a solution of **5** (0.040mmol) or **8** (0.040mmol) in DMF (0.4mL), COMU (20.7mg, 0.048mmol) and diisopropylethylamine (0.018mL, 0.101mmol) were added and stirred for 10 minutes at room temperature. The amine substrate (0.040mmol) was then added and the reaction run overnight until completion. The reaction mixture was then diluted with ethyl acetate and washed with water twice. The organic layer was extracted and washed twice more with 1M HCl, twice with 2M aqueous NaOH, and once with brine. The solvent was removed under reduced pressure, and the crude product was subject to NMR analysis.

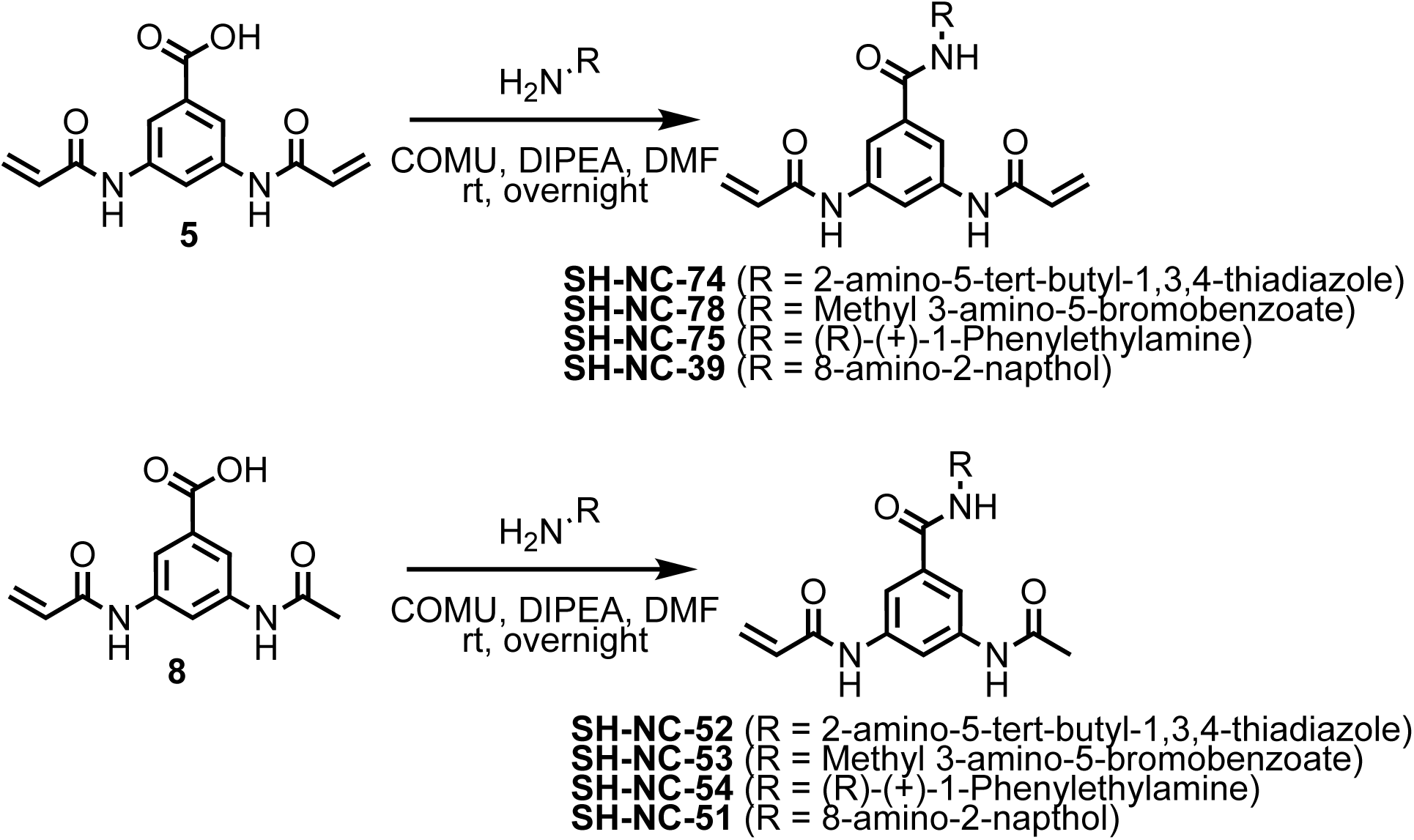

#### SH-NC-74 (*N*,*N*’-(5-((5-(*tert*-butyl)-1,3,4-thiadiazol-2-yl)carbamoyl)-1,3-phenylene)diacrylamide)

^1^H NMR (400 MHz, MeOD) *δ* 8.25 (dd, *J* = 2.0, 2.0 Hz, 1H), 8.02 (d, *J* = 1.9 Hz, 2H), 6.47 – 6.42 (m, 4H), 5.82 (dd, *J* = 9.0, 2.9 Hz, 2H), 1.50 (s, 9H).

#### SH-NC-52 (3-acetamido-5-acrylamido-*N*-(5-(*tert*-butyl)-1,3,4-thiadiazol-2-yl)benzamide)

^1^H NMR (400 MHz, MeOD) *δ* 8.16 (dd, *J* = 2.0, 2.0 Hz, 1H), 8.00 (dd, *J* = 1.8, 1.8 Hz, 1H), 7.92 (dd, *J* = 1.8, 1.8 Hz, 1H), 6.52 – 6.36 (m, 2H), 5.81 (dd, *J* = 8.9, 3.0 Hz, 1H), 2.17 (s, 3H), 1.50 (s, 9H).

#### SH-NC-78 (methyl 3-bromo-5-(3,5-diacrylamidobenzamido)benzoate)

^1^H NMR (400 MHz, MeOD) *δ* 8.34 (dd, *J* = 2.0, 1.4 Hz, 1H), 8.30 (dd, *J* = 2.0, 2.0 Hz, 1H), 8.21 (dd, *J* = 2.0, 2.0 Hz, 1H), 7.94 (d, *J* = 2.0 Hz, 2H), 7.91 (dd, *J* = 1.7, 1.7 Hz, 1H), 6.50 – 6.41 (m, 4H), 5.82 (dd, *J* = 9.1, 2.8 Hz, 2H), 3.94 (s, 3H).

#### SH-NC-53 (methyl 3-(3-acetamido-5-acrylamidobenzamido)-5-bromobenzoate)

^1^H NMR (400 MHz, MeOD) *δ* 8.31 (dd, *J* = 2.0, 1.5 Hz, 1H), 8.28 (dd, *J* = 2.0, 2.0 Hz, 1H), 8.11 (dd, *J* = 2.0, 2.0 Hz, 1H), 7.92 (dd, *J* = 1.8, 1.8 Hz, 1H), 7.90 (dd, *J* = 1.7, 1.7 Hz, 1H), 7.83 (dd, *J* = 1.8, 1.8 Hz, 1H), 6.52 – 6.35 (m, 2H), 5.81 (dd, *J* = 9.0, 2.9 Hz, 1H).

#### SH-NC-75 ((*R*)-*N*,*N*’-(5-((1-phenylethyl)carbamoyl)-1,3-phenylene)diacrylamide)

^1^H NMR (400 MHz, DMSO) *δ* 12.47 (s, 2H), 7.40 – 7.14 (m, 6H), 6.25 (dd, *J* = 17.2, 1.7 Hz, 2H), 6.08 (dd, *J* = 17.3, 10.3 Hz, 2H), 5.87 (dd, *J* = 10.2, 1.6 Hz, 2H), 1.65 (s, 3H + H_2_O).

#### SH-NC-54 ((*R*)-3-acetamido-5-acrylamido-*N*-(1-phenylethyl)benzamide)

^1^H NMR (400 MHz, MeOD) *δ* 8.08 (dd, *J* = 2.0, 2.0 Hz, 1H), 7.78 (dd, *J* = 1.7, 1.7 Hz, 1H), 7.69 (dd, *J* = 1.8, 1.8 Hz, 1H), 7.43 – 7.37 (m, 2H), 7.33 (dd, *J* = 8.5, 6.8 Hz, 2H), 7.28 – 7.19 (m, 1H), 6.50 – 6.33 (m, 2H), 5.79 (dd, *J* = 9.1, 2.8 Hz, 1H), 5.21 (q, *J* = 6.7 Hz, 1H), 2.14 (s, 3H), 1.56 (d, *J* = 7.0 Hz, 3H).

#### SH-NC-39 (*N*,*N*’-(5-((7-hydroxynaphthalen-1-yl)carbamoyl)-1,3-phenylene)diacrylamide)

^1^H NMR (400 MHz, MeOD) *δ* 8.24 (s, 1H), 8.09 (s, 2H), 7.79 (d, *J* = 8.9 Hz, 1H), 7.74 (d, *J* = 8.2 Hz, 1H), 7.51 (d, *J* = 7.2 Hz, 1H), 7.31 (t, *J* = 7.8 Hz, 1H), 7.28 (d, *J* = 2.2 Hz, 1H), 7.11 (dd, *J* = 8.8, 2.4 Hz, 1H), 6.53 – 6.38 (m, 4H), 5.82 (dd, *J* = 9.3, 2.6 Hz, 2H).

#### SH-NC-51 (3-acetamido-5-acrylamido-*N*-(7-hydroxynaphthalen-1-yl)benzamide)

^1^H NMR (400 MHz, MeOD) *δ* 8.15 (dd, *J* = 2.0, 2.0 Hz, 1H), 8.08 (dd, *J* = 1.9 Hz, 1H), 7.98 (dd, *J* = 1.8, 1.8 Hz, 1H), 7.79 (d, *J* = 8.9 Hz, 1H), 7.74 (d, *J* = 8.2 Hz, 1H), 7.51 (d, *J* = 7.2 Hz, 1H), 7.31 (dd, *J* = 8.2, 7.3 Hz, 1H), 7.27 (d, *J* = 2.4 Hz, 1H), 7.11 (dd, *J* = 8.9, 2.4 Hz, 1H), 6.53 – 6.37 (m, 2H), 5.81 (dd, *J* = 9.3, 2.5 Hz, 1H), 2.17 (s, 3H).

### Synthesis of SH-E4A-59 and SH-E4A-66 and SH-E4A-65

#### Step 1: Preparation of 3-acetamido-5-acrylamidobenzoic acid (10)

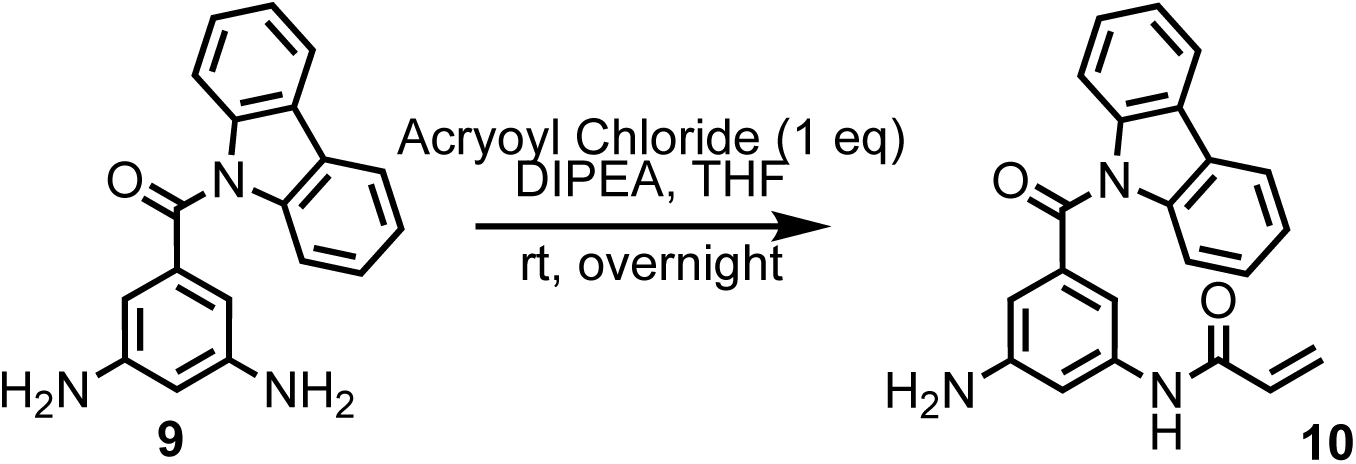

To a solution of **9** (50mg, 0.163mmol) dissolved in THF (0.82mL), diisopropylethylamine (0.057mL, 0.327mmol) and acryloyl chloride (0.013mL, 0.163mmol) were slowly added at −78℃. The reaction was stirred for 10 minutes, then allowed to warm to room temperature overnight. Upon completion, the solvent was removed under reduced pressure and the crude material purified by column chromatography (eluent: H_2_O/ACN with 0.1% formic acid). Yield: 17mg (25%).

^1^H NMR (400 MHz, CDCl_3_) *δ* 8.04 – 7.97 (m, 2H), 7.77 (s, 1H), 7.64 – 7.56 (m, 2H), 7.40 – 7.31 (m, 4H), 6.78 (dd, *J* = 25.0, 1.8 Hz, 2H), 6.46 – 6.38 (m, 1H), 6.18 (dd, *J* = 16.8, 10.3 Hz, 1H), 5.78 (d, *J* = 10.3 Hz, 1H), 3.94 (s, 2H).

#### Step 2a: Preparation of *N*-(3-(9*H*-carbazole-9-carbonyl)-5-(2-chloroacetamido)phenyl)acrylamide (SH-E4A-66)

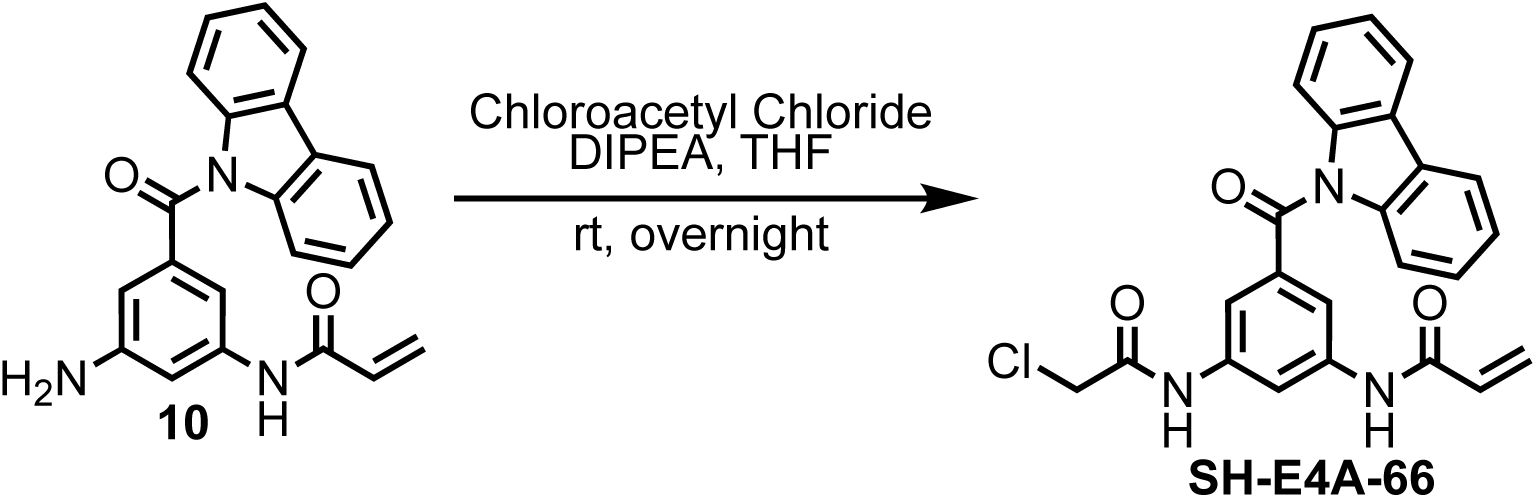

Chloroacetyl chloride (0.0023mL, 0.029mmol) was added to a solution of **10** (5.2mg, 0.015mmol) dissolved in THF (0.146mL) and diisopropylethylamine (0.005mL, 0.029mmol) and the reaction stirred overnight at room temperature. The solvent was removed under reduced pressure and the crude material purified by column chromatography (eluent: DCM/MeOH). Yield: 5.5mg (87%).

^1^H NMR (400 MHz, CDCl_3_) *δ* 8.39 (d, *J* = 1.9 Hz, 1H), 8.36 (s, 1H), 8.00 (dd, *J* = 6.5, 0.9 Hz, 2H), 7.70 (dd, *J* = 2.8, 1.9 Hz, 1H), 7.59 (dd, *J* = 1.7, 1.7 Hz, 1H), 7.55 (dd, *J* = 7.1, 1.8 Hz, 2H), 7.44 – 7.29 (m, 4H), 6.44 (dd, *J* = 16.9, 1.1 Hz, 1H), 6.21 (dd, *J* = 16.8, 10.3 Hz, 1H), 5.82 (dd, *J* = 10.1, 1.6 Hz, 1H), 4.18 (s, 2H).

#### Step 2b: Preparation of *N*-(3-acetamido-5-(9*H*-carbazole-9-carbonyl)phenyl)acrylamide (SH-E4A-65)

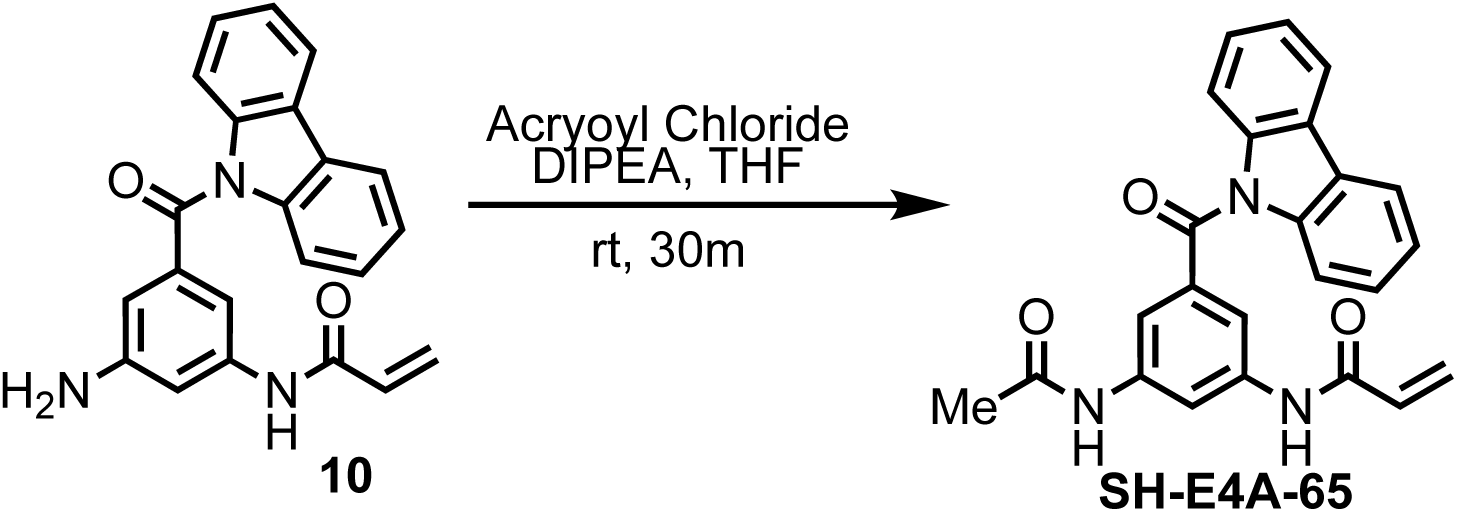

Acetyl chloride (0.0053mL, 0.075mmol) was added to a solution of **10** (13.3mg, 0.037mmol) dissolved in THF (0.2mL) and diisopropylethylamine (0.016mL, 0.094mmol) and the reaction stirred overnight at room temperature. The solvent was removed under reduced pressure and the crude material purified by column chromatography (eluent: DCM/MeOH). Yield: 9.3mg (63%).

^1^H NMR (400 MHz, CDCl_3_) *δ* 8.33 (s, 1H), 8.00 (dd, *J* = 5.7, 2.1 Hz, 2H), 7.64 (s, 1H), 7.59 – 7.52 (m, 3H), 7.40 – 7.29 (m, 5H), 6.42 (dd, *J* = 16.7, 0.9 Hz, 1H), 6.21 (dd, *J* = 16.8, 10.3 Hz, 1H), 5.80 (dd, *J* = 10.1, 1.0 Hz, 1H), 2.18 (s, 3H).

### Preparation of N,N’-(5-(9H-carbazole-9-carbonyl)-1,3-phenylene)diacrylamide (SH-E4A-59)

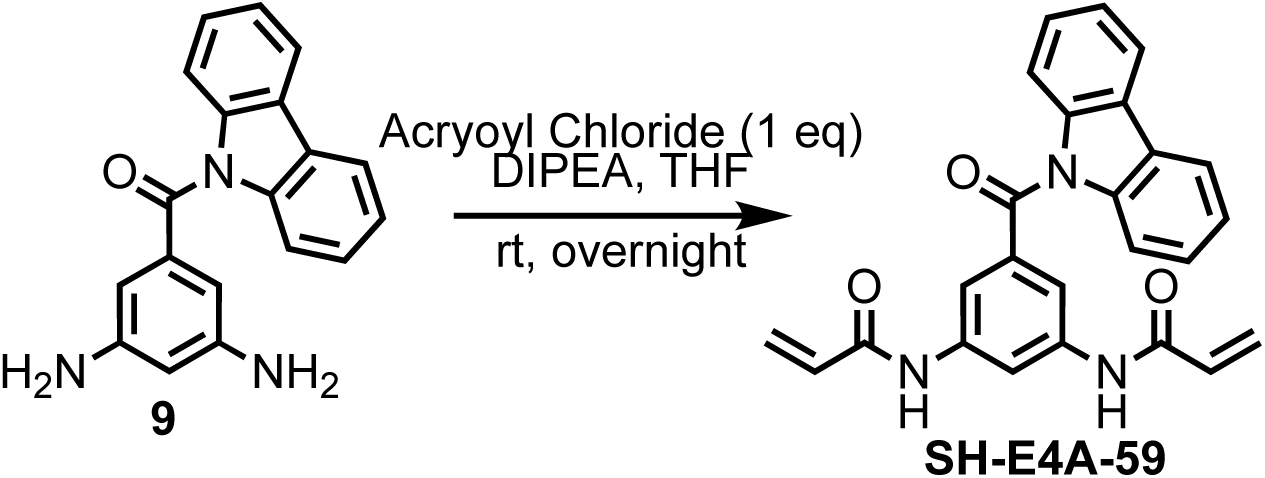

To a solution of **9** (50mg, 0.163mmol) dissolved in THF (0.82mL), diisopropylethylamine (0.057mL, 0.327mmol) and acryloyl chloride (0.026mL, 0.326mmol) were slowly added at −78℃. The reaction was stirred for 10 minutes, then allowed to warm to room temperature overnight. Upon completion, the solvent was removed under reduced pressure and the crude material purified by column chromatography (eluent: H_2_O/ACN with 0.1% formic acid). Yield: 30mg (50%).

^1^H NMR (400 MHz, MeOD) δ 8.48 (t, *J* = 2.0 Hz, 1H), 8.10-8.08 (m, 2H), 7.74-7.73 (m, 2H), 7.55-7.53 (m, 2H), 7.39-7.34 (m, 4H), 6.45-6.33 (m, 4H), 5.77 (dd, *J* = 9.3, 2.6 Hz, 2H).

### Synthesis of *N*-(3-(2-chloroacetamido)phenyl)acrylamide (SH-NC-27)

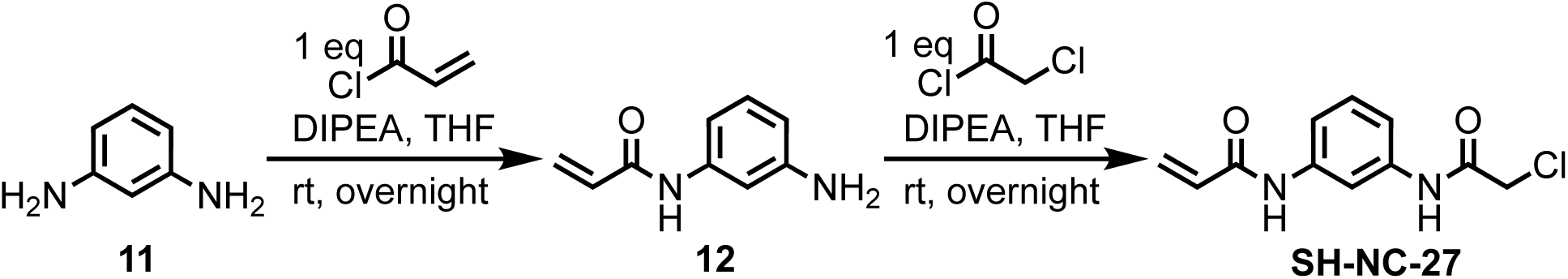

#### Step 1: Preparation of *N*-(3-aminophenyl)acrylamide (12)

Acryloyl chloride (0.075mL, 0.925mmol) was dripped to a solution of **11** (100mg, 0.925mmol) and diisopropylethylamine (0.322mL, 1.850mmol) in THF (4.6mL) at −78℃. The mixture was stirred at −78℃ for 10m, then allowed to warm to room temperature and stirred overnight. The solvent was removed under reduced pressure and the residual mixture purified by C18 column chromatography (eluent: H_2_O/ACN with 0.1% formic acid). Yield: 22mg (15%).

^1^H NMR (400 MHz, DMSO) *δ* 10.23 (s, 1H), 8.13 (s, 2H), 7.51 (s, 1H), 7.20 (d, *J* = 4.7 Hz, 2H), 6.71 (dd, *J* = 5.5, 5.4 Hz, 1H), 6.49 – 6.42 (m, 1H), 6.25 (dd, *J* = 17.0, 2.0 Hz, 1H), 5.75 (dd, *J* = 10.1, 2.0 Hz, 1H).

#### Step 2: Preparation of *N*-(3-(2-chloroacetamido)phenyl)acrylamide (SH-NC-27)

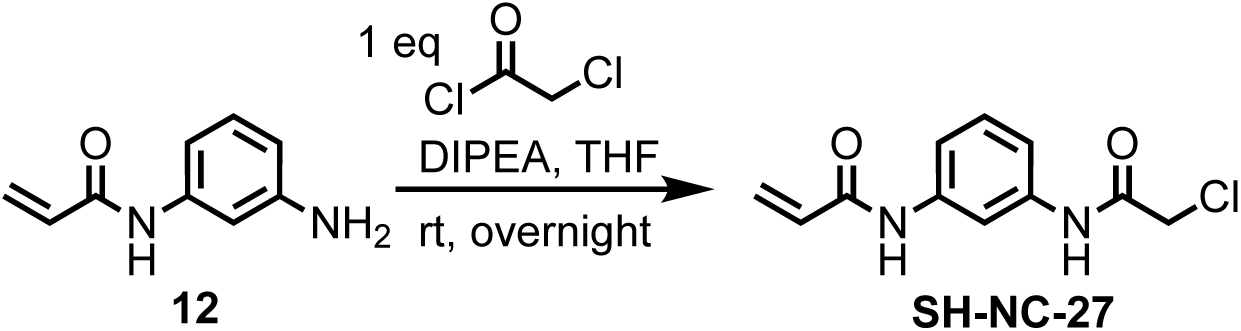

Chloroacetyl chloride (0.022mL, 0.271mmol) was added to a solution of **12** (22mg, 0.136mmol) in THF (1.36mL) and diisopropylethylamine (0.047mL, 0.271mmol). The reaction was stirred overnight until completion and the solvent removed under reduced pressure. The crude product was purified by column chromatography (eluent: Hexanes/Ethyl Acetate). Yield: 16mg (51%)

^1^H NMR (400 MHz, MeOD) *δ* 7.47 – 7.23 (m, 5H), 6.50 – 6.31 (m, 2H), 5.77 (dd, *J* = 9.7, 2.2 Hz, 1H), 4.15 (s, 2H).

### Synthesis of *N*-(3-acetamido-5-(9*H*-carbazole-9-carbonyl)phenyl)-2-chloroacetamide (SH-E4A-62)

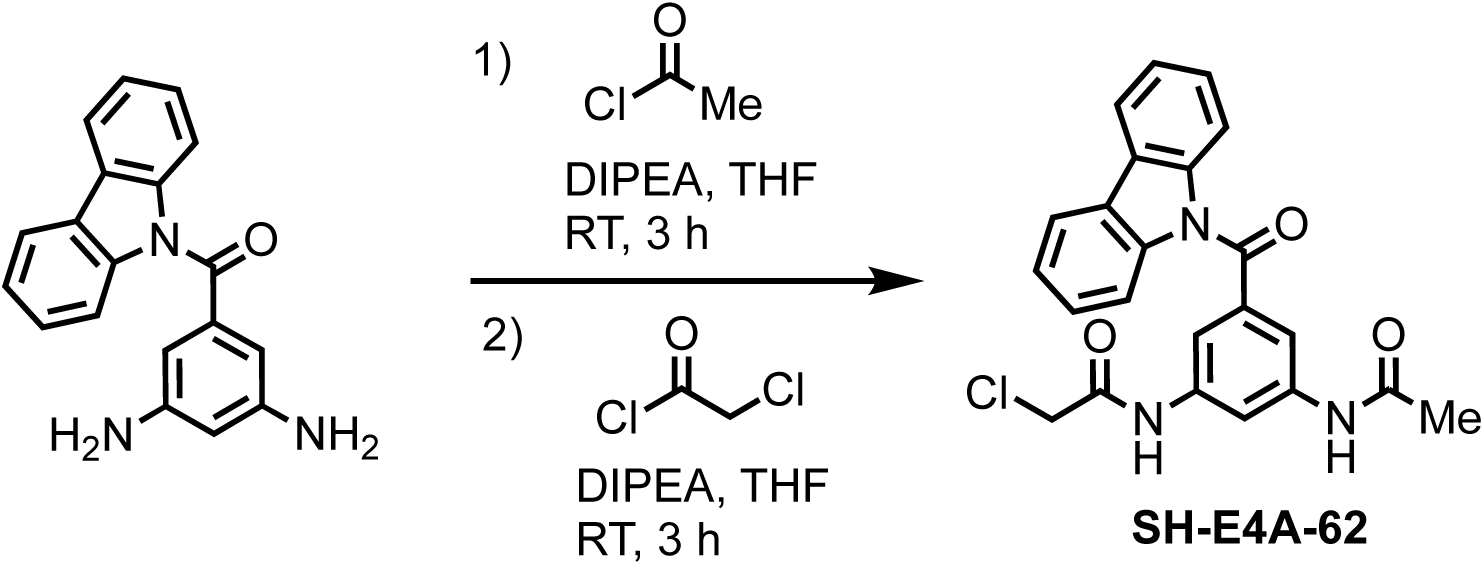

#### Step 1: Preparation of *N*,*N*’-(5-(9*H*-carbazole-9-carbonyl)-1,3-phenylene)diacetamide

Acetyl chloride (0.0053mL, 0.075mmol) was added to a solution of (9*H*-carbazol-9-yl)(3,5-diaminophenyl)methanone (11.7mg, 0.037mmol) dissolved in THF (0.2mL) and diisopropylethylamine (0.016mL, 0.094mmol) and the reaction stirred overnight at room temperature. The solvent was removed under reduced pressure and the crude material purified by column chromatography (eluent: DCM/MeOH). (41%).

#### Step 2: Preparation of *N*-(3-acetamido-5-(9*H*-carbazole-9-carbonyl)phenyl)-2-chloroacetamide (SH-E4A-62)

Chloroacetyl chloride (0.0023mL, 0.029mmol) was added to a solution of (5.1mg, 0.015mmol) dissolved in THF (0.146mL) and diisopropylethylamine (0.005mL, 0.029mmol) and the reaction stirred overnight at room temperature. The solvent was removed under reduced pressure and the crude material purified by column chromatography (eluent: DCM/MeOH). Yield: 5.5mg (87%).

^1^H NMR (400 MHz, MeOD) *δ* 8.28 (dd, *J* = 2.0, 2.0 Hz, 1H), 8.07 – 7.98 (m, 2H), 7.60 (dt, *J* = 4.0, 1.6 Hz, 2H), 7.51 – 7.40 (m, 2H), 7.36 – 7.25 (m, 4H), 4.12 (s, 2H), 2.07 (s, 3H).

### Synthesis of *N*,*N*’-(5-(9*H*-carbazole-9-carbonyl)-1,3-phenylene)bis(2-chloroacetamide) (SH-E4A-61)

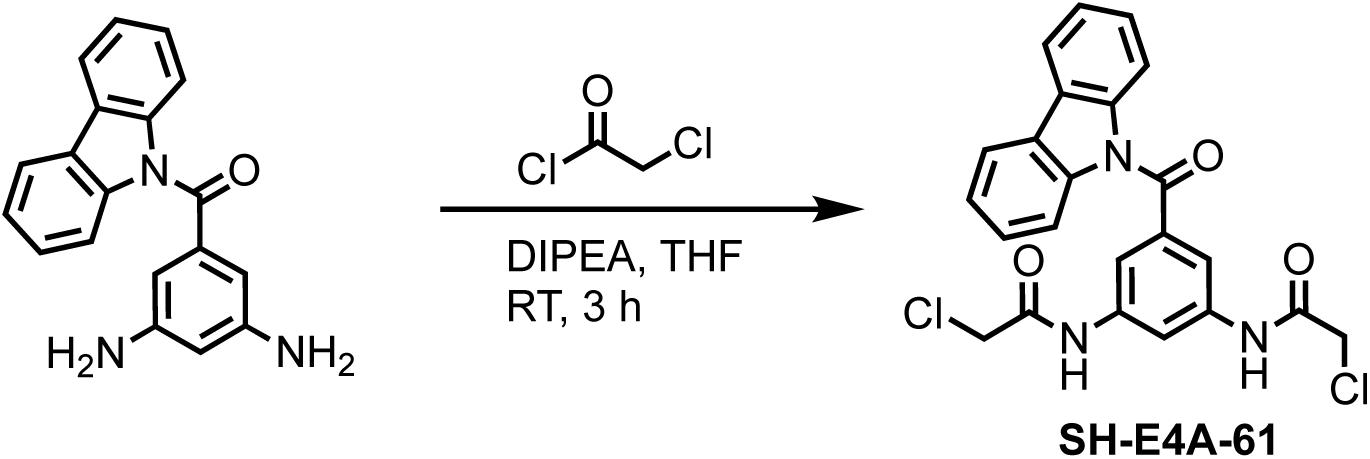

Chloroacetyl chloride (0.005mL, 0.060mmol) was added to a solution of (9*H*-carbazol-9-yl)(3,5-diaminophenyl)methanone (4.5mg, 0.015mmol) dissolved in THF (0.146mL) and diisopropylethylamine (0.01mL, 0.06mmol) and the reaction stirred overnight at room temperature. The solvent was removed under reduced pressure and the crude material purified by column chromatography (eluent: DCM/MeOH). Yield: 5.5mg (87%).

^1^H NMR (400 MHz, MeOD) δ 8.36 (dd, *J* = 2.0, 2.0 Hz, 1H), 8.10 – 8.00 (m, 2H), 7.66 (d, *J* = 2.0 Hz, 2H), 7.54 – 7.45 (m, 2H), 7.39 – 7.28 (m, 4H), 4.16 (s, 4H).

### Cell culture

BXPC3, HCC78, HCT116, HT1080, K562, NCCIT, NCI-H3122, OVTOKO, and UACC257 cell lines were maintained in RPMI-1640 media supplemented with 10% fetal bovine serum and 1% penicillin/ streptomycin (Thermo Fisher Scientific). MCF7, U251MG, and U2OS cell lines used DMEM media with 10% fetal bovine serum and 1% penicillin/ streptomycin (Thermo Fisher Scientific). The BaF3 cell line uses DMEM media supplemented with Mouse IL3 Recombinant-Protein (Thermo Fisher Scientific). For subculture, adherent cells were detached using 0.25% Trypsin-EDTA (Thermo Fisher Scientific) with a 5-minute incubation period at 37°C. All cell lines tested negative for mycoplasma prior to initiation of experiments. Please refer to Extended Data Table S2 for culture details.

### Cell Treatment, Lysis and Protein Complex Interactions

Cell treatment, lysis and protein-protein interactions were assessed as previously described^182^. For treatment of U2OS cells expressing the indicated cDNAs, cells were seeded in a 6-well plate and 1 µg of plasmid DNA was transfected with jetOPTIMUS transfection reagent (polyplus-transfection) overnight. 24 hrs post-transfection cells were treated with varying drug concentrations in 1% DMSO with time as described in the text. For all other cell treatments, please refer to text.

Unless otherwise described, cells were lysed as described in the CONNECT platform below. For cells transiently expressing genes of interested, cells were lysed in a buffer composed of 5 mM KCl, 2.5 mM MgCl_2_, 20 mM HEPES, pH 7.4, 0.5% TritonX-100, and 0.5x CST Buffer (Cell Signaling Technologies) and supplemented with protease inhibitor (Roche), phosphatase inhibitor (Roche), and a protease/phosphatase inhibitor cocktail (Cell Signaling Technologies). For Structure-activity relationship (SAR) screen of 75 compounds, H3122 cells were lysed in 0.1% CHAPS lysis buffer (150 mM KCl, 50 mM HEPES, 0.1% CHAPS w/v, pH 7.4), supplemented with protease inhibitor (Roche), and collected with method described in the CONNECT platform. Protein concentrations were measured using the BCA protein assay (Thermo Fisher Scientific). 50-80 µg of cell lysate was combined with Blue Loading Buffer Pack with DTT (Cell Signaling Technologies) and heated at 95°C for 5 minutes before loaded into a precast gel (GenScript Biotech).

For protein-protein interactions, HEK293 cells transiently expressed 2 µg of pRK5-HA-CPSF6, pRK5-CPSF7-FLAG, or pRK5-RTRAF (control protein) using jetOPTIMUS transfection reagent (polyplus-transfection) overnight. The cells were then lysed in Pierce IP lysis buffer (Thermo Fisher Scientific, 25 mM Tris, pH 7.4, 150 mM NaCl, 1 mM EDTA, 1% NP40, 5% glycerol) with the extraction method described above. Lysate concentrations were normalized before compound treatment as described above. After 1.5 hr incubation at 37°C, samples were either mixed with loading buffer, or HA-Tag immunoprecipitation was performed with Pierce Ha-Tag Magnetic IP/Co-IP Kit (Thermo Fisher Scientific) following the manufacture’s protocol. 15 µl of beads were used and IP lysis/wash buffer provided was used to wash the beads. Loading buffer was added to the immunoprecipitated beads to elute proteins which were subsequently denatured by boiling for 5 min. Proteins were resolved by SDS-PAGE and analyzed by immunoblotting.

### Immunoblotting

Samples were resolved on an SDS-PAGE (GenScript Biotech), and proteins were transferred to a PVDF membrane (Millipore Sigma) overnight at 45V in a 4°C cold room. The membranes were blocked in 5% milk in TBST for 1 hr at room temperature while shaking, followed by incubation in primary antibody, diluted in 5% BSA in TBST for 1 hr at room temperature or overnight in the 4°C room. Refer to the Extended Data Table S2 antibody information. Membranes were washed 3 times for 5 minutes with TBST, and they were incubated with secondary antibody, diluted in 5% milk in TBST for 1 hr at room temperature. The membranes were washed again 3 times for 5 minutes with TBST. Proteins were detected with Pierce™ ECL Western Blotting Substrate (Thermo Fisher Scientific), imaged and developed with autoradiography film. Band intensity was measured with ImageJ (v2.14.0). The percentage of dimerization of ALK and PSME1 was measured by dividing the dimer band intensities by the combined intensities of the dimer and monomer bands.

### cDNA cloning and mutagenesis

Single and double cysteine mutants were generated by site-directed mutagenesis were generated using Q5 Site-Directed Mutagenesis Kit (New England Biolabs). The all-cysteine mutant constructs were cloned by using customized synthesized gBlock (Integrated DNA Technologies) as PCR template. Restriction enzymes were from New England Biolabs. See Extended Data Table S2 for primer sequences. All constructs were verified by DNA sequencing.

### CONNECT Platform

#### Cysteine Focused Chemical Proteomics

Iso-TMT samples were prepared as previously described^83^. In brief, adherent or suspension cells were cultured until reaching ∼80% confluency. Cells were washed once with ice-cold PBS, snap-frozen in liquid nitrogen, and stored at −80°C until use (n=3 biological replicates). Frozen cell pellets were lysed in DPBS supplemented with Benzonase (Santacruz) and protease inhibitors (Roche) using a chilled bath sonicator (Q700,Qsonica) and centrifuged for 3 min at 300 x g. Proteins were quantified by BCA assay (Thermo Fisher Scientific) and a total of 50 μg protein extracts were used per compound treatment. Lysates were treated with vehicle (DMSO) or COUPLrs (500 µM) for 1 hr, followed by 1 mM DBIA treatment for 1 hr.

Following DBIA incubation, lysates were reduced with 5 mM 5-tris(2-carboxyethyl)phosphine hydrochloride (TCEP) (Sigma-Aldrich) for 2 min at room temperature, followed by alkylation using 20 mM chloroacetamide (Sigma-Aldrich) for 30 min in the dark at room temperature. Proteins were precipitated using SP3 magnetic beads. In brief, SP3 magnetic beads (Cytiva) were prewashed with LC-MS grade water (Sigma Aldrich), and 250 μg combined SP3 beads (1:1, hydrophobic:hydrophilic) and LC-MS grade ethanol (Sigma Aldrich)were added to each sample to reach a final concentration of 50% ethanol. SP3 incubation was performed for 30 min at room temperature, and beads were subsequently washed 3 times with 80% HPLC grade-ethanol (Sigma Aldrich) and then resuspended with 175 uL of Trypsin/Lys-C (1 μg, Thermo Fisher Scientific) in 200 mM EPPS (Sigma Aldrich) pH 8.4, 5 mM CaCl2. Proteins were digested overnight (16 h) at 37°C and digested peptides were enriched with streptavidin magnetic beads (Cytiva) for 1 hr at room temperature. Beads were subsequently washed three times with DPBS, twice with HPLC grade-water (Sigma Aldrich). Peptides were eluted with 50 % acetonitrile (Sigma Aldrich), 0.1% formic acid (Thermo Fisher Scientific), and dried using a Speedvac (Thermo Fisher Scientific). Cysteine-enriched peptides were reconstituted with 30 % acetonitrile, 70 % 200 mM EPPS pH 8.4 and labeled with 25 μg of TMT reagent (Thermo Fisher Scientific) per channel for 75 min at room temperature with rotation. Labeling was terminated by the addition of 5% hydroxylamine (Acros Organics) for 15 min followed by addition of 10% formic acid. Peptides were desalted with stage tips using the following procedure: peptides were reconstituted with 5% acetonitrile/0.1 % formic acid and loaded onto C18 Micro Spin columns (Nest Group) pre-equilibrated with LC-MS grade methanol (Fisher Chemical) and LC/MS-grade water containing 0.1% formic acid. C18 spin columns were washed 10 times with LC/MS grade water containing 0.1 % formic acid and subsequently eluted with 80% acetonitrile, 0.1% formic acid and dried using a Speedvac (Thermo Fisher Scientific).

#### Mobility shift proteomics

Samples were prepared as described^62^. In brief, cell pellets (n=3 biological replicates) were lysed with 50 mM HEPES, pH7.5, 0.1% of SDS, supplemented with *cOmplete* protease inhibitor (Roche) on ice. The reconstituted cell lysate was sonicated with 2 rounds of 20 pulses using a probe sonicator (Branson Digital Sonifier SFX 250), rotated in a cold room for 10 minutes, and spun down at 300 *g* for 5 minutes at 4°C in a tabletop centrifuge. Supernatants were transfer to a new tube to measure concentration with BCA protein assay following manufacturer protocol (Thermo Fisher Scientific). Protein concentrations were normalized and 300 µg of 100 µl of lysate were used to set up the *in vitro* treatment with the indicated molecular COUPLRs (500 µM, 1 hr 37°C, end-over-end rotation or as described in the text). Following treatment, samples were treated with 50 µl of 3x Blue Loading Buffer Pack with DTT (Cell Signaling Technologies). 40 µg of each sample with loading dye were boiled at 95°C in a heat block before loading into a precast 12-well 8% polyacrylamide gel (GenScript Biotech) with Color Prestained Protein Standard (New England Biolabs). Samples were run at 100V for 2 hrs and the gel was stained with GelCode Blue Stain Reagent (Thermo Fisher Scientific). For each lane, the gel was cut according to the protein standard into six pieces (>250, 250-180, 180-95, 95-72, 72-55, 55-43 kDa) and placed into wells of a 96-well plate.

Samples were digested as previously described^90^. In brief, gel pieces were washed 3 times with 75 μL of 1:1 100 mM NH₄HCO₃: acetonitrile for 15 minutes and dehydrated with 75 μL of 100% acetonitrile for 30-60 seconds. The gel pieces were then rehydrated with 75 μL of 10 mM DTT (Millipore Sigma), in 50 mM NH₄HCO₃ for 1 hr at 56°C. After rehydration, the gel pieces were alkylated with 75 μL of 55 mM chloroacetamide (Millipore Sigma), 50 mM NH₄HCO₃. The gel pieces were washed again with a 1:1 100 mM NH₄HCO₃: acetonitrile solution, and then dehydrated with 100% acetonitrile for 30-60 seconds. The gel pieces were then rehydrated in 60 μL of digestion buffer (50 mM NH₄HCO₃, 5 mM CaCl_2_, 5 ng/μL Trypsin) for 15 minutes on ice. The samples were then incubated in 70 μL of 50 mM NH₄HCO₃, 5 mM CaCl_2_ at 37°C overnight. To extract the peptides, the samples were centrifuged briefly, and the supernatant was transferred to a new 96 well plate for collection. 75 μL of 50% acetonitrile, 0.3% formic acid was added to the gel pieces for 15 minutes before being transferred to the collection plate. 75 μL of 80% acetonitrile, 0.3% formic acid was added to the gel pieces for 15 minutes before being transferred to the collection plate. The pooled samples were then frozen in the −80°C freezer before being dried in the speed vac. Peptides were then subjected to the desalting method described.

#### MS Data Acquisition and Processing

All the mass spectrometry samples were analyzed as described previously^83^ on an Orbitrap EclipseTM TribridTM Mass Spectrometer coupled with an Easy NanoLC-1200 system (Thermo Fisher Scientific). Peptides were separated on a 75-µm capillary column packed with 50 cm of C18 resin (2µm, 100Å; Thermo Fisher Scientific) using a 180 min gradient of 4-35% acetonitrile in 0.1% FA with a flow rate of 300nL/min. Eluted peptides were acquired by data-dependent acquisition and quantified using the synchronous precursor selection (DDA-SPS-MS3) method for TMT quantification. Briefly, MS1 spectra were acquired in the scan range of 400-1400 m/z at an orbitrap resolution of 120,000 with 50 ms as a maximum injection time with high-field asymmetric-waveform ion-mobility spectrometry (FAIMS) values at −40, −50, and −70 compensation voltage (CV). MS2 spectra were acquired by selection of the top twenty most abundant features via collisional induced dissociation in the ion trap using an automatic gain control (AGC) setting of 10 K, quadrupole isolation width of 0.7 *m*/*z,* and a maximum ion accumulation time of 50 ms. These spectra were passed in real time to the external computer for online database searching. Intelligent data acquisition using Comet real-time searching (RTS) was performed with a database including cell line mutations (DepMap)^183^ and human protein databases (release_20210506)^184, 185^. The same forward- and reversed-sequence human protein databases were used for both the RTS search and the final search (Uniprot)^186^. Next, peptides were filtered using simple, initial parameters that included the following: not a match to a reversed-sequence, containing TMTPro16 isobaric tags, maximum PPM error <50, minimum PPM error >5, minimum ΔCorr 0.10. If peptide spectra matched those above, an SPS–MS3 scan was performed using up to 20 *b*- and *y*-type fragment ions as precursors with an AGC of 250 K for a maximum of 250 ms, with a normalized HCD collision energy setting of 55 (TMTPro16)^185^.

Peptide search was performed in Proteome Discover (2.5, Thermo Fisher Scientific) against uniport human protein database (release_20210506) with the following parameters: up to two missed cleavages, 20 ppm precursor tolerance, 0.6 Da fragment ion tolerance and fully tryptic peptides were allowed with a minimum of 6 peptide length. Static modification of TMTPro16 on lysine and peptide N-termini (+304.2071 Da) and carbamidomethylation of cysteine residues (+57.0214 Da) whereas oxidation of methionine (+15.9949 Da) and DIBA on cysteine residues (for iso-TMT samples) (+239.1634), as variable modifications was allowed. Results were filtered to a peptide false discovery rate (FDR) of 1%. For the TMT reporter ion quantification, all the identified peptide spectral matches (PSMs) from the MS3 scans were extracted by an in-house program and the reporter ion intensities were adjusted for impurity correction according to the manufacturer’s specifications. For quantification of each MS3 spectrum, a total sum signal-to-noise of all reporter ions of 100 (TMT16-plex) was used.

### Live cell imaging, immunofluorescence staining, and analysis

#### NPM1 live cell imaging

The GFP-NPM1 cDNA was obtained from Addgene (#17578) and the NPM1c mutation was introduced and edited using Q5 Site-Directed Mutagenesis Kit (New England Biolabs). U2OS cells were transiently transfected with GPF-NPM1 or GFP-NPM1c overnight as described above, treated as described in the text and -stained with 5 µl/ml of Hoechst 33342 (R&D) in PBS at room temperature for 5 minutes. To assay how molecular COUPLrs altered NPM1c localization, live cell imaging was conducted on in Tokaihit stage incubator (37°C, 5% CO2, high humidity) equipped with a Yokogawa CSU-W1 spinning disc confocal attached to a Ti-2 Nikon inverted microscope. The plates were scanned every 15 minutes, with 6-7 field of interest, and for 6 hrs with 20x objective.

#### EML4-ALK Live cell imaging

U2OS-EML4-ALK(K589M)-mscarlet cells were seeded in 24-well glass-like bottom #1.5 cover glass plate (Cellvis) one day prior to treatment. Once attached, the cells were pre-stained with 5 µl/ml of Hoechst 33342 (R&D) in PBS at room temperature for 5 minutes then treated with the compounds as described in the text. 100 µg/µl of cycloheximide (Millipore Sigma) and 1 µM MG132 (Cell Signaling Technologies) were used to study ALK degradation. The imaging procedure was described in above. The plates were scanned every 15 minutes, with 6-7 field of interest, and for 24 hrs with 20x objective.

For cell segmentation and tracking, ND2 files for each treatment were processed using Fiji v2.11.0 ^187^ via PyImageJ v1.4.1^188^. Cells in each time series were segmented and tracked using the TrackMate-Cellpose plugin (v0.1.2)^189, 190^. Segmentation masks were produced using the “cyto” model, with mScarlet as the primary channel, DAPI as the secondary nuclear channel, a median diameter of 30 µm, and without simplifying cell segmentation mask contours. XML files containing cell segmentation mask and track information were saved for visual validation, and resulting CSV files of cell fluorescence intensity information for each time series were produced for further analysis in R v4.0.4. For data filtering and outlier removal, cell fluorescence data was filtered such that for each well, any cells that were not present in every frame in a time series were removed. Additionally, any cells with an mScarlet standard deviation that fell below the 0.02 quantile or above the 0.99 quantile for each well were removed as outliers. The number of cells remaining after filtering was recorded for each treatment. To correct for autofluorescence, on a frame-by-frame basis, the minimum cell mScarlet intensity in each frame was subtracted from each mean cell mScarlet intensity values. Adjusted mean intensities for each cell were then normalized to the cell’s value in the first frame. The average change in adjusted mean intensity and the 95% confidence interval was calculated for each frame for each well using plyr v1.8.8. Time out of 12 hrs was calculated using number of frames and was plotted against average change in adjusted mean intensity from time=0 with ggplot2 v3.4.2.

#### NPM1 immunofluorescence staining

For quantification of nuclear and cytosolic translocation of NPM1c, U2OS cells expressing FLAG-NPM1c were seeded in matrigel-coated 96-well glass-like bottom plate (Cellvis). The cells were then treated with SH-X-42 for 3 hrs, fixed with 2% PFA for 15 minutes at room temperature, The cells were first permeabilized with 50 µL PBST (0.05% Triton-X) for 10 minutes at room temperature. The PBST was then aspirated, and cells were rinsed twice for five minutes each time in 200 µL PBS. After rinsing, the cells were blocked with 50 µL of 3% BSA in PBS for 30 minutes at room temperature. The serum was then aspirated, and cells were incubated for one hr at room temperature with 43 µL of a 1:500 dilution FLAG antibody in PBS. After aspirating the primary antibody solution, cells were rinsed three times for five minutes in 200 µL PBS. Cells were then incubated for one hr at room temperature with Texas-red anti-mouse antibody (Vector Laboratories) in PBS. The secondary antibody solution was aspirated before rinsing cells three times for five minutes in 200 µL PBS. The cells were then stained with 50 µL of a 1:1000 dilution of DAPI (Thermo Fisher Scientific) in PBS. Plates were scanned with CSU-W1 SoRa (Nikon) and ND2 images were processed using QuPath v0.5.0^191^ and Cellpose v3.0.7^190^ via the qupath-extension cellpose v0.9.2 (https://github.com/BIOP/qupath-extension-cellpose). Cell cytoplasm and nuclei were segmented independently via the ‘cyto’ model. Cytoplasms were segmented using TRITC as the primary channel and DAPI as the secondary channel with the following parameters: pixelSize=0.5, normalizePercentilesGlobal=(0.2, 0.8, 1); tileSize=2048; simplify=0. Nuclei were segmented using only the DAPI channel with the following parameters: pixelSize=0.5; tileSize=2048; diameter=0.0; simplify=0. Full cell segmentation masks were calculated by taking pairwise intersection of all nuclei masks with all cytoplasm masks and combining the two with the largest intersection. Cell fluorescence summary statistics were recorded in one CSV per ND2 series. Cells with low NPM1 mean intensity (< 200) or with a DAPI nuclear mean / cytoplasmic mean ratio of < 2 were excluded, along with treatments with < 100 cells. The retained cells were used to calculate a median NPM1 nuclear mean / cytoplasmic mean ratio for each treatment. Additional experiment information, including plate number, time, treatment, and concentration, was added using R v4.3.3.

#### Purification EML4-C82 recombinant protein

Human EML4 (residues 7-86) in the pET28 vector (Novagen), containing N-terminal His-SUMO tag, was overexpressed in E. coli BL21 (DE3) and purified using affinity chromatography and size-exclusion chromatography. Briefly, cells were grown at 37°C in minimal medium supplemented with ^15^N-ammonium chloride (Cambridge Isotope Laboratories, Inc) in the presence of 50 μg/mL of kanamycin to an OD of 0.8, cooled to 17°C, induced with 400 μM isopropyl-1-thio-D-galactopyranoside (IPTG), incubated overnight for 20 hrs at 17°C, collected by centrifugation, and stored at −80°C. Cell pellets were lysed in buffer A (25 mM Tris, pH 8.0, 500 mM NaCl, 0.5 mM TCEP, and 20mM imidazole) using Microfluidizer (Microfluidics), and the resulting lysate was centrifuged at 30,000g for 40 min. Ni-NTA beads (Qiagen) were mixed with cleared lysate for 90 min and washed with buffer A. Beads were transferred to an FPLC-compatible column, and the bound protein was washed further with buffer A for 20 column volumes, followed by elution using 400 mM imidazole in buffer A for 10 column volumes. His-SUMO-tag was cleaved by adding ULP1 protease and incubating overnight in a cold room. The cleaved protein was concentrated and purified further using a Superdex 75 16/600 column (Cytiva) in buffer B (20 mM HEPES, pH 7.5, 200 mM NaCl, and 0.5 mM TCEP). The fractions containing cleaved EML4 protein were concentrated to ∼6 mg/mL and stored in −80°C.

#### NMR experiments and analysis

^15^N-labeled EML4 protein samples were incubated with 1:1 ratio of 20 µM SH-E4A-61 and SH-E4A-62 at 35°C for 16 hrs in 10 ml of 20 mM HEPES, 200 mM NaCl, 1 mM TCEP, pH-7.5 buffer. The samples were then concentrated to a final volume of 250 µl with 0.8 mM EML4 protein before adding 20 µl D_2_O and transferred to Shigemi NMR tubes. ^15^N-TROSY-HSQC data were collected on a Bruker Advance II 600 MHz spectrometer equipped with a Prodigy cryogenic probe at 25°C. The number of complex points acquired were 512 in the direct ^1^H dimension and 120 in the indirect ^15^N dimension, and the number of scans were 12. Data processing and analyzing were done using the Bruker Topspin software.

#### Thermal stability assays

Thermal stability assays were conducted as previously described^90^. In brief, to determine NRAS thermal stability, 1 µg of plasmid DNA were transfected to HEK-293T cells to transiently expressing each protein in the pRK5 vector were harvested and the pellets lysed with DPBS supplemented with protease inhibitor (Roche), Sodium Fluoride (Sigma Aldrich), and Sodium Orthovanadate (Sigma Aldrich), using a chilled bath sonicator (Q700, QSonica). The lysates were clarified by centrifugation at 300 *g* for 3 min. The supernatants were diluted to 1.25 mg/mL using DPBS and aliquoted at 50 µL/well in a PCR strip. The FLAG-NRAS lysates were treated with vehicle (DMSO), 100 µM SH-X-14 at room temperature for 1 hr. The samples were heated to the indicated temperature in a BioRad T100 Thermal Cycler (BioRad) for 3 min. Samples were then cool at room temperature for 3 min, incubated on ice for 3 min, transferred into 1.5 mL tubes, and centrifuged at 21,000 g for 1 hr. The soluble fractions were collected for immunoblot analysis.

For ALK thermal stability, H3122 cells were treated as described in the text for 6 hrs. Cells were lysed in 0.1% CHAPS lysis buffer supplemented with *cOmplete* protease inhibitor (Roche) and 100X CST protease/ phosphatase inhibitor (Cell Signaling Technologies). The samples were then sonicated with 20 pulses using a probe sonicator and spun down at 2,000 *g* at 4°C in a tabletop centrifuge for 5 minutes. The resulting clarified lysates were transferred to new 1.5-mL microcentrifuge tubes and protein concentrations were normalized to a concentration of 2 µg/µL. The samples were subsequently transferred to PCR strips, heated on a PCR block at their respective temperatures for exactly 3 minutes, then left at room temperature for 3 minutes, and finally placed on ice for 3 minutes. After briefly spinning down the PCR strips, samples were transferred to 1.5-mL microcentrifuge tubes and spun down at 14,000 rpm at 4°C in a tabletop centrifuge for 60 minutes. The soluble fractions were collected for immunoblot analysis.

The band intensity was assessed using ImageJ analysis and normalized to the value at the lowest temperature used for each experiment (NRAS: 42 °C, EML4-ALK: 41 °C). The melting curves of each protein were generated by Prism v10 (GraphPad) software.

#### Lentivirus production and generation of stable cell lines

Lentivirus generation was described previously^90^. In brief, U2OS or BaF3 cells were transduced with viral particles harboring the indicated genes to generate pCDH-EF1-EML4-ALK(K589M)-mScarlet (U2OS) or EML4-ALK-HA, EML4-ALK(G1202R)-HA, and EML4(cys-mut)-ALK-HA (BaF3). All cells were selected with puromycin (Thermo Fisher Scientific) after infection. The BaF3 EML4-ALK lines were cultured without 0.5 ng/ml IL3 after 3 weeks of transduction.

#### ALK *In vitro* kinase assay

Five million BaF3 cells expressing either EML4-ALK-HA or EML4-ALK G1202R-HA were treated with SH-E4A-66 or vehicle control for 3 hrs at 37°C as described in text. Cells were washed with chilled PBS and lysed in IP Lysis Buffer (25 mM Tris, pH 7.4, 150 mM NaCl, 1 mM EDTA, 1% NP40, 5% glycerol), supplemented with protease inhibitor (Roche), phosphatase inhibitor (Roche), and protease/phosphatase inhibitor cocktail (Cell Signaling Technologies). The cells were sonicated with 10 quick pulses and incubated for 15 minutes at 4°C with end-to-end rotation. The lysates were centrifuged at 13,300 rpm for 10 minutes at 4°C and the supernatant was transferred to a 1.5 mL tube. 200 µg of clarified lysate was incubated with 18 μL of anti-HA magnetic beads (Thermo Fisher Scientific) and incubated for 1 hr at room temperature with end- to-end rotation. The beads were washed 3 times with IP lysis buffer and 2 times with Milli-Q water.

The *in vitro* kinase assay was performed as method previously described^182^. In brief, magnetic beads harboring the indicated proteins were incubated in kinase buffer (10 mM HEPES, pH 7.5, 10 mM MnCl_2_, 1 mM DTT) with or without 200 µM ATP (New England Biolabs) for 30 minutes at room temperature with end-to-end rotation. Samples were washed once with kinase buffer followed by addition Blue Loading Buffer Pack with DTT (Cell Signaling Technologies) was added to each sample, which were subsequently boiled at 95°C for 5 minutes. Kinase activity was assessed by immunoblot analysis and band intensity corresponding to ALK phosphorylation activity (Tyr1278, Tyr1282, Tyr1283) was measured with ImageJ (v2.14.0). The IC_50_ was determined by quantifying p-ALK signals of BaF3 EML4-ALK cells treated with 0, 0.04, 0.08, 0.16, 0.31, 0.62, 1.25, 2.5, 5, and 10 µM of SH66. Prism 10 (GraphPad) non-linear fit, variable slope (four parameters) was used to calculate IC50.

#### Data Transformation

We calculated cysteine engagement as previously described^90^.

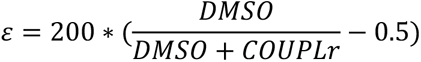

When possible, p-values were calculated with respect to one cysteine and molecular COUPLr at a time. Specifically, for a single molecular COUPLr, the engagement scores of a single cysteine were compared to a vector of zeros with length equal to the number of measurements made, i.e. the number of detections across n = 3 biological replicates. We tested the significance of cysteine engagement on the null hypothesis that a COUPLr, if completely inactive at a particular site, exhibits no competition with DBIA. For example, if an engagement score was measured for a particular cysteine three times, the significance of these three measurements would be tested against a vector of three zeroes (i.e. 0% engagement) via the MATLAB ttest2() function. A cysteine was considered as liganded by a molecular COUPLr if bound with ε ≥ 25% and p < 0.05.

Recalling that the mobility shift assay consists of serial mass spectrometric protein abundance measurements in both DMSO- and molecular COUPLr-treated conditions, where proteins are isolated from different molecular weight ranges prior to analysis by mass spectrometry, mobility shifting by a molecular COUPLr was quantified in reference to a defined molecular weight range or “band” at a time. On a per-band basis and with respect to one molecular COUPLr at a time, a protein can be assigned a “mobility shifting score” which is defined as the ratio of two protein abundances:

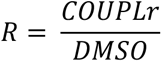

Considering a single molecular COUPLr, at most six mobility shifting scores can be assigned to a protein, where each score corresponds to a change in absolute protein abundance (i.e. MS^3^ reporter ion intensity) within one of six different molecular weight ranges which were subjected to proteomic profiling. To estimate confidence in a set of mobility shifting scores, p-values were calculated. The protein abundances which were measured for each condition (i.e. DMSO and SH-X-1C9) were first log-transformed individually (i.e. “within a column”) and then z-normalized to yield a set of relative protein abundances within a particular molecular weight range and with respect to different treatment conditions. These two sets of z-scores were then fed to the MATLAB ttest2() function to obtain a p-value estimating the significance of a given protein’s relative abundance change within a molecular weight range. A protein was considered to undergo a mobility shift by a COUPLr if its ratio R = COUPLr/DMSO ≥ 1.5 and p < 0.05 within a molecular weight distribution that is heavier than expected. A protein’s expected molecular weight is taken to be the weight, in kilo-Daltons, of its full-length monomer.

#### Circos Plot

To summarize the landscape of C^2^-coupling within the U2OS cell line as revealed by the CONNECT platform, we visualized cysteine engagement and proteomic mobility shifting data in a dual Circos plot, which summarizes information with increasing compactness as the radial component increases (i.e. reading the plot from “inside to outside”). Each of 33 molecular COUPLrs as well as two cross-linkers (C2 and C5) are arrayed angularly. The innermost disc depicts angularly compressed volcano plots mapping the cysteine engagement profiles of each compound. This is followed by an annulus of scatter plots depicting the molecular weight of a protein as a function of the confidence and degree of its mobility shifting by a given molecular COUPLr/cross-linker. The third layer of the circos plot summarizes protein domain targeting of the indicated molecule from the perspective of cysteine engagement. The outermost ring of the circos plot summarizes the frequency of protein coupling as understood among different protein classes. The categorization of cysteines, and their corresponding proteins, into different sets was previously reported^90^.

#### Visualizing the architecture of proteomic coupling across 12 cell lines

We sought to visualize the proteomic fingerprint of SH-X-1C9 induced mobility shifts across all 12 cell lines which were profiled with four molecular COUPLrs (SH-X-1C9, SH-X-34A, SH-X-48, SH-X-65). These C^2^-COUPLrs were chosen based on their varying reactivity as demonstrated in the initial screen of 33 COUPLrs in the U2OS cell line. Thus, for a single cell line at a time, we subsetted the mobility shifting scores on the subset of proteins which were detected in all six molecular weight ranges profiled. We then compressed this data to a two-dimensional manifold via the umap.UMAP function in the umap.umap_ Python package (n_neighbors=15, random_state=42, n_components=2).

To integrate these data with their companion cysteine engagement measurements, we first calculated the maximum engagement, minimum engagement, median engagement, and standard deviation for each cysteine across all cell lines, collapsing the cysteine engagement data for a given cell line into four intuitive features. These calculations were carried out with respect to the SH-X-65 probe, whose intermediate reactivity among the four COUPLrs allowed us to visualize a balanced representation of potential C^2^-coupling across the cysteinome. This compressed representation of the cysteine engagement data was then re-weighted by column-normalization and collapsed to a two-dimensional manifold via the umap.UMAP function in the umap.umap_ Python package (n_neighbors=15, random_state=42, n_components=2). This embedding was then projected onto a sphere as previously described^90^. Select examples of cysteine engagement and their corresponding molecular coupling are shown (Extended Data Fig. S5).

#### Integration of the CONNECT Platform with the Cancer Cell Line Encyclopedia

Given that 13 total cell lines profiled by the CONNECT platform herein have also been extensively subjected to multimodal molecular profiling in the Cancer Cell Line Encyclopedia^176^, we performed co-analyses of the mobility shifting data with publicly available records of proteinaceous somatic mutations (DepMap Public 24Q2;OmicsSomaticMutations.csv), RNA transcript expression levels (DepMap Public 24Q2;OmicsExpressionProteinCodingGenesTPMLogp1.csv), and genetic dependency estimates (DepMap Public 24Q2;CRISPRGeneEffect.csv).

#### Visualization of Protein Structures

Scenes containing representations of protein structures were generated in Chimera 1.17.3^192^.

#### Application of Cysteine Set Enrichment Analysis (CSEA)

When applied to an individual molecular COUPLr and evaluating the prevalence of molecular/biological features among coupled cysteines, cysteines which were liganded (ε ≥ 25% and p < 0.05) were searched for in a previously published CSEA library^90^, against a background set of cysteines comprising all other non-liganded and detected cysteines (Fig. 1g and Extended Data Fig. S3A-H). When searching for structural features which were frequent among highly-coupled cysteines, we focused on search on cysteines which demonstrated confident and strong binding (and ε ≥ 60% and p < 0.01 for Extended Data Fig. S4F) by at least 3 molecular COUPLrs, run against a background set of cysteines comprising all other cysteines which were detected but did not meet these stringent criteria.

#### Analysis of Protein Structures

In this study, we used the Euclidean distances between the geometric centroids of cysteines in three-dimensional space as an additional filter to understand the potential C^2^-coupling between two cysteine residues, whether within the same protein (i.e. intra-protein coupling) or between two different proteins (i.e. inter-protein coupling). Cysteine-cysteine distances were computed over a list of previously analyzed x-ray structures^90^, AlphaFold2^193^, and a set of structures containing the conformations of predicted^175^ binary protein-protein interactions (https://archive.bioinfo.se/huintaf2/).

#### Prediction of Multimeric Protein Structures

All multimeric protein structures, i.e. the NPM1 multimer (Figure 2), were predicted using ColabFold v1.5.5 with default settings as implemented in the AlphaFold2_mmseqs2 Jupyter notebook^194^.

#### Machine Learning based Classification of Protein Coupling

We sought an interpretable machine-learning based model which would in principle enable rational prediction of a particular protein’s amenability to molecular coupling. We first compiled a database of features which might influence a protein’s proclivity to interact with other proteins, namely: a protein’s molecular weight; the number of reported interactors a protein has in the BioGRID^180^ and IntACT^195^ databases; the number of turns, helices, strands, disordered regions, amino acid motifs, and domains a protein is predicted to have in the UniProt database^181^; the number of co-essential genes which are reported for a protein in the DepMap CRISPR screening database (Pearson ρ ≥ 0.4 across all cell lines); the number of labile cysteines a protein has (≥20% detection rate in a previous large scale cysteine-targeted covalent fragment screening study^90^), and the number of ligandable cysteines a protein has (≥20% detection rate and median quantile-normalized engagement ≥ 60% by ≥1 scout-probe e.g. KB05, KB03, KB02).

We then built a XGBoost (Python xgboost) tree-based classifier to evaluate whether a given protein is amenable to molecular coupling or not. The classifier was trained over observations of the previously described features among 5831 proteins which were detected at least once among the 12 cell lines profiled with four C^2^-COUPLrs via the CONNECT platform. We specifically applied the XGBoost architecture to predict a binary target class, where a protein which demonstrated at least once instance of significant mobility shifting (COUPLr/DMSO ratio ≥ 1.5 and p < 0.05 by SH-X-1C9, SH-X-34A, SH-X-48, or SH-X-64) in at least one of the six molecular weight distributions profiled was considered a 1, and all other proteins were treated as 0.

Hyperparameters were optimized using Bayesian optimization (Python bayes_opt;github.com/bayesian-optimization/BayesianOptimization) under a 70-30 train-test split regime. Class imbalance was compensated via SMOTE (Synthetic Minority Over-sampling Technique)^196^. Based on commonly investigated hyperparameter settings^197^, we explored a subset of the total possible hyperparameter space over 100 iterations. Specifically, we allowed the optimizer to converge within learning rate = 0.1-1, alpha = 0-100, lambda = 0-100, n_estimators = 100-250, and max_depth = 0-15. Optimization was governed by a 5-fold cross-validation score, using the mean AUC (Area Under the Curve) across the 5 folds as the evaluation metric. The optimal parameters were finally used to fit the model, and model statistics were computed across 1000 random train-test splits.

To find the features which drive model performance, we undertook a feature importance analysis^198^ on the test set. We first eliminated multicollinearity within the feature set, specifically to find a minimal basis set of qualitatively “independent” features. Specifically, we performed hierarchical clustering on the input data and determined empirical clusters of correlated features. From each cluster, a single representative feature was selected randomly, reducing redundancy and improving the interpretation of feature importance. I.e., the permutation of any one feature within this “basis” set will, by definition, more heavily influence model performance. To evaluate feature importance, the model was re-trained on the previously optimized parameters and the selected representative features, and permutation-based feature importance was evaluated via the Python scikit-learn package. We randomly permuted the values for one feature at a time and measured ΔROC-AUC to determine feature importance.

#### Tanimoto Similarity Coefficient, Similarity Maps Procedure

Using the cheminformatics package RDKit (version 2023.9.1), Tanimoto similarities were calculated between fragments. Structures inputted as SMILES strings were first converted into RDKit molecular objects, and RDKitFingerprints were generated for each molecule. Tanimoto coefficients were computed between fingerprints to assess molecular similarity. To generate SimilarityMaps between the linker and fragments, Morgan fingerprints (default radius 2 and 2048 bits) were generated for all fragments. Similarity maps were created using the default Dice similarity.

#### DFT Procedure

Density Functional Theory (DFT) calculations were performed using Gaussian 16. Geometric optimizations were performed in scf=water at b3lyp level of theory and 6-31+G(d,p) basis set. GaussView 6.1 was used to construct structures before optimization and visualize all output from the Gaussian 16 calculations. Please refer to **Extended Data Fig. 9** for DFT data.

---

**Extended Data Fig. 1:**
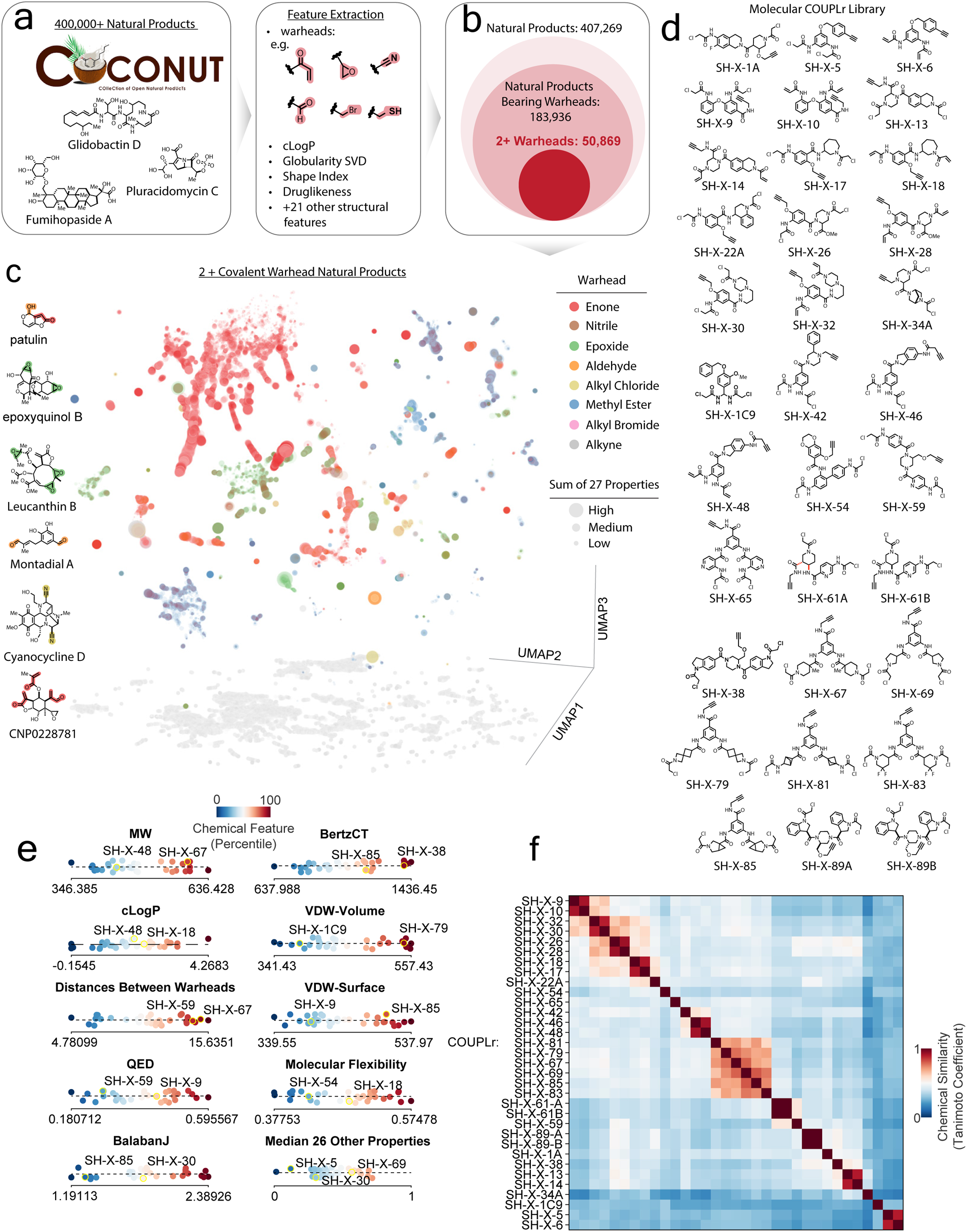
Dual covalent warheads are common in natural products. (a) Schematic for chemical feature exaction across a collection of 400,000+ natural products^179^. (b) Electrophilic warheads are common in natural products. Venn diagram enumerating natural products with one or more electrophilic warheads. (c) UMAP representation of natural products with two or more electrophilic groups. Representative examples of natural products from each cluster with cysteine reactive groups highlighted (see methods). (d) Structures of C^2^-COUPLr library. (e) Summary of chemical features of 33-member C^2^-COUPLr library. (f) Correlation heatmap of molecular fingerprints for 33 C^2^-COUPLrs.

**Extended Data Fig. 2:**
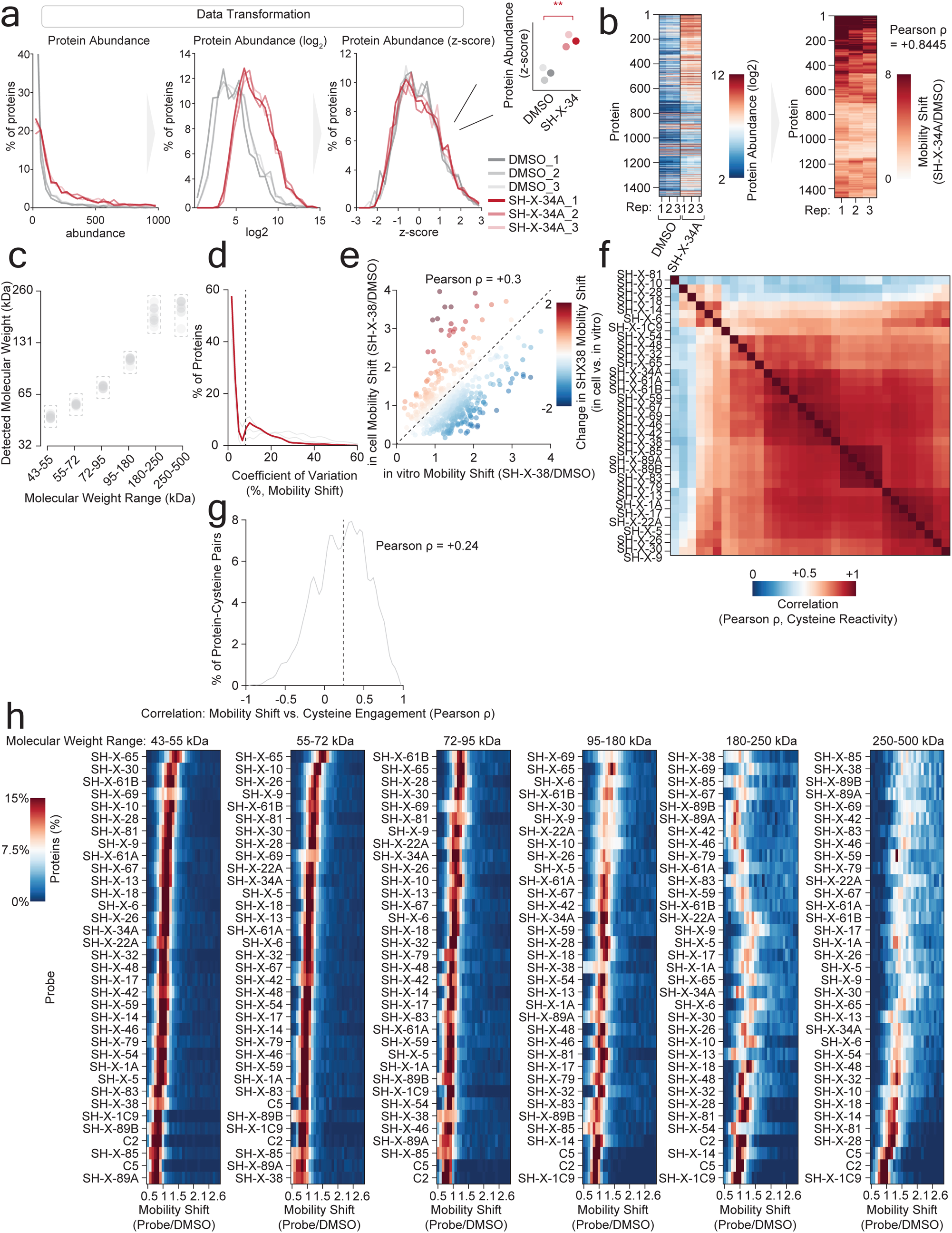
The CONNECT platform identifies protein targets of molecular COUPLrs. (a) Transformation of data output from the CONNECT platform (see also Methods). (b) Heatmap of protein abundances (left) and calculated mobility shift (right) following treatment with vehicle or SH-X-34A. Lysates isolated from U2OS cells were treated with vehicle or SH-X-34A and then run on a denaturing SDS-PAGE which was segmented into individual bands of defined molecular weight. The protein contents of each band were determined by mass spectrometry and a mobility shift is determined by movement of a protein from a gel band of its predicted molecular weight to a higher molecular weight (see Methods). (c) Average molecular weight of proteins in each band from cell lysates treated with DMSO and quantified by mobility shift proteomics. (d) Coefficient of variation across replicates of COUPLr treatment. (e) Comparison of in cell and lysate mobility shift following treatment with SH-X-38. (f) Heatmap depicts correlations of cysteine engagement profiles among probes in COUPLr library. (g) Positive correlation between the outputs from the two chemical proteomic platforms that comprise CONNECT. Distribution of correlations between cysteine engagement and mobility shift measurements across all COUPLrs. (h) Heatmaps display distribution of mobility shifts for each member of the 33 C^2^-COUPLr library for a given band. *p< 0.05. Statistical significance determined by MATLAB ttest2() function.

**Extended Data Fig. 3:**
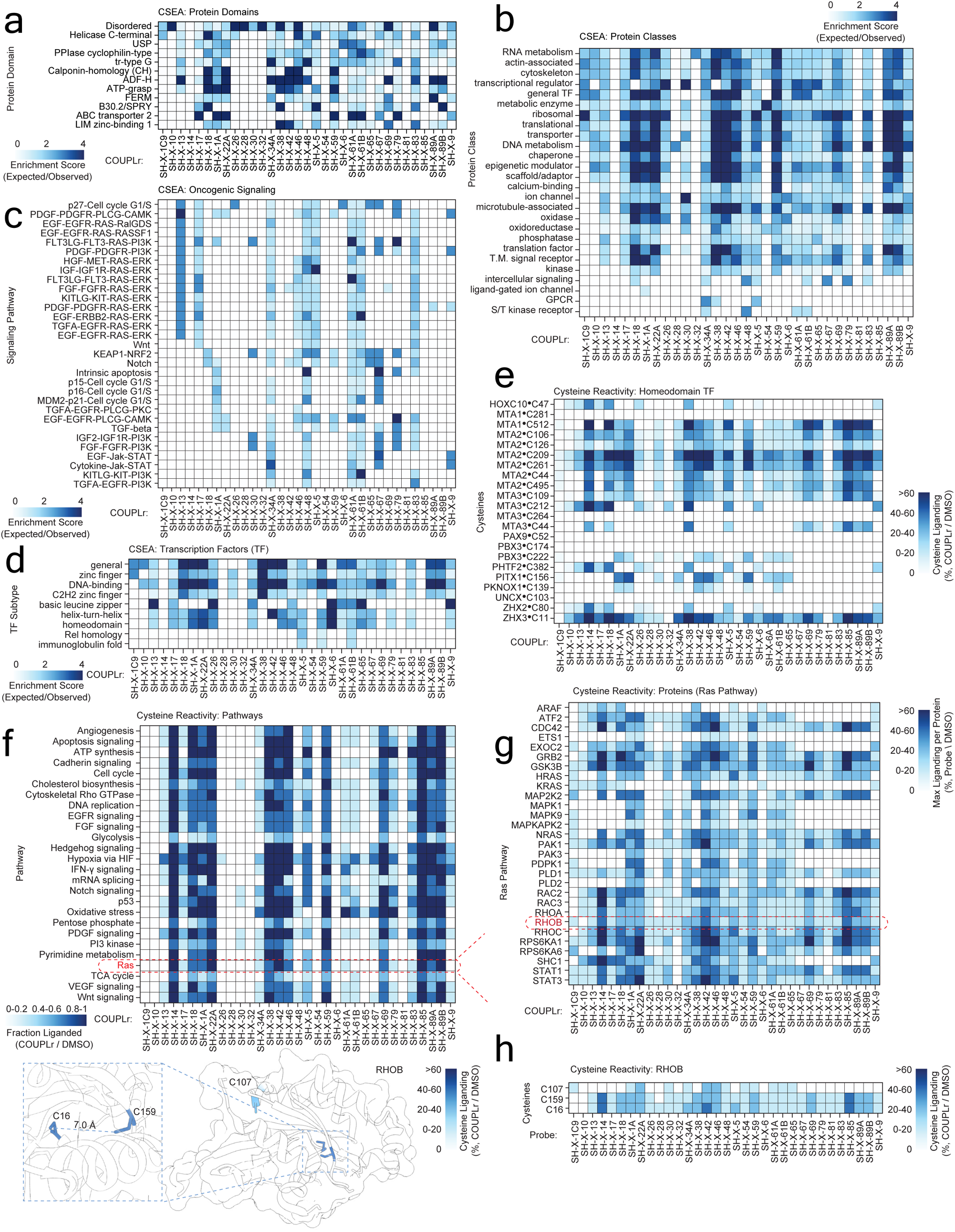
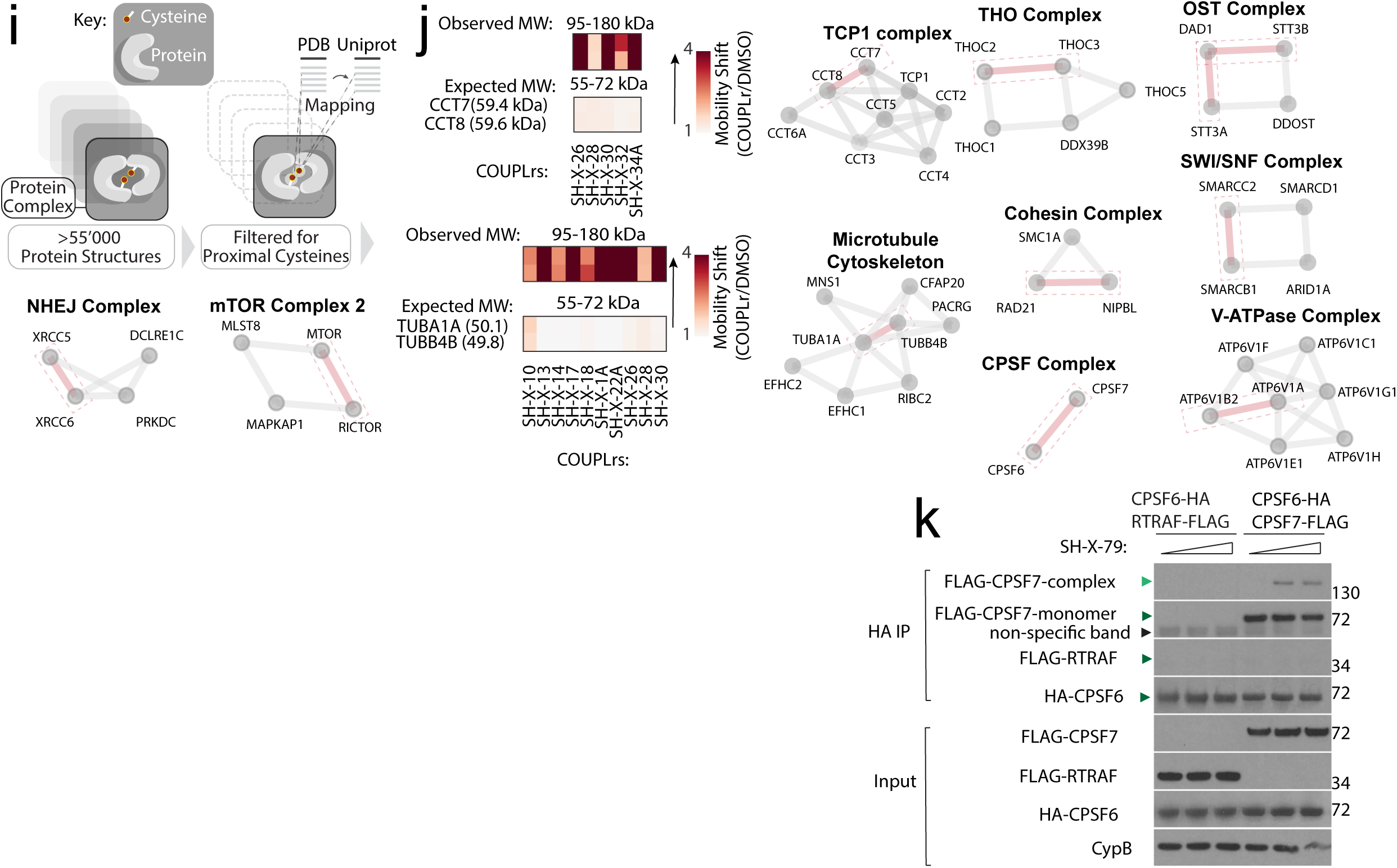
Identification of coupled targets in protein domains, classes and pathways. (a-c) Heatmaps for CSEA enrichment displaying protein domains (a), classes (b) and (c) oncogenic pathways for each C^2^-COUPLr profiled in U2OS cells. (d-e) Identification of transcription factor classes (d) that are targets of COUPLrs, highlighting cysteines engaged in homedomain containing transcription factors (e). (f-h) Pathway/protein/cysteine-level analysis of COUPLr targets in U2OS cells. Heatmap of cysteine engagement in biological pathways, with a focus on RAS signaling (f). Heatmap of engaged proteins involved in the RAS signaling pathway (g). Cysteines engaged within RhoB (PDB=2FV8) (h). (i) Schematic depiction of complexes that are potentially coupled. (j) Map of potentially coupled protein complexes. Mobility shift following molecular COUPLr treatment in U2OS cells was overlaid onto crystallized or predicted protein-protein interactions, prioritizing interfaces with proximal cysteines on respective interactors (see Methods)^175^. (k) CPSF6 and CPSF7 interact. Lysates isolated from HEK-293T cells transiently expressing HA or FLAG-tagged proteins were treated with SH-X-79 (0, 50 and 100 µM, 1.5 hr) and protein-protein interactions were monitored by immunoblot following enrichment with anti-FLAG M2-affinity resin. Here, RTRAF-FLAG functions as a negative control as it does not interact with CPSF6 and CPSF7.

**Extended Data Fig. 4:**
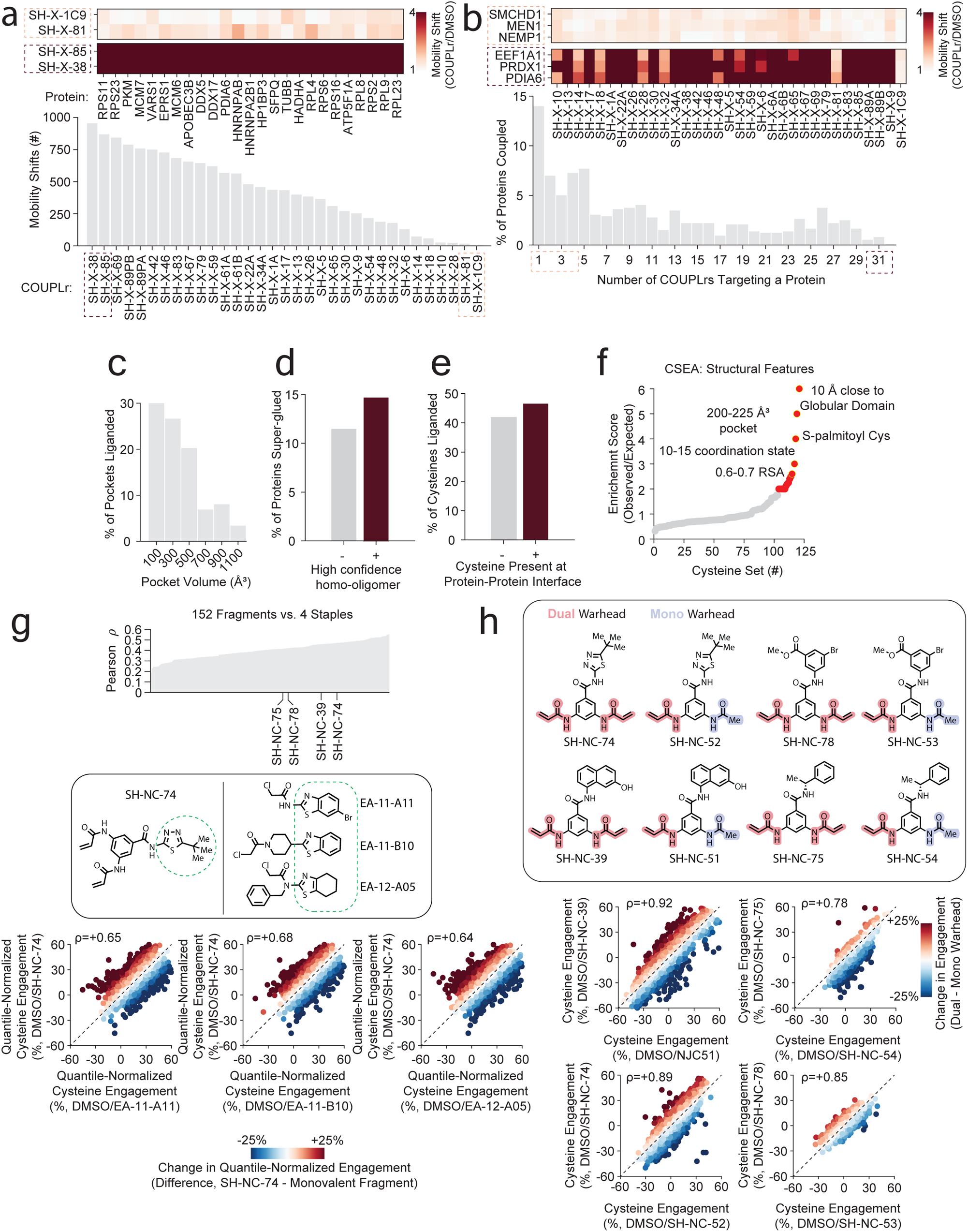
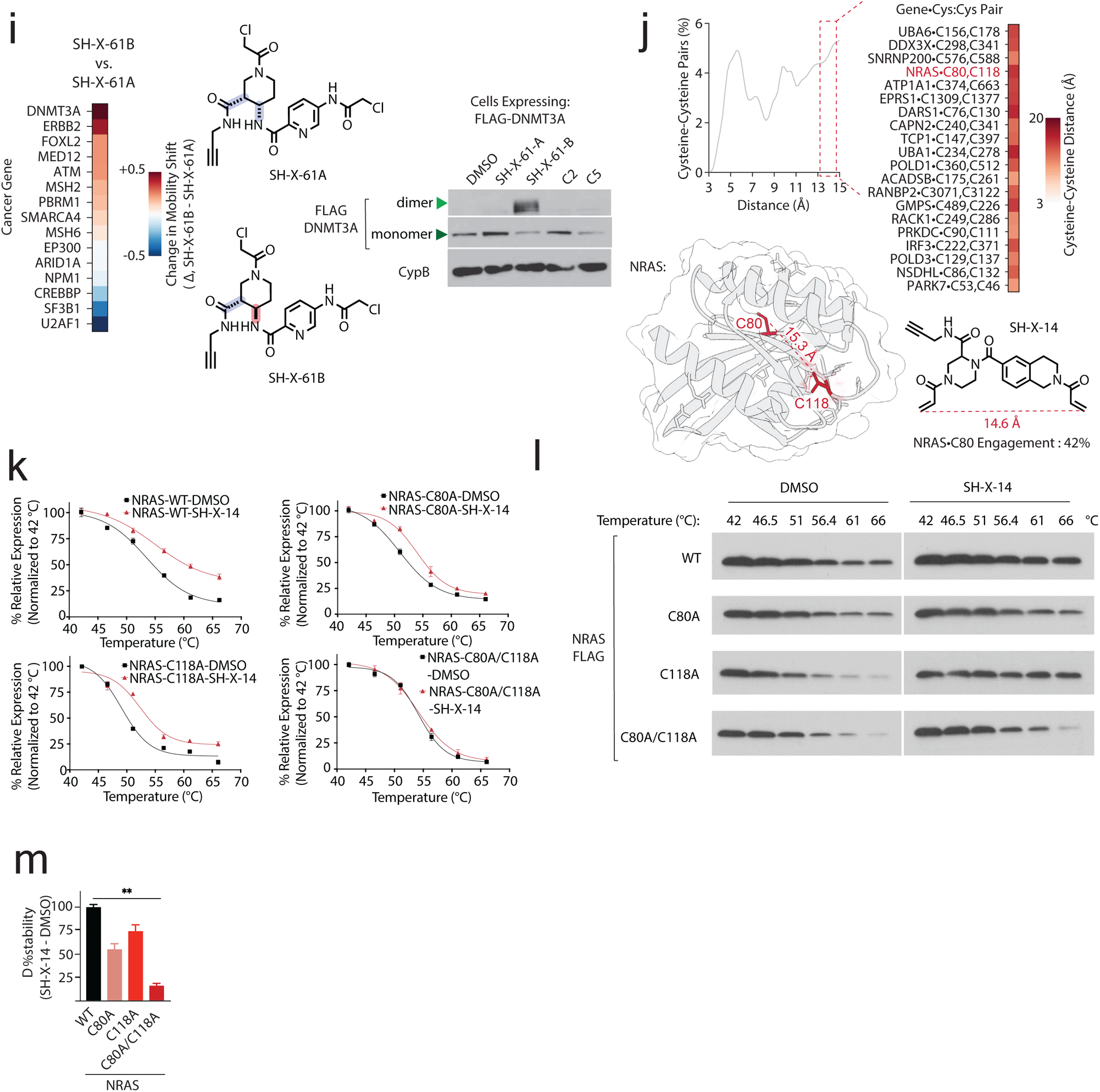
Molecular COUPLrs can induce intra-protein coupling and engage targets in a stereospecific manner. (a) Distribution of protein targets for each C^2^-COUPLr. Insets, proteins coupled by select compounds. (b) Distribution of coupled proteins across C^2^-COUPLR library identifies proteins which are common targets (inset). (c) Pocket volume distribution targeted by COUPLrs (see Methods). (d-e) Proteins annotated^180,181^ as homo-oligomers^102^ (d), and cysteines located at protein-protein interfaces (e) are targeted by COUPLrs (see Methods). (f) Cysteine set enrichment analysis (CSEA) for structural features that are frequent among COUPLr targeted cysteines. (g) COUPLrs show correlated reactivity with mono-warhead fragments. Plot summarizing cysteine-cysteine reactivity correlations between COUPLrs profiled in K562 cells and 152 electrophilic fragments using cysteine-focused chemical proteomics^90^. Inset displays 3 mono-warhead compounds which share cysteine reactivity profiles which are similar with SH-X-74 (top) and corresponding cysteine-engagement plots (bottom). (h) Mono- and dual-warhead compounds share similar cysteine reactivity profiles. Top, structures of paired mono and dual warhead compounds. Bottom, cysteine engagement plots for paired compounds. Lysates from K562 cells were treated with each compound or vehicle control, and changes in cysteine reactivity were determined by cysteine-focused chemical proteomics (see methods). (i) COUPLrs can target proteins in a stereoisomer-specific manner. Left, heatmap comparing coupling between the SH-X-61A and SH-X-61B stereoisomer pair. Right, DNMT3A coupling is stereospecific. U2OS cell was treated with the indicated compounds (20 µM, 6 hrs), and DNMT3A coupling was determined by immunoblot. (j) Comparison of distances (Å) between cys-cys pairs based on the analysis of PDB and AlphaFold-predicted structures (see Methods). Inset, left, proteins containing cys-cys pairs within the indicated range. Bottom, structure of NRAS highlighting Cys80 and Cys118 (PDB=3CON). (k-m) Intra-protein coupling within NRAS. (k) Quantification of thermal stability for NRAS or selected cysteine mutants following treatment with vehicle (black) or SH-X-14 (red). (l) immunoblot analysis of NRAS levels. Lysates from HEK-293T cells expressing the indicated proteins were treated with vehicle control or 100 µM SH-X-14, heated to the indicated temperatures and the soluble fraction of NRAS following centrifugation was quantified by immunoblot. (m) Quantification of change in thermal stability between SH-X-14 and DMSO for each of the indicated NRAS variants. **p< 0.001. Statistical significance determined by Student’s t-test (two-tailed, unpaired).

**Extended Data Fig. 5:**
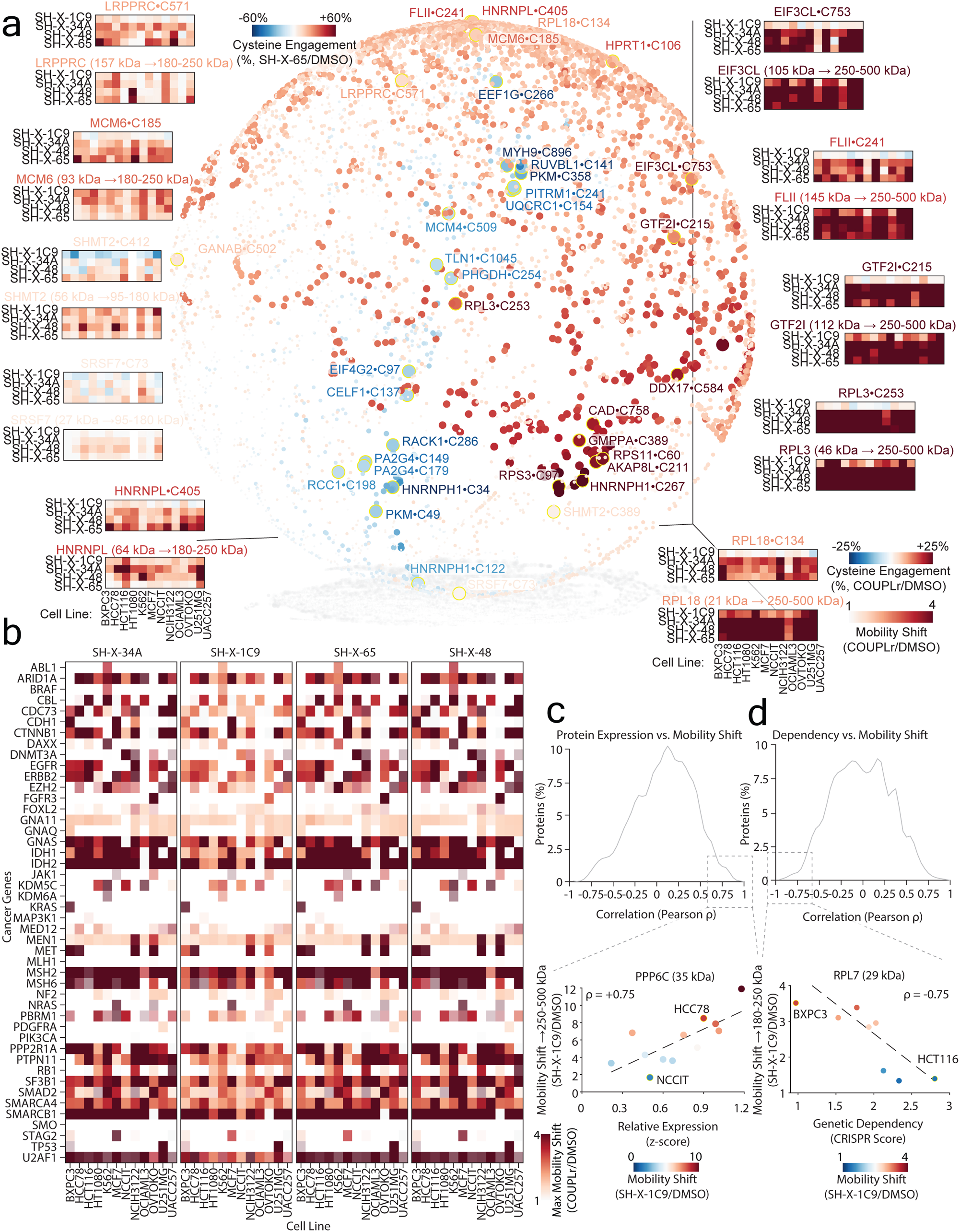
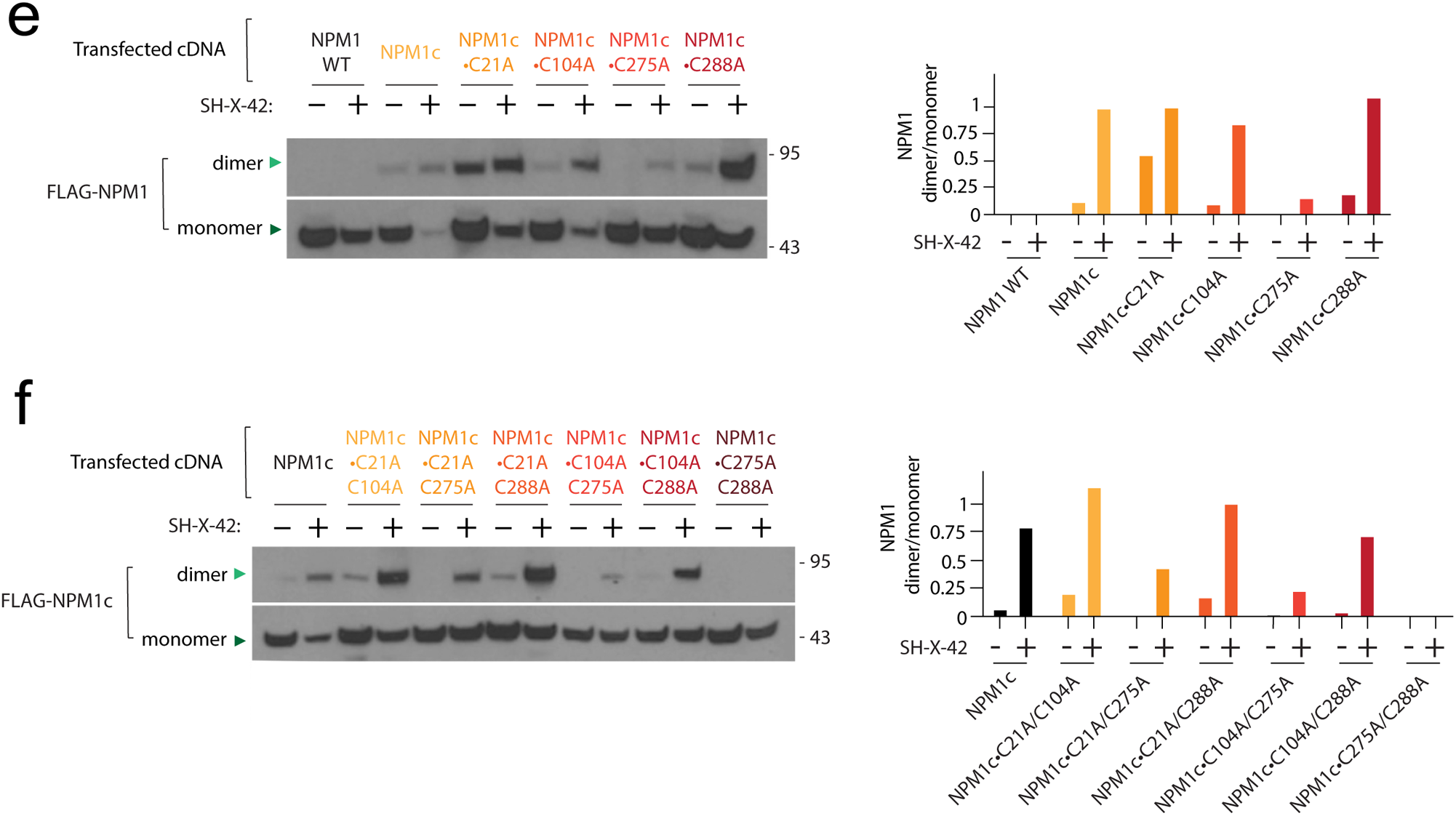
Molecular COUPLr profiling across 12 cancer cell lines. (a) Spherical representation of coupled cysteines across 12 cell lines when treated with SH-X-65. Inset heatmaps, highlight cysteine engagement (top) and mobility shift (bottom) for four *informer* COUPLrs across the different cell lines profiled (see methods). (b) Common cancer related proteins are targets of COUPLrs. Heatmap depicting coupling of the indicated gene products by each scout COUPLr across 12 cell lines. (c-d) Pearson correlation histogram depicting the relationship between coupling and protein expression levels (c) or genetic dependency (d) across 12 cell lines. (e) Cys275 within NPM1c is primarily responsible for coupling by SH-X-42. Left, lysates isolated from HEK-293T cells expressing the indicated NPM1c cysteine mutants were treated with 20 µM SH-X-42 for 3 hrs or vehicle and coupling was determined by immunoblot. Right, quantification of coupling. (f) Compound cysteine mutations within NPM1c disrupt coupling. Left, mutation of additional cysteines within NPM1c was evaluated in the presence of the NPM1c•C275A variant as described in (e). Right, quantification of coupling.

**Extended Data Fig. 6:**
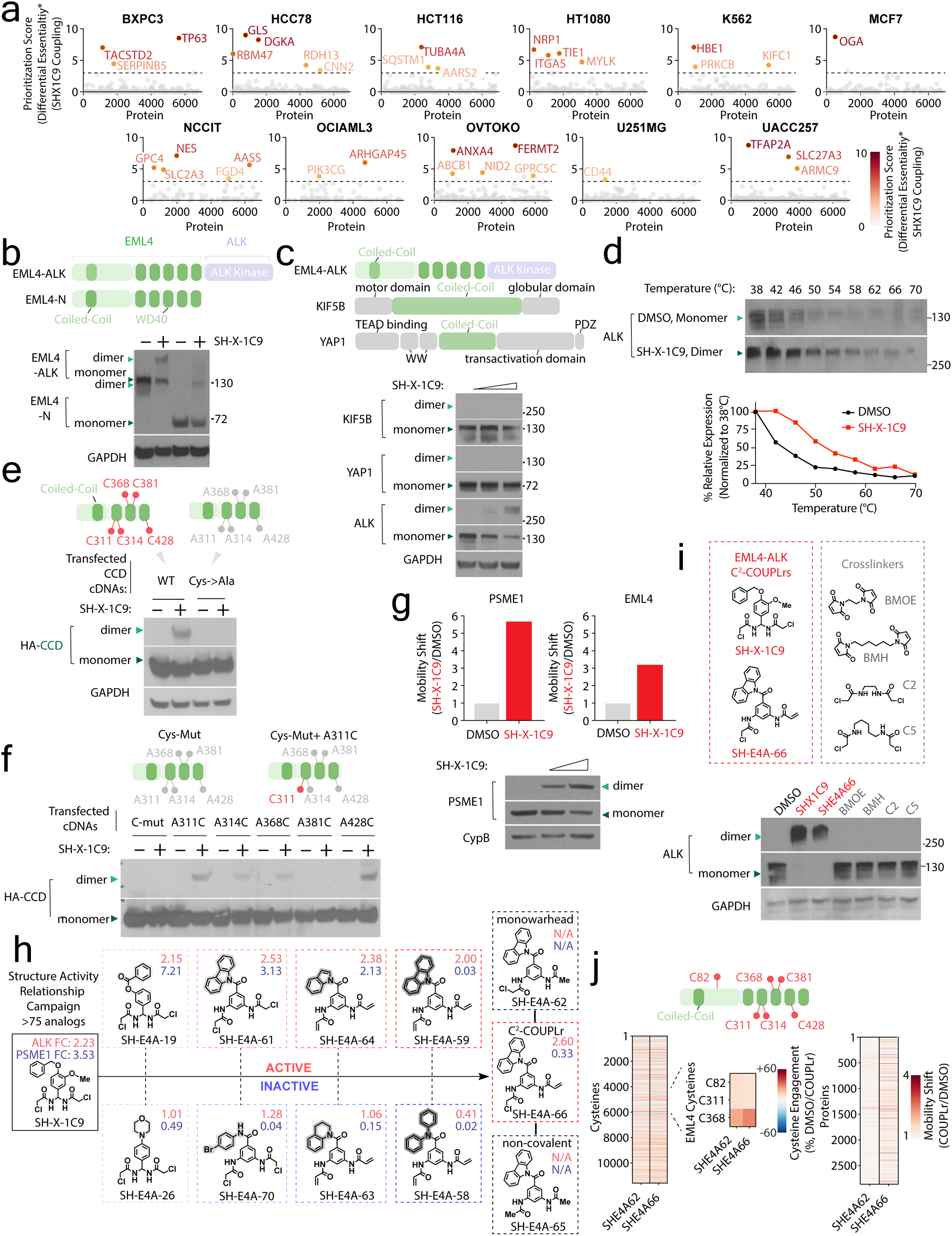
Development of an EML4-ALK COUPLr. (a) Identification of coupled dependencies across lineages. Plots depicting prioritization score for proteins identified in each cell line. A prioritization score for each protein was determined based on coupling by SH-X-1C9 and its corresponding gene essentiality. (b) EML4 within EML4-ALK is sufficient to mediate coupling by SH-X-1C9. Lysates isolated from U2OS cells expressing full length EML4-ALK or the EML4 portion of the fusion (EML4-N) were treated with 10 µM SH-X-1C9 for 3 hrs and coupling was assessed by immunoblot. (c) Targeting specificity of SH-X-1C9 for coil-coil domain (CCD) containing proteins. H3122 cells were treated with 0, 5 and 10 µM of SH-X-1C9 for 3 hrs, and coupling was assessed for the indicated CCD containing proteins by immunoblot. (d) SH-X-1C9 coupling of EML4-ALK results in thermal stabilization. Top, H3122 cells expressing EML4-ALK treated with DMSO or 10 µM of SH-X-1C9 for 6 hr followed by incubation at the indicated temperatures for 3 min. Following centrifugation, the relative levels of EML4-ALK were assessed by immunoblot. Bottom, quantification of thermal stability of EML4-ALK following treatment with SH-X-1C9 or vehicle control DMSO. (e) EML4-ALK coupling is cysteine-dependent. U2OS cells expressing either wild-type or Cys–>Ala mutant EML4 were treated with 10 µM SH-X-1C9 for 3 hr and coupling was determined as described in (b). (f) Role of individual cysteines within EML4-ALK CCD that mediate coupling. U2OS cells expressing the indicated EML4-ALK CCD variants were treated with SH-X-1C9 (10 µM) for 3 hrs, and coupling was evaluated by immunoblot. (g) PSME1 is a target of SH-X-1C9. Top, plots depicting PSME1 and EML4-ALK coupling by SH-X-1C9 in H3122 cells (see also Extended Table S1). Bottom, immunoblot analysis of PSME1 coupling in H3122 cells treated with SH-X-1C9 (0, 5 and 10 µM) for 3 hrs. (h) Development of an advanced EML4-ALK COUPLr. Structures and activities of SH-X-1C9 and related derivatives, highlighting chemical features that promote or negate coupling. EML4-ALK and PSME1 fold change (FC) coupling was assessed in H3122 cell lysate after treatment with the indicated compounds (5 µM for 3 hrs, see also Fig. 3d). (i) EML4-ALK coupling requires advanced binding elements. H3122 cell lysate was treated with 10 µM for 3 hrs of EML4-ALK COUPLrs (SH-X-1C9 or SH-E4A-66) or crosslinkers (C2, C5, BMOE, BMH) and coupling was assessed by immunoblot. (j) Low proteome-wide reactivity for advanced EML4-ALK COUPLrs. Heatmap depicting cysteine engagement (left) and mobility shift (right) following treatment of H3122 cells with 20 µM of the indicated compounds for 3 hrs.

**Extended Data Fig. 7:**
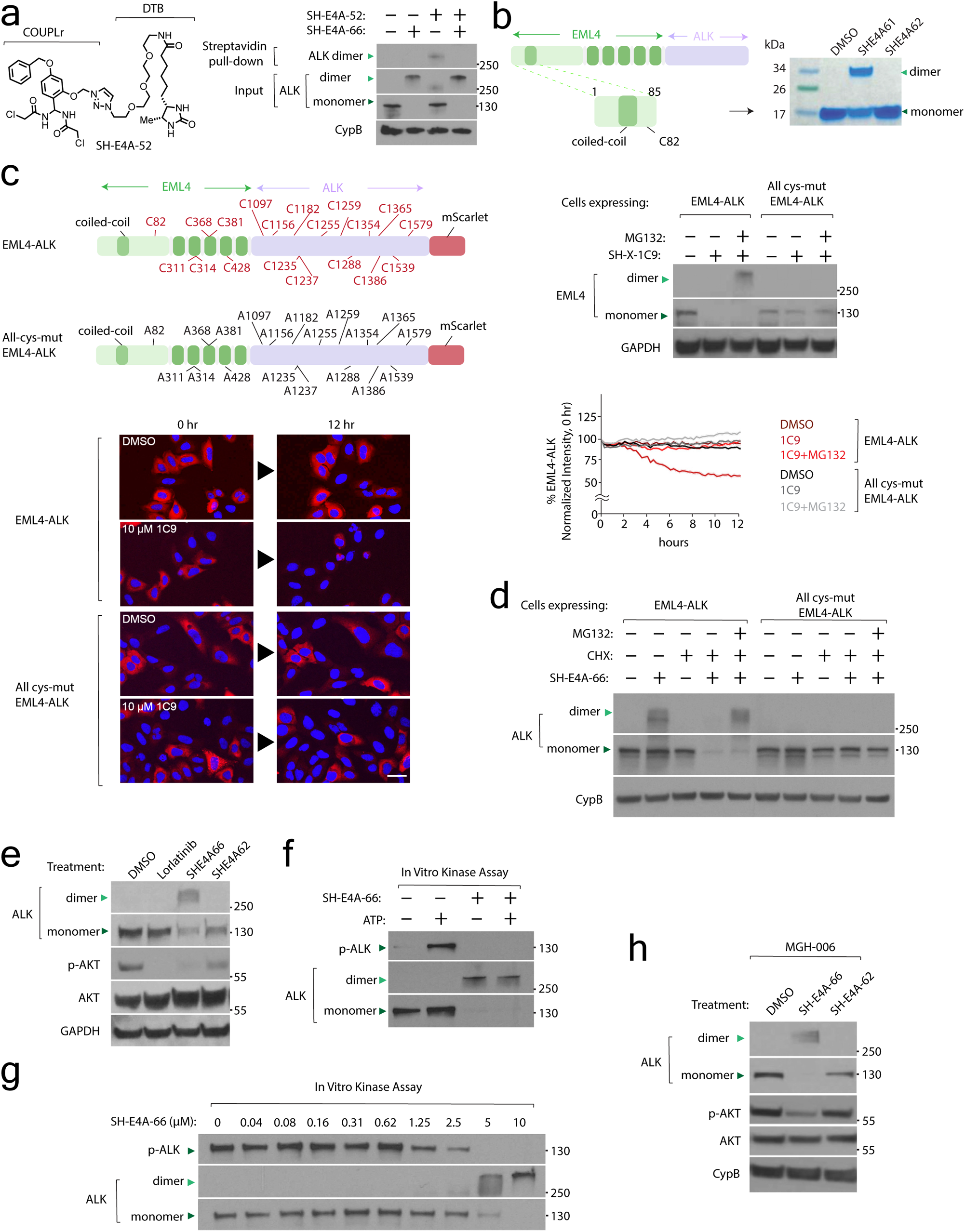
EML4-ALK coupling blocks signaling and leads to degradation. (a) SH-E4A-66 ligands EML4-ALK. Left, structure of a biotinylated analog of SH-X-1C9 (SH-X-E4A-52). Right, lysate isolate from H3122 cells was treated with vehicle or SH-E4A-66 (50 µM, 2.5 hrs) followed by a chase with SH-X-E4A-52 (100 µM, 2.5 hrs). EML4-ALK levels were determined by immunoblot following streptavidin enrichment. (b) Recombinant EML4-ALK CCD domain is sufficient for coupling by SH-E4A-61. 5 µg of recombinant protein (see Methods) was incubated with the indicated compounds and coupling was visualized on SDS-PAGE with Coomassie staining. (c) SH-X-1C9 degradation of EML4-ALK is cysteine and proteosome dependent. Top, schematic of EML4-ALK variants expressed in U2OS cells. Following cellular treatment with SH-X-1C9 (10 µM) and cycloheximide (100 µg/µl), the latter to block new translation, cells were subsequently treated with proteosome inhibitor MG132 (1 µM) and the relative expression of EML4-ALK was determined by immunoblot. Bottom, representative immunofluorescence images and quantification of WT and cysteine-mutant EML4-ALK following treatment with SH-X-1C9 (10 µM for 12 hrs). (d) SH-E4A-66 induces proteasome-dependent degradation of EML4-ALK in a cysteine-dependent manner. U2OS cells expressing the indicated proteins were treated as with 5 µM SH-E4A-66 and 100 µg/µl cycloheximide for 18 hrs followed by 1 µM MG132 and EML4-ALK levels were determined by immunoblot. (e) SH-E4A-66 but not SH-E4A-62 disrupts EML4-ALK signaling. BaF3 cell lines expressing EML4-ALK were treated with the indicated compounds (3 µM of SH-E4A-62 and SH-E4A-66, 20 nM of Lorlatinib) for 6 hrs, and EML4-ALK signaling was determined by immunoblotting for the indicated proteins. (f) Development of an in vitro kinase assay for EML4-ALK. Immunopurified HA-EML4-ALK from BaF3 cells treated with vehicle control or 5 µM SH-E4A-66 was incubated with 10 mM ATP and kinase activity was assessed by immunoblotting for phosphorylated ALK (p-Tyr1278/1282/1283) (see Methods). (g) SH-E4A-66 disrupts EML4-ALK kinase activity in a dose-dependent manner. EML4-ALK activity was measured as described in (f). (h) EML4-ALK coupling disrupts signaling in patient-derived lung cancer cell line. The MGH006 cell line was treated with vehicle, SH-E4A-62, or SH-E4A-66 (concentration 20 µM, 2 hrs), and signaling was determined as described in (e). Scale bar= 50 µm.

